# MarkerMatch: A Proximity-Based Probe-Matching Algorithm for Joint Analysis of Copy-Number Variants from Different Genotyping Arrays

**DOI:** 10.1101/2025.06.30.662249

**Authors:** Franjo Ivankovic, Dongmei Yu, James Shen, Lingyu Zhan, Maria Niarchou, Ariadne Kaylor, Laura Domènech, Tyne W Miller-Fleming, Luz M Porras, Paola Giusti-Rodríguez, Roel A Ophoff, Jeremiah M Scharf, Carol A Mathews

## Abstract

**Motivation:** Copy-number variants (CNVs) are a form of genetic structural variation with increasing importance in complex human disorders. Both DNA sequencing and microarray data can be used to call CNVs, which can be used in association tests, such as association between CNV number and disease status. Unlike genotypes, CNV detection in microarrays requires the use of observed intensity signals at each probe, which limits the imputability for analyses that span multiple array types. Thus far, a consensus set of probes (the intersection encompassing the probes that occur in common on all arrays) has been used to circumvent the problem of differing array-specific sensitivities. This has, however, led to excessive reduction in overall sensitivity of CNV calls as arrays can have an undesirably low overlap of probe sets. To overcome this limitation, we developed MarkerMatch, a proximity-based algorithm that matches probes across different genotyping microarrays to maximize the number of probes considered in the CNV calling algorithm, thereby increasing the resolution and sensitivity while preserving precision.

**Results:** By analyzing CNV calls from 4,906 individuals genotyped across three different arrays (Global Screening Array, Omni2.5 array, and Omni Express Exome array), we show that the MarkerMatch approach improves sensitivity by increasing the density of probes available for CNV calling while maintaining precision or improving it relative to the current practice (e.g., use of consensus probes only). We further demonstrate that MarkerMatch exceeds the output from current practice in terms of F1 score, Fowlkes-Mallows index, and Jaccard index. We also optimize MarkerMatch parameters, *D_MAX_* and *Method*, and find an optimal *D_MAX_* setting at 10kb, with no clear optimal candidate based on *Method*, indicating that parameters for this metric should be determined on a use case basis.

## Introduction

Copy-number variants (CNVs) are a form of structural variant involving unbalanced rearrangements leading to increased (duplication) or decreased (deletion) DNA content (Zarrei *et al*. 2015). CNVs have been studied in the context of various complex human disorders to better understand their underlying pathobiology (Lionel *et al*. 2011; Olson *et al*. 2014; Huang *et al*. 2017; Marshall *et al*. 2017: 17; Wang *et al*. 2018; Nakatochi, Kushima and Ozaki 2021; Fu *et al*. 2022). Microarrays are one of the most commonly used technologies to assay CNVs due to their relatively low cost and widespread use in genome-wide association studies and biobanks such as All of Us and the UK Biobank. (Bycroft *et al*. 2018; The All of Us Research Program Genomics Investigators *et al*. 2024).

While microarrays were not designed for the specific purpose of assaying CNVs, several algorithms have been developed to accurately assess CNV events using microarray data. Some software, e.g., PennCNV, QuantiSNP, Birdseye, and GenoCN, employ Bayesian approaches, such as Hidden Markov Models (HMM), to call CNVs using the intensity data from genotyping microarrays (Colella *et al*. 2007; Wang *et al*. 2007; McCarroll *et al*. 2008; Sun *et al*. 2009). Other approaches like cnvPartition or iPattern employ recursive partitioning and/or clustering to determine copy-number states (Pinto *et al*. 2011; Illumina 2017). More recent methods have focused on combining or building on existing approaches to fine-tune CNV calling performance (Zhang *et al*. 2019; Lavrichenko *et al*. 2021).

Genotyping microarrays vary in the density and selection of probes, with some arrays designed to capture variation specific to populations, diseases, genomic regions, etc (Ehli *et al*. 2017). This variability presents challenges to meta-analysis efforts, which are crucial for aggregating sufficiently powered datasets to detect genetic associations in complex traits (Zeggini and Ioannidis 2009).

By leveraging the effects of linkage disequilibrium, accurate imputation of non-genotyped markers is possible to enable joint and meta-analysis across different genotyping arrays (Scheet and Stephens 2006; Marchini and Howie 2010). However, CNV detection algorithms rely on direct probe intensity measures that may vary across arrays, and which cannot be imputed for CNV analysis.

Probe density and distribution vary across different array products, which may lead to array-type biases in CNV call sensitivity and specificity (Wang *et al*. 2007). Traditionally, to avoid such biases, researchers have taken manifest intersections and focused only on markers that were genotyped on all SNP arrays for CNV detection and analyses (Huang *et al*. 2017). However, this approach works only if the genotyping microarrays considered are highly similar; if the overlap between probes across the genotyping microarrays is low, then the overall resolution and sensitivity of CNV detection and analyses can be severely impacted.

To overcome this limitation, we developed MarkerMatch. MarkerMatch is a proximity-based algorithm that matches probes across different genotyping microarrays within specified genomic distance (a distance between a probe on reference array and those on the matched array) to maximize the number of probes considered in the CNV calling algorithm, thereby increasing the resolution and sensitivity of subsequent genome-wide CNV association analyses without affecting their specificity. MarkerMatch returns a list of probes for each of the matched arrays to be used in CNV calling. We tested MarkerMatch in two independent experiments: a within-array experiment to test the effects of probe downsampling on CNV calling, and a cross-array experiment to test the reliability of CNV calling in samples genotyped on multiple arrays.

## Materials and Methods

### Samples and Software

Samples from two independent cohorts (totalling 4,906 individuals) and three different Illumina array products (Global Screening Array, GSA; Omni2.5 array, OMNI; and Omni Express Exome array, OEE) were used to test and validate MarkerMatch algorithms. A summary of the cohorts, specific genotyping platforms, sample sizes, and probe density information is shown in Table 1. Briefly, we used existing genomic data from the Simons Simplex Collection (SSC) and Tourette Association of America International Consortium for Genetics (TAAICG), detailed descriptions of which are provided in the Supplementary Information (Fischbach and Lord 2010; Scharf *et al*. 2013; Sanders *et al*. 2015; Huang *et al*. 2017). Table 2 summarizes software used in this study.

**Table 1.**
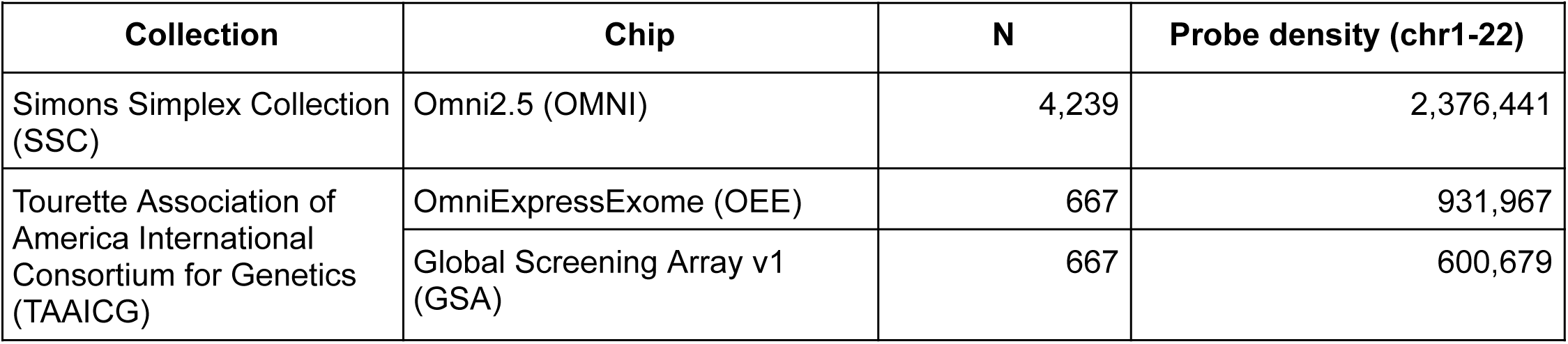
Samples used in this study.

| **Table 1.** Samples used in this study. |  |  |  |
| --- | --- | --- | --- |
| **Collection** | **Chip** | **N** | **Probe density (chr1-22)** |
| Simons Simplex Collection (SSC) | Omni2.5 (OMNI) | 4,239 | 2,376,441 |
| Tourette Association of America International Consortium for Genetics (TAAICG) | OmniExpressExome (OEE) | 667 | 931,967 |
|  | Global Screening Array v1 (GSA) | 667 | 600,679 |

**Table 2.** Software used in this study.

| **Table 2.** Software used in this study. |  |  |
| --- | --- | --- |
| **Software** | **Version** | **Reference** |
| betareg | 3.2-1 | (Cribari-Neto and Zeileis 2010) |
| GenomeStudio | 2.0.5 | (Illumina 2020) |
| ggpubr | 0.6.0 | (Kassambara 2023) |
| matrixStats | 0.52.2 | (Bengtsson *et al.* 2017) |
| numpy | 2.1.3 | (Harris *et al.* 2020) |
| PennCNV | 1.0.5 | (Wang *et al.* 2007, 2008; Diskin *et al.* 2008) |
| pandas | 2.2.3 | (The pandas development team 2024) |
| plink | 1.90 beta 7.4 | (Purcell *et al.* 2007) |
| python | 3.11.5 | (Python Software Foundation 2023) |
| R | 4.1.2 | (R Core Team 2021) |
| R Studio | 2024.12.1 | (Posit Team 2024) |
| tidyverse | 1.3.2 | (Wickham *et al.* 2019) |

### Development of MarkerMatch

MarkerMatch follows a simple loop and match algorithm (Figure 1A) to identify the best-matching probes between two manifests at a time. MarkerMatch takes in two annotated manifests, the smaller of which is considered the *reference* manifest and the larger the *matching* manifest. Both reference and matching manifests must contain the following information: (1) probe name, (2) chromosome, (3) genomic position, (4)

**Figure 1.**
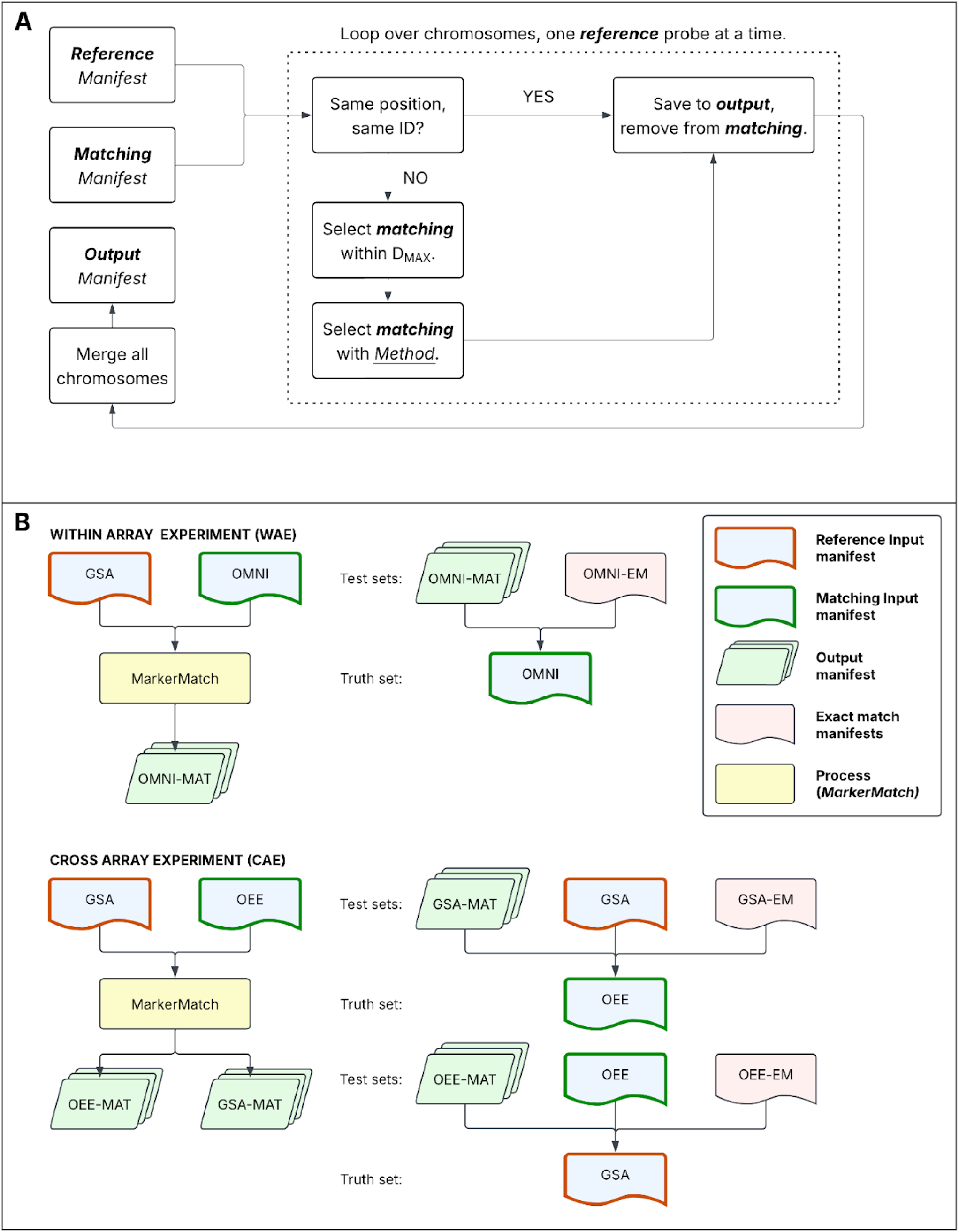
Panel. **A**. Diagram depicting the MarkerMatch algorithm. MarkerMatch follows a step-by-step process to identify the best match for each probe across the two selected manifests, while ensuring no duplicates. Specifically, in the first step (exact matching) MarkerMatch will take an intersection of probes from two manifests and keep them in the ***output***. The second step (*<u>Method</u>* matching): MarkerMatch will take a probe from the ***reference*** manifest and match it with all the remaining probes (those not used in the first step) in the ***matching*** *manifest* within the specified *D_MAX_*distance. A probe from the ***matched*** manifest that has the smallest difference in the selected *<u>Method</u>* from the selected probe from the ***reference*** manifest will be retained. Once a probe from the matching manifest is paired with a reference probe, it is removed from the ***matching*** manifest. This prevents it from being matched again, avoiding repetitive matching of identical probes from the *matched* manifest. This process will continue until all *reference* probes have been considered. **Panel B**. Graphical representation of experimental setup. Blue boxes represent unprocessed array data, with red borders representing ***reference*** manifests and green borders representing ***matching*** manifests. Yellow boxes represent the MarkerMatch algorithm for WAE (for all *Methods* and 10bp < *D_MAX_* < 5Mb) and CAE (for all *Methods* and *D_MAX_* = 10kb). Green boxes represent ***output*** manifests of the MarkerMatch algorithm (-MAT suffix indicates output manifests from *MarkerMatch*). Red boxes represent ***exact match*** manifests as currently used in CNV association analyses (intersections, also consensus manifests and-EM suffix). In WAE, we compared matched OMNI callsets to full OMNI as a truth set (also *Full Set*). In CAE, we compared matched GSA callsets to full OEE as a truth set, as well as matched OEE callsets to full GSA as a truthset. OMNI: Omni2.5 array, GSA: Global Screening Array, OEE: Omni Express Exome array. These processes have been repeated for each iteration of *Method* and *D_MAX_* combination.

B-allele frequency (BAF), (5) mean of the log-R ratio (LRR mean), and (6) standard deviation of LRR (LRR sd). In addition to the two manifests, the MarkerMatch function requires two pre-specified parameters: *D_MAX_*(which determines the maximum allowable distance from which a marker can be selected) and *Method* (which determines what metric should be prioritized for matching), and returns a 1:1 set of matched probes from the two manifests. There are 4 options for the *Method* parameter in the MarkerMatch function that can be used to select probes: *position*, *BAF*, *LRR mean*, and *LRR sd*.

MarkerMatch follows a step-by-step process to identify the best match for each probe across the two selected manifests, while ensuring no duplicates. Specifically, in the first step (exact matching) MarkerMatch will take an intersection of probes from two manifests and keep them. In the second step (nearby matching), MarkerMatch will take a probe from the *reference* manifest and match it with all the remaining probes (those not used in the first step) in the *matching manifest* within the specified *D_MAX_* distance. A probe from this set that has the smallest difference in the selected *Method* from the identified probe in the *reference* manifest will be selected and saved into the *output manifest*. Once a probe from the matching manifest is paired with a reference probe, it is removed from the matching manifest to avoid it being matched again. This process will continue until all *reference* probes have been considered.

MarkerMatch was written as an R script dependent only on *tidyverse* packages (Wickham *et al*. 2019) and is easy and flexible to implement. An alternative implementation in Python has also been written.

### CNV Calling

Briefly, the Illumina GenomeStudio final reports were exported from for each array and passed into PennCNV to call CNVs (Wang *et al*. 2007; Illumina 2020). Data preprocessing, array clustering, genotyping quality control, CNV calling and quality control are described in detail in the Supplementary Information.

For the Within-Array Experiment (WAE) utilizing SSC data, we performed CNV calling for the GSA-matched OMNI manifest at variable MarkerMatch matching metrics (position, LRR mean, LRR standard deviation, and BAF) and variable maximum allowable distances (10bp, 50bp, 100bp, 500bp, 1kb, 5kb, 10kb, 50kb, 100kb, 500kb, 1Mb, and 5Mb), as well as for the full OMNI manifest (full set) and for the intersection of the OMNI manifest with the GSA manifest (exact match). We additionally modeled validation metric performance of MarkerMatch callsets relative to the *Full Set* to determine the optimal *D_MAX_* parameter setting.

For the Cross-Array Experiment (CAE) utilizing TAAICG data, we performed CNV calling for the full OEE manifest (full OEE set), the full GSA manifest (full GSA set), an intersection of the OEE manifest with the GSA manifest (exact match), as well as the GSA-matched OEE manifest at a fixed maximum allowable distance of 10kb and variable MarkerMatch matching metrics (position, LRR mean, LRR standard deviation, and BAF).

### Validation

We performed two independent validations of the MarkerMatch: the Within-Array Experiment (WAE) examined MarkerMatch’s performance in probe reductions within the same array (OMNI data from SSC), and the Cross-Array Experiment (CAE), which examined MarkerMatch’s performance across arrays using the OEE and GSA data from TAAICG. A detailed explanation of these two experiments is provided in the Supplementary Information and the graphical representation is shown in Figure 1B.

For each experiment, we derived a partial confusion matrix including true positive, false positive, and false negative counts. Truth sets were full set OMNI, OEE, and GSA CNV callsets. True negative counts were impossible to determine as we do not know the true copy-number states for the examined genomes. Based on the partial confusion matrix, we derived the following metrics: true positive rate (sensitivity, recall), false negative rate (FNR), positive predictive value (PPV, precision), false discovery rate (FDR), F1 score (F1; harmonic mean of precision and recall), Fowlkes–Mallows index (FMI; geometric mean of precision and recall); and Jaccard index (JI; ratio of the intersection to the union of the two sets). These data were used to visually and quantitatively inspect performance of specific *D_MAX_* and *Method* parameter configurations in MarkerMatch callsets, and to inform decisions for optimal parameter selection. The full methodology is available in the Supplementary Information.

## Results

### Implementation of MarkerMatch

The MarkerMatch algorithm runs about 4 times faster in Python than R (Figure 2) across all *Methods*, with an average R run time of 2.19 minutes and an average Python run time of 0.54 minutes on chromosome 22. The total genome-wide run for MarkerMatch with parameters *D_MAX_* = 10kb and *Method* = Distance was 20.52 hours in R and 7.22 hours in Python.

**Figure 2.**
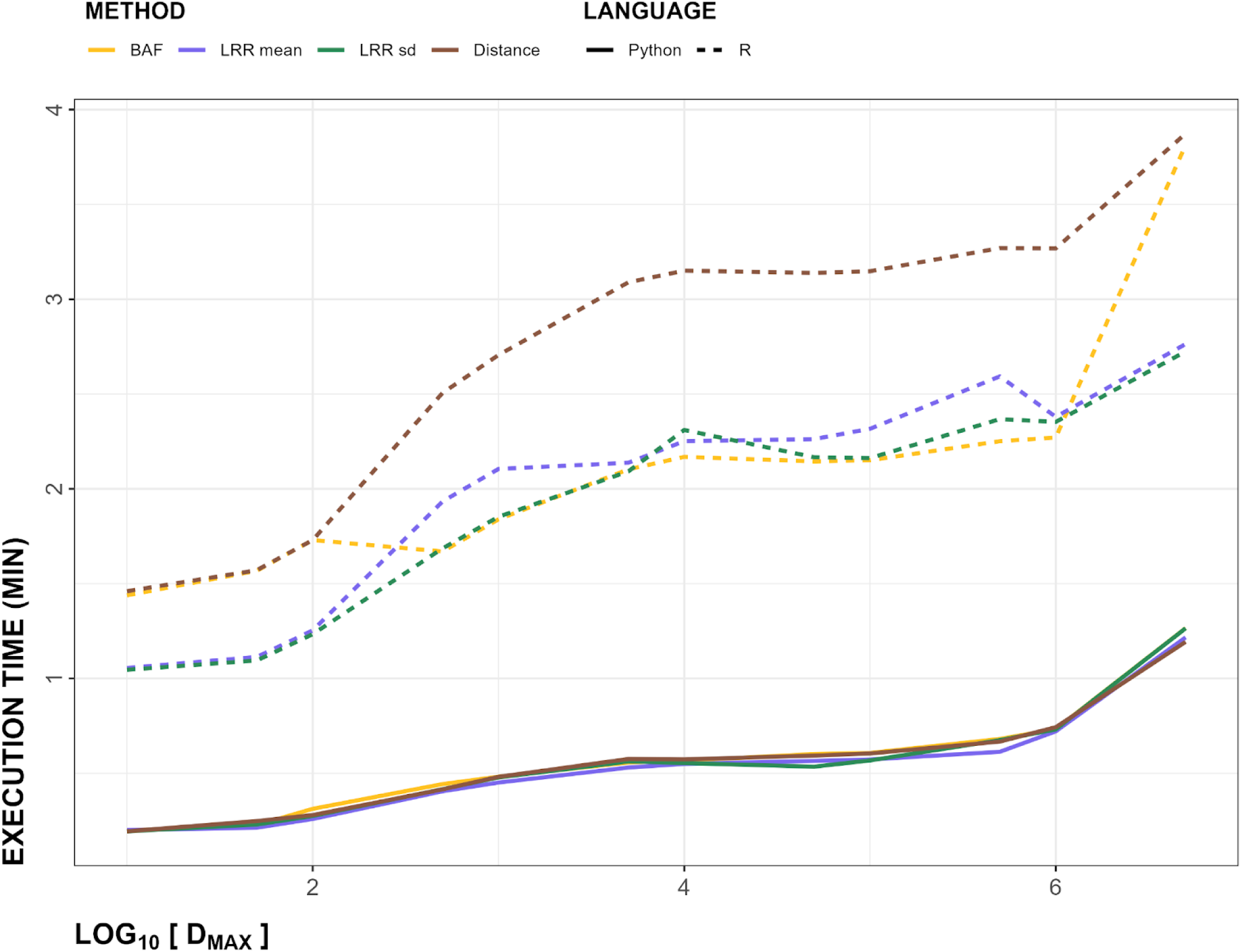
Execution time curves for chromosome 22, shown as function of time (in minutes) on the y-axis, and log_10_ of maximum matching distance *D_MAX_* (in bp) on x-axis. Solid lines represent Python, while dashed lines represent R execution times. *Method* is shown in colors (BAF in yellow, LRR mean in purple, LRR sd in green, and Distance in brown).

Analysis of array coverage shows successful recovery of GSA coverage when matched with both the OMNI and OEE arrays (Figure 3A-D, Table 3, Supplemental Table S1B). For the OMNI array, coverage plateaued at a D_MAX_ value of 10kb, with 28% coverage of the OMNI array (Figure 3A) and 98% coverage of the GSA array (Figure 3B). In contrast, the exact match approach resulted in retention of 5% of markers from the OMNI array (Figure 3A) and 19% of markers from the GSA array (Figure 3B).

**Figure 3.**
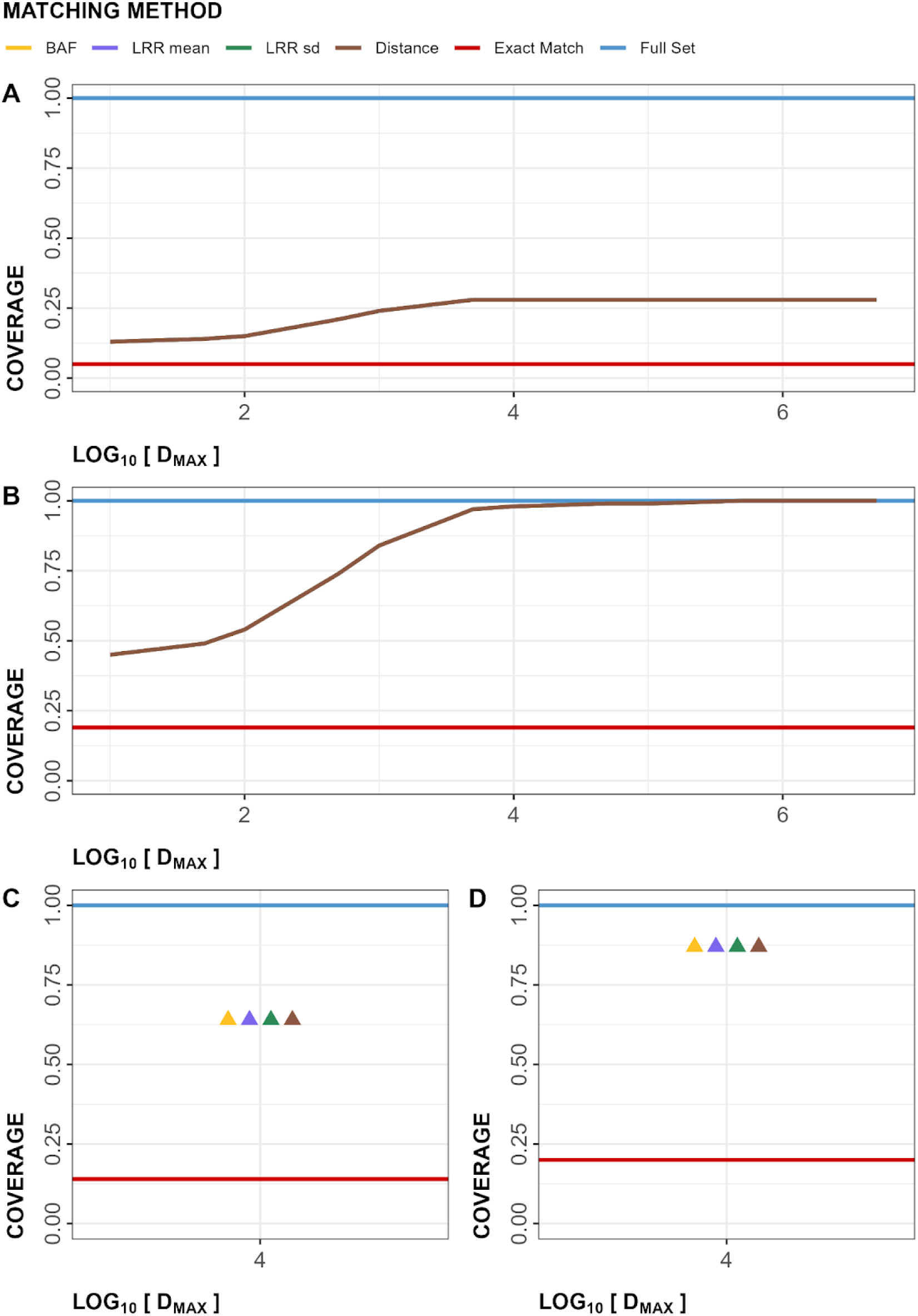
Figure showing coverage of arrays in Within-Array Experiment (WAE; A: OMNI, B: GSA) and Cross-Array Experiment (CAE; C: OEE, D: GSA). In all graphs, y-axis is showing the coverage rates and the x-axis is showing maximum allowable distances in base-pairs, log_10_(D_MAX_). Note: Lines for all matching Methods in panels A and B are overlapped. Points on panels C and D are horizontally jittered for visibility, but log_10_(D_MAX_) is 4 for all matching Methods. OMNI: Omni2.5 array, GSA: Global Screening Array, OEE: Omni Express Exome array.

**Table 3.** Summaries of *MarkerMatch* outcomes at *D_MAX_ = 10kb* across all *Method* parameters (BAF, LRR mean, LRR sd, and Distance), as well as *Full Set* and *Exact Match* reference comparisons. Coverage is displayed in rate; gaps and LRR sds are displayed in log_10_[median] (log_10_[IQR]); BAFs and LRR means are displayed in median (IQR). Within-array experiment outcomes are detailed in *OMNI array, OMNI matched to GSA* and *GSA array, OMNI matched to GSA* segments. Cross-array experiment outcomes are detailed in *OEE array, OEE matched to GSA* and *GSA array, OEE matched to GSA* segments.

| Measure | Full Set | BAF | LRR mean | LRR sd | Distance | Exact Match |
| --- | --- | --- | --- | --- | --- | --- |
| ***OMNI array, OMNI matched to GSA*** |  |  |  |  |  |  |
| Coverage | 1.00 | 0.28 | 0.28 | 0.28 | 0.28 | 0.05 |
| Gaps | 2.79 (0.79) | 3.36 (0.80) | 3.36 (0.80) | 3.36 (0.80) | 3.36 (0.82) | 4.04 (0.74) |
| BAF | 0.07 (0.25) | 0.04 (0.20) | 0.08 (0.24) | 0.11 (0.27) | 0.11 (0.26) | 0.26 (0.22) |
| LRR sd | 0.10 (0.04) | 0.10 (0.04) | 0.10 (0.05) | 0.10 (0.05) | 0.10 (0.04) | 0.10 (0.04) |
| LRR mean | -0.002 (0.004) | -0.002 (0.003) | -0.002 (0.004) | -0.002 (0.004) | -0.002 (0.004) | -0.001 (0.004) |
| ***GSA array, OMNI matched to GSA*** |  |  |  |  |  |  |
| Coverage | 1.00 | 0.98 | 0.98 | 0.98 | 0.98 | 0.19 |
| ***OEE array, OEE matched to GSA*** |  |  |  |  |  |  |
| Coverage | 1.00 | 0.64 | 0.64 | 0.64 | 0.64 | 0.14 |
| Gaps | 3.14 (1.15) | 3.38 (0.95) | 3.38 (0.93) | 3.38 (0.91) | 3.39 (0.91) | 4.03 (0.74) |
| BAF | 0.14 (0.31) | 0.13 (0.28) | 0.16 (0.29) | 0.16 (0.30) | 0.17 (0.29) | 0.25 (0.22) |
| LRR sd | 0.13 (0.06) | 0.13 (0.06) | 0.13 (0.06) | 0.12 (0.06) | 0.13 (0.06) | 0.13 (0.05) |
| LRR mean | -0.003 (0.009) | -0.003 (0.009) | -0.003 (0.009) | -0.003 (0.008) | -0.003 (0.008) | -0.003 (0.009) |
| ***GSA array, OEE matched to GSA*** |  |  |  |  |  |  |
| Coverage | 1.00 | 0.87 | 0.87 | 0.87 | 0.87 | 0.20 |
| Gaps | 3.35 (0.86) | 3.40 (0.85) | 3.40 (0.85) | 3.40 (0.85) | 3.40 (0.85) | 4.03 (0.74) |
| BAF | 0.04 (0.18) | 0.04 (0.20) | 0.04 (0.200) | 0.04 (0.20) | 0.04 (0.20) | 0.25 (0.22) |
| LRR sd | 0.11 (0.07) | 0.11 (0.07) | 0.11 (0.07) | 0.11 (0.07) | 0.11 (0.07) | 0.11 (0.05) |
| LRR mean | -0.003 (0.006) | -0.003 (0.006) | -0.003 (0.006) | -0.003 (0.006) | -0.003 (0.006) | -0.002 (0.006) |

For the OEE array, coverage at D_MAX_ = 10kb was 63% of the OEE array (Figure 3C) and 88% of the GSA array (Figure 3D). The exact match approach resulted in the retention of 14% markers from the OEE array (Figure 3C) and 20% markers from the GSA array (Figure 3D).

Inter-marker gaps (i.e., gaps between SNPs used by PennCNV to make CNV calls) were also reduced in size on both the OMNI (median 2.2kb at *D_MAX_* = 10kb) and OEE (median 2.2kb at *D_MAX_*= 10kb) arrays matched to the GSA array, relative to their *Exact Match* counterparts with a median of about 11kb (Supplemental Figure S1A-C, Table 3, Supplemental Table S1C).

BAF distributions of *Full Set* arrays were better approximated by MarkerMatched than *Exact Match* configurations (Supplemental Figure S2A-C, Table 3, Supplemental Table S1D). Notably, the median BAF differed for each *Method*, with the median BAF being 0.04 for *Method* = BAF, 0.08 for *Method* = LRR mean, 0.11 for *Method* = LRR sd, and 0.11 for *Method* = Distance arrays, whereas the median BAF values for *Exact Match* and *Full Set* were 0.26 and 0.07, respectively.

Conversely, LRR sds and LRR means showed relatively little variability between various *Method* and *D_MAX_* configurations of MarkerMatch, as well as between MarkerMatch configurations and *Full Set* and *Exact Match*, with median LRR sd values around 1.26 and LRR mean values around-0.002 (Supplemental Figures S3A-C, S5A-C, S6A-C, Table 3, Supplemental Tables S1E, S1F).

### Within-Array Experiment (WAE)

Within-Array Experiment (WAE) results are summarized in Table 4, and in Supplemental Tables S1G-I. Briefly, the *Full Set* callset resulted in the most CNV calls (low-stringency QC = 288,602; medium-stringency QC = 56,017) and *Exact Match* resulted in the fewest (low-stringency QC = 11,173; medium-stringency QC = 6,776). MarkerMatched callsets counted 3-8 times more CNV calls relative to *Exact Match* across all four *Methods* at *D_MAX_* = 10kb (low-stringency QC range 77,132 - 84,140; medium-stringency QC range 31,430 - 34,272).

**Table 4.**
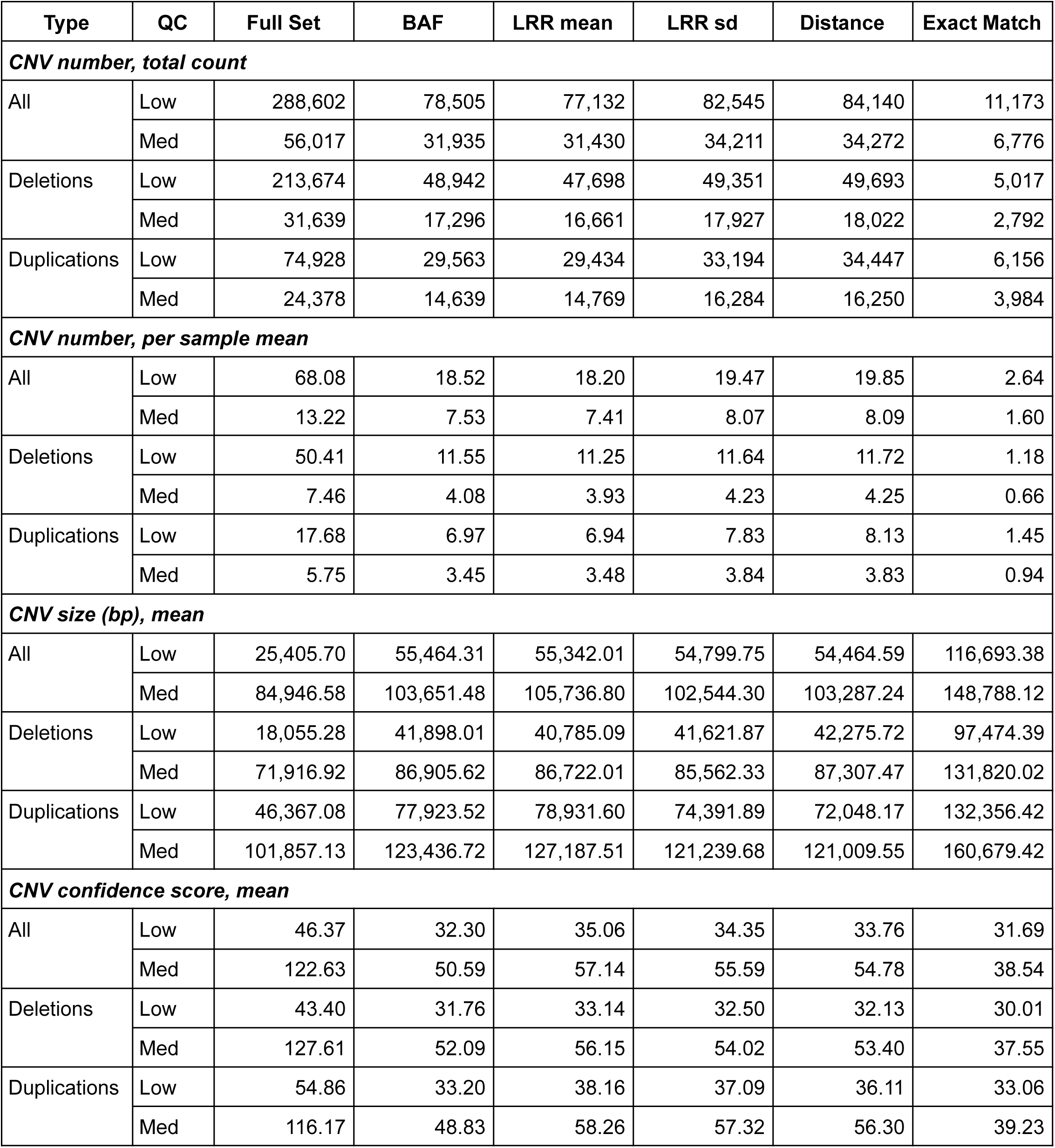
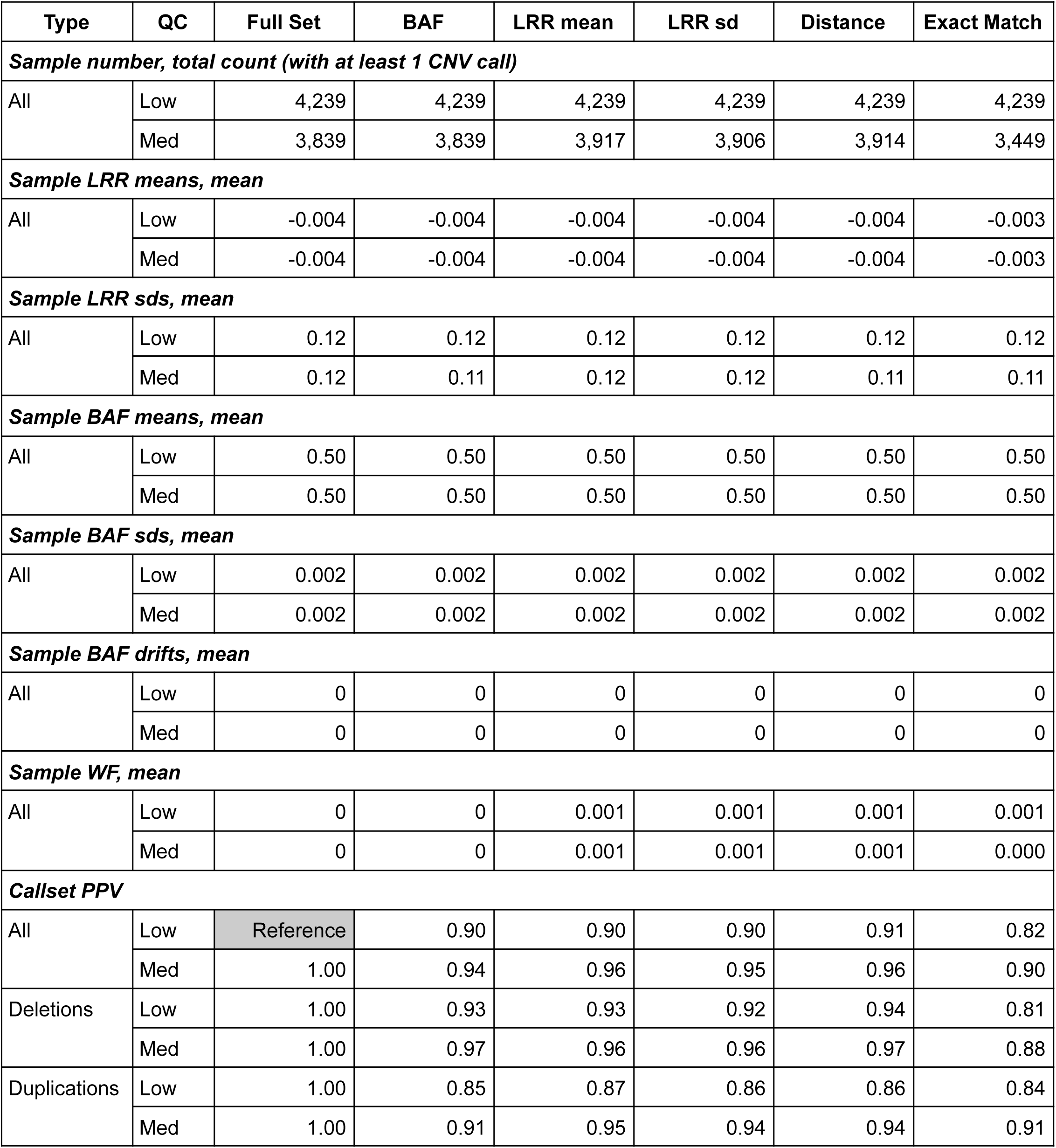
Summaries of *MarkerMatch* CNV callsets at *D_MAX_ = 10kb* across all *Method* parameters (BAF, LRR mean, LRR sd, and Distance), as well as *Full Set* and *Exact Match* reference comparisons for OMNI array. Full tables including other *D_MAX_* parameter values and specific size bins, as well as other statistics like medians and IQRs, are available in the Supplemental Tables S1G-I. Note: callset PPVs are based on comparisons to OMNI array *Full Set* callset. In the QC column, *low* stands for low-stringency QC and *med* stands for medium-stringency QC.

| Type | QC | Full Set | BAF | LRR mean | LRR sd | Distance | Exact Match |
| --- | --- | --- | --- | --- | --- | --- | --- |
| **CNV number, total count** |  |  |  |  |  |  |  |
| All | Low | 288,602 | 78,505 | 77,132 | 82,545 | 84,140 | 11,173 |
|  | Med | 56,017 | 31,935 | 31,430 | 34,211 | 34,272 | 6,776 |
| Deletions | Low | 213,674 | 48,942 | 47,698 | 49,351 | 49,693 | 5,017 |
|  | Med | 31,639 | 17,296 | 16,661 | 17,927 | 18,022 | 2,792 |
| Duplications | Low | 74,928 | 29,563 | 29,434 | 33,194 | 34,447 | 6,156 |
|  | Med | 24,378 | 14,639 | 14,769 | 16,284 | 16,250 | 3,984 |
| **CNV number, per sample mean** |  |  |  |  |  |  |  |
| All | Low | 68.08 | 18.52 | 18.20 | 19.47 | 19.85 | 2.64 |
|  | Med | 13.22 | 7.53 | 7.41 | 8.07 | 8.09 | 1.60 |
| Deletions | Low | 50.41 | 11.55 | 11.25 | 11.64 | 11.72 | 1.18 |
|  | Med | 7.46 | 4.08 | 3.93 | 4.23 | 4.25 | 0.66 |
| Duplications | Low | 17.68 | 6.97 | 6.94 | 7.83 | 8.13 | 1.45 |
|  | Med | 5.75 | 3.45 | 3.48 | 3.84 | 3.83 | 0.94 |
| **CNV size (bp), mean** |  |  |  |  |  |  |  |
| All | Low | 25,405.70 | 55,464.31 | 55,342.01 | 54,799.75 | 54,464.59 | 116,693.38 |
|  | Med | 84,946.58 | 103,651.48 | 105,736.80 | 102,544.30 | 103,287.24 | 148,788.12 |
| Deletions | Low | 18,055.28 | 41,898.01 | 40,785.09 | 41,621.87 | 42,275.72 | 97,474.39 |
|  | Med | 71,916.92 | 86,905.62 | 86,722.01 | 85,562.33 | 87,307.47 | 131,820.02 |
| Duplications | Low | 46,367.08 | 77,923.52 | 78,931.60 | 74,391.89 | 72,048.17 | 132,356.42 |
|  | Med | 101,857.13 | 123,436.72 | 127,187.51 | 121,239.68 | 121,009.55 | 160,679.42 |
| **CNV confidence score, mean** |  |  |  |  |  |  |  |
| All | Low | 46.37 | 32.30 | 35.06 | 34.35 | 33.76 | 31.69 |
|  | Med | 122.63 | 50.59 | 57.14 | 55.59 | 54.78 | 38.54 |
| Deletions | Low | 43.40 | 31.76 | 33.14 | 32.50 | 32.13 | 30.01 |
|  | Med | 127.61 | 52.09 | 56.15 | 54.02 | 53.40 | 37.55 |
| Duplications | Low | 54.86 | 33.20 | 38.16 | 37.09 | 36.11 | 33.06 |
|  | Med | 116.17 | 48.83 | 58.26 | 57.32 | 56.30 | 39.23 |

| **Table 4.** (continued) |  |  |  |  |  |  |  |
| --- | --- | --- | --- | --- | --- | --- | --- |
| Type | QC | Full Set | BAF | LRR mean | LRR sd | Distance | Exact Match |
| **Sample number, total count (with at least 1 CNV call)** |  |  |  |  |  |  |  |
| All | Low | 4,239 | 4,239 | 4,239 | 4,239 | 4,239 | 4,239 |
|  | Med | 3,839 | 3,839 | 3,917 | 3,906 | 3,914 | 3,449 |
| **Sample LRR means, mean** |  |  |  |  |  |  |  |
| All | Low | -0.004 | -0.004 | -0.004 | -0.004 | -0.004 | -0.003 |
|  | Med | -0.004 | -0.004 | -0.004 | -0.004 | -0.004 | -0.003 |
| **Sample LRR sds, mean** |  |  |  |  |  |  |  |
| All | Low | 0.12 | 0.12 | 0.12 | 0.12 | 0.12 | 0.12 |
|  | Med | 0.12 | 0.11 | 0.12 | 0.12 | 0.11 | 0.11 |
| **Sample BAF means, mean** |  |  |  |  |  |  |  |
| All | Low | 0.50 | 0.50 | 0.50 | 0.50 | 0.50 | 0.50 |
|  | Med | 0.50 | 0.50 | 0.50 | 0.50 | 0.50 | 0.50 |
| **Sample BAF sds, mean** |  |  |  |  |  |  |  |
| All | Low | 0.002 | 0.002 | 0.002 | 0.002 | 0.002 | 0.002 |
|  | Med | 0.002 | 0.002 | 0.002 | 0.002 | 0.002 | 0.002 |
| **Sample BAF drifts, mean** |  |  |  |  |  |  |  |
| All | Low | 0 | 0 | 0 | 0 | 0 | 0 |
|  | Med | 0 | 0 | 0 | 0 | 0 | 0 |
| **Sample WF, mean** |  |  |  |  |  |  |  |
| All | Low | 0 | 0 | 0.001 | 0.001 | 0.001 | 0.001 |
|  | Med | 0 | 0 | 0.001 | 0.001 | 0.001 | 0.000 |
| **Callset PPV** |  |  |  |  |  |  |  |
| All | Low | Reference | 0.90 | 0.90 | 0.90 | 0.91 | 0.82 |
|  | Med | 1.00 | 0.94 | 0.96 | 0.95 | 0.96 | 0.90 |
| Deletions | Low | 1.00 | 0.93 | 0.93 | 0.92 | 0.94 | 0.81 |
|  | Med | 1.00 | 0.97 | 0.96 | 0.96 | 0.97 | 0.88 |
| Duplications | Low | 1.00 | 0.85 | 0.87 | 0.86 | 0.86 | 0.84 |
|  | Med | 1.00 | 0.91 | 0.95 | 0.94 | 0.94 | 0.91 |

WAE per-sample CNV calls, summarized in Table 4 and Supplemental Table S1G, were highest for the *Full Set* callset (low-stringency QC = 68.1; medium-stringency QC = 13.2), lowest for *Exact Match* (low-stringency QC = 2.6, medium-stringency QC = 1.6), and about 3-7 times the *Exact Match* in MarkerMatched callsets across all four *Methods* at *D_MAX_* = 10kb (low-stringency QC range 18.2 - 19.9; medium-stringency QC range 7.4 - 8.1).

The average CNV sizes (in bp) identified in WAE were smaller on denser *MarkerMatched* configurations (Table 4, Supplemental Table S1G). *Full Set* averages were the smallest (low-stringency QC 25,405.7 bp; medium-stringency QC 95,946.6 bp), *Exact Match* were the largest (low-stringency QC 116,693.4bp; medium-stringency QC 148,788.1bp), and the MarkerMatched callsets were intermediate (low-stringency QC range 55,342.0bp - 54,799.7bp; medium-stringency QC 102,544.3bp - 105,736.8bp).

Average confidence scores in WAE were larger on denser configurations (Table 4, Supplemental Table S1G), ranging from 122.6 on medium-stringency QC *Full Set* to 31.7 on low-stringency QC *Exact Match*. As with other metrics, CNV confidence scores were intermediate for MarkerMatch callsets (low-stringency QC range 32.3 - 35.1; medium-stringency QC range 50.6 - 57.1).

Overall number of samples with at least one CNV call were consistent after low-stringency QC across all approaches (N = 4,239), although they did vary somewhat after medium-stringency QC (N range 3,449 - 3,917) as shown in Table 4 and Supplemental Table S1H.

Sample-specific metrics did not vary substantially across low-stringency QC/medium-stringency QC or *Full Set*/*Exact Match*/MarkerMatched configurations, with average *LRR means* at-0.004, *LRR sds* at 0.114-0.121, *BAF means* at 0.503, *BAF sds* at 0.002, *BAF drifts* at 0.000, and *WFs* at 0.000-0.001 (Table 4, Supplemental Table S1H).

### Genome-Wide Validation

WAE genome-wide validation resulted in higher PPVs for MarkerMatch callsets matched at *D_MAX_*= 10kb for all *Method* parameters (ranges 0.896 - 0.907 and 0.943 - 0.958 for low-and medium-stringency QC, respectively) compared to *Exact Match* (0.824 and 0.900 for low-and medium-stringency QC, respectively), and for low-stringency QC and medium-stringency QC callsets, respectively (Figure 4, Table 4, Supplemental Table S1I).

**Figure 4.**
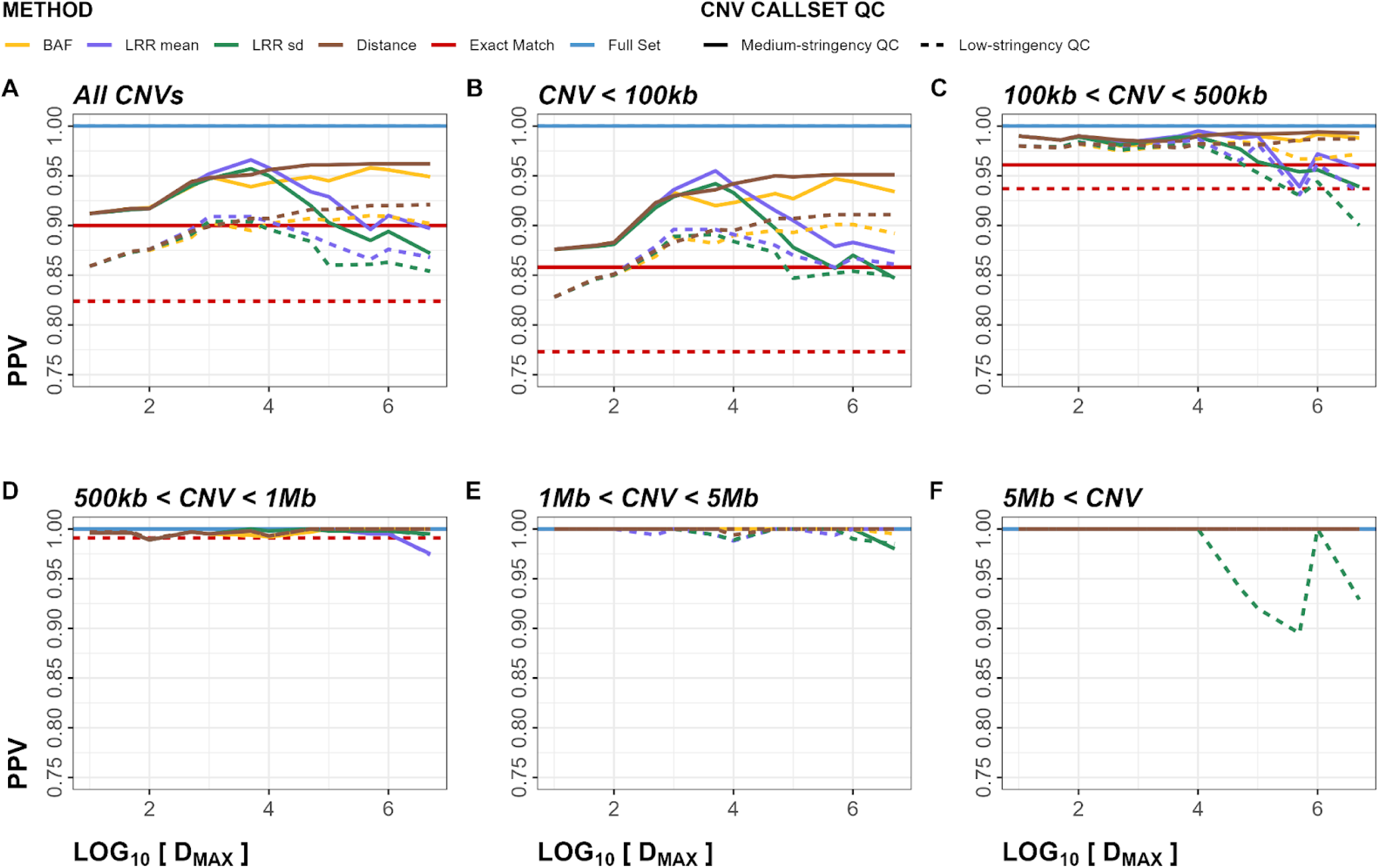
Within-Array Experiment (WAE) positive predictive value (PPV) plots for both deletions and duplications. Panels A-F show PPV metrics stratified by CNV size. Dashed line represents low-stringency QC, solid line represents medium-stringency QC.

Other metrics performed similarly, with larger CNVs having consistently better performance than smaller CNVs (Supplemental Figures S7-12, Supplemental Table S1I). Additionally, duplications had a slightly better performance than deletions (Supplemental Figure S12-25, Supplemental Table S1I). The genome-wide PPV plots (Figure 4) indicated that, in the majority of *D_MAX_* and *Method* parameter configurations, MarkerMatch somewhat or substantially outperformed the *Exact Match* approach.

### Regional Validation

Filtering on the medium-stringency QC *Exact Match* callset resulted in region-specific PPVs of 0.99 (telomeric), 0.90 (centromeric), and 0.96 (segmental duplications), whereas the genome-wide, unfiltered callset had a PPV of 0.90. Performance of MarkerMatch callsets at *D_MAX_* = 10kb was consistent across various *Method* parameters in the medium-stringency QC callset, with region-specific average PPVs of 0.98 (telomeric), 0.93 (centromeric), and 0.87 (segmental duplications).

Detailed reports for other *D_MAX_* parameter settings and performance metrics are available in the Supplemental Table S1J. These data are graphically shown for all validation metrics in the Supplemental Figures S26-32.

### Selection of D_MAX_

Visual inspection of plotted validation metrics after *loess*-smoothing and *Full Set* scaling (see the supplement equation Eq. 8) indicates that, for the majority of metrics (across sensitivity, PPV, F1, FMI, and JI), regardless of CNV size (all, CNV < 100kb, 100kb < CNV < 500kb, or 500kb < CNV < 1Mb), CNV type (all, deletions, or duplications), or *Method* parameter (BAF, LRR mean, LRR sd, or Distance), the peak, plateau, and/or inflection point occurred in the range of *D_MAX_* between 10kb and 100kb (Supplemental Figures S32-37). Further inspection indicated that *D_MAX_*= 10kb was the optimal maximum allowable distance to match within, and was thus chosen as the *D_MAX_* setting for the Cross-Array Experiment (CAE).

### Selection of Method

Visual inspection of the plotted validation metrics after *Full Set* scaling (see the supplement equation Eq. 8) indicated that, for the majority of metrics (across sensitivity, PPV, F1, FMI, and JI), regardless of CNV size (all, CNV < 100kb, 100kb < CNV < 500kb, or 500kb < CNV < 1Mb), CNV type (all, deletions, or duplications), the highest median performance occurred with Distance *Method* (Supplemental Figures S38-42). This was particularly evident in the PPV metrics, where Distance *Method* outperformed BAF, and substantially outperformed LRR mean and LRR sd metrics, except in the 500kb < CNV < 1Mb bin where all four methods appeared to perform about the same (Supplemental Figure S39). The mean-differences between PPV values, unstratified by CNV type or CNV size, of Distance *Method*, and LRR sd and LRR mean were nominally significant in the Welch two sample t-test (*p* = 0.006 and *p* = 0.027, for LRR sd and LRR mean, respectively). However, no significant differences were observed between any two *Method*’s metrics, for any CNV type and CNV size strata after FDR correction (Supplemental Table S1K). We thus opted to examine all *Method* parameters in the Cross-Array Experiment (CAE).

### Cross-Array Experiment (CAE)

Cross-Array Experiment (CAE) results are summarized in Tables 5 and 6, Supplemental Tables S1L-N.

**Table 5.**
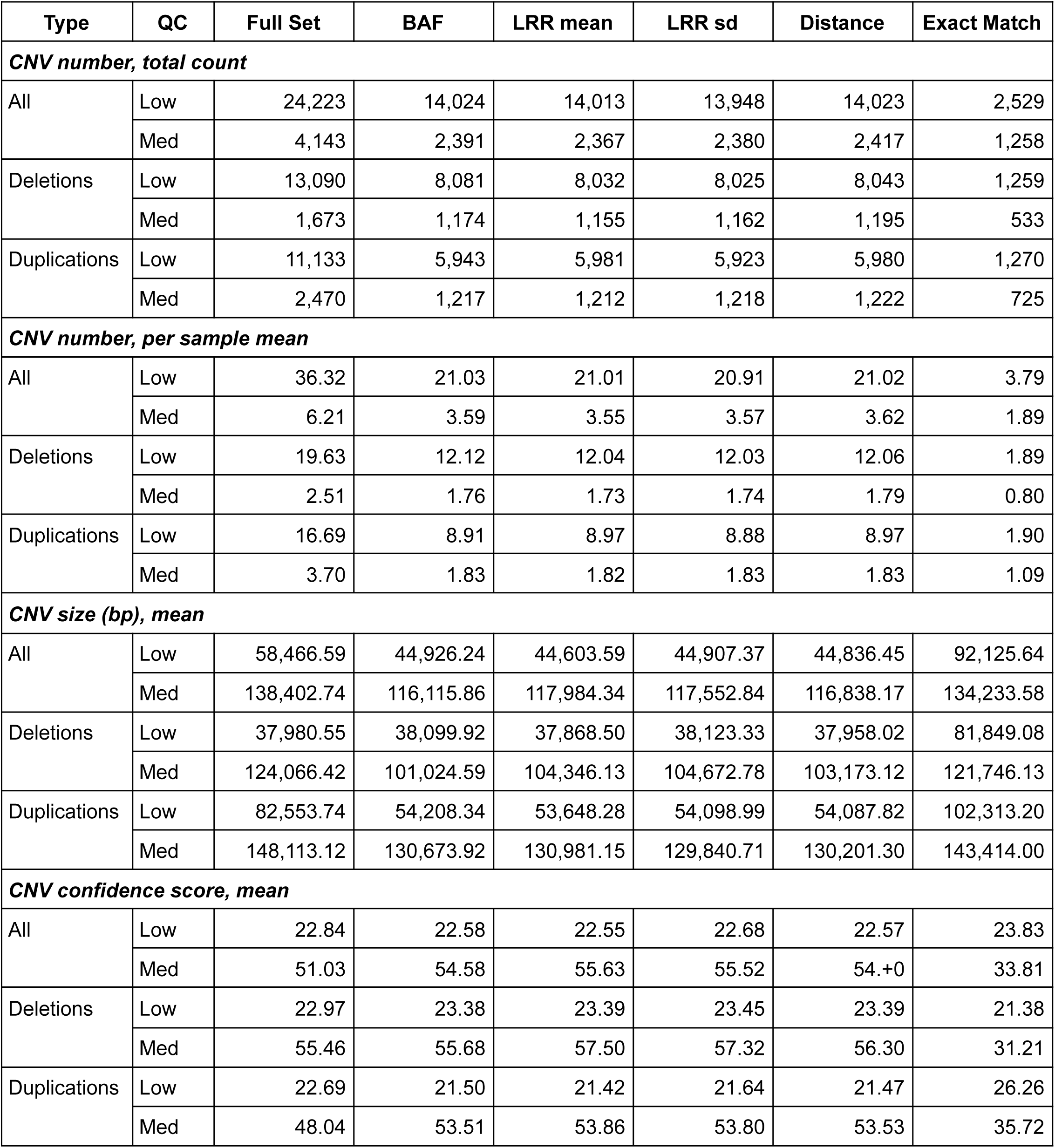
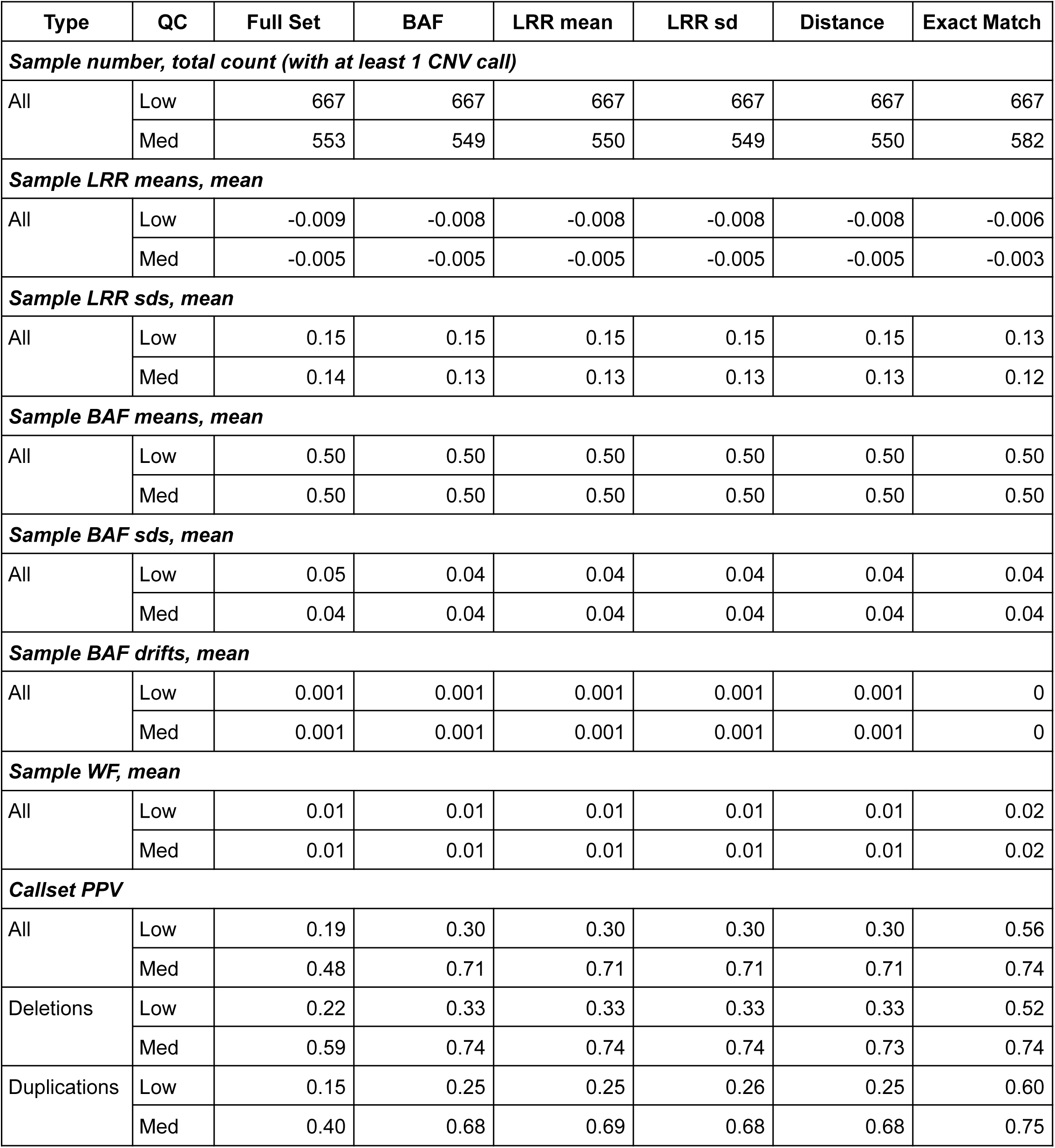
Summaries of *MarkerMatch* CNV callsets at *D_MAX_ = 10kb* across all *Method* parameters (BAF, LRR mean, LRR sd, and Distance), as well as *Full Set* and *Exact Match* reference comparisons for GSA array (*Ref* = OEE). Full tables including specific size bins, as well as other statistics like medians and IQRs, are available in the Supplemental Tables S1L-N. Note: callset PPVs are based on comparisons to OEE array *Full Set* callset. In the QC column, *low* stands for low-stringency QC and *med* stands for medium-stringency QC.

**Table 6.** Summaries of *MarkerMatch* CNV callsets at *D_MAX_ = 10kb* across all *Method* parameters (BAF, LRR mean, LRR sd, and Distance), as well as *Full Set* and *Exact Match* reference comparisons for OEE array (*Ref* = GSA). Full tables including specific size bins, as well as other statistics like medians and IQRs, are available in the Supplemental Tables S1L-N. Note: callset PPVs are based on comparisons to GSA array *Full Set* callset. In the QC column, *low* stands for low-stringency QC and *med* stands for medium-stringency QC.

| Type | QC | Full Set | BAF | LRR mean | LRR sd | Distance | Exact Match |
| --- | --- | --- | --- | --- | --- | --- | --- |
| **CNV number, total count** |  |  |  |  |  |  |  |
| All | Low | 21,399 | 11,811 | 10,491 | 10,790 | 11,033 | 1,883 |
|  | Med | 1,974 | 1,808 | 2,192 | 2,311 | 2,002 | 936 |
| Deletions | Low | 15,623 | 8,243 | 7,157 | 7,350 | 7,536 | 1,066 |
|  | Med | 995 | 886 | 1,049 | 1,086 | 927 | 420 |
| Duplications | Low | 5,776 | 3,568 | 3,334 | 3,440 | 3,497 | 817 |
|  | Med | 979 | 922 | 1,143 | 1,225 | 1,075 | 516 |
| **CNV number, per sample mean** |  |  |  |  |  |  |  |
| All | Low | 32.08 | 17.71 | 15.73 | 16.18 | 16.54 | 2.82 |
|  | Med | 2.96 | 2.71 | 3.29 | 3.47 | 3.00 | 1.40 |
| Deletions | Low | 23.42 | 12.36 | 10.73 | 11.02 | 11.30 | 1.60 |
|  | Med | 1.49 | 1.33 | 1.57 | 1.63 | 1.39 | 0.63 |
| Duplications | Low | 8.66 | 5.35 | 5.00 | 5.16 | 5.24 | 1.23 |
|  | Med | 1.47 | 1.38 | 1.71 | 1.84 | 1.61 | 0.77 |
| **CNV size (bp), mean** |  |  |  |  |  |  |  |
| All | Low | 40,102.77 | 57,901.15 | 62,706.29 | 62,385.87 | 62,007.95 | 104,828.30 |
|  | Med | 112,111.80 | 114,975.27 | 118,664.14 | 119,784.10 | 119,742.39 | 138,323.57 |
| Deletions | Low | 32,090.91 | 45,983.16 | 49,280.68 | 50,044.79 | 49,170.34 | 84,850.61 |
|  | Med | 90,390.63 | 95,544.90 | 99,967.46 | 100,387.38 | 104,106.94 | 121,483.52 |
| Duplications | Low | 61,773.34 | 85,434.76 | 91,526.64 | 88,754.15 | 89,672.88 | 130,894.68 |
|  | Med | 134,187.96 | 133,646.98 | 135,823.21 | 136,979.88 | 133,225.23 | 152,030.59 |
| **CNV confidence score, mean** |  |  |  |  |  |  |  |
| All | Low | 32.15 | 29.74 | 32.46 | 31.566 | 31.64 | 27.88 |
|  | Med | 99.27 | 69.98 | 77.63 | 75.12 | 77.25 | 36.79 |
| Deletions | Low | 29.29 | 26.69 | 28.24 | 27.70 | 27.57 | 23.17 |
|  | Med | 94.56 | 67.28 | 73.84 | 72.50 | 75.88 | 32.88 |
| Duplications | Low | 39.90 | 36.79 | 41.52 | 39.82 | 40.40 | 34.03 |
|  | Med | 104.04 | 72.57 | 81.10 | 77.44 | 78.43 | 39.98 |

| **Table 6.** (continued) |  |  |  |  |  |  |  |
| --- | --- | --- | --- | --- | --- | --- | --- |
| **Type** | **QC** | **Full Set** | **BAF** | **LRR mean** | **LRR sd** | **Distance** | **Exact Match** |
| **Sample number, total count (with at least 1 CNV call)** |  |  |  |  |  |  |  |
| All | Low | 667 | 667 | 667 | 667 | 667 | 667 |
|  | Med | 400 | 408 | 584 | 608 | 512 | 515 |
| **Sample LRR means, mean** |  |  |  |  |  |  |  |
| All | Low | -0.003 | -0.003 | -0.003 | -0.003 | -0.003 | -0.005 |
|  | Med | -0.002 | -0.002 | -0.002 | -0.002 | -0.002 | -0.003 |
| **Sample LRR sds, mean** |  |  |  |  |  |  |  |
| All | Low | 0.12 | 0.12 | 0.12 | 0.11 | 0.12 | 0.13 |
|  | Med | 0.11 | 0.11 | 0.11 | 0.11 | 0.11 | 0.12 |
| **Sample BAF means, mean** |  |  |  |  |  |  |  |
| All | Low | 0.50 | 0.50 | 0.50 | 0.50 | 0.50 | 0.50 |
|  | Med | 0.50 | 0.50 | 0.50 | 0.50 | 0.50 | 0.50 |
| **Sample BAF sds, mean** |  |  |  |  |  |  |  |
| All | Low | 0.03 | 0.03 | 0.03 | 0.03 | 0.03 | 0.03 |
|  | Med | 0.03 | 0.03 | 0.03 | 0.03 | 0.03 | 0.03 |
| **Sample BAF drifts, mean** |  |  |  |  |  |  |  |
| All | Low | 0 | 0 | 0 | 0 | 0 | 0 |
|  | Med | 0 | 0 | 0 | 0 | 0 | 0 |
| **Sample WF, mean** |  |  |  |  |  |  |  |
| All | Low | 0.01 | 0.01 | 0.009 | 0.01 | 0.01 | 0.02 |
|  | Med | 0.01 | 0.01 | 0.007 | 0.01 | 0.01 | 0.01 |
| **Callset PPV** |  |  |  |  |  |  |  |
| All | Low | 0.21 | 0.35 | 0.40 | 0.39 | 0.38 | 0.65 |
|  | Med | 0.75 | 0.81 | 0.84 | 0.84 | 0.82 | 0.82 |
| Deletions | Low | 0.19 | 0.31 | 0.36 | 0.36 | 0.35 | 0.52 |
|  | Med | 0.79 | 0.87 | 0.90 | 0.90 | 0.88 | 0.71 |
| Duplications | Low | 0.29 | 0.43 | 0.47 | 0.46 | 0.45 | 0.83 |
|  | Med | 0.71 | 0.76 | 0.79 | 0.79 | 0.76 | 0.91 |

Briefly, similarly to WAE, *Full Sets* in both the OEE and GSA array resulted in the most CNV calls (low-stringency QC 24,223 calls and 21,399 calls; medium-stringency QC 4,143 and 1,974 calls on GSA and OEE, respectively) and *Exact Match* the fewest (low-stringency QC on GSA array 2,529 and 1,883 calls; medium-stringency QC 1,258 and 936 calls on GSA and OEE, respectively). MarkerMatched callsets counted 2-6 times more CNV calls relative to *Exact Match* across all four *Methods* at *D_MAX_* = 10kb and both arrays (low-stringency QC range 10,491 - 14,024 calls; medium-stringency QC range 1,802 - 2,417 calls). While the average number of low-stringency QC calls overall was not substantially different between the OEE and GSA arrays (GSA overall callsets counted up to 1.3 times more CNVs), the differences between the two arrays observed in medium-stringency QC callsets were more variable and extreme (GSA overall callsets identified up to 2 times more CNVs).

CAE per-sample CNV calls, summarized in Tables 5 and 6, Supplemental Table S1L, were highest for the *Full Set* callset (low-stringency QC 36.3 and 32.1; medium stringency QC 6.2 and 3.0 for the GSA and OEE arrays, respectively), lowest for *Exact Match* (low-stringency QC 3.8 and 2.8; medium-stringency QC 1.9 and 1.4 for the GSA and OEE arrays, respectively), and about 2-6 times the *Exact Match* in MarkerMatched callsets across all *Methods* at *D_MAX_*= 10kb (low-stringency QC ranges of 20.9 - 21.0 and 15.7 - 17.7; medium-stringency QC ranges of 3.6 - 3.6 and 2.7 - 3.5 for GSA and OEE arrays, respectively).

In CAE, the average CNV sizes were smaller on the denser configurations (Tables 5 and 6, Supplemental Table S1L). The averages CNV sizes for GSA were smallest for MarkerMatch callsets (low-stringency QC range 44.6kb - 44.9kb; medium-stringency QC range 116.1kb - 118.0kb), followed by *Full Set* (low-stringency QC 58.5kb; medium-stringency QC 138.4kb). The average CNV sizes for OEE were smallest for *Full Set* (low-stringency QC 40.1kb; medium-stringency QC 112.1kb), followed by MarkerMatch callsets (low-stringency QC range 57.9kb - 62.7kb; medium-stringency QC range 115.0kb - 119.8b). *Exact Match* in both arrays had the largest average CNV sizes (low-stringency QC 92.1kb and 104.8kb; medium-stringency QC 134.2kb and 138.3kb for GSA and OEE arrays, respectively).

Average CNV confidence scores for the GSA array were about the same across all low-stringency QC configurations (Tables 5 and 6, Supplemental Table S1L). This included *Full Set*, *Exact Match*, and various *Method* configurations of MarkerMatched callsets (range 22.6 - 22.8), however, in the medium-stringency QC, MarkerMatched callsets had the highest CNV confidence scores (range 54.6 - 55.6), followed by *Full Set* (51.0) and *Exact Match* (33.8). For the OEE array, the spread was a bit wider among the low-stringency QC callsets (range 27.9 - 32.2), without clear segregation among the *Full Set* and MarkerMatch callsets, with *Exact Match* still being lowest. For medium-stringency QC callsets in the OEE array, we saw the highest average CNV confidence scores for *Full Set* (99.3), followed by MarkerMatch (range 70.0 - 77.6), and *Exact Match* (36.8).

Overall number of samples with at least one CNV call were consistent after low-stringency QC callsets across the board in both GSA and OEE array (N = 677, Tables 5 and 6, Supplemental Table S1M). The number of samples passing the medium-stringency QC varied somewhat across the callsets (N ranges 549 - 582 and 400 - 608, for GSA and OEE, respectively), with the OEE array showing a much higher range after medium-stringency QC.

Sample-specific metrics did not vary substantially across low-or medium-stringency QC, or *Full Set/Exact Match*/MarkerMatched configurations, or GSA/OEE arrays (Tables 5 and 6, Supplemental Table S1M).

### Genome-Wide Validation

CAE genome-wide validation of the GSA callsets (using low-stringency *Full Set* OEE as the truth set) resulted in PPVs that were the lowest for the *Full Set* (0.19), highest in *Exact Match* (0.56), and intermediate in the MarkerMatched (range 0.295 - 0.298) low-stringency QC callsets (Figure 5A, Table 5, Supplemental Table S1N). In medium-stringency QC GSA callsets, the performance was significantly lower in *Full Set* (0.48) than the rest of the callsets (range 0.72 - 0.74). CAE genome-wide validation of OEE callsets (using low-stringency *Full Set* GSA as truth set) has resulted in PPVs that were comparable to those in GSA callsets, however overall slightly larger, by an average factor of 1.2 (range 1.1 - 1.6), however *Methods* LRR mean and LRR sd had a slightly better performance than *Exact Match* in medium-stringency QC callsets (PPVs of 0.84, 0.84, and 0.82, respectively). Visual inspection of the genome-wide PPV plots (Figure 5) indicates that, in the majority of *Method* parameter configurations, MarkerMatch performed about as well as the *Exact Match* approach.

**Figure 5.**
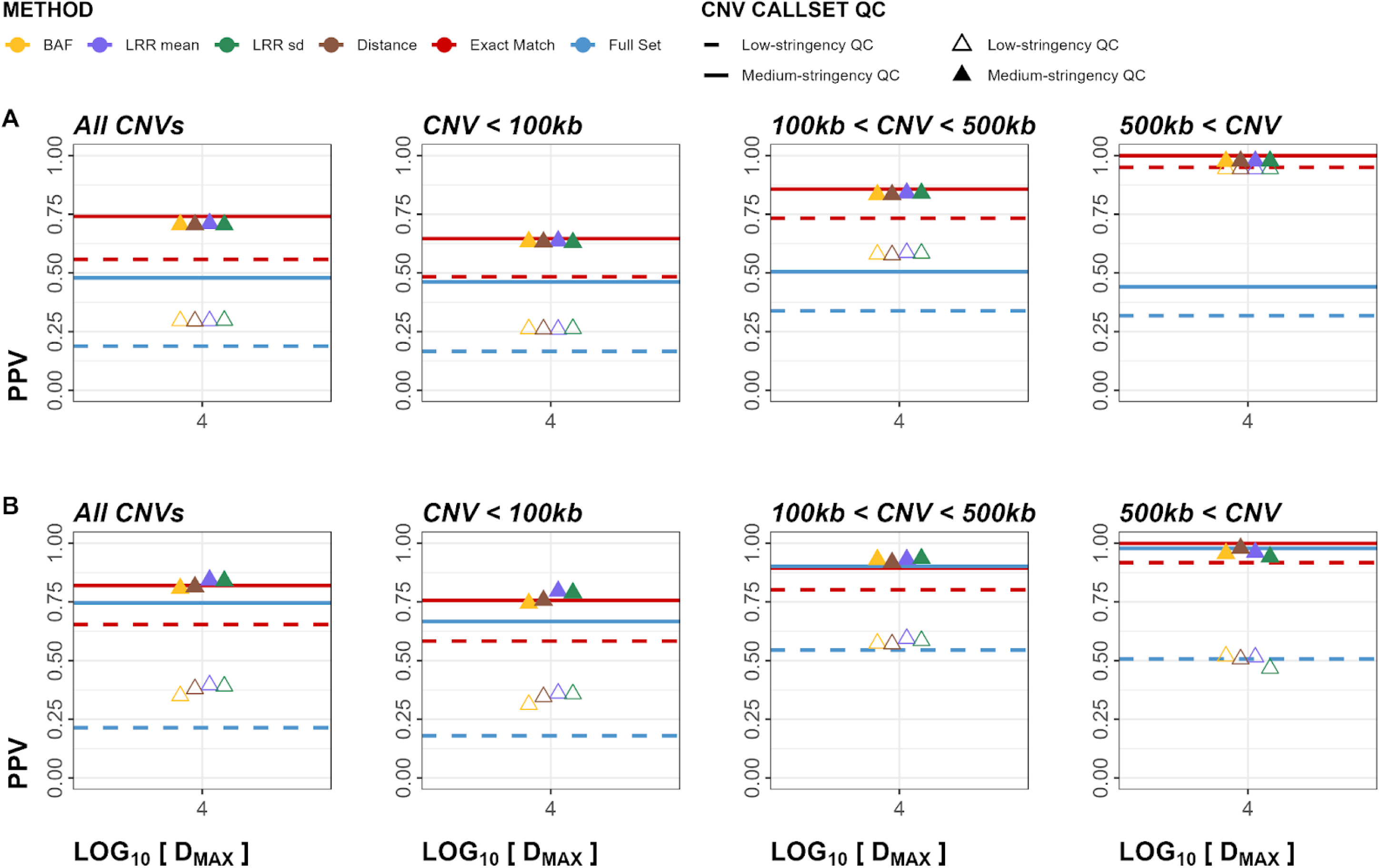
Cross-Array Experiment (CAE) positive predictive value (PPV) plots for both deletions and duplications. Panel A represents calls in Global Screening Array (GSA) validated in Omni Express Exome (OEE). Panel B represents calls in OEE validated in GSA. Empty triangles and dashed lines represent low-stringency QC callsets, whereas solid lines and filled triangles represent medium-stringency QC callsets.

Other metrics performed similarly, with larger CNVs having consistently better performance than smaller CNVs (Supplemental Figures S43-48, Supplemental Table S1N). Additionally, duplications had a noticeably better performance than deletions (Supplemental Figure S49-62, Supplemental Table S1N).

### Regional Validation

Filtering on medium-stringency QC indicated that *Exact Match* outperformed MarkerMatch in variable genomic regions including telomeric, centromeric, and segmental duplication regions (Supplemental Figures S63-69, Supplemental Table S1O) in terms of PPV. However, when accounting for the higher sensitivity of the MarkerMatch using the F1, FMI, and JI metrics, MarkerMatch either matched or slightly outperformed *Exact Match* across the board.

### Determination of Optimal Minimum CNV Size and SNP Coverage Thresholds

Beta regression of PPV on the GSA array resulted in significant associations with CNV length cutoff, marker coverage cutoff, and their interaction terms ([OR_PPV_ = 1.07, p_PPV_ < 0.001]; [OR_PPV_ = 1.02, p_PPV_ < 0.001]; [OR_PPV_ = 1.00, p_PPV_ < 0.001] respectively). In terms of PPV, LRR sd ([OR_PPV_ = 0.97, p_PPV_ = 0.009]) and BAF ([OR_PPV_ = 0.98, p_PPV_ = 0.03]) seemed to underperform LRR mean. The Distance method was not significantly different. The model’s pseudo R2 was 0.58.

In OEE, somewhat counterintuitively, the *marker coverage* cutoff seemed to actually decrease the PPV ([*OR_ppv_* = 0.98, *p_PPV_* < 0.001]) whereas the *CNV length cutoff* and the two terms’ interaction seemed to increase it ([*OR_PPV_* = 1.02, *p_PPV_* < 0.001]; [*OR_PPV_* = 1.00, *p_PPV_* < 0.001] respectively). Unlike with sensitivity, *Method* terms in PPV were significant in OEE array, with LRR sd overperforming ([*OR_PPV_* = 1.07, *p_PPV_* = 0.007]), and BAF underperforming ([*OR_PPV_* = 0.91, *p_PPV_*< 0.001]) relative to LRR mean. Distance was not significantly different from LRR mean. The model’s pseudo *R^2^* was 0.64.

Graphical representations are shown in Supplemental Figures S70-75.

### Sample-Wise Performance

The plots of performance by sample suggested that a substantial number of false positive calls were driven by poorly-performing samples (Figure 6). For example, considering *Exact Match*, 113 individuals with PPV < 0.1 accounted for 20.2% of false positive calls in the SSC dataset, 39 individuals with PPV < 0.1 accounted for 22.6% of false positive calls in GSA, and 29 individuals with PPV < 0.1 accounted for 32.5% of false positive calls in OEE (Table S2A, Supplemental Table S1P-Q). Because F1 scores are affected by the changes in sensitivity, and by the QC process in particular, we did not observe similar trends when examining F1 scores (Table S2B).

**Figure 6.**
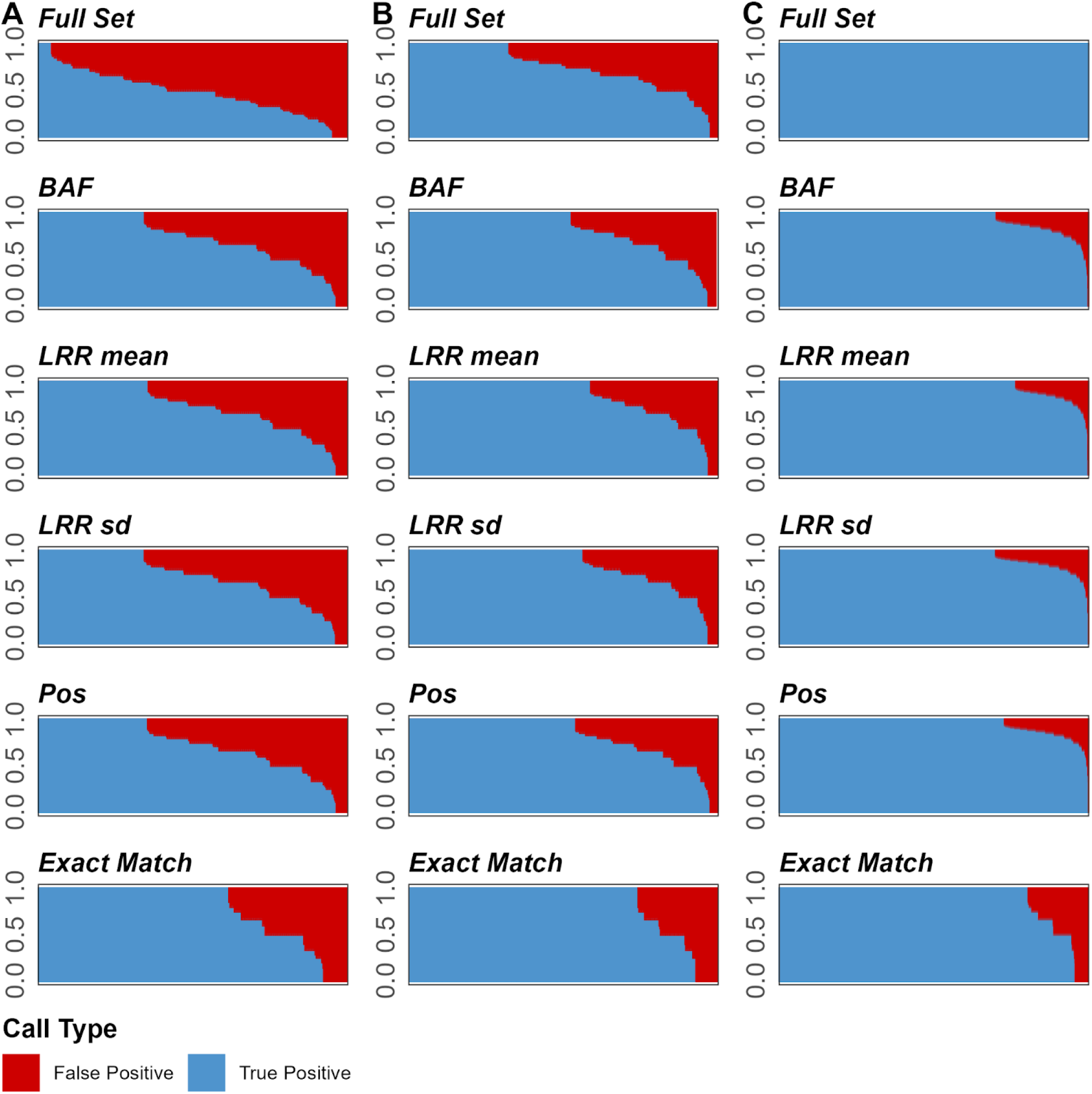
Figure showing proportions of true positive (blue) to false positive (red) CNV calls across medium-stringency QC callsets for Cross-Array Experiment (Global Screening Array and Omni Express Exome array on columns A and B, respectively) and Within-Array Experiment (Omni2.5 array on panel C). Y-axis shows positive predictive value (PPV), x-axis shows samples (ordered in descending PPV).

Plotting the curves along this sample-wise PPV thresholding indicated that, overall, conducting this CNV sample QC step may substantially improve callset PPV (Supplemental Figure S76), however, thresholds that are conservative may result in excessive drop-offs in F1 scores due to associated reductions in sensitivity (Supplemental Figure S77).

## Discussion

We describe a new approach to CNV pre-calling quality control to increase sensitivity in cross-array CNV studies. Instead of exclusively using consensus markers across all arrays considered (an intersection of common markers, or an approach that we call *Exact Match* in this study), we postulated that using markers in the same genomic neighborhood of the reference marker should result in an identical CNV state call. This ability to rely on similar genomic regions as opposed to identical markers would thus result in improved sensitivity of CNV calling by allowing higher coverage of available array markers (Figure 3).

Using simulated (within array experiment; WAE) data from the OMNI array, we show that using MarkerMatch not only substantially increased the sensitivity of CNV calling (4-fold increase in sensitivity from 0.031 in *Exact Match* to average sensitivity of 0.124 in MarkerMatch approach), it did so without a negative impact to PPV (0.900 in *Exact Match* compared to 0.943-958 in MarkerMatch) shown in Figure 4 and Table 4. Fluctuations in apparent performance are dependent on the number of CNVs in the truth set, for example, in specific subsets of CNV callsets such as CNV size cutoff > 5Mb only have between 2 and 25 CNV calls, therefore, a single false positive call may greatly affect observed PPV (Figure 4F). We identified a 10,000bp *D_MAX<_*parameter as a reasonable setting, and note that none of the specific *Method* parameters had a clear performance edge over the others. Noticeably, however, performance metrics that take into account both sensitivity and PPV, such as F1 score, Fowlkes-Mallows index, and Jaccard index, showed MarkerMatch substantially overperforming *Exact Match* (Supplemental Figures S9-11).

In the cross array experiment (CAE), we used TAAICG samples for which data were generated on two different arrays, and show that both *Exact Match* and MarkerMatch reduced some of the batch effects associated with the use of different arrays (PPV range of *Full Set* callset 0.479 vs. 0.705 - 0.741 in *Exact Match* and MarkerMatch callsets). CAE also demonstrated that MarkerMatch performed about as well as or better than *Exact Match* in terms of PPV (Figure 5, Tables 5 and 6). Similar to the WAE, performance metrics that take into account both sensitivity and PPV, such as F1, FMI, and JI, showed MarkerMatch outperforming *Exact Match* (Supplemental Figures S46-48). We also determined that increases in CNV length and marker coverage cutoffs drive improvements in PPV, but may cause reduction in overall sensitivity, as expected (Supplemental Figures S70-75).

We further inspected sample-wise performance rates to determine whether low PPVs were driven by individuals with low PPV (Figure 6), with individual samples with sample-wise PPV < 0.1 (ranging 0-14.4% of samples in a given callset) accounting for 20.2-32.5% of false positive CNV calls. We further examined whether eliminating these samples would substantially improve quality of the callsets, and found that while removing them may lead to noticeable improvements in PPV by a few percentage points, excessively conservative (high) thresholding hurt sensitivity and overall performance as measured by F1 scores (Supplemental Figures S76 and S77). Notably, excluding samples with low PPV values (< 0.1) does not decrease sensitivity or F1 scores.

While the MarkerMatch approach is successful in its primary function by increasing/rescuing sensitivity without reducing/sacrificing PPV, MarkerMatch does not eliminate batch effects attributable to the use of different arrays. We found some evidence of the potential reduction in batch and array effects in this study, but this needs to be further explored. Further research needs to be done to determine how significant array-specific batch effects really are, how much do *Exact Match* or MarkerMatch approaches really alleviate them, and what their quantifiable consequence to downstream CNV analyses might be. Thus, it is noteworthy that batch and array effects, albeit demonstrably reduced by MarkerMatch, still remain an important consideration in downstream CNV association analyses. Additionally, because we lacked access to adequate ancestrally diverse data, we did not examine the effects of ancestry composition on the MarkerMatch algorithm.

## Abbreviations

ASD: Autism spectrum disorder
BAF: B allele frequency
CNV: Copy-number variant
D_MAX: Maximum (allowable) distance
F1: F1 score
FDR: False discovery rate
FMI: Fowlkes–Mallows index
GSA: Global Screening Array
LRR: Log R ratio
MAD: Median absolute deviation
OEE: Omni Express Exome
OMNI: Omni 2.5
PPV: Positive predictive value
QC: Quality control
SSC: Simons Simplex Cohort
TAAICG: Tourette Association of America International Consortium for Genetics
TS: Tourette syndrome

## Acknowledgements

The authors would like to thank the Simons Foundation for Autism Research and the Tourette Association of America International Consortium for Genetics for contributing data for the analysis. The authors would also like to thank professor Lea Davis and Sharon Johnson for their feedback in the project.

## Funding

This work was supported by the National Institute for Neurological Disorders and Stroke [NS082168 to F.I.; NS105746 to R.O, L.D., J.S., and C.M.; NS102371 to R.O., L.D., J.S., and C.M.; MH115676 and MH125042 to R.O.; Tourette Association of America Young Investigator Award to T.M.F.].

## Notes

https://github.com/FranjoIM/MarkerMatch.

### Competing Interest Statement

Dr. Scharf is a member of the Tourette Association of America Scientific Advisory Board

