## Supplementary Information for "MarkerMatch: A Proximity-Based Probe-Matching Algorithm for Joint Analysis of Copy-Number Variants from Different Genotyping Arrays"

### S1 Methods

#### S1.1 Samples and Software

##### S1.1.1 Simons Simplex Collection (SSC)

The Simons Simplex Collection (SSC) was collected by the Simons Foundation Autism Research Initiative (SFARI) and contains over 2000 families in which probands have moderate to severe symptoms of autism spectrum disorder (ASD) and relatively little intellectual disability (Fischbach and Lord 2010). Data from 1,069 such families (N = 4,239) were generated using Illumina Infinium Omni2.5 (OMNI) assay with ~2,440,000 genome-wide probes (Sanders *et al.* 2015).

##### S1.1.2 Tourette Association of America International Consortium for Genetics (TAAICG)

The Tourette Association of America International Consortium for Genetics (TAAICG) is a collection of individuals with Tourette syndrome (TS) and their families, primarily from TS specialty clinics throughout North America, Europe, and Israel (Scharf *et al.* 2013). From this collection, we analyzed 3,920 individuals genotyped on Illumina Infinium OmniExpressExome (OEE) with ~960,000 probes (Huang *et al.* 2017) and 4,389 individuals genotyped on Illumina Infinium Global Screening Array v1 (GSA) with ~700,000 probes.

#### S1.2 Data Preprocessing

##### S1.2.1 Clustering and Data Export

Intensity files from genotyping microarrays were imported and clustered *de novo* in Illumina GenomeStudio software (Illumina 2020). Subsequently, we exported log-R ratio (LRR) and B-allele frequency (BAF) intensity data for all markers and all samples for downstream analysis into an Illumina final report format. We also exported the manifest containing probe name, chromosome, position, and probe BAF, and mean and standard deviation of LRR. The resulting files were passed into the MarkerMatch script for matching. The code available in the GitHub repository.

##### S1.2.2 Genotyping QC and Processing

In addition to processing data for CNV calling, genetic data were exported in Plink format from Illumina GenomeStudio for additional processing. The resulting Plink files were converted to *bfile* format. All datasets were processed through basic genotyping QC steps (Purcell *et al.* 2007; Marees *et al.* 2018). Namely, data were filtered to remove individuals with problematic sex information. For all datasets, only the autosomal SNPs were kept for further analysis. Serial, stepwise filtering was performed to retain only samples and SNPs with genotyping rate > 0.98. We then removed SNPs in Hardy-Weinberg disequilibrium by filtering out SNPs with

$P_{HWE} < 10^{-6}$. Lastly, we removed individuals with unusually high or low heterozygosity by filtering out individuals with $F_{HET}$ values outside of the three standard deviations of the cohorts' mean. Ultimately, we filtered out SNPs with MAF < 0.01. Only the samples passing this QC were retained in the validation study.

For the Cross-Array Experiment, we merged the individual batches into two TAAICG datasets: OEE and GSA (as determined by the array), each dataset filtered to retain only the intersection of SNPs from the respective array's batches. We then updated the datasets to only keep SNPs common to both OEE and GSA arrays, alleles of some SNPs in GSA were flipped to have the same allele configuration as those in OEE and rsIDs in GSA were updated to have the same rsID as those in OEE if the positions and alleles were matching. We then used the identity-by-descent approach to identify and remove potential cross-contamination and monozygotic twins in each dataset. Finally, we merged OEE and GSA into a single Plink file, and ran a single-line QC to remove samples and SNPs with genotyping rate < 0.98, MAF < 0.01, and $P_{HWE} < 10^{-6}$. The resulting dataset was processed through identity-by-descent to identify "monozygotic twins" who in our case actually represented the same individuals genotyped on two different arrays. We used the resulting ID matches to update IDs to be identical in both TAAICG/OEE and TAAICG/GSA CNV callsets.

#### S1.3 CNV Calling

Final reports exported from Illumina GenomeStudio were split into sample intensity files. We GC wave-adjusted the intensity files prior to CNV calling to reduce artifacts due to genomic waves in GC-content (Diskin *et al.* 2008). We then called CNVs using PennCNV as described in previous CNV studies (Huang *et al.* 2017).

CNVs were called individually, followed by merging of artificially broken up large CNVs. The resulting PennCNV callset was frozen as the low-stringency QC callset. The final dataset included 62 PennCNV callsets.

##### S1.3.1 CNV Quality Control

We applied the following QC metrics to derive the medium-stringency QC callset by removing samples with: (1) failing genotyping QC; (2) LRR mean outside of median +/- 4 median absolute deviations (MAD); (3) LRR standard deviation above median + 4 MAD; (4) BAF mean outside of median +/- 4 MAD; (5) BAF standard deviation above median + 4MAD; (6) BAF drift above median + 4 MAD; (7) GCWF outside median +/- 4 MAD; and (8) CNV count above callset median + 4 MAD.

We also removed CNVs based on the following criteria: (1) CNV spanning < 10 probes; and (2) CNV length < 20,000bp. CNV overlaps with telomeric, centromeric, immunoglobulin/T-cell receptor regions were removed, while segmental duplications were annotated for further examination.

CNVs were additionally annotated by the PennCNV confidence decile, thresholds for which were determined by the full set, low stringency QC, Within-Array Experiment CNV callset.

#### S1.4 CNV Call Validation

##### S1.4.1 Partial Confusion Matrix and Performance Metrics

In the Within-Array Experiment (WAE), we used the full OMNI manifest CNV callset as the ground truth to which we compared intersection of OMNI manifest with GSA manifest and different configurations of GSA-matched OMNI manifests. In the Cross-Array Experiment (CAE), we used the full GSA and OEE manifest CNV callsets as external and internal ground truths, respectively, to which we compared intersection of OEE manifest with GSA manifest and different configurations of GSA-matched OEE manifests.

To facilitate accurate measurement, Plink was used to identify samples which have been genotyped using both OEE and GSA arrays; the remaining samples were removed from the analysis. For the WAE, the low-stringency QC full set was considered a reference callset. For the CAE, the low-stringency QC OEE full set was considered a reference callset for GSA marker-matched callsets, and the low-stringency QC GSA full set was considered a reference callset for OEE marker-matched callsets. Supplemental **Figure 1B** shows the graphical representation of both validation studies.

True positives (TP) were defined as CNVs in test callsets that overlapped CNVs called in the *Full Set*, low-stringency QC callset. False positives (FP) were defined as CNVs in test callsets that did not overlap CNVs called in the *Full Set*, low-stringency QC callset. False negatives (FN) were defined as CNVs in the *Full Set*, low-stringency QC callset that did not overlap CNVs in test callsets. Here, we use test callsets to describe all callsets except the *Full Set*, low-stringency QC callset.

The following validation metrics were considered: true positive rate (sensitivity), false negative rate (FNR), positive predictive value (PPV; precision), false discovery rate (FDR), F1 score (F1; harmonic mean of precision and recall), Fowlkes–Mallows index (FMI; geometric mean of precision and recall); and Jaccard index (JI; ratio of the intersection to the union of the two sets).. To calculate these metrics, we used the equations **Eq. 1-7**.

In both WAE and CAE, we performed these analyses genome-wide and for CNVs overlapping centromeric, telomeric, and segmental duplications regions.

$$Sensitivity = TP / ( TP + FN ) = 1 - FNR \quad \text{(Eq. 1)}$$

$$FNR = FN / ( TP + FN ) = 1 - Sensitivity \quad \text{(Eq. 2)}$$

$$PPV = TP / ( TP + FP ) = 1 - FDR \quad \text{(Eq. 3)}$$

$$FDR = FP / ( TP + FP ) = 1 - PPV \quad \text{(Eq. 4)}$$

$$F1 = 2 * PPV * Sensitivity / ( PPV + Sensitivity ) \quad \text{(Eq. 5)}$$

$$FMI = sqrt( PPV * Sensitivity ) \quad \text{(Eq. 6)}$$

$$JI = TP / ( TP + FN + FP ) \quad \text{(Eq. 7)}$$

##### S1.4.2. Optimizing $D_{MAX}$ Parameter Selection

Only medium-stringency QC WAE callsets were used to optimize the $D_{MAX}$ parameter selection. In order to do that, we have first re-calibrated the performance metrics by scaling them to the respective medium-stringency QC *Full Set* callset values, following the equation **Eq. 8**, where $M_{CP}$ is a scaled performance metric, $M_{CO}$ is original performance metric for that (subset of) callset, and $M_{FS}$ is performance metric of (that subset's) *Full Set*. Only sensitivity, PPV, F1, FMI, and JI were included in this analysis. FNR and FDR were omitted as they are directly related to sensitivity and PPV, respectively.

$$M_{CP} = M_{CO} \ / \ M_{FS} \qquad \text{(Eq. 8)}$$

This was done to compensate for metric fluctuations of the *Full Set* performance metrics due to the quality control process. Once the metrics were scaled, they were re-plotted with *loess* smoothing function, for all CNV sizes except very large CNVs (< 1Mb), as there were too few to have a proper *loess* linear model fit. The smoothing was done to enable easier detection of major inflection points in validity metrics along the $D_{MAX}$ parameter values.

##### S1.4.3. Optimizing Method Parameter Selection

Only medium-stringency QC WAE and CAE callsets were used to optimize the *Method* parameter selection. In order to do that, we have first re-calibrated the performance metrics according to the method described above. Once the metrics were scaled, they were-replotted as *Method*-specific boxplots, stratified by CNV type and CNV size (except very large CNVs, < 1Mb). Box plots and medians were inspected visually to determine which *Methods* were performing better.

Following that, we performed CNV type and CNV size-stratified Welch two-sample t-test, assuming unequal variances between the scaled metric values (sensitivity', PPV', F1', FMI', and JI') between each combination of the four *Method* parameters, to evaluate whether the visual distribution shifts are statistically significant. We report both nominal *p* values, as well as FDR-adjusted *q* values.

##### S1.4.4. Optimizing Minimum CNV Length and Marker Coverage Thresholds

We also performed analysis to evaluate the optimal CNV length and marker coverage thresholds for the quality control purposes using the CAE callsets. While the performance metric data may contain 0 and 1 values, there is nothing intrinsically important about such extreme metric values to warrant zero/one inflated beta modeling approach. In other words, 1 and 0.9999 and 0 and 0.0001 are essentially the same for our modeling purposes. Thus, we transformed the metrics to fit (0, 1) distribution as opposed to [0, 1] distribution using the equation **Eq. 9** (Smithson and Verkuilen 2006). This will allow us to model metrics using regular beta distribution, as opposed to piecewise zero-one inflated beta modeling.

$$\textit{Metric (scaled)} = (\textit{Metric} * (N - 1) + 0.5) / N \qquad \text{(Eq. 9)}$$

The multivariable beta regression models were fit associating various scaled and transformed performance metric values to *CNV length cutoff* (varied from 20kb to 500kb in 20kb increments), *marker* *coverage cutoff* (varied from 10 to 50 in the increments of 5), and *Method* (measured relative to LRR mean for all MarkerMatch *Method* parameter values), according to the equation **Eq. 10**, across the medium-stringency QC callsets for both GSA and OEE arrays (where each array was analyzed independently). Notably, for the mean model, we included an interaction term between *CNV length cutoff* and *marker coverage cutoff*, while for the precision model we did not include any interaction terms.

$$\begin{gathered} \textit{Metric} \sim \textit{CNV length cutoff} * \textit{marker coverage cutoff} + \textit{Method} \mid \\ \textit{CNV length cutoff} + \textit{marker coverage cutoff} + \textit{Method} \end{gathered} \quad \text{(Eq. 10)}$$

In addition to regression models, we have plotted graphs showing scaled performance metrics relative to different combinations of *CNV length cutoff* and *marker coverage cutoff*, stratified by *Method*, $D_{MAX}$, and QC. Note units of *CNV length cutoff* are 10,000bp, e.g. value of 1=10,000bp, in the models. We also plotted performance metrics, specifically PPV and F1 score, at different *CNV length* and *marker coverage* cutoffs to see if most optimal cutoffs can be determined visually.

###### S1.4.5. Sample-Wise Performance

In order to determine whether there are samples driving high rates of false-positives, we plotted sample-wise stacked TP and FP rates for visual examination. We then re-calculated performance metrics to determine whether this as a QC step would lead to dramatic improvements in performance. This step was done on both WAE and CAE data.

### S2 Results

#### S2.1 Marker Matching

##### S2.1.1 Inter-Marker Gaps

Analysis of inter-marker gaps (i.e., gaps between SNPs used by PennCNV to make CNV calls) shows successful reduction of gap-sizes between SNPs on both OMNI and OEE when matched with GSA, relative to the typical *Exact Match* approach (**Supplemental Figure S1, Table 3, Supplemental Table S1C**). The reduction of inter-marker gaps in OMNI array when MarkerMatched to GSA at various *Method* and $D_{MAX}$ parameters appears to plateau at about $D_{MAX}$ = 10kb, where the median gaps were around 2.2kb, as compared to median gaps at *Exact Match* at about 11kb and *Full Set* at about 0.6kb (**Table 3, Supplemental Figure S1A**).

Similarly, when matching OEE to GSA, *Exact Match* resulted in gaps with a median of about 11kb, $D_{MAX}$ = 10kb MarkerMatch sets resulted in gaps with a median of about 2.4kb, and *Full Set* had gaps with a median of about 1.3kb on the OEE array (**Table 3, Supplemental Figure S1B**). *Exact Match* resulted in gaps with a median of about 11kb, $D_{MAX}$ = 10kb MarkerMatch sets resulted in gaps with a median of about 2.5kb, and *Full Set* had gaps with a median of about 2.2kb on the GSA array (**Table 3, Supplemental Figure S1C**).

###### S2.1.2 BAF

Analysis of BAFs shows that MarkerMatch approximates *Full Set* BAF distributions better than *Exact Match*. In the case of OMNI array, improvements in BAF plateaued at $D_{MAX}$ = 10kb for all *Method* settings (**Supplemental Figure S2, Table 3, Supplemental Table S1D**). Notably, the median BAF differed for each *Method*, with median BAF being 0.041 for *Method* = BAF, 0.080 for *Method* = LRR mean, 0.111 for *Method* = LRR sd, and 0.105 for *Method* = Distance arrays, whereas the median BAF values for *Exact Match* and *Full Set* were 0.257 and 0.074, respectively (**Table 3, Supplemental Figure S2A**). Notably, it appears that setting the *Method* parameter to BAF may result in a subset array with deflated BAF values.

When matching OEE to GSA, the median BAF on OEE array also differed for each *Method*, with median BAF being 0.133 for *Method* = BAF, 0.164 for *Method* = LRR mean, 0.169 for *Method* = LRR sd, and 0.172 for *Method* = Distance arrays, whereas the median BAF values for *Exact Match* and *Full Set* were 0.251 and 0.141, respectively (**Table 3, Supplemental Figure S2B**). The median BAF on the GSA array was identical for each *Method* with the value of 0.039, whereas the median BAF values for *Exact Match* and *Full Set* were 0.251 and 0.035, respectively (**Table 3, Supplemental Figure S2C**).

###### S2.1.3 LRR sd

Analysis of LRR sds has shown very little variability between the *Full Set*, *Exact Match*, and various configurations of the $D_{MAX}$ and *Method* parameters of MarkerMatched arrays in OMNI, OEE, and GSA datasets (**Supplemental Figure S3, Table 3, Supplemental Table S1E**), with median LRR sd values all being around 1.255.

###### S2.1.4 LRR mean

Analysis of LRR means has shown very little variability between the *Full Set*, *Exact Match*, and various configurations of the $D_{MAX}$ and *Method* parameters of MarkerMatched arrays in OMNI, OEE, and GSA datasets (**Supplemental Figures S4-5, Table 3, Supplemental Table S1F**), with median LRR mean values all being around -0.002.

#### S2.2 Within-Array Experiment

###### S2.2.1 CNV Call Rate Summaries

*Full Set*, low-stringency QC callset had a per-sample average of 68.1 (*st.dev* = 24.2) CNV calls. The CNV call rate for deletions was substantially higher than that for duplications, 50.4 ± 22.0 and 17.7 ± 8.8 respectively. The results for medium-stringency callset followed similar trends: 13.2 ± 5.9 (*Full Set* deletions and duplications), 7.5 ± 3.9 (*Full Set* deletions), 5.8 ± 3.6 (*Full Set* duplications).

*Exact Match*, low-stringency QC callset had a per-sample average of 2.6 (*st.dev* = 1.8) CNV calls. The CNV call rate for deletions was slightly lower than that for duplications, 1.2 ± 1.2 and 1.5 ± 1.4 respectively. The results for medium-stringency callset followed similar trends: 1.6 ± 1.5 (*Exact Match* deletions and duplications), 0.7 ± 0.9 (*Exact Match* deletions), 0.9 ± 1.1 (*Exact Match* duplications).

$D_{MAX}$ = 10kb, *Method* = BAF, low-stringency QC callset had a per-sample average of 18.5 (*st.dev* = 7.0). The CNV call rate for deletions was substantially higher than that for duplications, 11.6 ± 5.8 and 7.0 ± 4.0 respectively. The results for medium-stringency callset followed similar trends: 7.5 ± 3.8 ($D_{MAX}$ = 10kb, *Method* = BAF deletions and duplications), 4.1 ± 2.5 ($D_{MAX}$ = 10kb, *Method* = BAF deletions), 3.5 ± 2.4 ($D_{MAX}$ = 10kb, *Method* = BAF duplications).

$D_{MAX}$ = 10kb, *Method* = LRR mean, low-stringency QC callset had a per-sample average of 18.2 (*st.dev* = 7.9). The CNV call rate for deletions was substantially higher than that for duplications, 11.3 ± 6.9 and 6.9 ± 3.7 respectively. The results for medium-stringency callset followed similar trends: 7.4 ± 3.6 ($D_{MAX}$ = 10kb, *Method* = LRR mean deletions and duplications), 3.9 ± 2.4 ($D_{MAX}$ = 10kb, *Method* = LRR mean deletions), 3.5 ± 2.3 ($D_{MAX}$ = 10kb, *Method* = LRR mean duplications).

$D_{MAX}$ = 10kb, *Method* = LRR sd, low-stringency QC callset had a per-sample average of 19.5 (*st.dev* = 7.5). The CNV call rate for deletions was substantially higher than that for duplications, 11.7 ± 6.3 and 7.8 ± 3.8 respectively. The results for medium-stringency callset followed similar trends: 8.1 ± 3.8 ($D_{MAX}$ = 10kb, *Method* = LRR sd deletions and duplications), 4.3 ± 2.5 ($D_{MAX}$ = 10kb, *Method* = LRR sd deletions), 3.8 ± 2.4 ($D_{MAX}$ = 10kb, *Method* = LRR sd duplications).

$D_{MAX}$ = 10kb, *Method* = Distance, low-stringency QC callset had a per-sample average of 19.8 (*st.dev* = 7.5). The CNV call rate for deletions was substantially higher than that for duplications, 11.7 ± 6.3 and 8.1 ± 3.9 respectively. The results for medium-stringency callset followed similar trends: 8.1 ± 3.8 ($D_{MAX}$ = 10kb, *Method* = Distance deletions and duplications), 4.3 ± 2.5 ($D_{MAX}$ = 10kb, *Method* = Distance deletions), 3.8 ± 2.4 ($D_{MAX}$ = 10kb, *Method* = Distance duplications).

Detailed reports for other $D_{MAX}$ parameter settings (10bp, 50bp, 100bp, 500bp, 1kb, 5kb, 50kb, 100kb, 500kb, 1Mb, and 5Mb) are available in the **Supplemental Table S1G**.

##### S2.2.2 CNV Size Summaries

*Full Set*, low-stringency QC callset had an average CNV size of 25.4kb (*st.dev* = 96.9kb). The average sizes of deletions were substantially smaller than those of duplications, 18.1kb ± 75.5kb and 46.4kb ± 139.2kb respectively. The results for medium-stringency callset followed similar trends: 84.9kb ± 184.6kb (*Full Set* deletions and duplications), 71.9kb ± 176.4kb (*Full Set* deletions), 101.9kb ± 193.1kb (*Full Set* duplications).

*Exact Match*, low-stringency QC callset had an average CNV size of 116.7 kb (*st.dev* = 336.3kb). The average sizes of deletions were substantially smaller than those of duplications, 97.5kb ± 389.1kb and 132.4kb ± 285.3kb respectively. The results for medium-stringency callset followed similar trends: 148.8kb ± 399.5kb (*Exact Match* deletions and duplications), 131.8kb ± 484.1kb (*Exact Match* deletions), 160.7kb ± 326.9kb (*Exact Match* duplications).

$D_{MAX}$ = 10kb, *Method* = BAF, low-stringency QC callset had an average CNV size of 55.5kb (*st.dev* = 169.3kb). The average sizes of deletions were substantially smaller than those of duplications, 41.9kb ± 155.8kb and 77.9kb ± 187.4kb respectively. The results for medium-stringency callset followed similar trends:

103.7kb ± 239.5kb ($D_{MAX}$ = 10kb, *Method* = BAF deletions and duplications), 86.9kb ± 242.4kb ($D_{MAX}$ = 10kb, *Method* = BAF deletions), 123.4kb ± 234.5kb ($D_{MAX}$ = 10kb, *Method* = BAF duplications).

$D_{MAX}$ = 10kb, *Method* = LRR mean, low-stringency QC callset had an average CNV size of 55.3kb (*st.dev* = 172.2kb). The average sizes of deletions were substantially smaller than those of duplications, 40.8kb ± 157.7kb and 78.9kb ± 191.1kb respectively. The results for medium-stringency callset followed similar trends: 105.7kb ± 243.7kb ($D_{MAX}$ = 10kb, *Method* = LRR mean deletions and duplications), 86.7kb ± 247.2kb ($D_{MAX}$ = 10kb, *Method* = LRR mean deletions), 127.2kb ± 237.8kb ($D_{MAX}$ = 10kb, *Method* = LRR mean duplications).

$D_{MAX}$ = 10kb, *Method* = LRR sd, low-stringency QC callset had an average CNV size of 54.8kb (*st.dev* = 168.2kb). The average sizes of deletions were substantially smaller than those of duplications, 41.6kb ± 155.3kb and 74.4kb ± 184.1kb respectively. The results for medium-stringency callset followed similar trends: 102.5kb ± 236.9kb ($D_{MAX}$ = 10kb, *Method* = LRR sd deletions and duplications), 85.6kb ± 243.8kb ($D_{MAX}$ = 10kb, *Method* = LRR sd deletions), 121.2kb ± 227.6kb ($D_{MAX}$ = 10kb, *Method* = LRR sd duplications).

$D_{MAX}$ = 10kb, *Method* = Distance, low-stringency QC callset had an average CNV size of 54.5kb (*st.dev* = 166.7kb). The average sizes of deletions were substantially smaller than those of duplications, 42.3kb ± 155.4kb and 72kb ± 180.5kb respectively. The results for medium-stringency callset followed similar trends: 103.3kb ± 231.5kb ($D_{MAX}$ = 10kb, *Method* = Distance deletions and duplications), 87.3kb ± 247.3kb ($D_{MAX}$ = 10kb, *Method* = Distance deletions), 121kb ± 211.1kb ($D_{MAX}$ = 10kb, *Method* = Distance duplications).

Detailed reports for other $D_{MAX}$ parameter settings (10bp, 50bp, 100bp, 500bp, 1kb, 5kb, 50kb, 100kb, 500kb, 1Mb, and 5Mb) are available in the **Supplemental Table S1G**.

##### S2.2.3 CNV Confidence Score Summaries

*Full Set*, low-stringency QC callset had an average CNV confidence score of 46.4 (*st.dev* = 168.9). The average confidence scores were roughly similar for deletions and duplications, 43.4 ± 174.6 and 54.9 ± 151.1 respectively. The results for medium-stringency callset followed similar trends, except with overall higher confidence scores: 122.6 ± 347.5 (*Full Set* deletions and duplications), 127.6 ± 410.4 (*Full Set* deletions), 116.2 ± 242.4 (*Full Set* duplications).

*Exact Match*, low-stringency QC callset had an average CNV confidence score of 31.7 (*st.dev* = 70.5). The average confidence scores were roughly similar for deletions and duplications, 30.0 ± 86.6 and 33.1 ± 53.9 respectively. The results for medium-stringency callset followed similar trends, except with overall higher confidence scores: 38.5 ± 85.4 (*Exact Match* deletions and duplications), 37.6 ± 110.2 (*Exact Match* deletions), 39.2 ± 62.4 (*Exact Match* duplications).

$D_{MAX}$ = 10kb, *Method* = BAF, low-stringency QC callset had an average CNV confidence score of 32.3 (*st.dev* = 90.2). The average confidence scores were roughly similar for deletions and duplications, 31.8 ± 102.8 and 33.2 ± 64.2 respectively. The results for medium-stringency callset followed similar trends, except with overall higher confidence scores: 50.6 ± 124.4 ($D_{MAX}$ = 10kb, *Method* = BAF deletions and duplications), 52.1 ± 149.9 ($D_{MAX}$ = 10kb, *Method* = BAF deletions), 48.8 ± 85.0 ($D_{MAX}$ = 10kb, *Method* = BAF duplications).

$D_{MAX}$ = 10kb, *Method* = LRR mean, low-stringency QC callset had an average CNV confidence score of 35.4 (*st.dev* = 105.1). The average confidence scores were roughly similar for deletions and duplications, 33.1 ± 119.8 and 38.2 ± 75.2 respectively. The results for medium-stringency callset followed similar trends, except with overall higher confidence scores: 57.1 ± 148.2 ($D_{MAX}$ = 10kb, *Method* = LRR mean deletions and duplications), 56.2 ± 181.3 ($D_{MAX}$ = 10kb, *Method* = LRR mean deletions), 58.3 ± 98.3 ($D_{MAX}$ = 10kb, *Method* = LRR mean duplications).

$D_{MAX}$ = 10kb, *Method* = LRR sd, low-stringency QC callset had an average CNV confidence score of 34.4 (*st.dev* = 102.9). The average confidence scores were roughly similar for deletions and duplications, 32.5 ± 117.7 and 37.1 ± 75.5 respectively. The results for medium-stringency callset followed similar trends, except with overall higher confidence scores: 55.6 ± 153.0 ($D_{MAX}$ = 10kb, *Method* = LRR sd deletions and duplications), 54.0 ± 188.4 ($D_{MAX}$ = 10kb, *Method* = LRR sd deletions), 57.3 ± 100.4 ($D_{MAX}$ = 10kb, *Method* = LRR sd duplications).

$D_{MAX}$ = 10kb, *Method* = Distance, low-stringency QC callset had an average CNV confidence score of 33.8 (*st.dev* = 94.5). The average confidence scores were roughly similar for deletions and duplications, 32.1 ± 107.5 and 36.1 ± 71.7 respectively. The results for medium-stringency callset followed similar trends, except with overall higher confidence scores: 54.8 ± 143.3 ($D_{MAX}$ = 10kb, *Method* = Distance deletions and duplications), 53.4 ± 174.9 ($D_{MAX}$ = 10kb, *Method* = Distance deletions), 56.3 ± 96.9 ($D_{MAX}$ = 10kb, *Method* = Distance duplications).

Detailed reports for other $D_{MAX}$ parameter settings (10bp, 50bp, 100bp, 500bp, 1kb, 5kb, 50kb, 100kb, 500kb, 1Mb, and 5Mb) are available in the **Supplemental Table S1G**.

##### S2.2.4 Sensitivity Analyses

The overall sensitivity of the *Exact Match* approach was 0.037 and 0.031, for low- and medium-stringency QC callsets, respectively. Sensitivity was higher for larger CNVs, with the medium-stringency QC sensitivities of 0.019 (under 100kb), 0.275 (100kb to 500kb), 0.538 (500kb to 1Mb), 0.685 (1Mb to 5Mb), and 0.909 (over 5Mb).

The MarkerMatch algorithm shows sensitivity to be similar across the different *Method* parameters. The sensitivity seems to grow before plateauing at about $D_{MAX}$ = 10kb across all size bins, at about 0.124 (all sizes), 0.091 (under 100kb), 0.725 (100kb to 500kb), 0.725 (500kb to 1Mb), 0.820 (1Mb to 5Mb), and 1.000 (over 5Mb), averaged across all four *Method* parameters, at medium-stringency QC.

*Full Set*-specific analysis indicates a significant potential drop in sensitivity due to quality control process (medium-stringency QC callset shows over 75% reduction in sensitivity, at 0.247), however this effect seems to taper off among larger CNVs, with size-bin specific sensitivity reductions of about 19.5% (under 100kb), 8.5% (100kb to 500kb), 2.8% (500kb to 1Mb), 16.2% (1Mb to 5Mb), and 0% (over 5Mb).

Detailed reports for other $D_{MAX}$ parameter settings (10bp, 50bp, 100bp, 500bp, 1kb, 5kb, 50kb, 100kb, 500kb, 1Mb, and 5Mb) are available in the **Supplemental Table S1I**. Sensitivity WAE plots are available in

**Supplemental Figure S6** for all CNV types, as well as **Supplemental Figures S12** and **S19** for deletions and duplications, respectively.

##### S2.2.5 Positive Predictive Value Analyses

The overall positive predictive value (PPV) of the *Exact Match* approach was 0.824 and 0.900, for low- and medium-stringency QC callsets, respectively. PPV was higher for larger CNVs, with the medium-stringency QC PPVs of 0.858 (under 100kb), 0.961 (100kb to 500kb), 1.000 (500kb to 1Mb), 1.000 (1Mb to 5Mb), and 1.000 (over 5Mb).

The MarkerMatch algorithm shows PPV to be similar across the different *Method* parameters. The PPV seems to grow before plateauing at about $D_{MAX}$ = 10kb across all size bins, at about 0.952 (all sizes), 0.935 (under 100kb), 0.992 (100kb to 500kb), 0.995 (500kb to 1Mb), 1.000 (1Mb to 5Mb), and 1.000 (over 5Mb), averaged across all four *Method* parameters, at medium-stringency QC. At $D_{MAX}$ = 10kb, the overall all performance was the best for *Method* = LRR mean ($PPV_{LRR\ mean,\ 19kb}$ = 0.958), followed by *Method* = Distance ($PPV_{Distance,\ 19kb}$ = 0.956), *Method* = LRR sd ($PPV_{LRR\ sd,\ 19kb}$ = 0.950), and finally *Method* = BAF ($PPV_{BAF,\ 19kb}$ = 0.943).

The highest PPV of 0.966 in MarkerMatch WAE callsets was observed at *Method* = LRR mean and $D_{MAX}$ = 5kb across all CNV types and sizes, in the medium-stringency QC callset.PPVs generally appeared to peak at around $D_{MAX}$ = 5-10kb, followed by plateau (in case of Distance and BAF *Methods*) or rapidly decrease (in case of LRR mean and  LRR sd *Methods*). PPVs for CNVs over the size of 500kb, however, remained constant.

Detailed reports for other $D_{MAX}$ parameter settings (10bp, 50bp, 100bp, 500bp, 1kb, 5kb, 50kb, 100kb, 500kb, 1Mb, and 5Mb) are available in the **Supplemental Table S1I**. PPV WAE plots are available in **Figure 4** for all CNV types, as well as **Supplemental Figures S13** and **S20** for deletions and duplications, respectively.

##### S2.2.6 False Negative Rate Analyses

Since the false negative rate (FNR) is a complement to sensitivity, as per equations **Eq. 1** and **Eq. 3**. Overall, FNRs appear to be highest for the *Exact Match*, at 0.963 and 0.969, for low- and medium-stringency QC callsets, respectively. FNR tends to drop with larger-size CNV bins.

A detailed FNR report is available in the **Supplemental Table S1I**. FNR WAE plots are available in **Supplemental Figure S7** for all CNV types, as well as **Supplemental Figures S14** and **S21** for deletions and duplications, respectively.

##### S2.2.7 False Discovery Rate Analyses

Since the false discovery rate (FDR) is a complement to PPV, as per equations **Eq. 2** and **Eq. 4**. Similarly, but inversely, FDRs were lower for MarkerMatch callsets than *Exact Match* for all configurations of *Method* parameters and $D_{MAX}$ < 100kb.

A detailed FDR report is available in the **Supplemental Table S1I**. FDR WAE plots are available in **Supplemental Figure S8** for all CNV types, as well as **Supplemental Figures S15** and **S22** for deletions and duplications, respectively.

##### S2.2.8 F1 Score Analyses

F1 score for MarkerMatch continuously outperformed the *Exact Match* callset, when compared to the *Full Set* callset, with an exception of large CNV subsets of $D_{MAX}$ < 100bp callsets. Similarly to other metrics, F1 scores plateau around $D_{MAX}$ = 10kb and appear to be consistent across different *Method* parameters.

The overall F1 score (F1) of the *Exact Match* approach was 0.071 and 0.060, for low- and medium-stringency QC callsets, respectively. The F1 seems to grow before plateauing at about $D_{MAX}$ = 10kb across all size bins, at about 0.220 (all sizes), 0.165 (under 100kb), 0.838 (100kb to 500kb), 0.964 (500kb to 1Mb), 0.901 (1Mb to 5Mb), and 1.000 (over 5Mb), averaged across all four *Method* parameters, at medium-stringency QC.

A detailed F1 report is available in the **Supplemental Table S1I**. F1 WAE plots are available in **Supplemental Figure S9** for all CNV types, as well as **Supplemental Figures S16** and **S23** for deletions and duplications, respectively.

##### S2.2.9 Fowlkes-Mallows Index Analyses

Similarly to F1, the Fowlkes-Mallows index (FMI) continuously outperformed the *Exact Match* callset, when compared to the *Full Set* callset, with an exception of large CNV subsets of $D_{MAX}$ < 100bp callsets. Similarly to other metrics, FMIs plateau around $D_{MAX}$ = 10kb and appear to be consistent across different *Method* parameters.

The overall FMIs of the *Exact Match* approach were 0.174 and 0.167, for low- and medium-stringency QC callsets, respectively. The FMI seems to grow before plateauing at about $D_{MAX}$ = 10kb across all size bins, at about 0.344 (all sizes), 0.291 (under 100kb), 0.848 (100kb to 500kb), 0.965 (500kb to 1Mb), 0.906 (1Mb to 5Mb), and 1.000 (over 5Mb), averaged across all four *Method* parameters, at medium-stringency QC.

A detailed FMI report is available in the **Supplemental Table S1I**. FMI WAE plots are available in **Supplemental Figure S10** for all CNV types, as well as **Supplemental Figures S17** and **S24** for deletions and duplications, respectively.

##### S2.2.10 Jaccard Index Analyses

Similarly to both FMI and F1, and sensitivity, the Jaccard index (JI) continuously outperformed the *Exact Match* callset, when compared to the *Full Set* callset, with an exception of large CNV subsets of $D_{MAX}$ < 100bp callsets. Similarly to other metrics, JIs plateau around $D_{MAX}$ = 10kb and appear to be consistent across different *Method* parameters.

The overall JIs of the *Exact Match* approach were 0.037 and 0.031, for low- and medium-stringency QC callsets, respectively. The JI seems to grow before plateauing at about $D_{MAX}$ = 10kb across all size bins, at about 0.124 (all sizes), 0.090 (under 100kb), 0.721 (100kb to 500kb), 0.932 (500kb to 1Mb), 0.820 (1Mb to 5Mb), and 1.000 (over 5Mb), averaged across all four *Method* parameters, at medium-stringency QC.

A detailed JI report is available in the **Supplemental Table S1I**. JIWAE plots are available in **Supplemental Figure S11** for all CNV types, as well as **Supplemental Figures S18** and **S25** for deletions and duplications, respectively.

###### S2.2.11 Deletions vs. Duplications

Deletions and duplications followed roughly the same trend when it came to validation metrics, with some exceptions. Namely, in *Exact Match*, in the low-stringency QC callset, duplications had about 3.7-fold higher sensitivity and JI, about 3.5-fold higher F1, and about 2.0-fold higher FMI. In the medium-stringency QC callset, the difference was more extreme, with duplications having 4.1-fold higher sensitivity and JI, about 3.8-fold higher F1, and about 2.1-fold higher FMI.

The similar was the case with MarkerMatch callsets for all *Method* parameters and $D_{MAX}$ = 10kb, specifically, values for sensitivity, FDR, F1, FMI, and JI, which were between 1.5 and 2.4 times higher for deletions than duplications, on average. Values for PPV and FNR were roughly the same for deletions and duplications.

Detailed reports for other $D_{MAX}$ parameter settings (10bp, 50bp, 100bp, 500bp, 1kb, 5kb, 50kb, 100kb, 500kb, 1Mb, and 5Mb) are available in the **Supplemental Table S1I**. These data have been graphically shown for deletions and duplications in the **Supplemental Figures S12-25**.

##### S2.3 Cross-Array Experiment

###### S2.3.1 CNV Call Rate Summaries

*Full Set* low-stringency QC callset from the GSA array had a per-sample average of 36.3 (*st.dev* = 31.7) CNV calls. The CNV call rate for deletions was about the same as that for duplications, 19.6 ± 26.9 and 16.7 ± 19.0, respectively. The results for medium-stringency QC callset followed similar trends: 6.2 ± 4.4 (*Full Set* deletions and duplications), 2.5 ± 2.7 (*Full Set* deletions), 3.7 ± 3.2 (*Full Set* duplications). The low-stringency QC callset from the OEE array had a per-sample average of 32.1 (*st.dev* = 61.6) CNV calls. The CNV call rate for deletions was noticeably higher than that for duplications, 23.4 ± 59.7 and 8.7 ± 8.6, respectively. The results for the medium-stringency QC callset followed somewhat different trends, with deletions and duplications being more similar: 3.0 ± 3.0 (*Full Set* deletions and duplications), 1.5 ± 1.7 (*Full Set* deletions), 1.5 ± 1.8 (*Full Set* duplications).

*Exact Match* low-stringency QC callset from the GSA array had a per-sample average of 3.8 (*st.dev* = 2.7) CNV calls. The CNV call rate for deletions was about the same as that for duplications, 1.9 ± 2.3 and 1.9 ± 1.7, respectively. The results for medium-stringency QC callset followed similar trends: 1.9 ± 1.6 (*Exact Match* deletions and duplications), 0.8 ± 1.1 (*Exact Match* deletions), 1.1 ± 1.2 (*Exact Match* duplications). The low-stringency QC callset from the OEE array had a per-sample average of 2.8 (st.dev = 2.3) CNV calls. The CNV call rate for deletions was somewhat higher than that for duplications, 1.6 ± 2.1 and 1.2 ± 1.1, respectively. The results for the medium-stringency QC callset followed somewhat different trends, with deletions and duplications being more similar: 1.4 ± 1.4 (*Exact Match* deletions and duplications), 0.6 ± 0.9 (*Exact Match* deletions), 0.8 ± 0.9 (*Exact Match* duplications).

$D_{MAX}$ = 10kb, *Method* = BAF, low-stringency QC callset from the GSA array had a per-sample average of 21.0 (*st.dev* = 25.9) CNV calls. The CNV call rate for deletions was somewhat higher than that for duplications, 12.1 ± 22.3 and 8.9 ± 14.4, respectively. The results for medium-stringency QC callset followed somewhat different trends, with deletions and duplications being more similar: 3.6 ± 2.8 ($D_{MAX}$ = 10kb, *Method* = BAF, deletions and duplications), 1.8 ± 2.0 ($D_{MAX}$ = 10kb, *Method* = BAF, deletions), 1.8 ± 1.9 ($D_{MAX}$ = 10kb, *Method* = BAF, duplications). The low-stringency QC callset from the OEE array had a per-sample average of 17.7 (st.dev = 29.6) CNV calls. The CNV call rate for deletions was noticeably higher than that for duplications, 12.4 ± 29.0 and 5.3 ± 4.3, respectively. The results for the medium-stringency QC callset followed somewhat different trends, with deletions and duplications being more similar: 2.7 ± 2.7 ($D_{MAX}$ = 10kb, *Method* = BAF, deletions and duplications), 1.3 ± 1.6 ($D_{MAX}$ = 10kb, *Method* = BAF, deletions), 1.4 ± 1.7 ($D_{MAX}$ = 10kb, *Method* = BAF, duplications).

$D_{MAX}$ = 10kb, *Method* = LRR mean, low-stringency QC callset from the GSA array had a per-sample average of 21.0 (*st.dev* = 25.6) CNV calls. The CNV call rate for deletions was somewhat higher than that for duplications, 12.0 ± 22.0 and 9.0 ± 14.3, respectively. The results for medium-stringency QC callset followed somewhat different trends, with deletions and duplications being more similar: 3.5 ± 2.8 ($D_{MAX}$ = 10kb, *Method* = LRR mean, deletions and duplications), 1.7 ± 1.9 ($D_{MAX}$ = 10kb, *Method* = LRR mean, deletions), 1.8 ± 1.9 ($D_{MAX}$ = 10kb, *Method* = LRR mean, duplications). The low-stringency QC callset from the OEE array had a per-sample average of 15.7 (*st.dev* = 25.6) CNV calls. The CNV call rate for deletions was noticeably higher than that for duplications, 10.7 ± 25.1 and 5.0 ± 3.6, respectively. The results for the medium-stringency QC callset followed somewhat different trends, with deletions and duplications being more similar: 3.3 ± 2.2 ($D_{MAX}$ = 10kb, *Method* = LRR mean, deletions and duplications), 1.6 ± 1.4 ($D_{MAX}$ = 10kb, *Method* = LRR mean, deletions), 1.7 ± 1.5 ($D_{MAX}$ = 10kb, *Method* = LRR mean, duplications).

$D_{MAX}$ = 10kb, *Method* = LRR sd, low-stringency QC callset from the GSA array had a per-sample average of 20.9 (*st.dev* = 25.6) CNV calls. The CNV call rate for deletions was somewhat higher than that for duplications, 12.0 ± 22.0 and 8.9 ± 14.3, respectively. The results for medium-stringency QC callset followed somewhat different trends, with deletions and duplications being more similar: 3.6 ± 2.8 ($D_{MAX}$ = 10kb, *Method* = LRR sd, deletions and duplications), 1.7 ± 1.9 ($D_{MAX}$ = 10kb, *Method* = LRR sd, deletions), 1.8 ± 1.9 ($D_{MAX}$ = 10kb, *Method* = LRR sd, duplications). The low-stringency QC callset from the OEE array had a per-sample average of 16.2 (*st.dev* = 27.6) CNV calls. The CNV call rate for deletions was somewhat higher than that for duplications, 11.0 ± 27.2 and 5.2 ± 3.6, respectively. The results for the Medium-stringency QC callset followed somewhat different trends, with deletions and duplications being more similar: 3.5 ± 2.3 ($D_{MAX}$ = 10kb, *Method* = LRR sd, deletions and duplications), 1.6 ± 1.4 ($D_{MAX}$ = 10kb, *Method* = LRR sd, deletions), 1.8 ± 1.6 ($D_{MAX}$ = 10kb, *Method* = LRR sd, duplications).

$D_{MAX}$ = 10kb, *Method* = Distance, low-stringency QC callset from the GSA array had a per-sample average of 21.0 (*st.dev* = 25.8) CNV calls. The CNV call rate for deletions was substantially higher than that for duplications, 12.1 ± 22.1 and 9.0 ± 14.5, respectively. The results for medium-stringency QC callset followed somewhat different trends, with deletions and duplications being more similar: 3.6 ± 2.9 ($D_{MAX}$ = 10kb, *Method* = Distance, deletions and duplications), 1.8 ± 2.1 ($D_{MAX}$ = 10kb, *Method* = Distance, deletions), 1.8 ± 1.9 ($D_{MAX}$ = 10kb, *Method* = Distance, duplications). The low-stringency QC callset from the OEE array had a per-sample average of 16.5 (st.dev = 27.5) CNV calls. The CNV call rate for deletions was substantially higher than that for duplications, 11.3 ± 27.1 and 5.2 ± 3.8, respectively. The results for the medium-stringency QC callset followed somewhat different trends, with deletions and duplications being more similar: 3.0 ± 2.5 ($D_{MAX}$ = 10kb, *Method* = Distance, deletions and duplications), 1.4 ± 1.5 ($D_{MAX}$ = 10kb, *Method* = Distance, deletions), 1.6 ± 1.7 ($D_{MAX}$ = 10kb, *Method* = Distance, duplications).

Detailed reports for size-stratified bins are available in the **Supplemental Table S1L**.

##### S2.3.2 CNV Size Summaries

*Full Set* low-stringency QC callset from the GSA array had an average CNV size of 58.5kb (*st.dev* = 404kb). The average sizes of deletions were substantially smaller than those for duplications, 38kb ± 243.2kb and 82.6kb ± 533.4kb, respectively. The results for medium-stringency QC callset followed similar trends: 138.4kb ± 439.1kb (*Full Set* deletions and duplications), 124.1kb ± 658.0kb (*Full Set* deletions), 148.1kb ± 173.1kb (*Full Set* duplications). The low-stringency QC callset from the OEE array had an average CNV size of 40.1kb (*st.dev* = 139.1kb). The average sizes of deletions were substantially smaller than those for duplications, 32.1kb ± 89.3kb and 61.8kb ± 222.5kb, respectively. The results for the medium-stringency QC callset followed similar trends: 112.1kb ± 157.7kb (*Full Set* deletions and duplications), 90.4kb ± 126.7kb (*Full Set* deletions), 134.2kb ± 181.3kb (*Full Set* duplications).

*Exact Match* low-stringency QC callset from the GSA array had an average CNV size of 92.1kb (*st.dev* = 167.6kb). The average sizes of deletions were substantially smaller than those for duplications, 81.8kb ± 164.5kb and 102.3kb ± 170.2kb, respectively. The results for medium-stringency QC callset followed similar trends: 134.2kb ± 216.8kb (*Exact Match* deletions and duplications), 121.7kb ± 229kb (*Exact Match* deletions), 143.4kb ± 207.1kb (*Exact Match* duplications). The low-stringency QC callset from the OEE array had an average CNV size of 104.8kb (*st.dev* = 190.6kb). The average sizes of deletions were substantially smaller than those for duplications, 84.9kb ± 179.6kb and 130.9kb ± 201.2kb, respectively. The results for the medium-stringency QC callset followed similar trends: 138.3kb ± 220.6kb (*Exact Match* deletions and duplications), 121.5kb ± 249.7kb (*Exact Match* deletions), 152kb ± 192.9kb (*Exact Match* duplications).

$D_{MAX}$ = 10kb, *Method* = BAF, low-stringency QC callset from the GSA array had an average CNV size of 44.9kb (*st.dev* = 94.2kb). The average sizes of deletions were somewhat smaller than those for duplications, 38.1kb ± 81.7kb and 54.2kb ± 108.2kb, respectively. The results for medium-stringency QC callset followed similar trends: 116.1kb ± 163.1kb ($D_{MAX}$ = 10kb, *Method* = BAF, deletions and duplications), 101kb ± 138.8kb ($D_{MAX}$ = 10kb, *Method* = BAF, deletions), 130.7kb ± 182.4kb ($D_{MAX}$ = 10kb, *Method* = BAF, duplications). The low-stringency QC callset from the OEE array had an average CNV size of 57.9kb (*st.dev* = 179.5kb). The average sizes of deletions were substantially smaller than those for duplications, 46kb ± 110.8kb and 85.4kb ± 277.9kb, respectively. The results for the medium-stringency QC callset followed similar trends: 115kb ± 176.5kb ($D_{MAX}$ = 10kb, *Method* = BAF, deletions and duplications), 95.5kb ± 133.7kb ($D_{MAX}$ = 10kb, *Method* = BAF, deletions), 133.6kb ± 208.0kb ($D_{MAX}$ = 10kb, *Method* = BAF, duplications).

$D_{MAX}$ = 10kb, *Method* = LRR mean, low-stringency QC callset from the GSA array had an average CNV size of 44.6kb (*st.dev* = 93.9kb). The average sizes of deletions were somewhat smaller than those for duplications, 37.9kb ± 81.8kb and 53.6kb ± 107.5kb, respectively. The results for medium-stringency QC callset followed similar trends: 118.0kb ± 179.4kb ($D_{MAX}$ = 10kb, *Method* = LRR mean, deletions and duplications), 104.3kb ± 176.0kb ($D_{MAX}$ = 10kb, *Method* = LRR mean, deletions), 131.0kb ± 181.7kb ($D_{MAX}$ = 10kb, *Method* = LRR mean, duplications). The low-stringency QC callset from the OEE array had an average CNV size of 62.7kb (*st.dev* = 186.3kb). The average sizes of deletions were substantially smaller than those for duplications, 49.3kb ± 114.3kb and 91.5kb ± 282.7kb, respectively. The results for the medium-stringency QC callset followed similar trends: 118.7kb ± 185.5kb ($D_{MAX}$ = 10kb, *Method* = LRR mean, deletions and duplications), 100.0kb ± 169.3kb ($D_{MAX}$ = 10kb, *Method* = LRR mean, deletions), 135.8kb ± 197.7kb ($D_{MAX}$ = 10kb, *Method* = LRR mean, duplications).

$D_{MAX}$ = 10kb, *Method* = LRR sd, low-stringency QC callset from the GSA array had an average CNV size of 44.9kb (*st.dev* = 94.0kb). The average sizes of deletions were somewhat smaller than those for duplications, 38.1kb ± 81.8kb and 54.1kb ± 107.7kb, respectively. The results for medium-stringency QC callset followed similar trends: 117.6kb ± 178.5kb ($D_{MAX}$ = 10kb, *Method* = LRR sd, deletions and duplications), 104.7kb ± 175.6kb ($D_{MAX}$ = 10kb, *Method* = LRR sd, deletions), 129.8kb ± 180.5kb ($D_{MAX}$ = 10kb, *Method* = LRR sd, duplications). The low-stringency QC callset from the OEE array had an average CNV size of 62.4kb (*st.dev* = 187.2kb). The average sizes of deletions were somewhat smaller than those for duplications, 50.0kb ± 115.4kb and 88.8kb ± 283.6kb, respectively. The results for the medium-stringency QC callset followed similar trends: 119.8kb ± 205.3kb ($D_{MAX}$ = 10kb, *Method* = LRR sd, deletions and duplications), 100.4kb ± 167.6kb ($D_{MAX}$ = 10kb, *Method* = LRR sd, deletions), 137.0kb ± 232.5kb ($D_{MAX}$ = 10kb, *Method* = LRR sd, duplications).

$D_{MAX}$ = 10kb, *Method* = Distance, low-stringency QC callset from the GSA array had an average CNV size of 44.8kb (*st.dev* = 93.9kb). The average sizes of deletions were somewhat smaller than those for duplications, 38.0kb ± 81.5kb and 54.1kb ± 107.7kb, respectively. The results for medium-stringency QC callset followed similar trends: 116.8kb ± 177.5kb ($D_{MAX}$ = 10kb, *Method* = Distance, deletions and duplications), 103.2kb ± 173.2kb ($D_{MAX}$ = 10kb, *Method* = Distance, deletions), 130.2kb ± 180.8kb ($D_{MAX}$ = 10kb, *Method* = Distance, duplications). The low-stringency QC callset from the OEE array had an average CNV size of 62.0kb (*st.dev* = 186.9kb). The average sizes of deletions were somewhat smaller than those for duplications, 49.2kb ± 113.5kb and 89.7kb ± 285.3kb, respectively. The results for the medium-stringency QC callset followed similar trends: 119.7kb ± 190.7kb ($D_{MAX}$ = 10kb, *Method* = Distance, deletions and duplications), 104.1kb ± 179.3kb ($D_{MAX}$ = 10kb, *Method* = Distance, deletions), 133.2kb ± 199.1kb ($D_{MAX}$ = 10kb, *Method* = Distance, duplications).

Detailed reports for size-stratified bins are available in the **Supplemental Table S1L**.

##### S2.3.3 CNV Confidence Score Summaries

*Full Set* low-stringency QC callset from the GSA array had an average CNV confidence score of 22.8 (*st.dev* = 40.3). The confidence scores of deletions were similar to those for duplications, 23.0 ± 40.2 and 22.7 ± 40.4, respectively. The results for medium-stringency QC callset followed slightly different trends, with deletions having somewhat higher scores: 51.0 ± 79.8 (*Full Set* deletions and duplications), 55.5 ± 86.8 (*Full Set* deletions), 48.0 ± 74.5 (*Full Set* duplications). The low-stringency QC callset from the OEE array had an average CNV confidence score of 32.2 (*st.dev* = 60.7). The average confidence scores of deletions were somewhat lower than those for duplications, 29.3 ± 50.3 and 39.9 ± 81.9, respectively. The results for the medium-stringency QC callset followed similar trends: 99.3 ± 150.7 (*Full Set* deletions and duplications), 94.6 ± 139.0 (*Full Set* deletions), 104.0 ± 161.6 (*Full Set* duplications).

*Exact Match* low-stringency QC callset from the GSA array had an average CNV confidence score of 23.8 (*st.dev* = 38.4). The confidence scores of deletions were similar to those for duplications, 21.4 ± 34.2 and 26.3 ± 42.0, respectively. The results for medium-stringency QC callset followed similar trends: 33.8 ± 50.7 (*Exact Match* deletions and duplications), 31.2 ± 47.4 (*Exact Match* deletions), 35.7 ± 52.8 (*Exact Match* duplications). The low-stringency QC callset from the OEE array had an average CNV confidence score of 27.9 (*st.dev* = 40.1). The average confidence scores of deletions were noticeably lower than those for duplications, 23.2 ± 32.4 and 34.0 ± 47.7, respectively. The results for the medium-stringency QC callset followed similar trends: 36.8 ± 48.2 (*Exact Match* deletions and duplications), 32.9 ± 43.2 (*Exact Match* deletions), 40.0 ± 51.7 (*Exact Match* duplications).

$D_{MAX}$ = 10kb, *Method* = BAF, low-stringency QC callset from the GSA array had an average CNV confidence score of 22.6 (*st.dev* = 45.8). The confidence scores of deletions were similar to those for duplications, 23.4 ± 44.6 and 21.5 ± 47.4, respectively. The results for medium-stringency QC callset followed similar trends: 54.6 ± 95.4 ($D_{MAX}$ = 10kb, *Method* = BAF, deletions and duplications), 55.7 ± 94.8 ($D_{MAX}$ = 10kb, *Method* = BAF, deletions), 53.5 ± 96.1 ($D_{MAX}$ = 10kb, *Method* = BAF, duplications). The low-stringency QC callset from the OEE array had an average CNV confidence score of 29.7 (*st.dev* = 53.7). The average confidence scores of deletions were noticeably lower than those for duplications, 26.7 ± 44.5 and 36.8 ± 70.0, respectively. The results for the medium-stringency QC callset followed similar trends: 70.0 ± 106.2 ($D_{MAX}$ = 10kb, *Method* = BAF, deletions and duplications), 67.3 ± 97.8 ($D_{MAX}$ = 10kb, *Method* = BAF, deletions), 72.6 ± 113.7 ($D_{MAX}$ = 10kb, *Method* = BAF, duplications).

$D_{MAX}$ = 10kb, *Method* = LRR mean, low-stringency QC callset from the GSA array had an average CNV confidence score of 22.5 (*st.dev* = 45.9). The confidence scores of deletions were similar to those for duplications, 23.4 ± 44.6 and 21.4 ± 47.4, respectively. The results for medium-stringency QC callset followed similar trends: 55.6 ± 100.4 ($D_{MAX}$ = 10kb, *Method* = LRR mean, deletions and duplications), 57.5 ± 104.3 ($D_{MAX}$

= 10kb, *Method* = LRR mean, deletions), 53.9 ± 96.6 ($D_{MAX}$ = 10kb, *Method* = LRR mean, duplications). The low-stringency QC callset from the OEE array had an average CNV confidence score of 32.5 (*st.dev* = 63.0). The average confidence scores of deletions were noticeably lower than those for duplications, 28.2 ± 51.7 and 41.5 ± 81.3, respectively. The results for the medium-stringency QC callset followed similar trends: 77.6 ± 117.1 ($D_{MAX}$ = 10kb, *Method* = LRR mean, deletions and duplications), 73.8 ± 112.9 ($D_{MAX}$ = 10kb, *Method* = LRR mean, deletions), 81.1 ± 120.8 ($D_{MAX}$ = 10kb, *Method* = LRR mean, duplications).

$D_{MAX}$ = 10kb, *Method* = LRR sd, low-stringency QC callset from the GSA array had an average CNV confidence score of 22.7 (*st.dev* = 46.0). The confidence scores of deletions were similar to those for duplications, 23.5 ± 44.8 and 21.6 ± 47.7, respectively. The results for medium-stringency QC callset followed similar trends: 55.5 ± 100.3 ($D_{MAX}$ = 10kb, *Method* = LRR sd, deletions and duplications), 57.3 ± 104.2 ($D_{MAX}$ = 10kb, *Method* = LRR sd, deletions), 53.8 ± 96.4 ($D_{MAX}$ = 10kb, *Method* = LRR sd, duplications). The low-stringency QC callset from the OEE array had an average CNV confidence score of 31.6 (*st.dev* = 59.2). The average confidence scores of deletions were noticeably lower than those for duplications, 27.7 ± 47.8 and 39.8 ± 77.7, respectively. The results for the medium-stringency QC callset followed somewhat different trends, with deletions and duplications having rather similar confidence scores: 75.1 ± 110.7 ($D_{MAX}$ = 10kb, *Method* = LRR sd, deletions and duplications), 72.5 ± 104.8 ($D_{MAX}$ = 10kb, *Method* = LRR sd, deletions), 77.4 ± 115.7 ($D_{MAX}$ = 10kb, *Method* = LRR sd, duplications).

$D_{MAX}$ = 10kb, *Method* = Distance, low-stringency QC callset from the GSA array had an average CNV confidence score of 22.6 (*st.dev* = 45.9). The confidence scores of deletions were similar to those for duplications, 23.4 ± 44.7 and 21.5 ± 47.4, respectively. The results for medium-stringency QC callset followed similar trends: 54.9 ± 99.6 ($D_{MAX}$ = 10kb, *Method* = Distance, deletions and duplications), 56.3 ± 103.0 ($D_{MAX}$ = 10kb, *Method* = Distance, deletions), 53.5 ± 96.3 ($D_{MAX}$ = 10kb, *Method* = Distance, duplications). The low-stringency QC callset from the OEE array had an average CNV confidence score of 31.6 (*st.dev* = 61.0). The average confidence scores of deletions were noticeably lower than those for duplications, 27.6 ± 49.7 and 40.4 ± 79.4, respectively. The results for the medium-stringency QC callset followed somewhat different trends, with deletions and duplications having rather similar confidence scores: 77.3 ± 120.2 ($D_{MAX}$ = 10kb, *Method* = Distance, deletions and duplications), 75.9 ± 118.9 ($D_{MAX}$ = 10kb, *Method* = Distance, deletions), 78.4 ± 121.3 ($D_{MAX}$ = 10kb, *Method* = Distance, duplications).

Detailed reports for size-stratified bins are available in the **Supplemental Table S1L**.

##### S2.3.4 Sensitivity Analyses

The overall sensitivity of the *Exact Match* approach on the GSA array was 0.074 and 0.059, and on the OEE array was 0.059 and 0.049, for low- and medium-stringency QC callsets respectively. In the medium-stringency QC GSA callset, the sensitivity was higher for larger CNVs: 0.033 (under 100kb), 0.295 (100kb to 500kb), 0.415 (over 500kb). The medium-stringency QC OEE callset followed similar trends: 0.029 (under 100kb), 0.179 (100kb to 500kb), 0.348 (over 500kb).

The MarkerMatch algorithm shows sensitivity to be similar across the different *Method* parameters on the GSA array: 0.070 (under 100kb), 0.472 (100kb to 500kb), 0.527 (over 500kb), averaged across all four *Method* parameters, at medium-stringency QC. Similar was true for the OEE array: 0.069 (under 100kb), 0.295 (100kb to 500kb), 0.423 (over 500kb).

*Full Set* sensitivity analysis indicates just how small the overlap of surveyed genomic regions is between the two arrays, with GSA array only ever capturing 32.2% of CNVs called on OEE array, and OEE capturing only 26.5% of CNVs called on GSA array. In addition to the batch-wise reduction in sensitivity, there also seem to be QC-related sensitivity reductions (e.g., 57% lower sensitivity in OEE low- vs. medium-stringency QC callset).

Detailed report is available in the **Supplemental Table S1N**. Sensitivity CAE plots are available in **Supplemental Figure S43** for all CNV types, as well as **Supplemental Figures S49** and **S56** for deletions and duplications, respectively.

##### S2.3.5 Positive Predictive Value Analyses

The overall PPV of the *Exact Match* approach on the GSA array was 0.558 and 0.741, and on the OEE array was 0.654 and 0.820, for low- and medium-stringency QC callsets respectively. In the medium-stringency QC GSA callset, the PPV was higher for larger CNVs: 0.646 (under 100kb), 0.857 (100kb to 500kb), 1.000 (over 500kb). The medium-stringency QC OEE callset followed similar trends: 0.757 (under 100kb), 0.895 (100kb to 500kb), 1.000 (over 500kb).

The MarkerMatch algorithm shows PPV to be similar across the different *Method* parameters on the GSA array: 0.633 (under 100kb), 0.837 (100kb to 500kb), 0.977 (over 500kb), averaged across all four *Method* parameters, at medium-stringency QC. Similar was true for the OEE array: 0.772 (under 100kb), 0.929 (100kb to 500kb), 0.960 (over 500kb). The best-performing *Method* was LRR mean for both GSA and OEE, with overall PPVs of 0.711 and 0.844, respectively.

There also appeared to be an array-specific effect in terms of PPV, with GSA *Full Set* performing at PPV reductions of 12.0% and 35.8% compared to OEE, for low- and medium-stringency QC callsets, respectively. Detailed report is available in the **Supplemental Table S1N**. Sensitivity CAE plots are available in **Figure 5** for all CNV types, as well as **Supplemental Figures S50** and **S57** for deletions and duplications, respectively.

##### S2.3.6 False Negative Rate Analyses

Since the false negative rate (FNR) is a complement to sensitivity, as per equations **Eq. 1** and **Eq. 3**. Overall, FNRs appear to be highest for the *Exact Match*, at 0.926 and 0.941 on GSA, and 0.941 and 0.951 on OEE, for low- and medium-stringency QC callsets, respectively. FNR tends to drop with larger-size CNV bins.

A detailed FNR report is available in the **Supplemental Table S1N**. FNR CAE plots are available in **Supplemental Figure S44** for all CNV types, as well as **Supplemental Figures S51** and **S58** for deletions and duplications, respectively.

##### S2.3.7 False Discovery Rate Analyses

Since the false discovery rate (FDR) is a complement to PPV, as per equations **Eq. 2** and **Eq. 4**. Similarly, but inversely, FDRs were lower for MarkerMatch callsets than *Exact Match* for all configurations of *Method* parameters and $D_{MAX}$ < 100kb.

A detailed FDR report is available in the **Supplemental Table S1N**. FDR CAE plots are available in **Supplemental Figure S45** for all CNV types, as well as **Supplemental Figures S52** and **S59** for deletions and duplications, respectively.

##### S2.3.8 F1 Score Analyses

F1 score for MarkerMatch performed either as well as *Exact Match* and *Full Set*, or better, for both GSA and OEE arrays. This effect seemed to follow in CNV size-stratified cases for all CNV-size and array configurations, with the exception of large CNVs (over 500kb) on the OEE array, where *Full Set* performed noticeably better than MarkerMatch or *Exact Match* callsets (*F1* = 0.682)

A detailed F1 report is available in the **Supplemental Table S1N**. F1 CAE plots are available in **Supplemental Figure S46** for all CNV types, as well as **Supplemental Figures S53** and **S60** for deletions and duplications, respectively.

##### S2.3.9 Fowlkes-Mallows Index Analyses

The results for FMI were similar to those of F1 in the CAE. A detailed FMI report is available in the **Supplemental Table S1N**. FMI CAE plots are available in **Supplemental Figure S47** for all CNV types, as well as **Supplemental Figures S54** and **S61** for deletions and duplications, respectively.

##### S2.3.10 Jaccard Index Analyses

The results for JI were similar to those of F1 and FMI in the CAE. A detailed JI report is available in the **Supplemental Table S1N**. JI CAE plots are available in **Supplemental Figure S48** for all CNV types, as well as **Supplemental Figures S55** and **S62** for deletions and duplications, respectively.

##### S2.3.11 Deletions vs. Duplications

Deletions and duplications followed roughly the same trend when it came to most metrics, however, some interesting patterns emerge when looking at sensitivity in large CNV sizes. Namely, it appears that GSA overperformed OEE for large duplications, whereas OEE overperformed GSA for large deletions. MarkerMatch, however, tended to perform equally well or better than *Exact Match*. For example, among large deletions OEE sensitivity was 2.1-fold that of GSE, whereas among duplications that sensitivity was 0.6-fold.

A detailed report is available in the **Supplemental Table S1N**. These data have been graphically shown for deletions and duplications in the **Supplemental Figures S49-62**.

##### S2.3.12 Determination of Optimal Minimum CNV Size and SNP Coverage Thresholds

Beta regression of sensitivity on the GSA array has shown *CNV length cutoff*, *marker coverage cutoff*, and their interaction terms to be significantly associated ([$OR_{TPR}$ = 1.05, $p_{TPR}$ < 0.001]; [$OR_{TPR}$ = 1.04, $p_{TPR}$ <

0.001]; [$OR_{TPR}$ = 1.00, $p_{TPR}$ < 0.001] respectively). In the OEE array, we observed similar associations ([$OR_{TPR}$ = 1.05, $p_{TPR}$ < 0.001]; [$OR_{TPR}$ = 1.02, $p_{TPR}$ < 0.001]; [$OR_{TPR}$ = 1.00, $p_{TPR}$ < 0.001] respectively). While the associations were significant, the effect sizes were rather small. The model's pseudo $R^2$ was 0.70.

Unlike in GSA array, *Method* terms were significant in OEE array, with LRR sd underperforming ([$OR_{TPR}$ = 0.97, $p_{TPR}$ = 0.007]), and BAF and Distance overperforming ([$OR_{TPR}$ = 1.23, $p_{TPR}$ < 0.001]; [$OR_{TPR}$= 1.11, $p_{TPR}$ < 0.001], respectively) relative to LRR mean. The model's pseudo $R^2$ was 0.96.

Beta regressions of F1, FMI, and JI scores on the GSA array have indicated trends similar to PPV and sensitivity above. Namely, *CNV length cutoff* ([$OR_{F1}$ = 1.05. $p_{F1}$ < 0.001], [$OR_{FMI}$ = 1.05. $p_{FMI}$ < 0.001], [$OR_{JI}$ = 1.05. $p_{JI}$ < 0.001]), *marker coverage cutoff* ([$OR_{F1}$ = 1.03. $p_{F1}$ < 0.001], [$OR_{FMI}$ = 1.03. $p_{FMI}$ < 0.001], [$OR_{JI}$ = 1.03. $p_{JI}$ < 0.001]), and their interaction terms ([$OR_{F1}$ = 1.00. $p_{F1}$ = 0.002], [$OR_{FMI}$ = 1.00. $p_{FMI}$ = 0.01], [$OR_{JI}$ = 1.00. $p_{JI}$ = 0.01]) were all significantly associated with the measurement metrics, while the individual *Methods* were not with a couple of exceptions. The pseudo $R^2$ were 0.80 (F1 model), 0.81 (FMI model), and 0.80 (JI model). No significant *Method* terms were present in any of the F1, FMI, nor JI models.

Beta regressions of F1, FMI, and JI scores on the OEE array have resulted in similar trends as those on GSA array, but with a better fit, with pseudo $R^2$ values of 0.95 for all three (F1, FMI, and JI) models. *CNV length cutoff* ([$OR_{F1}$ = 1.03. $p_{F1}$ < 0.001], [$OR_{FMI}$ = 1.03. $p_{FMI}$ < 0.001], [$OR_{JI}$ = 1.04. $p_{JI}$ < 0.001]), *marker coverage cutoff* ([$OR_{F1}$ = 1.01. $p_{F1}$ < 0.001], [$OR_{FMI}$ = 1.01. $p_{FMI}$ < 0.001], [$OR_{JI}$ = 1.01. $p_{JI}$ < 0.001]), and their interaction terms ([$OR_{F1}$ = 1.00. $p_{F1}$ = 0.006], [$OR_{FMI}$ = 1.00. $p_{FMI}$ = 0.01], [$OR_{JI}$ = 1.00. $p_{JI}$ = 0.002]) were all significantly associated with the measurement metrics. In all three models, BAF ([$OR_{F1}$ = 1.34. $p_{F1}$ < 0.001], [$OR_{FMI}$ = 1.10. $p_{FMI}$ < 0.001], [$OR_{JI}$ = 1.14. $p_{JI}$ < 0.001]) and Distance ([$OR_{F1}$ = 1.08. $p_{F1}$ < 0.001], [$OR_{FMI}$ = 1.07. $p_{FMI}$ < 0.001], [$OR_{JI}$ = 1.08. $p_{JI}$ < 0.001]) *Methods* also performed better than LRR mean. LRR sd was not outperforming LRR mean significantly in any of the three models.

An inspection of the plots displaying PPV and F1 scores across various breakdowns of *CNV length* and *marker coverage cutoffs* shows size-dependent diminishing of batch (array) and MarkerMatch/*Method* effects (**Supplemental Figures S70-75**). While it is evident larger CNV sizes and higher marker coverage are overall favorable, there are no specific recommendable cutoff values that drastically improve either PPV or F1, and any improvements are far from perfect.

### S4 Supplementary Tables

| **Supplementary Table S1** |
|---|
| **Table S1A**. Tabulated results of execution times (*S1A: Execution Times*). |
| **Table S1B**. Tabulated array coverage metrics (*S1B: Array Coverage*). |
| **Table S1C**. Tabulated inter-marker gap statistics (*S1C: Inter-Marker Gaps*). |
| **Table S1D**. Tabulated BAF metric statistics (*S1D: BAF Metric*). |
| **Table S1E**. Tabulated LRR sd metric statistics (*S1E: LRR sd Metric*). |
| **Table S1F**. Tabulated LRR mean metric statistics (*S1F: LRR mean Metric*). |
| **Table S1G**. Tabulated Within-Array Experiment callset summaries (*S1G: WAE Callset Summaries*). |
| **Table S1H**. Tabulated Within-Array Experiment callset sample summaries (*S1H: WAE Callset Sample Summaries*). |
| **Table S1I**. Tabulated Within-Array Experiment callset analytics (*S1I: WAE Genome-Wide Callset Analytics*). |
| **Table S1J**. Tabulated Within-Array Experiment callset analytics (*S1J: WAE Regional Callset Analytics*). |
| **Table S1K**. Tabulated Within-Array Experiment Method modeling (*S1K: WAE Method Modeling*). |
| **Table S1L**. Tabulated Cross-Array Experiment callset summaries (*S1L: CAE Callset Summaries*). |
| **Table S1M**. Tabulated Cross-Array Experiment callset sample summaries (*S1M: CAE Callset Sample Summaries*). |
| **Table S1N**. Tabulated Cross-Array Experiment callset analytics (*S1N: CAE Genome-Wide Callset Analytics*). |
| **Table S1O**. Tabulated Cross-Array Experiment callset analytics (*S1O: CAE Regional Callset Analytics*). |
| **Table S1P**. Tabulated sample-wise performance (*S1P: Sample-WIse Performance*). |
| **Table S1Q**. Tabulated cumulative metric-wise performance (*S1Q: Cumulative Metric-Wise Performance*). |

**Table 2A**. Table summarizing sample counts within specified ranges of PPV for medium-stringency QC callsets.

| PPV Values | Full Set | | | BAF | | | LRR mean | | | LRR sd | | | Distance | | | Exact Match | | |
|---|---|---|---|---|---|---|---|---|---|---|---|---|---|---|---|---|---|---|
| | GSA | OEE | SSC | GSA | OEE | SSC | GSA | OEE | SSC | GSA | OEE | SSC | GSA | OEE | SSC | GSA | OEE | SSC |
| 0.0 | 26 | 9 | 0 | 20 | 9 | 4 | 20 | 16 | 3 | 21 | 18 | 3 | 20 | 12 | 1 | 39 | 29 | 113 |
| (0.0, 0.1) | 4 | 0 | 0 | 0 | 0 | 0 | 0 | 0 | 0 | 0 | 0 | 0 | 0 | 0 | 0 | 0 | 0 | 0 |
| [0.1, 0.2) | 20 | 2 | 0 | 6 | 2 | 0 | 6 | 0 | 1 | 4 | 0 | 0 | 4 | 2 | 0 | 1 | 0 | 0 |
| [0.2, 0.3) | 63 | 8 | 0 | 13 | 7 | 2 | 12 | 6 | 0 | 15 | 8 | 1 | 16 | 6 | 0 | 13 | 4 | 7 |
| [0.3, 0.4) | 59 | 10 | 0 | 23 | 8 | 8 | 24 | 7 | 2 | 22 | 8 | 3 | 23 | 7 | 3 | 17 | 10 | 20 |
| [0.4, 0.5) | 62 | 9 | 0 | 19 | 8 | 10 | 17 | 6 | 3 | 20 | 3 | 1 | 15 | 5 | 3 | 2 | 1 | 2 |
| [0.5, 0.6) | 115 | 45 | 0 | 59 | 31 | 37 | 54 | 38 | 26 | 57 | 42 | 27 | 62 | 42 | 18 | 64 | 35 | 174 |
| [0.6, 0.7) | 91 | 66 | 0 | 81 | 40 | 81 | 84 | 56 | 54 | 83 | 52 | 67 | 82 | 55 | 51 | 39 | 21 | 148 |
| [0.7, 0.8) | 51 | 49 | 0 | 57 | 24 | 185 | 59 | 39 | 110 | 60 | 52 | 152 | 52 | 42 | 134 | 12 | 8 | 63 |
| [0.8, 0.9) | 35 | 65 | 0 | 70 | 56 | 598 | 66 | 63 | 527 | 67 | 71 | 628 | 70 | 58 | 555 | 9 | 1 | 35 |
| [0.9, 1.0) | 1 | 3 | 0 | 1 | 0 | 216 | 2 | 0 | 191 | 1 | 1 | 287 | 1 | 0 | 293 | 0 | 0 | 0 |
| 1.0 | 25 | 129 | 3839 | 184 | 210 | 2694 | 191 | 334 | 2996 | 184 | 332 | 2735 | 190 | 271 | 2854 | 317 | 319 | 2390 |

**Table 2B**. Table summarizing sample counts within specified ranges of F1 for medium-stringency QC callsets.

| F1 Values | Full Set | | | BAF | | | LRR mean | | | LRR sd | | | Distance | | | Exact Match | | |
|---|---|---|---|---|---|---|---|---|---|---|---|---|---|---|---|---|---|---|
| | GSA | OEE | SSC | GSA | OEE | SSC | GSA | OEE | SSC | GSA | OEE | SSC | GSA | OEE | SSC | GSA | OEE | SSC |
| 0.0 | 27 | 10 | 0 | 20 | 13 | 6 | 20 | 18 | 6 | 21 | 19 | 3 | 20 | 13 | 2 | 40 | 31 | 128 |
| (0.0, 0.1) | 64 | 67 | 2 | 64 | 70 | 247 | 64 | 114 | 258 | 62 | 121 | 181 | 63 | 93 | 151 | 127 | 181 | 2375 |
| [0.1, 0.2) | 133 | 102 | 50 | 120 | 122 | 1421 | 120 | 168 | 1496 | 121 | 167 | 1337 | 122 | 160 | 1324 | 204 | 167 | 458 |
| [0.2, 0.3) | 165 | 127 | 532 | 157 | 112 | 1592 | 160 | 171 | 1627 | 157 | 172 | 1681 | 154 | 146 | 1687 | 105 | 43 | 6 |
| [0.3, 0.4) | 100 | 60 | 1366 | 99 | 56 | 512 | 98 | 64 | 480 | 102 | 73 | 617 | 103 | 65 | 656 | 26 | 6 | 0 |
| [0.4, 0.5) | 47 | 20 | 1306 | 46 | 20 | 55 | 47 | 26 | 49 | 45 | 29 | 80 | 47 | 17 | 91 | 10 | 2 | 0 |
| [0.5, 0.6) | 14 | 10 | 515 | 21 | 6 | 4 | 19 | 6 | 0 | 20 | 6 | 5 | 20 | 7 | 2 | 1 | 0 | 0 |
| [0.6, 0.7) | 2 | 0 | 59 | 4 | 0 | 0 | 5 | 0 | 0 | 4 | 1 | 0 | 4 | 0 | 0 | 0 | 0 | 0 |
| [0.7, 0.8) | 1 | 0 | 9 | 2 | 0 | 0 | 2 | 0 | 0 | 2 | 0 | 0 | 2 | 0 | 0 | 1 | 0 | 0 |
| [0.8, 0.9) | 0 | 0 | 0 | 0 | 0 | 0 | 0 | 0 | 0 | 0 | 0 | 0 | 0 | 0 | 0 | 0 | 0 | 0 |
| [0.9, 1.0) | 0 | 0 | 0 | 0 | 0 | 0 | 0 | 0 | 0 | 0 | 0 | 0 | 0 | 0 | 0 | 0 | 0 | 0 |
| 1.0 | 0 | 0 | 0 | 0 | 0 | 0 | 0 | 0 | 0 | 0 | 0 | 0 | 0 | 0 | 0 | 0 | 0 | 0 |

#### S5 Supplementary Figures

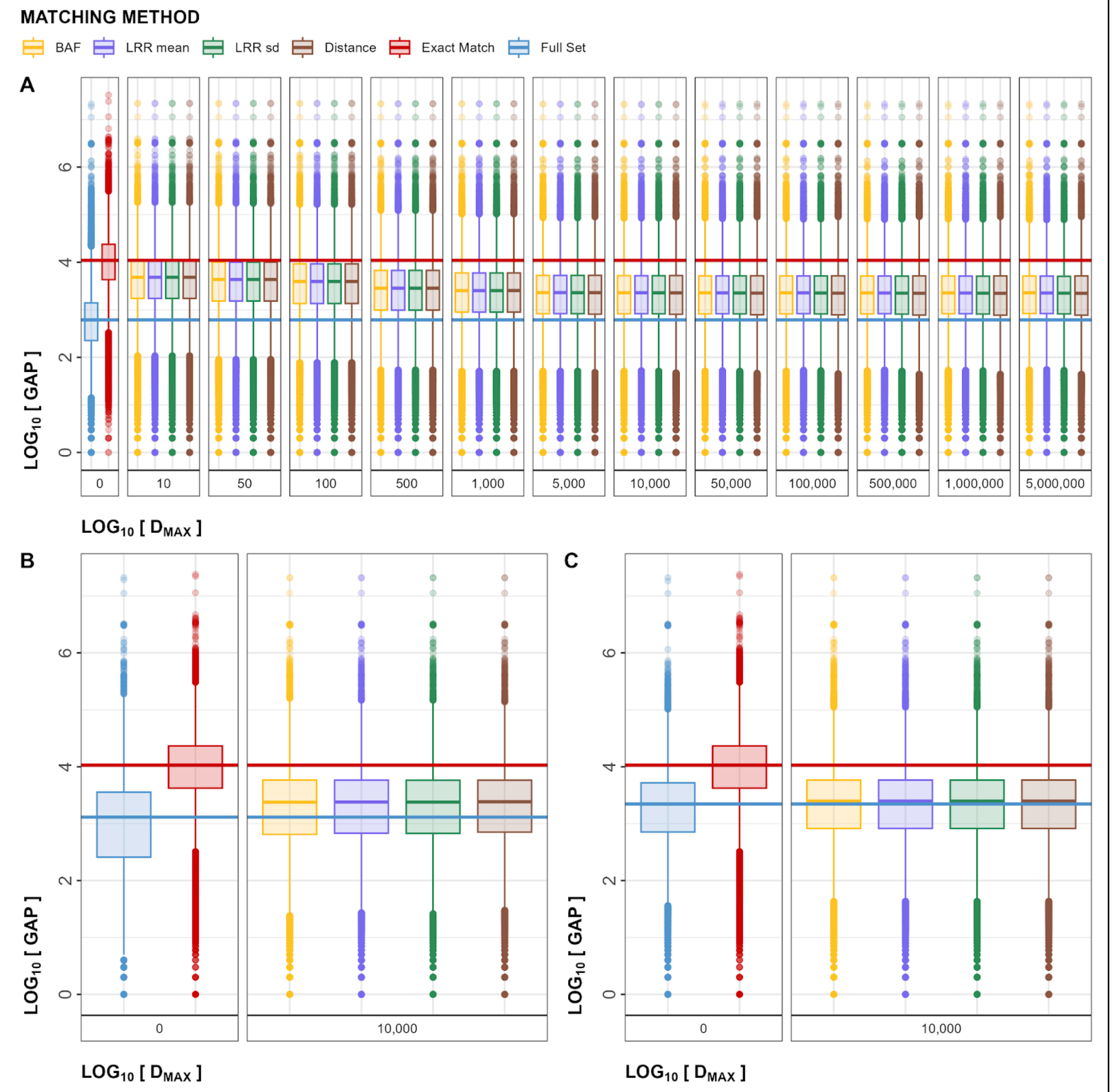


**Figure S1**. Inter-marker gaps in the Within-Array Experiment (A) and Cross-Array Experiment (B, C). Box plots of $\log_{10}$ transformations of the gap size between two adjacent markers are shown for full set (in blue), *Exact Match* (in red), and an array of MarkerMatch configurations (BAF based in yellow, LRR mean based in purple, LRR standard deviation based in green, and distance based in brown). X-axis shows *Full Set* and *Exact Match* for reference (with median lines spanning the entire graph), as well as different $D_{MAX}$ parameters (ranging from 10bp to 5,000,000bp). Gaps are shown for OMNI array (A), OEE array (B), and GSA array (C). OMNI: Omni2.5 array, GSA: Global Screening Array, OEE: Omni Express Exome array.

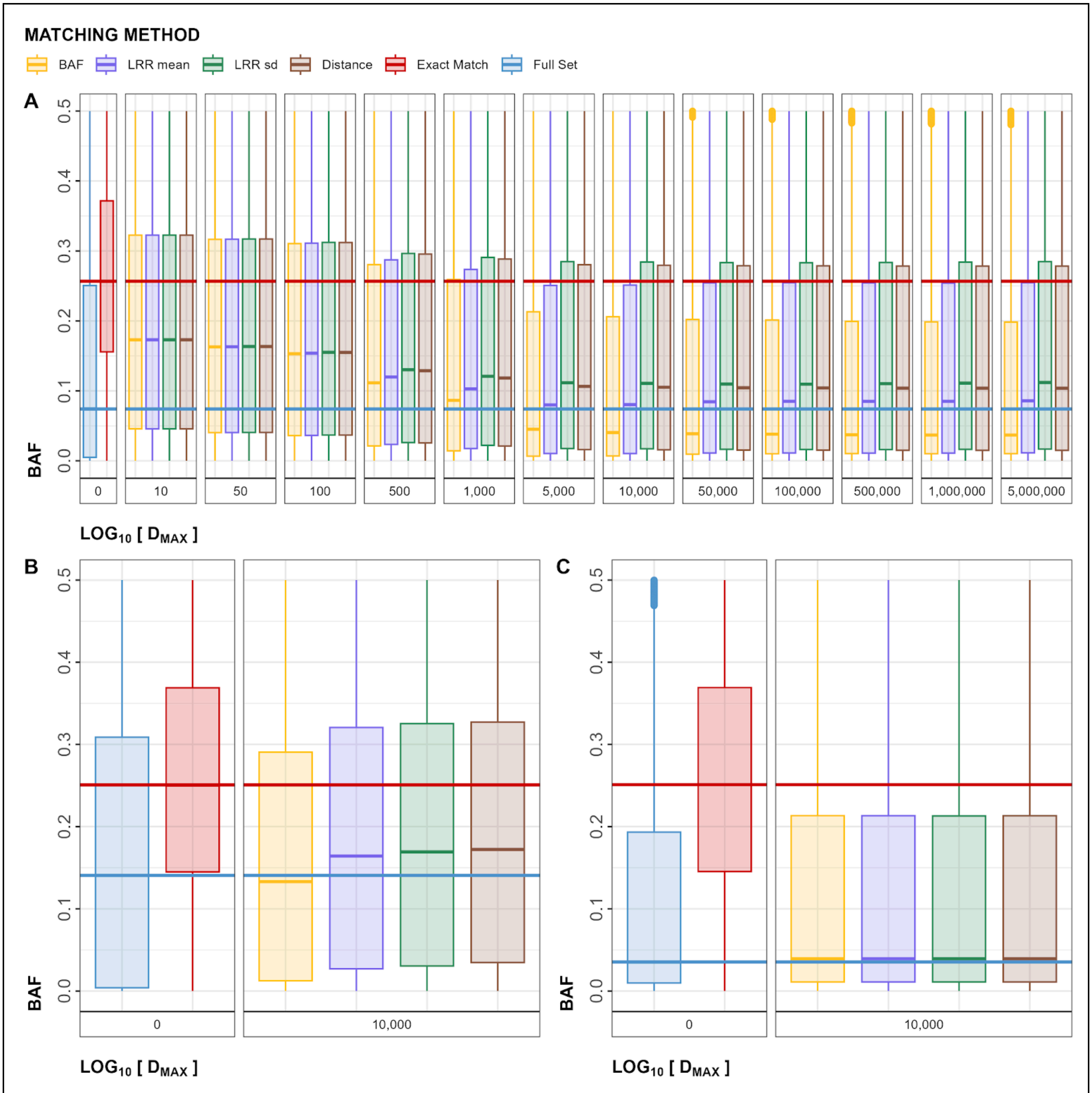


**Figure S2**. BAF metrics in the Within-Array Experiment (A) and Cross-Array Experiment (B, C). Box plots of marker-wise BAF are shown for *Full Set* (in blue), *Exact Match* (in red), and an array of MarkerMatch configurations (BAF based in yellow, LRR mean based in purple, LRR standard deviation based in green, and distance based in brown). X-axis shows *Full Set* and *Exact Match* for reference (with median lines spanning the entire graph), as well as different $D_{MAX}$ parameters (ranging from 10bp to 5,000,000bp). BAFs are shown for OMNI array (A), OEE array (B), and GSA array (C). OMNI: Omni2.5 array, GSA: Global Screening Array, OEE: Omni Express Exome array.

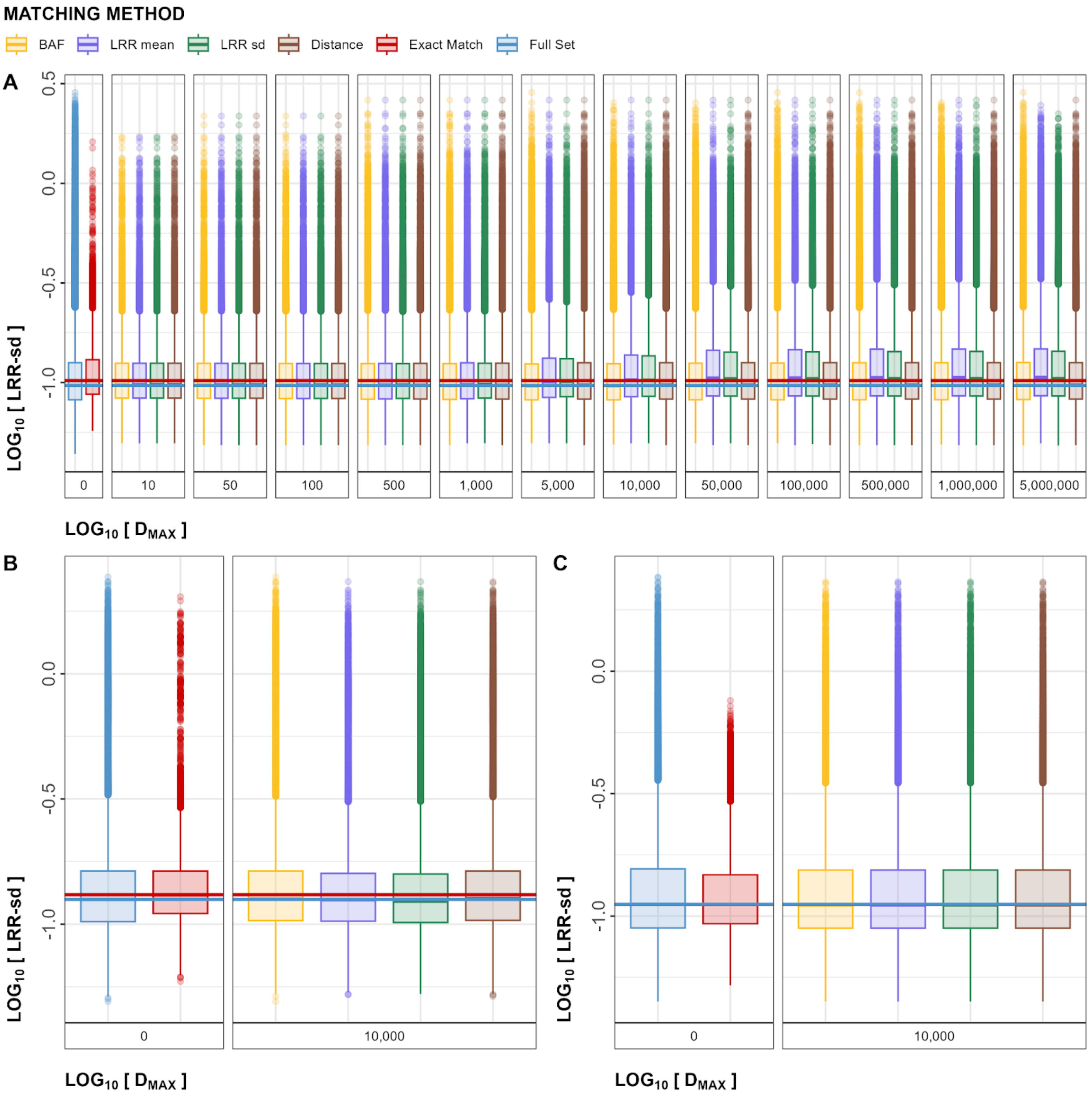


**Figure S3**. LRR sd metrics in the Within-Array Experiment (A) and Cross-Array Experiment (B, C). Box plots of $\log_{10}$ transformations of marker-wise LRR sds are shown for full set (in blue), Exact match (in red), and an array of MarkerMatch configurations (BAF based in yellow, LRR mean based in purple, LRR standard deviation based in green, and distance based in brown). X-axis shows *Full Set* and *Exact Match* for reference (with median lines spanning the entire graph), as well as different $D_{MAX}$ parameters (ranging from 10bp to 5,000,000bp). LRR sds are shown for OMNI array (A), OEE array (B), and GSA array (C). OMNI: Omni2.5 array, GSA: Global Screening Array, OEE: Omni Express Exome array.

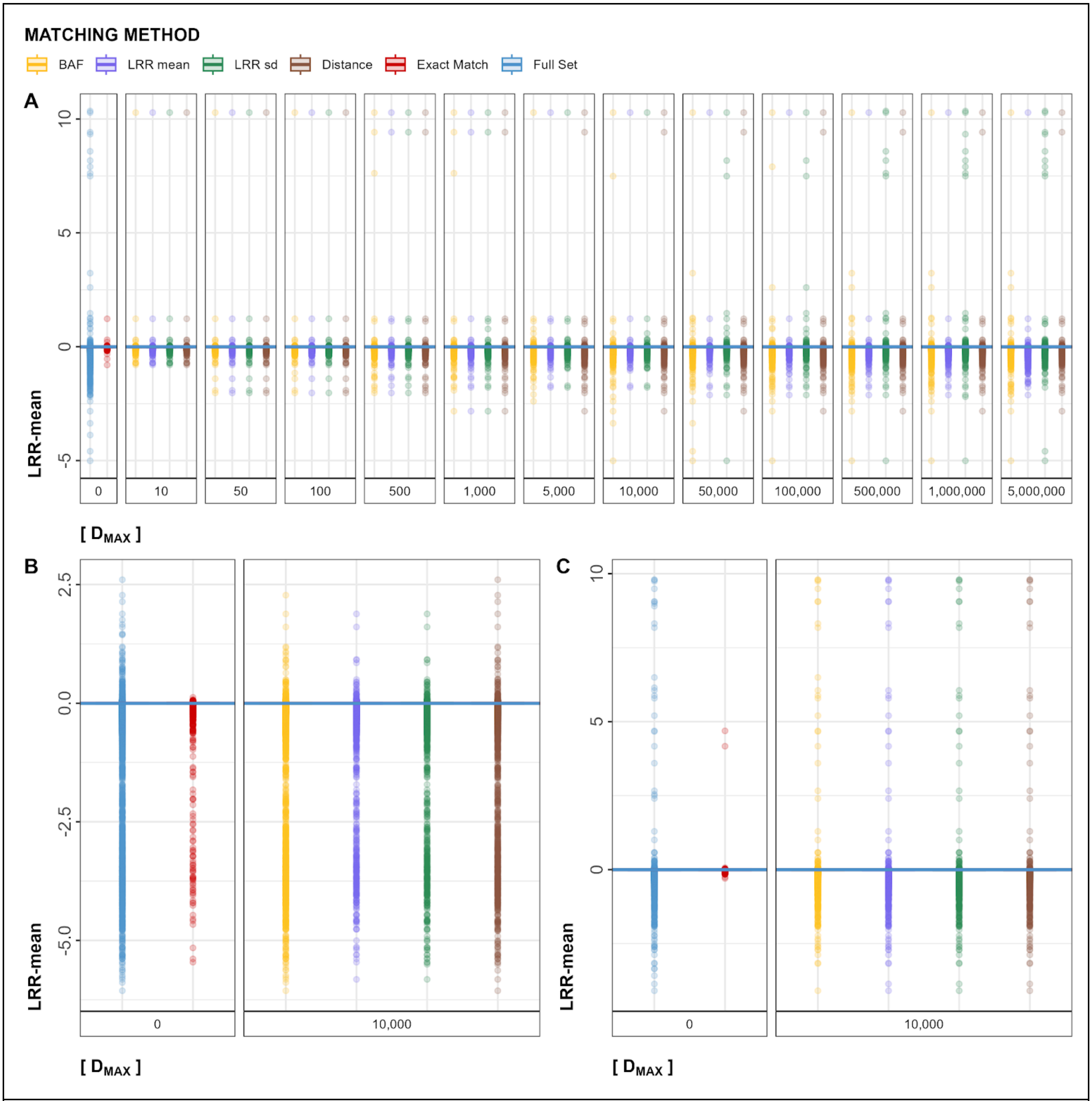


**Figure S4**. LRR mean metrics in the Within-Array Experiment (A) and Cross-Array Experiment (B, C). Box plots of marker-wise LRR means are shown for full set (in blue), Exact match (in red), and an array of MarkerMatch configurations (BAF based in yellow, LRR mean based in purple, LRR standard deviation based in green, and distance based in brown). X-axis shows *Full Set* and *Exact Match* for reference (with median lines spanning the entire graph), as well as different $D_{MAX}$ parameters (ranging from 10bp to 5,000,000bp). LRR means are shown for OMNI array (A), OEE array (B), and GSA array (C). OMNI: Omni2.5 array, GSA: Global Screening Array, OEE: Omni Express Exome array.

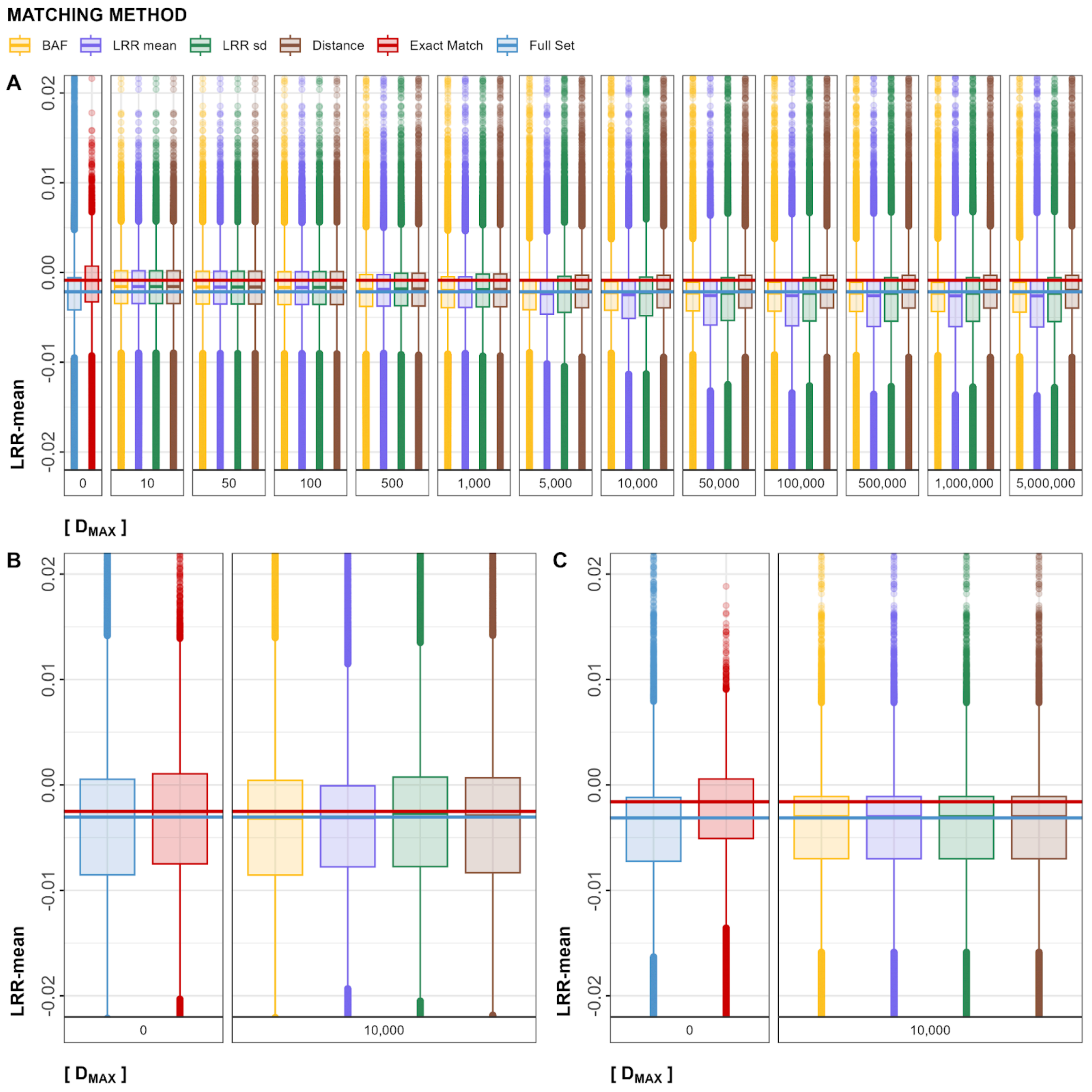


**Figure S5**. LRR mean metrics in the Within-Array Experiment (A) and Cross-Array Experiment (B, C). Box plots of marker-wise LRR means are shown for full set (in blue), Exact match (in red), and an array of MarkerMatch configurations (BAF based in yellow, LRR mean based in purple, LRR standard deviation based in green, and distance based in brown). X-axis shows *Full Set* and *Exact Match* for reference (with median lines spanning the entire graph), as well as different $D_{MAX}$ parameters (ranging from 10bp to 5,000,000bp). LRR means are shown for OMNI array (A), OEE array (B), and GSA array (C). The plots are zoomed to LRR mean range -0.02 to 0.02. OMNI: Omni2.5 array, GSA: Global Screening Array, OEE: Omni Express Exome array.

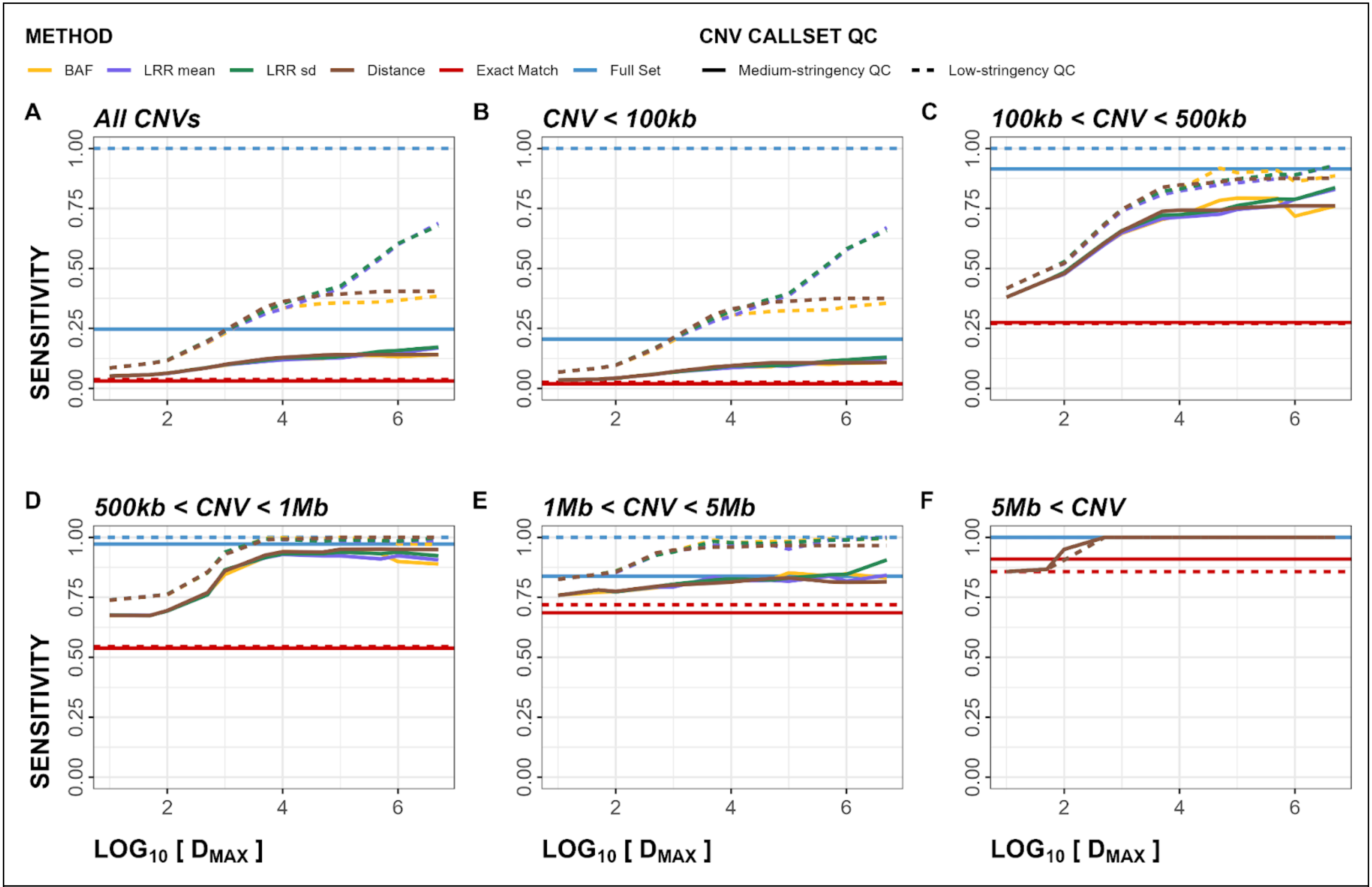


**Figure S6**. Within-Array Experiment sensitivity plots for both deletions and duplications. Panels A-F show sensitivity metrics stratified by CNV size. Dashed line represents low-stringency QC, solid line represents medium-stringency QC.

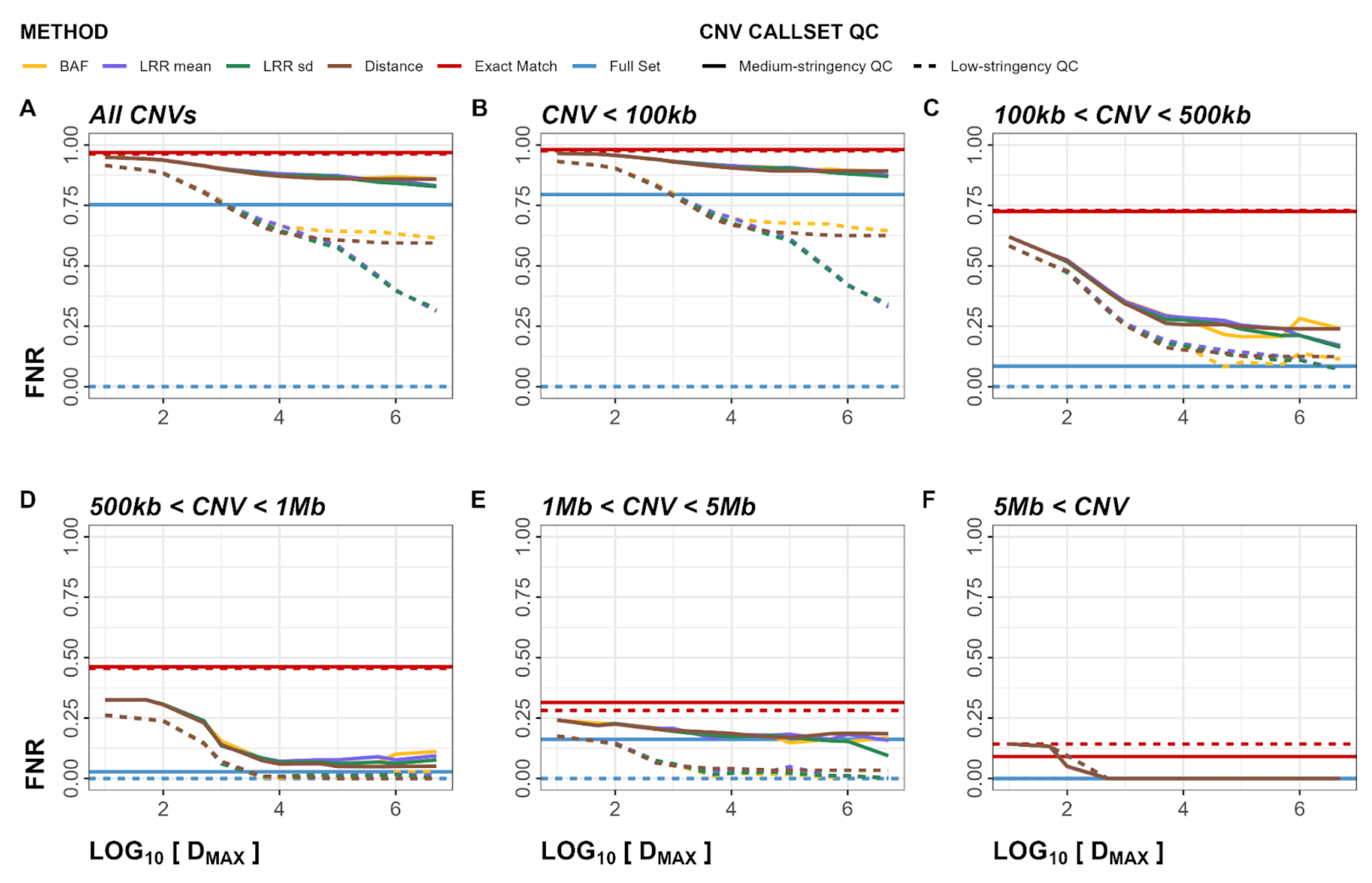


**Figure S7**. Within-Array Experiment false negative rate (FNR) plots for both deletions and duplications. Panels A-F show FNR metrics stratified by CNV size. Dashed line represents low-stringency QC, solid line represents medium-stringency QC.

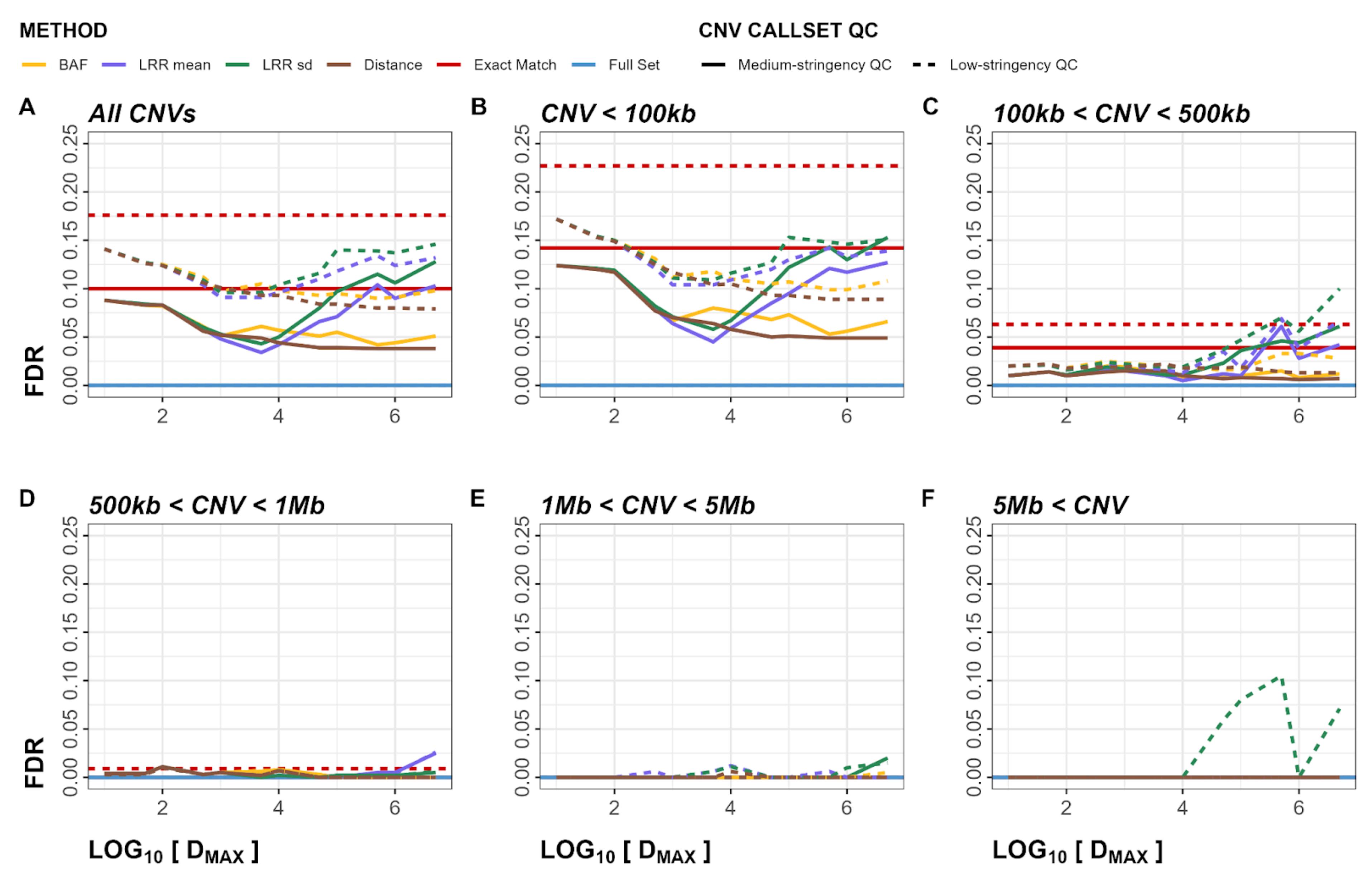


**Figure S8**. Within-Array Experiment false discovery rate (FDR) plots for both deletions and duplications. Panels A-F show FDR metrics stratified by CNV size. Dashed line represents low-stringency QC, solid line represents medium-stringency QC..

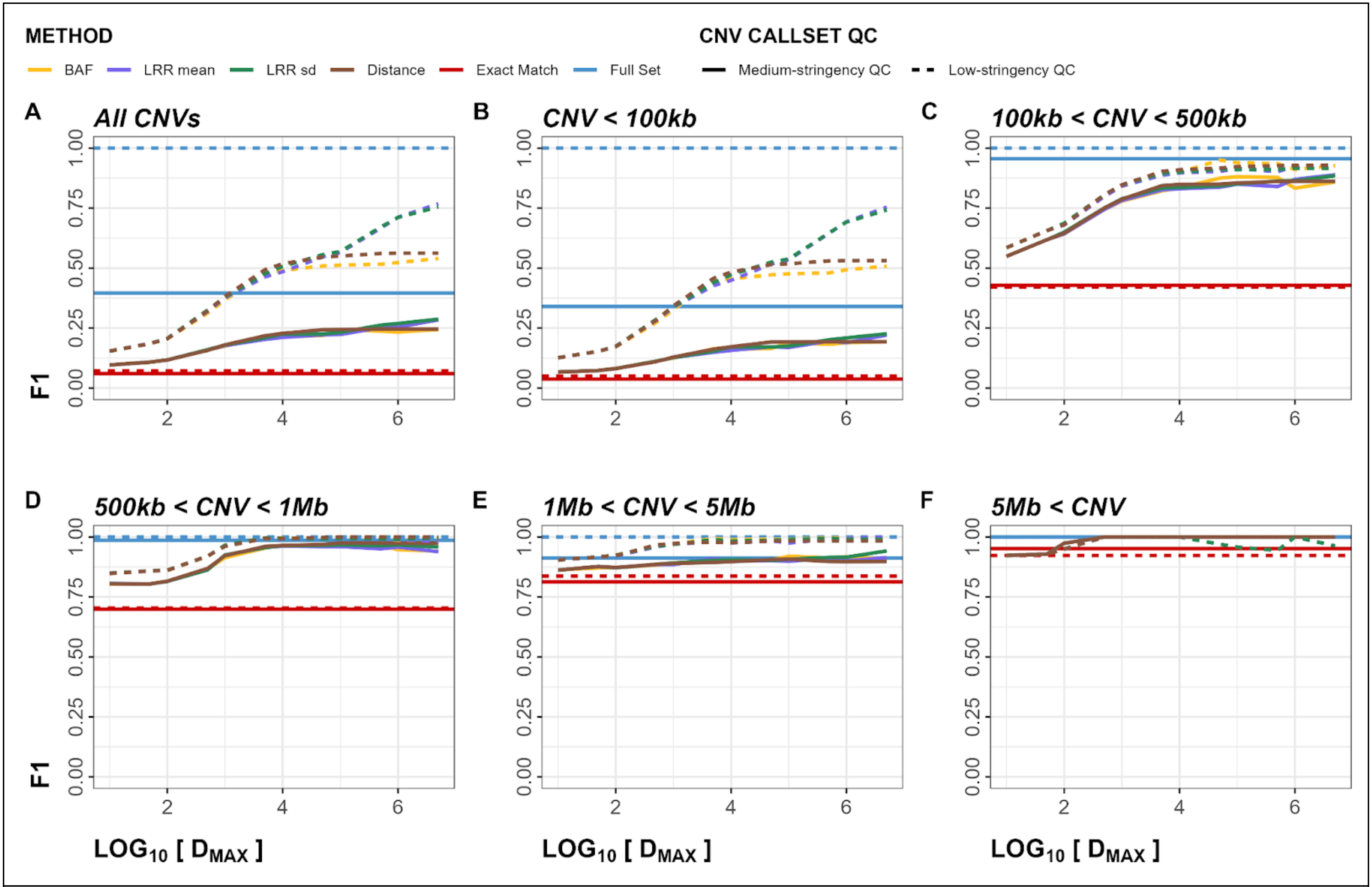


**Figure S9**. Within-Array Experiment F1 score plots for both deletions and duplications. Panels A-F show F1 metrics stratified by CNV size. Dashed line represents low-stringency QC, solid line represents medium-stringency QC.

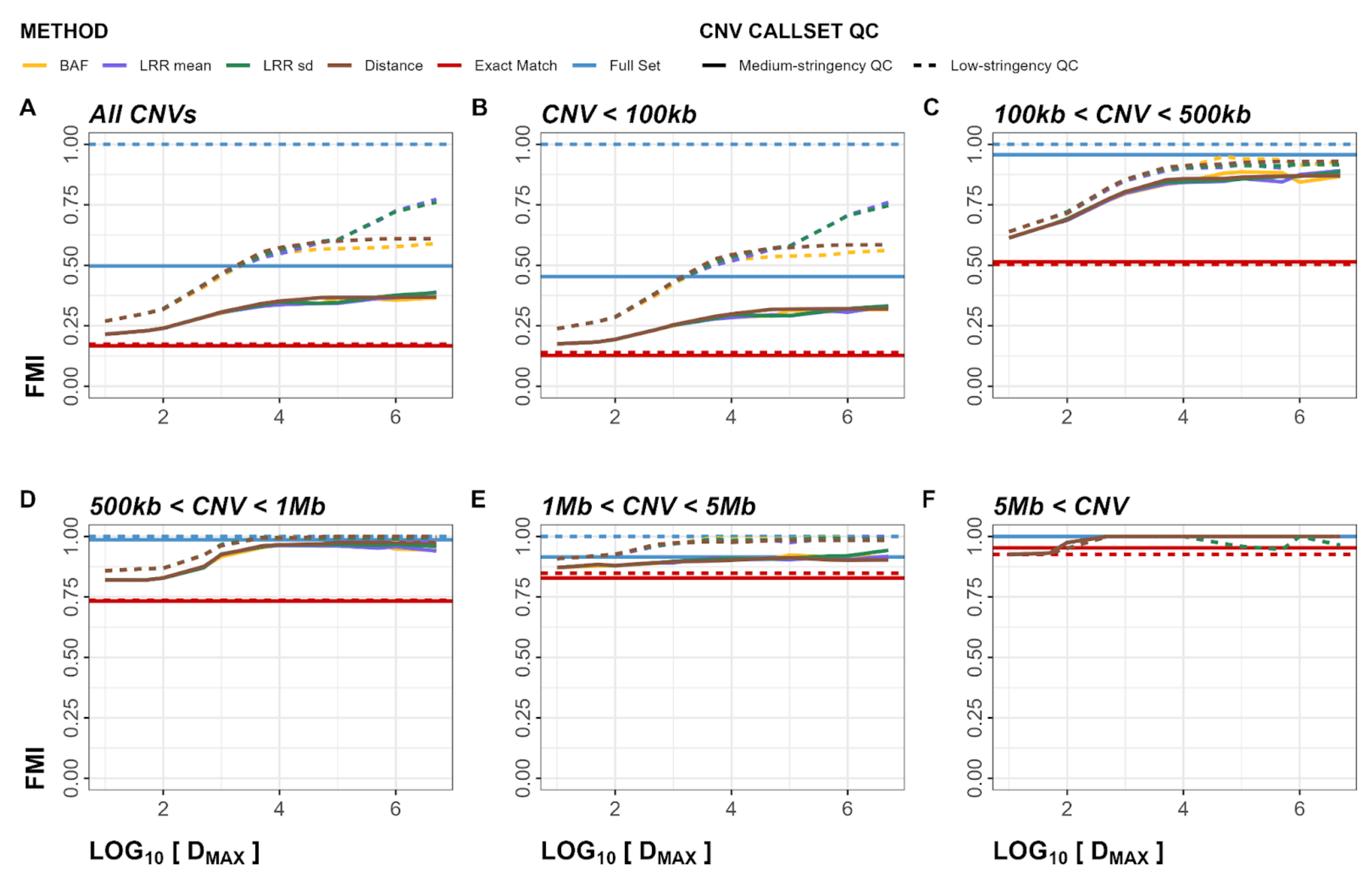


**Figure S10**. Within-Array Experiment Fowlkes-Mallows index (FMI) plots for both deletions and duplications. Panels A-F show FMI metrics stratified by CNV size. Dashed line represents low-stringency QC, solid line represents medium-stringency QC.

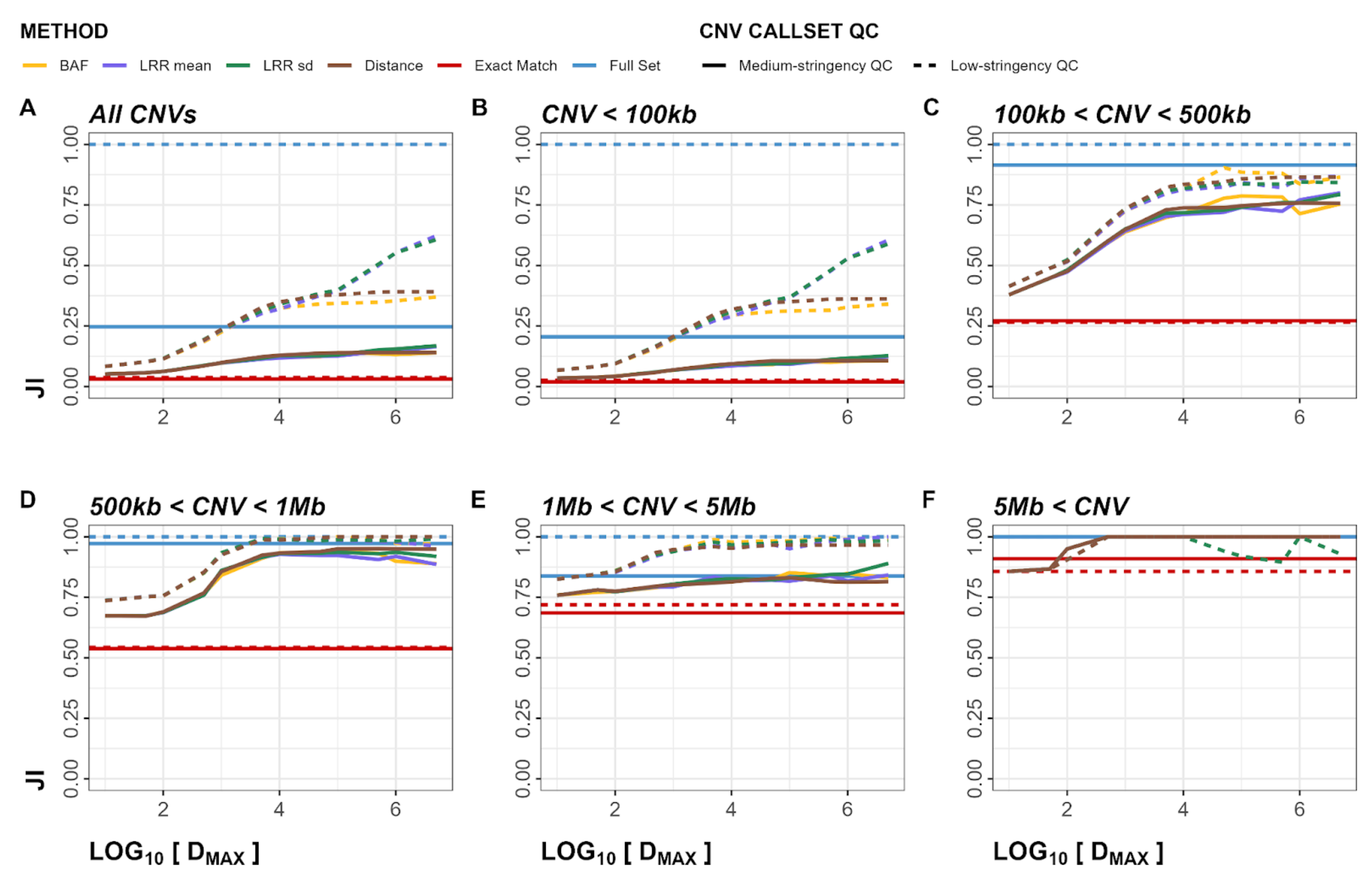


**Figure S11**. Within-Array Experiment Jaccard index (JI) plots for both deletions and duplications. Panels A-F show JI metrics stratified by CNV size. Dashed line represents low-stringency QC, solid line represents medium-stringency QC.

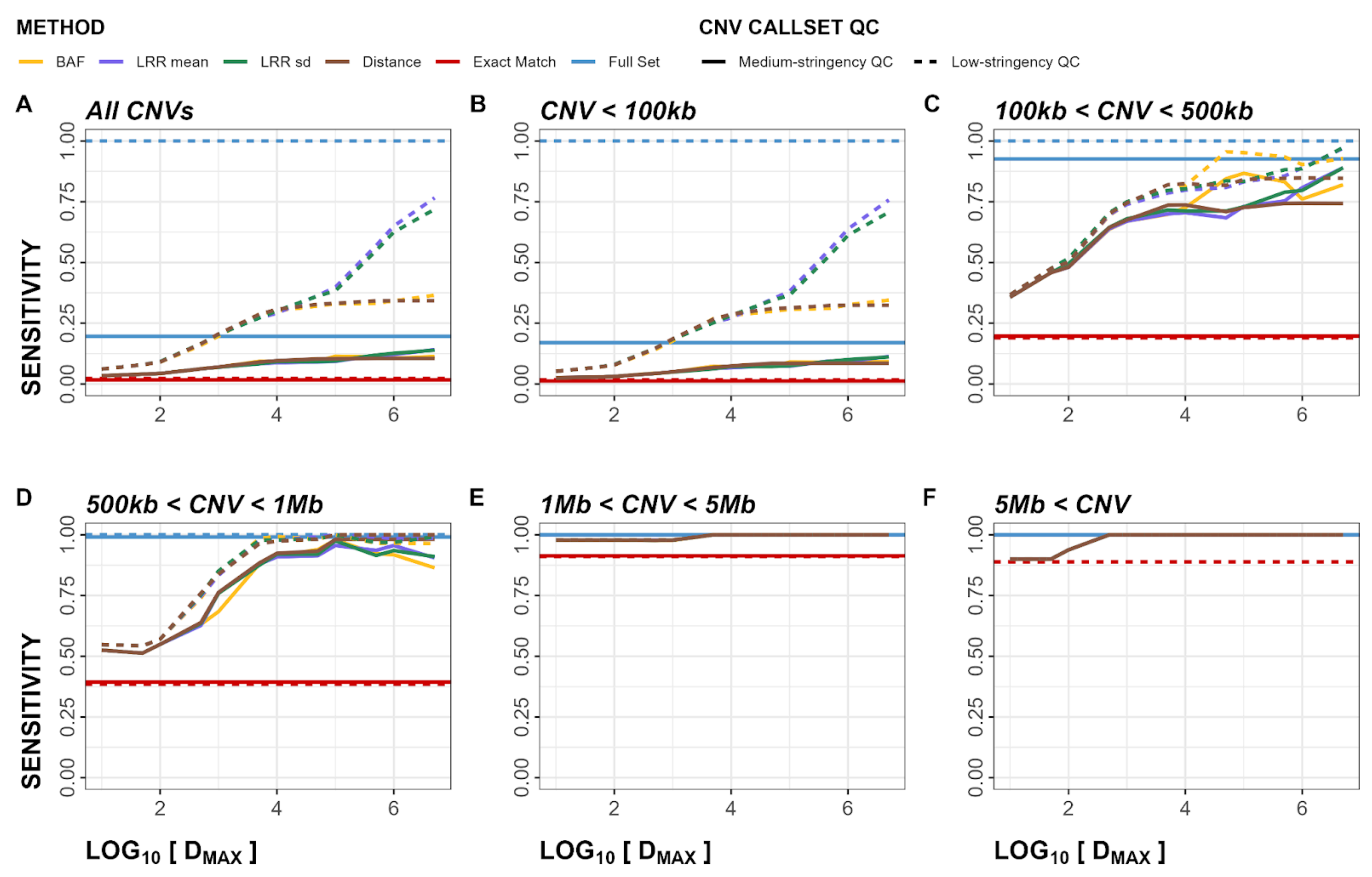


**Figure S12**. Within-Array Experiment sensitivity plots for deletions. Panels A-F show sensitivity metrics stratified by CNV size. Dashed line represents low-stringency QC, solid line represents medium-stringency QC.

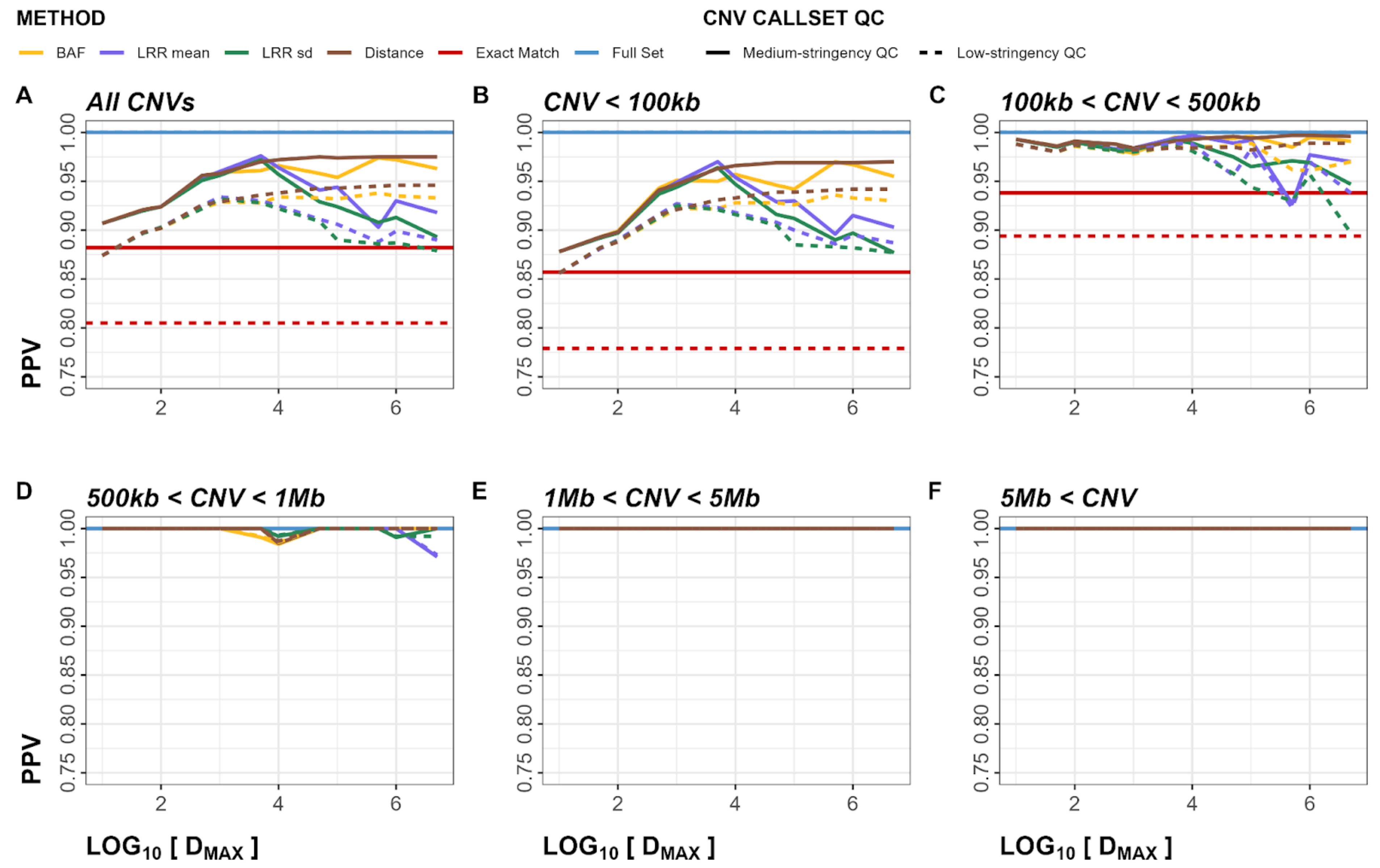


**Figure S13**. Within-Array Experiment positive predictive values (PPV) plots for deletions. Panels A-F show PPV metrics stratified by CNV size. Dashed line represents low-stringency QC, solid line represents medium-stringency QC.

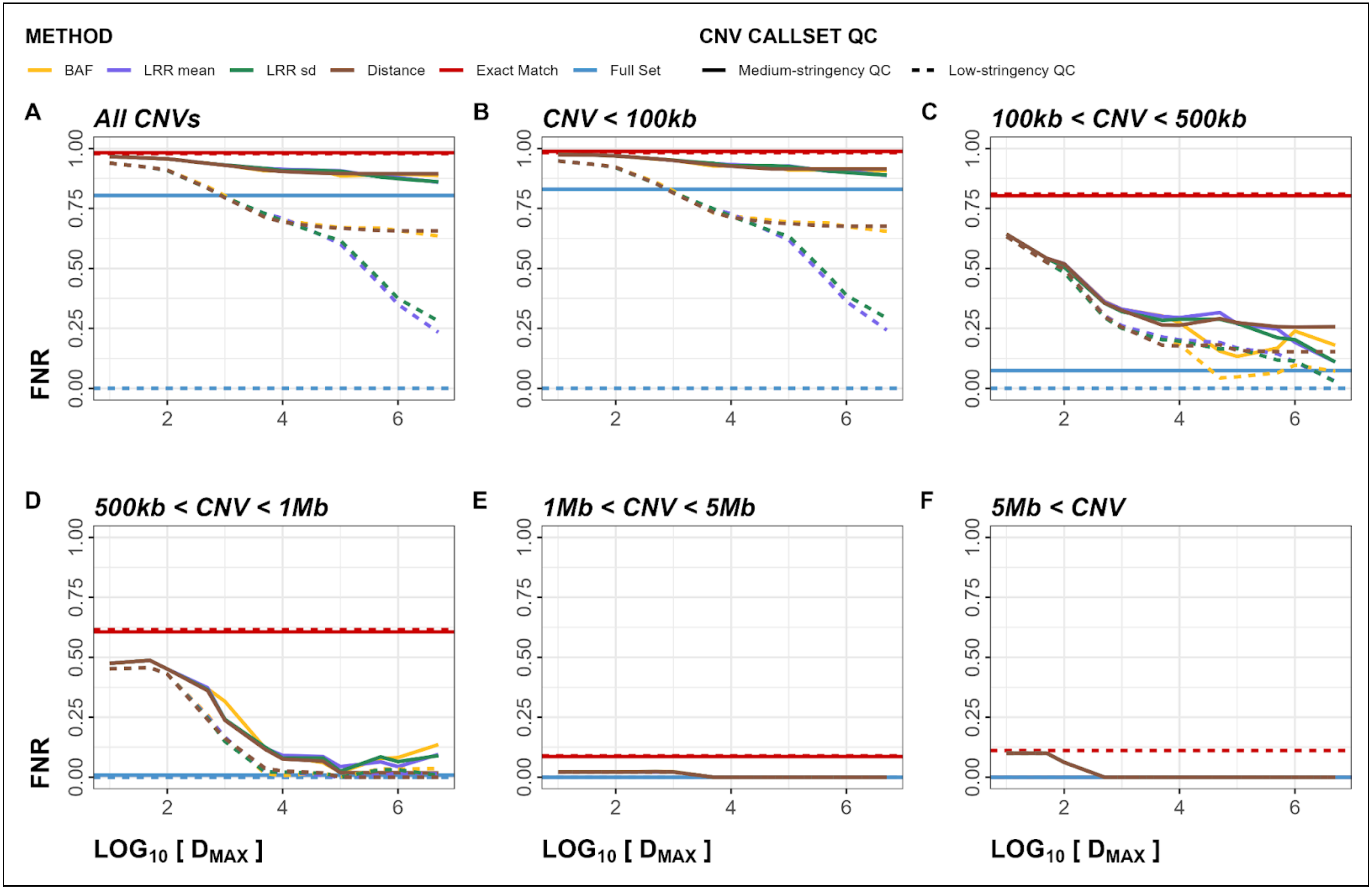


**Figure S14**. Within-Array Experiment false negative rate (FNR) plots for deletions. Panels A-F show FNR metrics stratified by CNV size. Dashed line represents low-stringency QC, solid line represents medium-stringency QC.

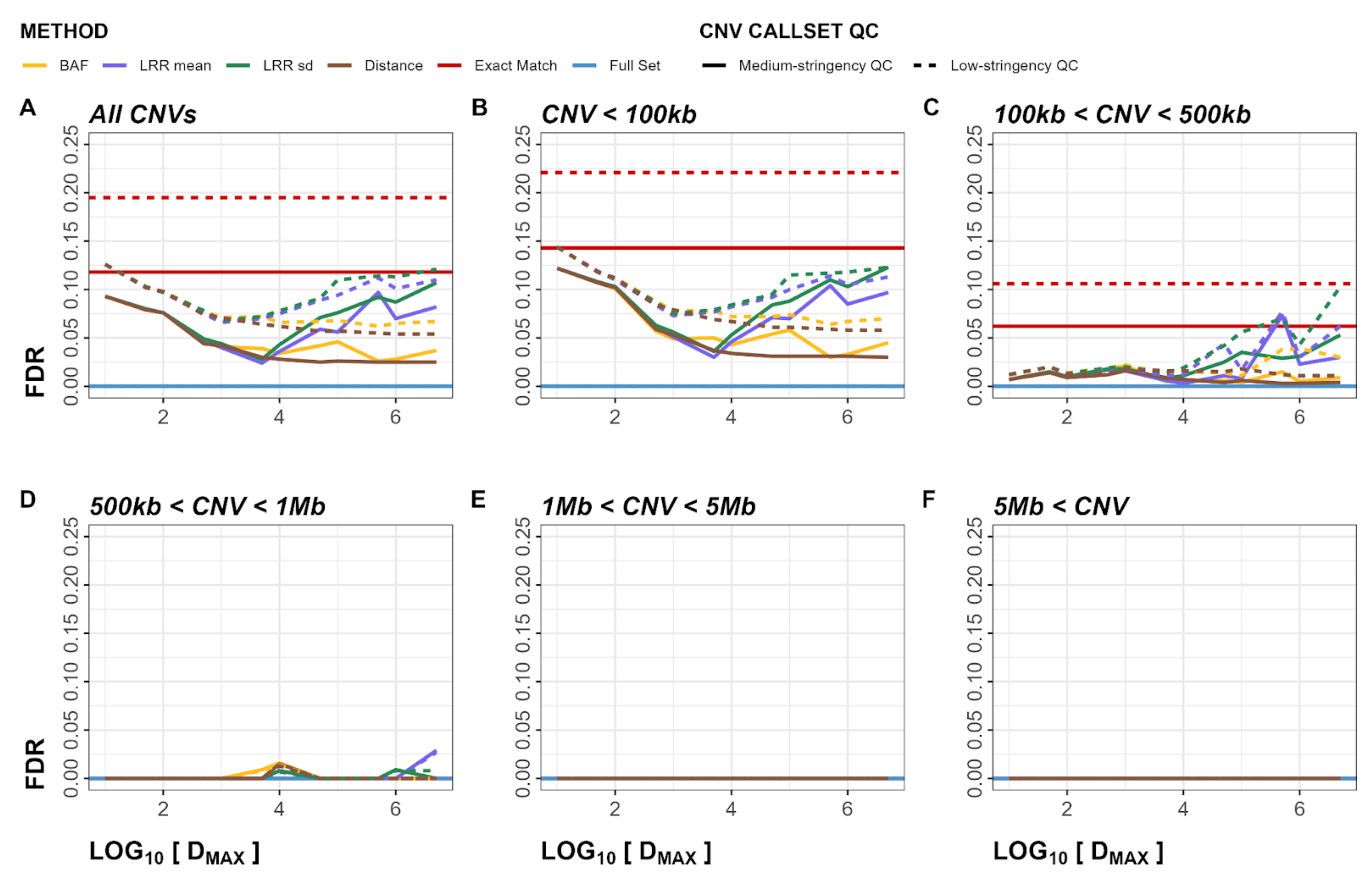


**Figure S15**. Within-Array Experiment false discovery rate (FDR) plots for deletions. Panels A-F show FDR metrics stratified by CNV size. Dashed line represents low-stringency QC, solid line represents medium-stringency QC..

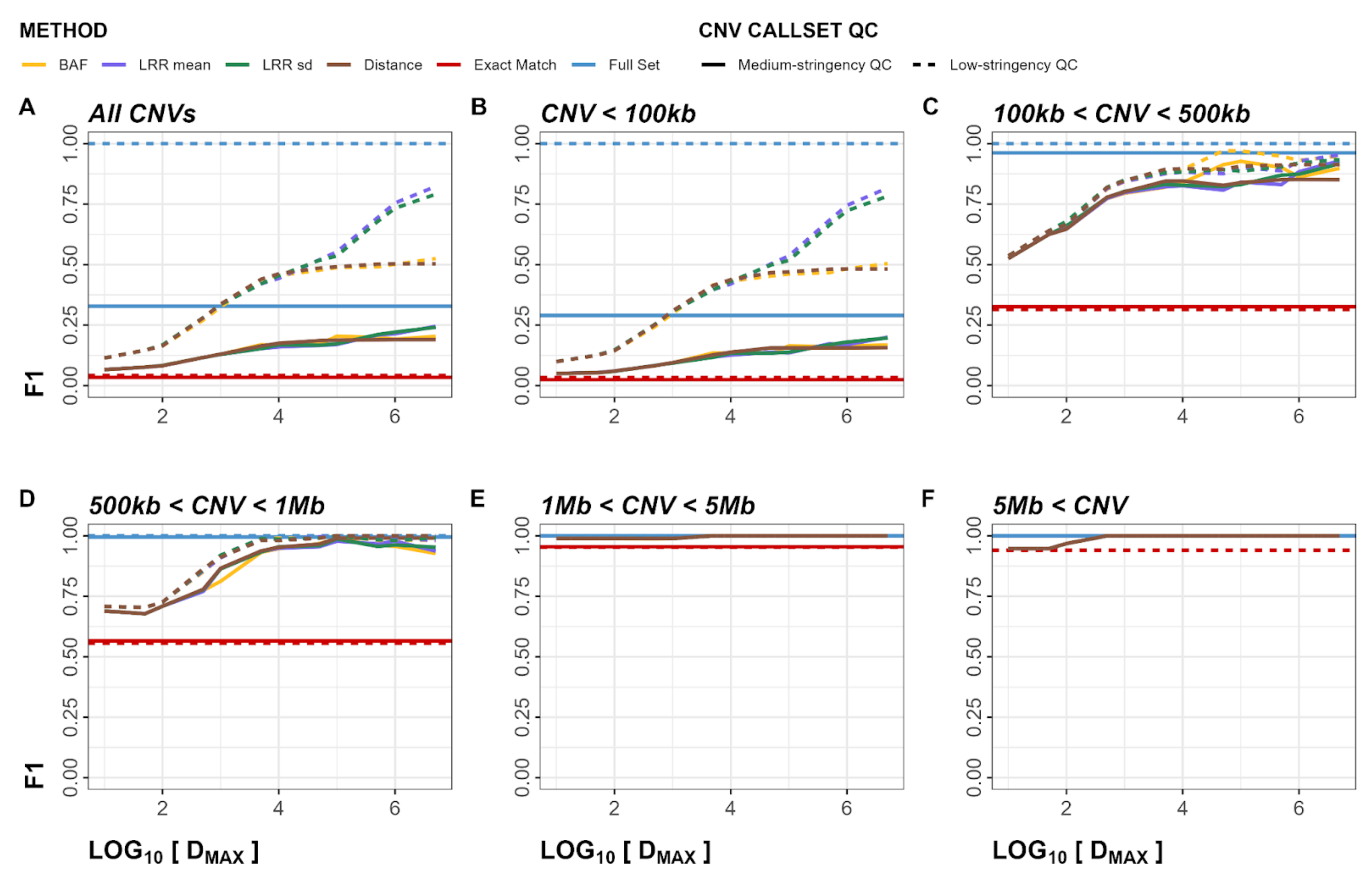


**Figure S16**. Within-Array Experiment F1 score plots for deletions. Panels A-F show F1 metrics stratified by CNV size. Dashed line represents low-stringency QC, solid line represents medium-stringency QC.

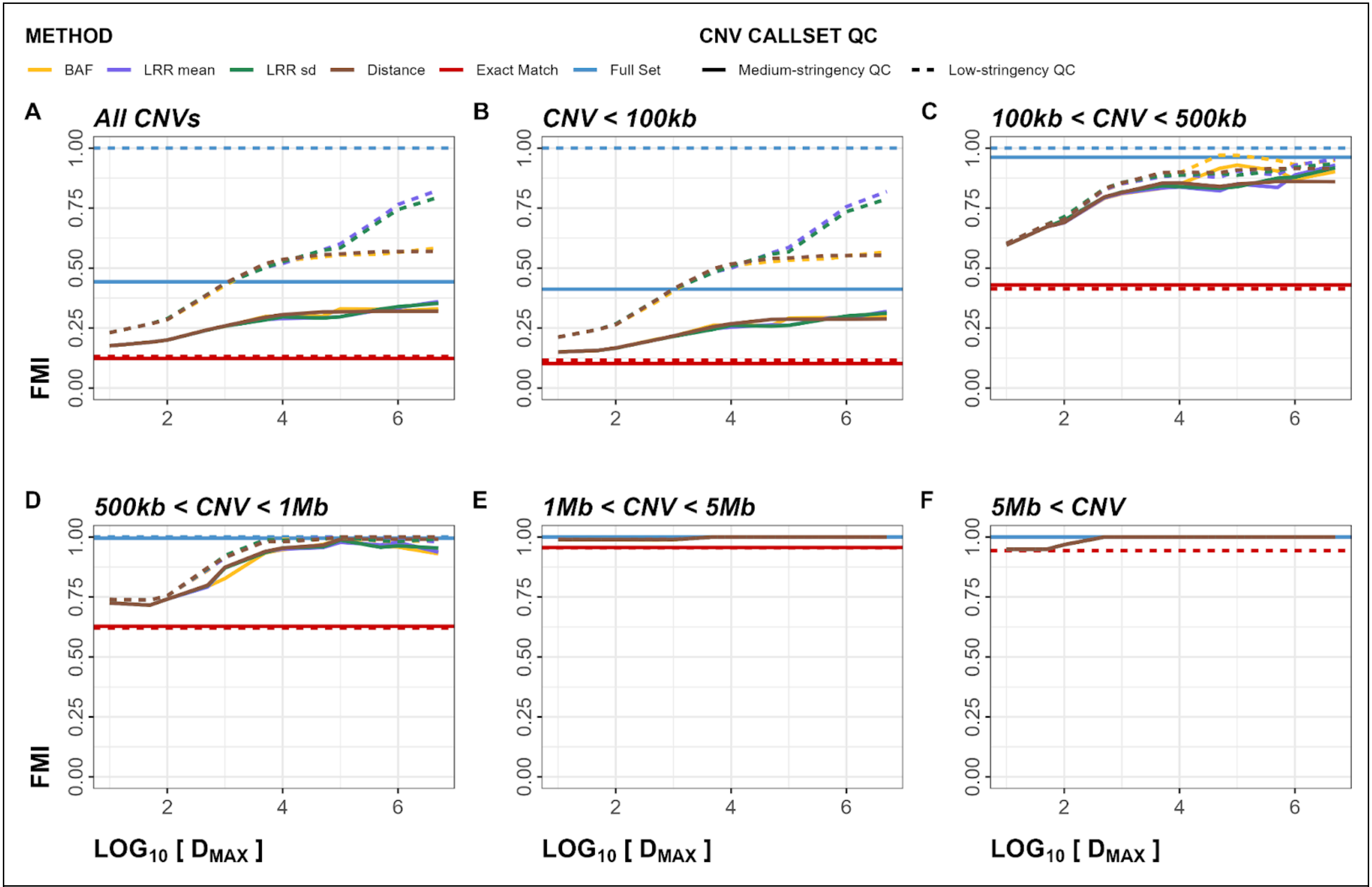


**Figure S17**. Within-Array Experiment Fowlkes-Mallows index (FMI) plots for deletions. Panels A-F show FMI metrics stratified by CNV size. Dashed line represents low-stringency QC, solid line represents medium-stringency QC.

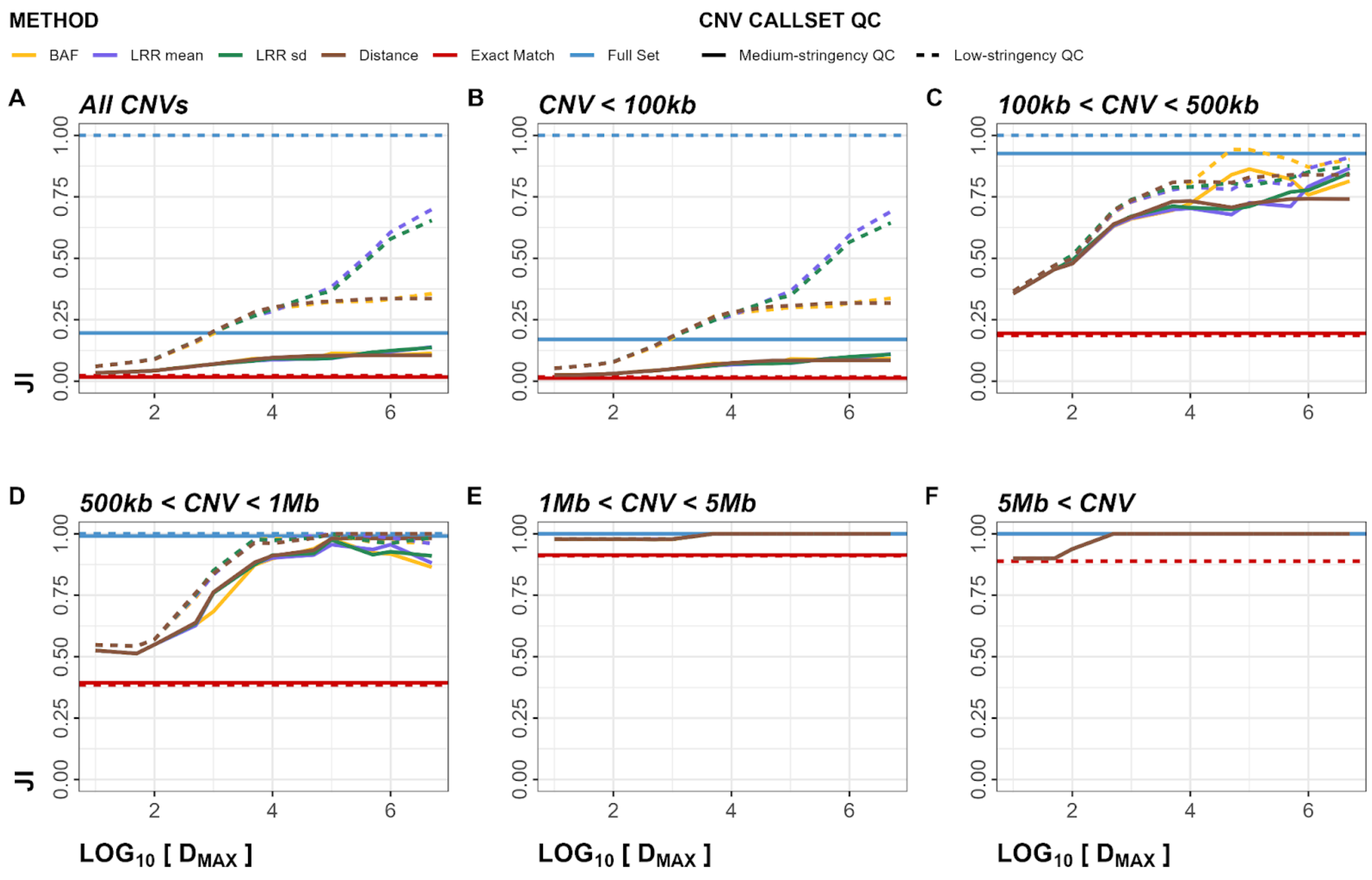


**Figure S18**. Within-Array Experiment Jaccard index (JI) plots for deletions. Panels A-F show JI metrics stratified by CNV size. Dashed line represents low-stringency QC, solid line represents medium-stringency QC.

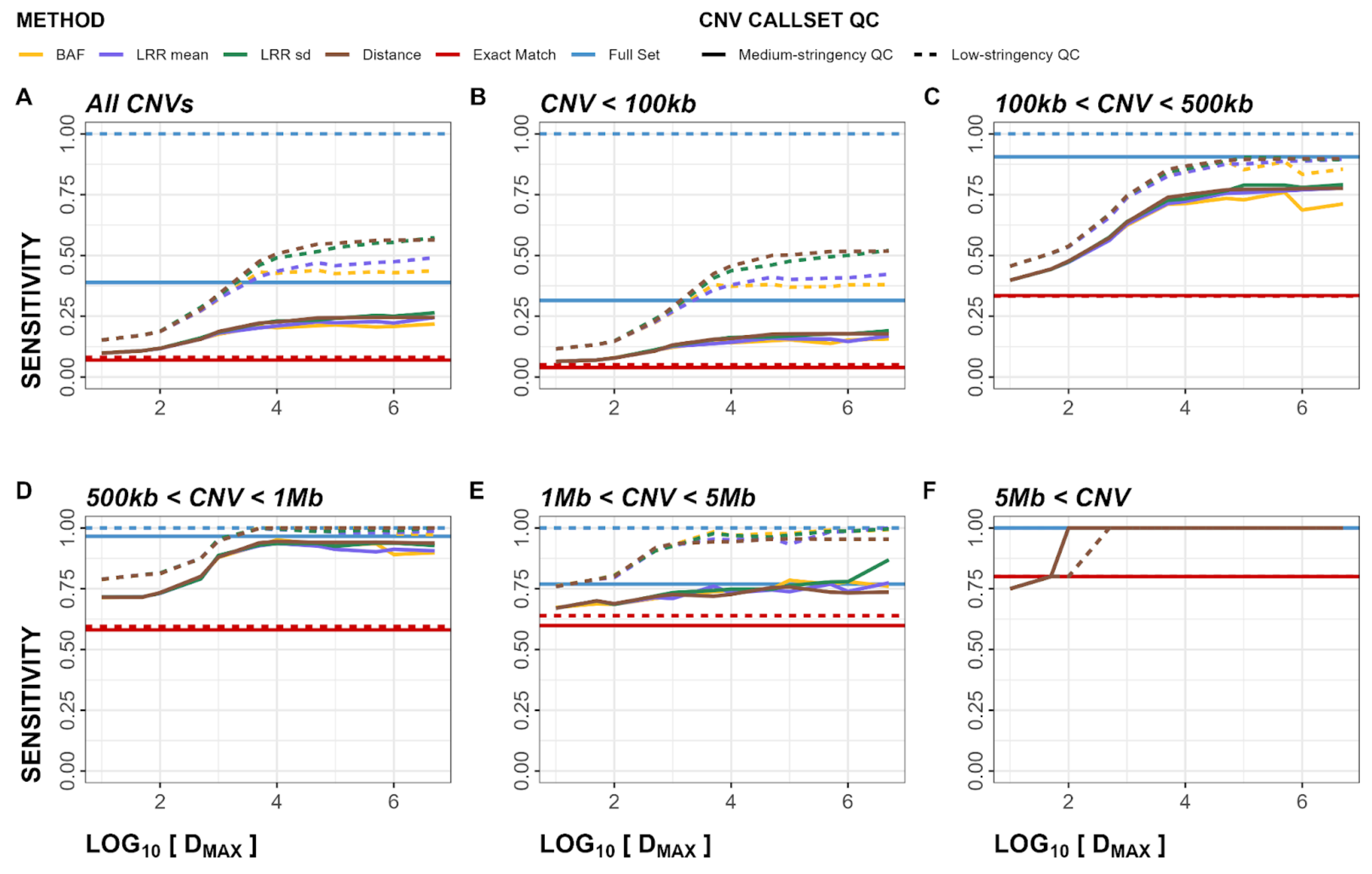


**Figure S19**. Within-Array Experiment sensitivity plots for duplications. Panels A-F show sensitivity metrics stratified by CNV size. Dashed line represents low-stringency QC, solid line represents medium-stringency QC.

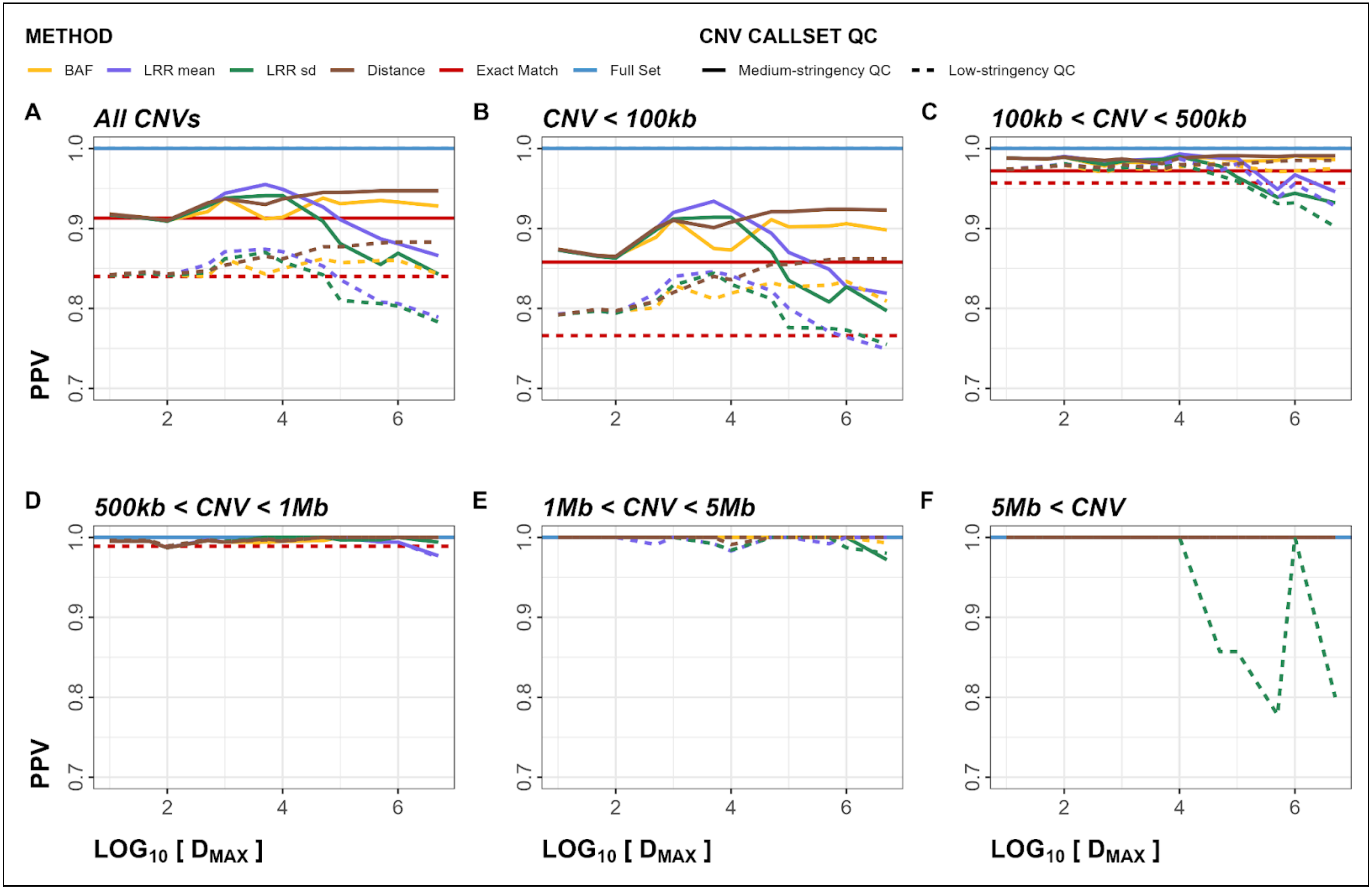


**Figure S20**. Within-Array Experiment positive predictive values (PPV) plots for duplications. Panels A-F show PPV metrics stratified by CNV size. Dashed line represents low-stringency QC, solid line represents medium-stringency QC.

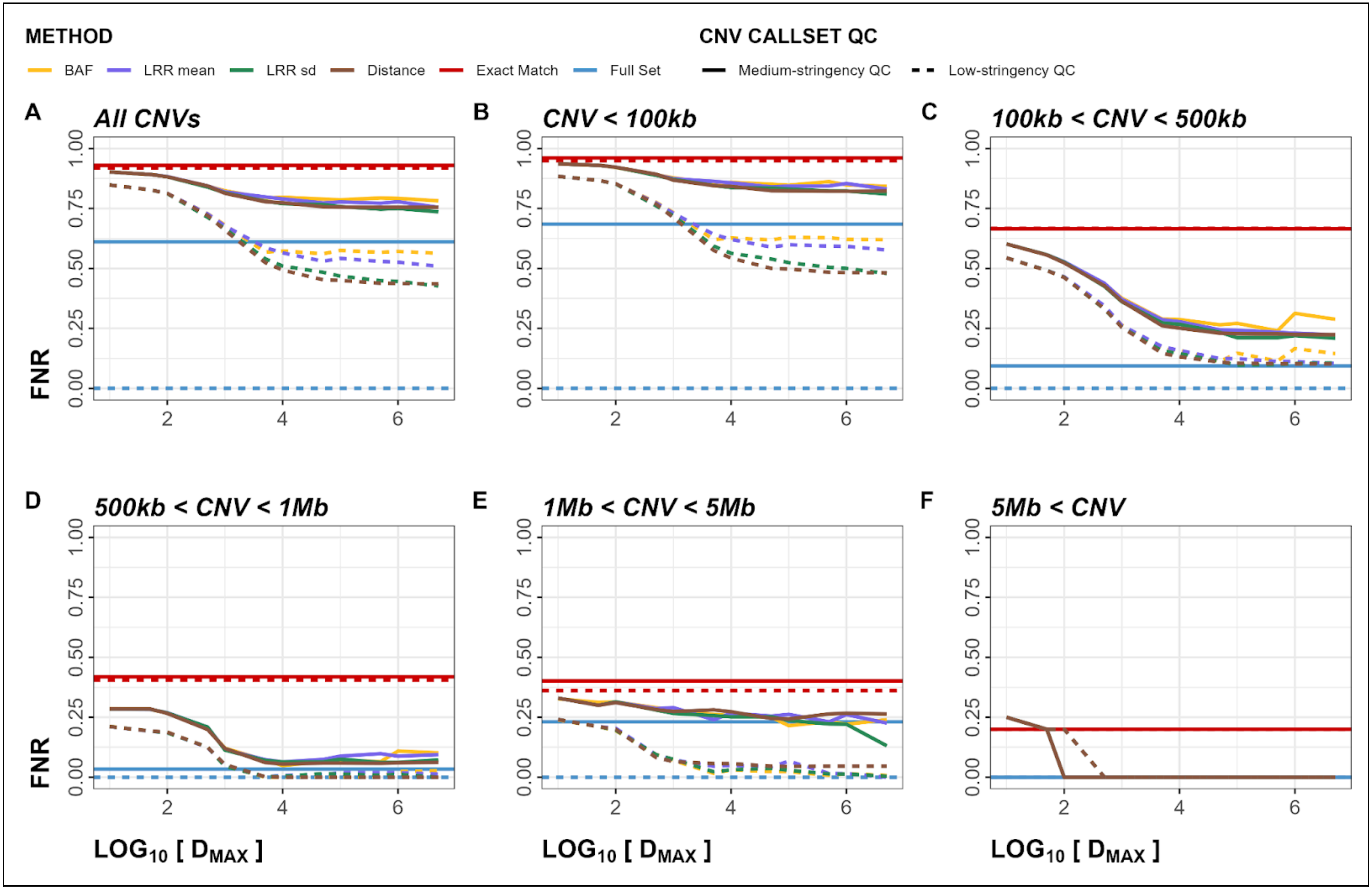


**Figure S21**. Within-Array Experiment false negative rate (FNR) plots for duplications. Panels A-F show FNR metrics stratified by CNV size. Dashed line represents low-stringency QC, solid line represents medium-stringency QC.

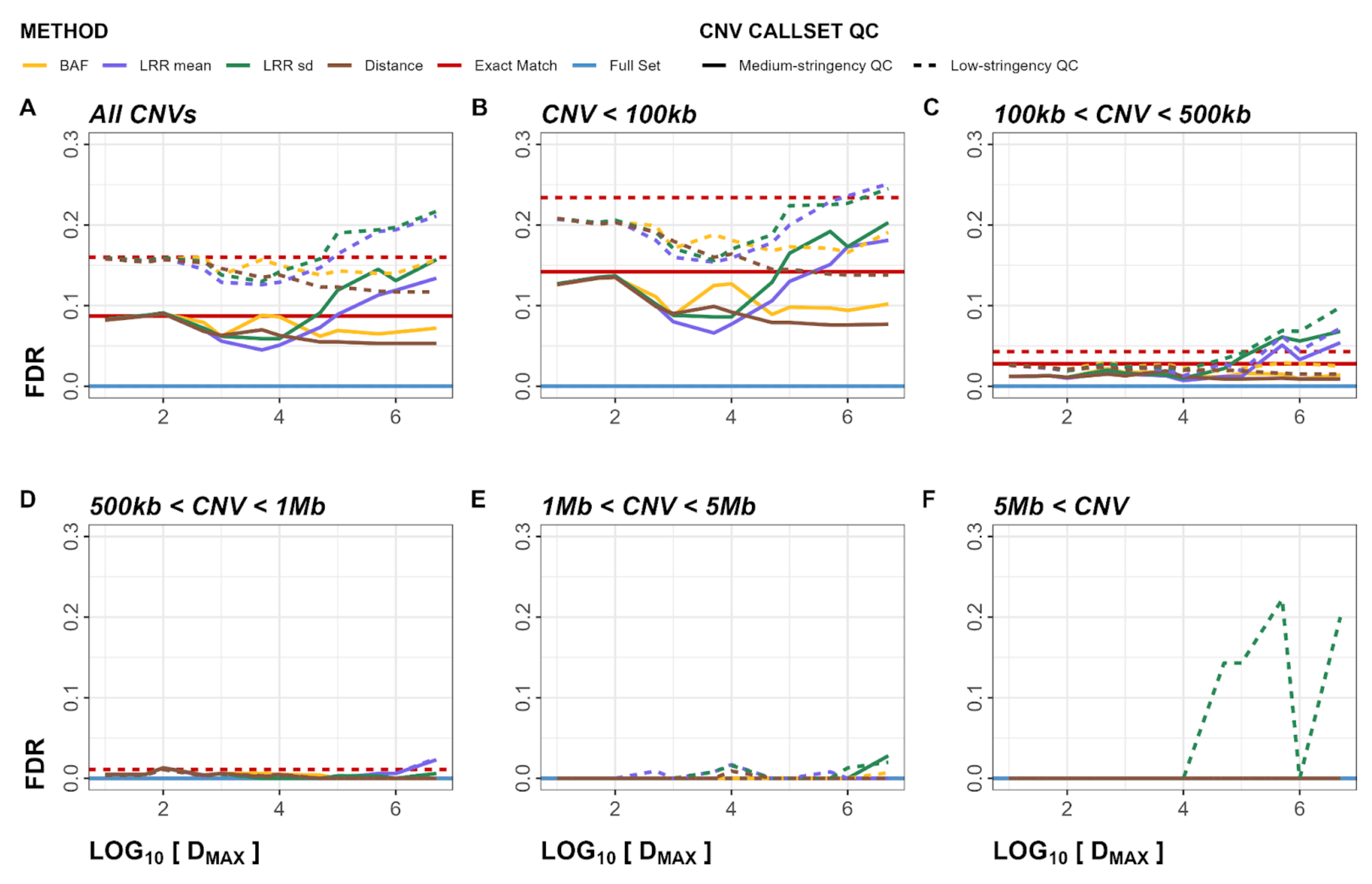


**Figure S22**. Within-Array Experiment false discovery rate (FDR) plots for duplications. Panels A-F show FDR metrics stratified by CNV size. Dashed line represents low-stringency QC, solid line represents medium-stringency QC..

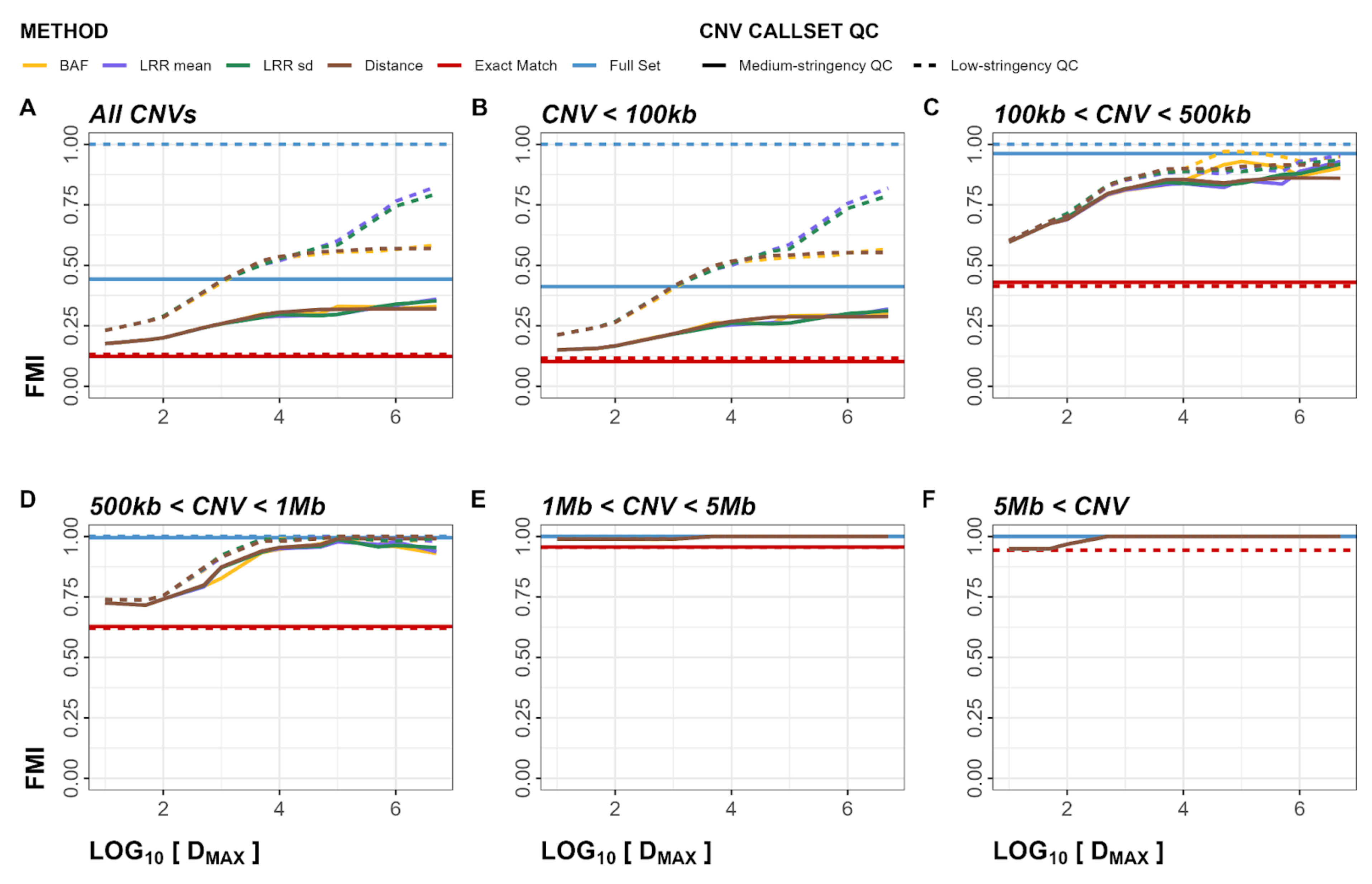


**Figure S23**. Within-Array Experiment F1 score plots for duplications. Panels A-F show F1 metrics stratified by CNV size. Dashed line represents low-stringency QC, solid line represents medium-stringency QC.

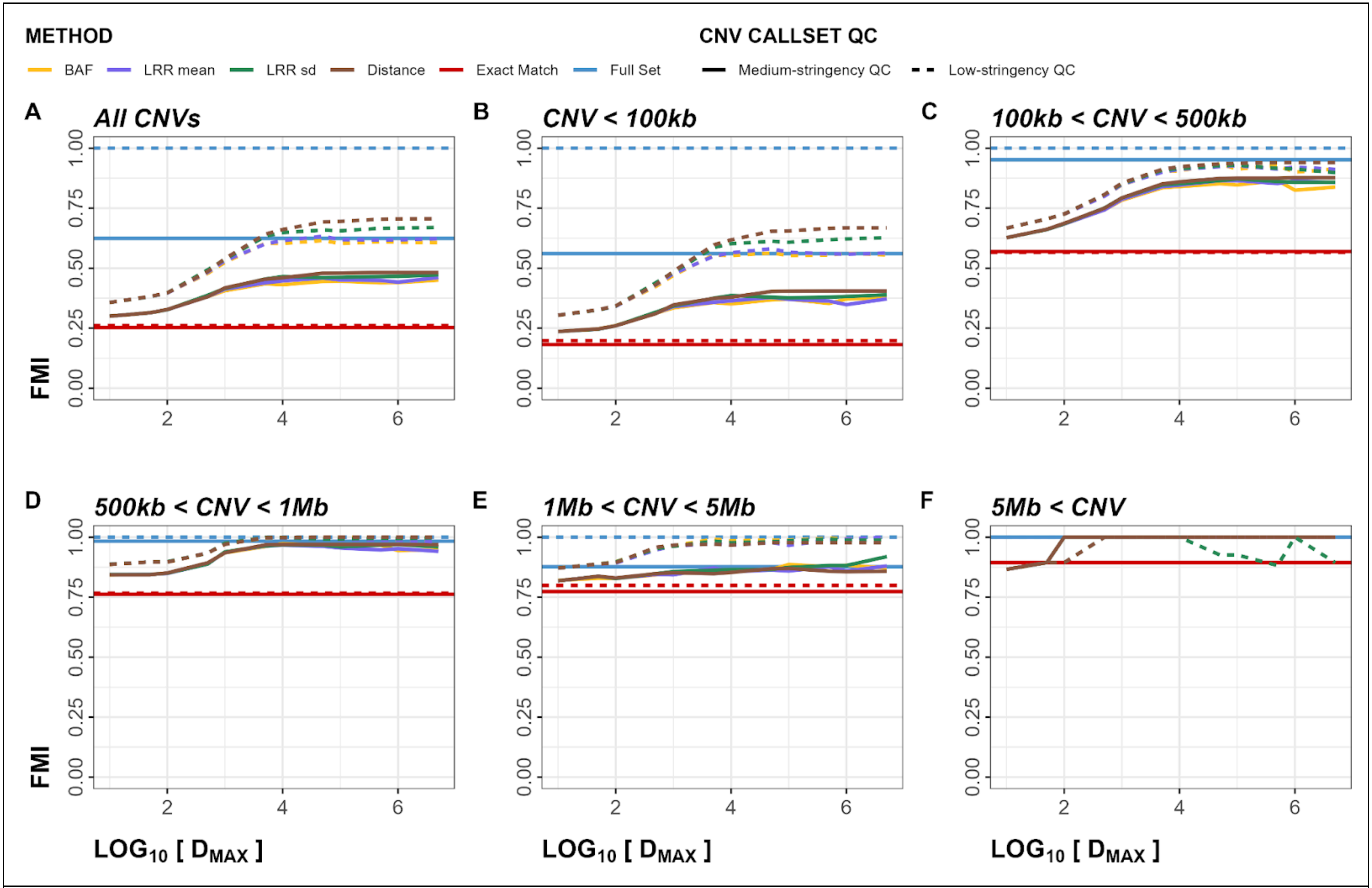


**Figure S24**. Within-Array Experiment Fowlkes-Mallows index (FMI) plots for duplications. Panels A-F show FMI metrics stratified by CNV size. Dashed line represents low-stringency QC, solid line represents medium-stringency QC.

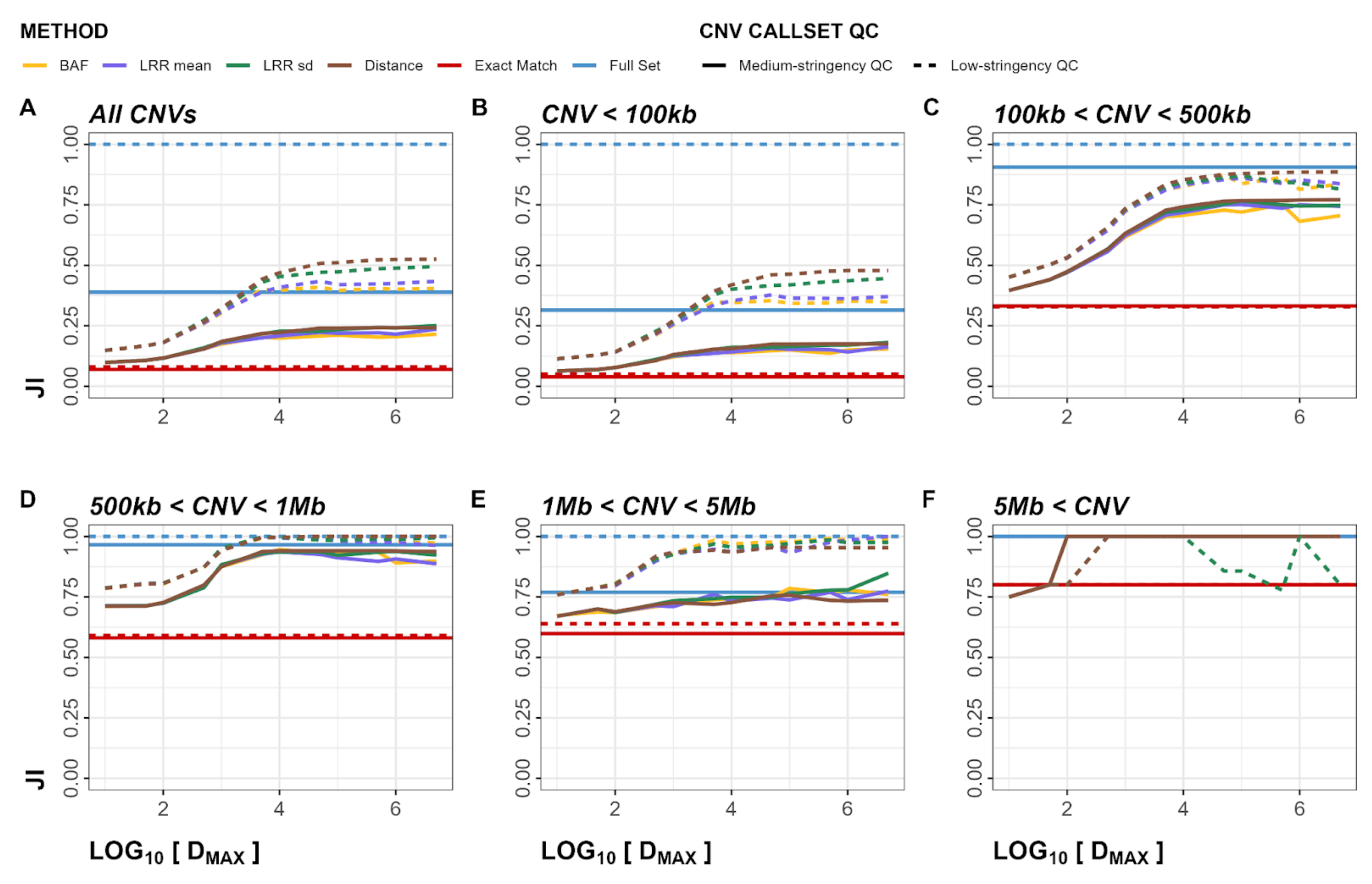


**Figure S25**. Within-Array Experiment Jaccard index (JI) plots for duplications. Panels A-F show JI metrics stratified by CNV size. Dashed line represents low-stringency QC, solid line represents medium-stringency QC.

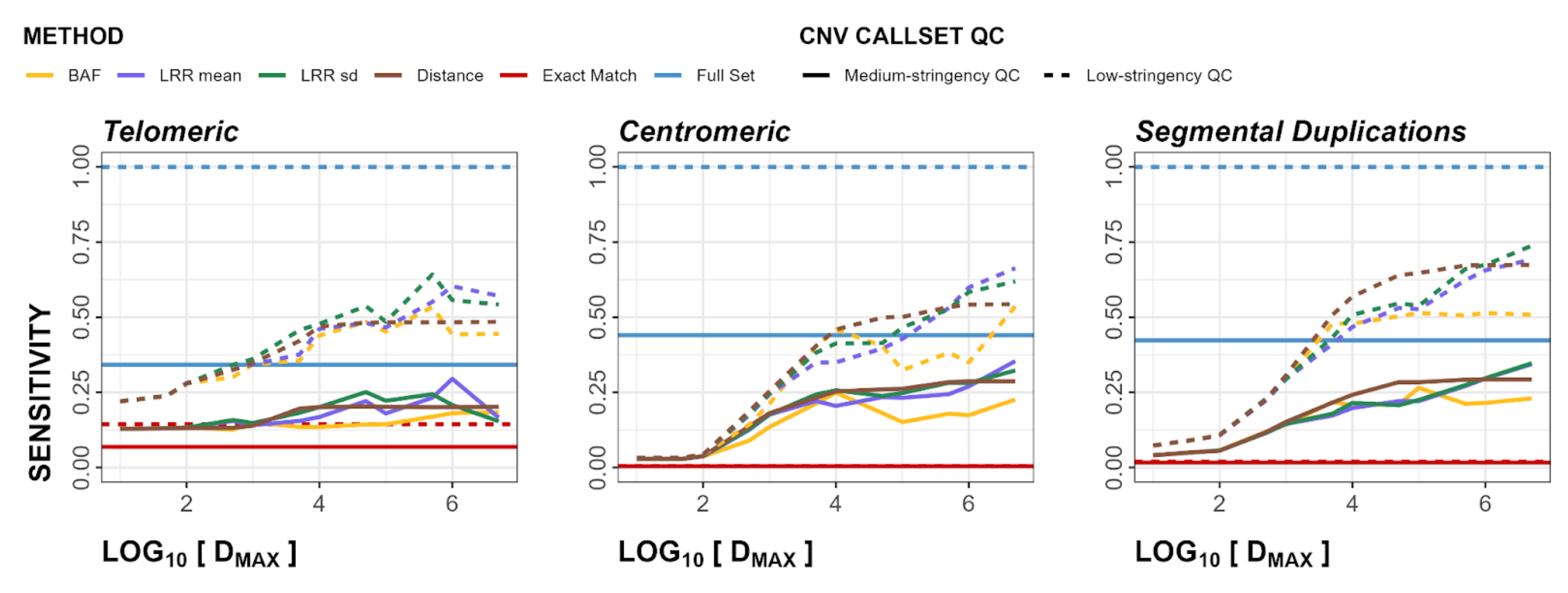


**Figure S26**. Within-Array Experiment sensitivity regional plots for both deletions and duplications. Panels show telomeric, centromeric, and segmental duplication regions in order, left to right. Dashed line represents low-stringency QC, solid line represents medium stringency QC.

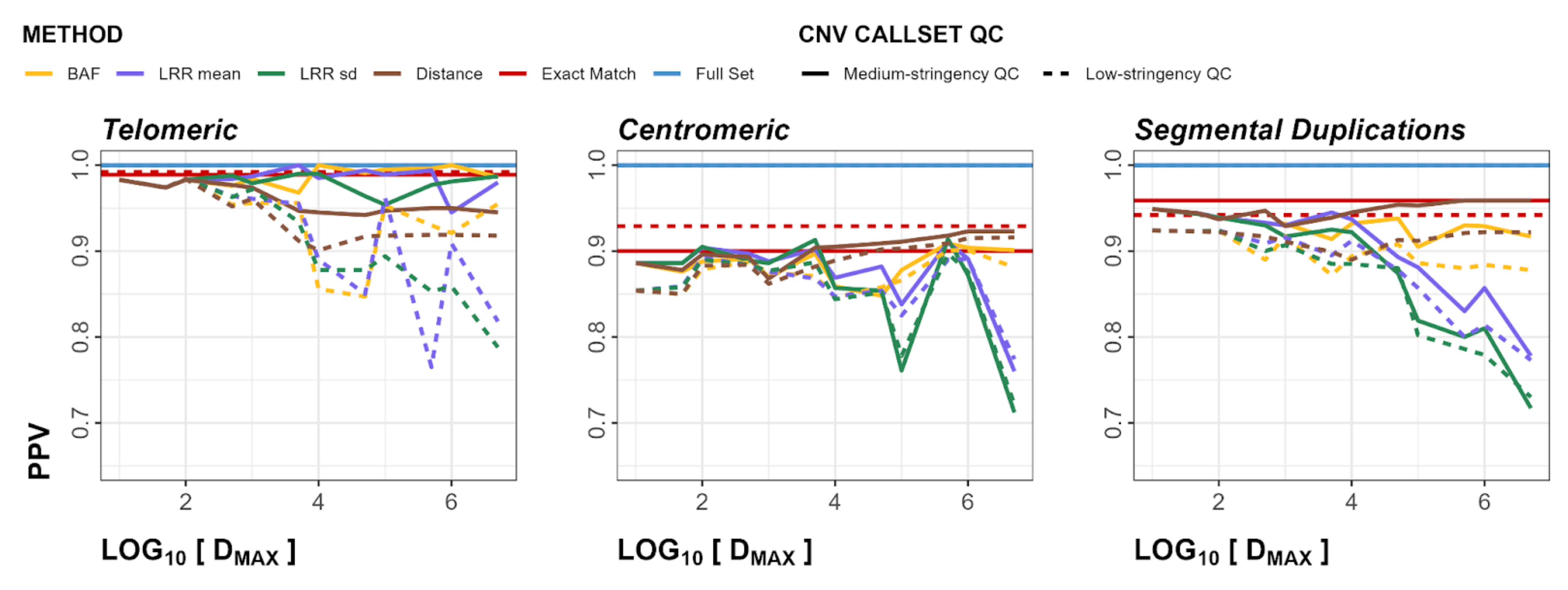


**Figure S27**. Within-Array Experiment PPV regional plots for both deletions and duplications. Panels show telomeric, centromeric, and segmental duplication regions in order, left to right. Dashed line represents low-stringency QC, solid line represents medium stringency QC.

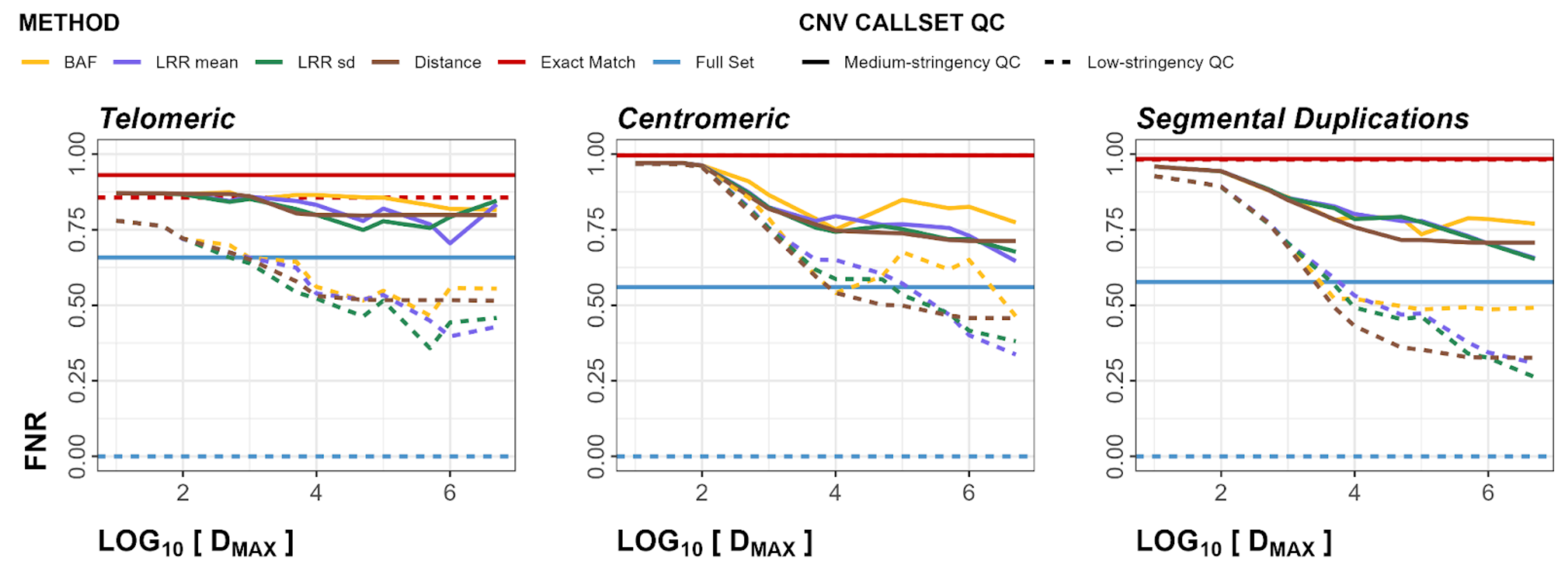


**Figure S28**. Within-Array Experiment FNR regional plots for both deletions and duplications. Panels show telomeric, centromeric, and segmental duplication regions in order, left to right. Dashed line represents low-stringency QC, solid line represents medium stringency QC.

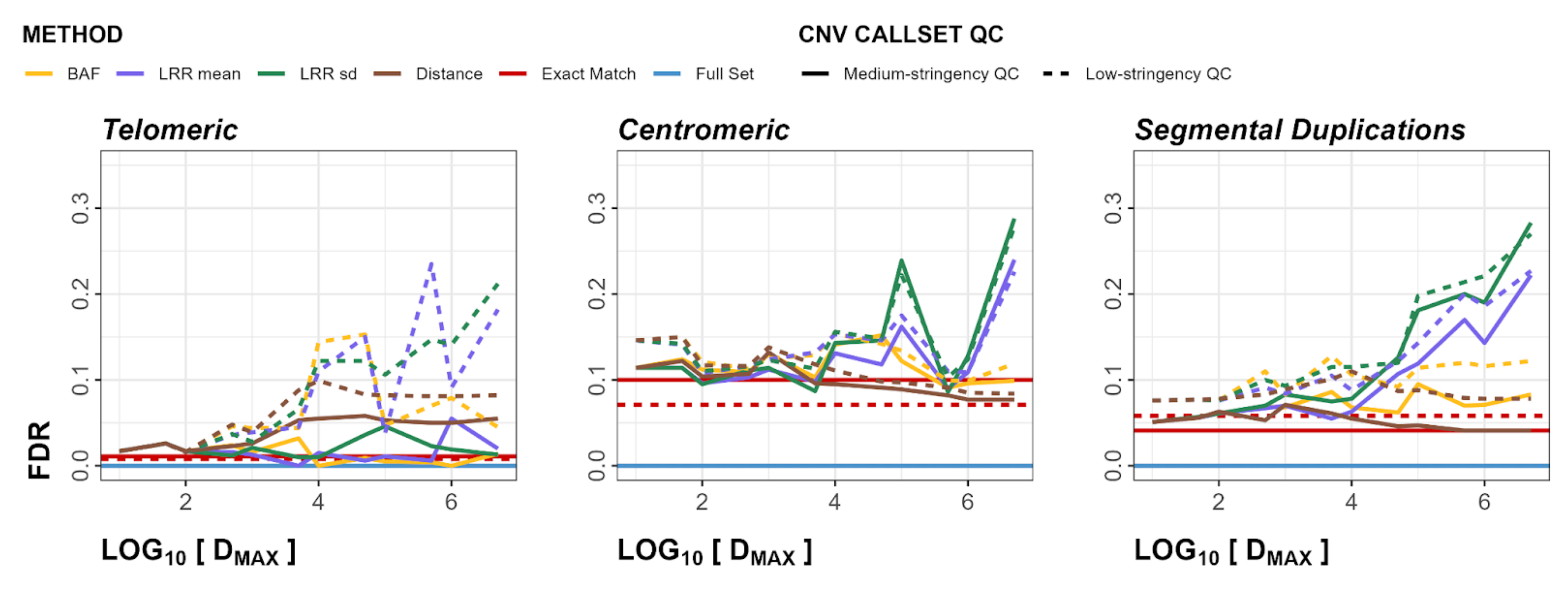


**Figure S29**. Within-Array Experiment FDR regional plots for both deletions and duplications. Panels show telomeric, centromeric, and segmental duplication regions in order, left to right. Dashed line represents low-stringency QC, solid line represents medium stringency QC.

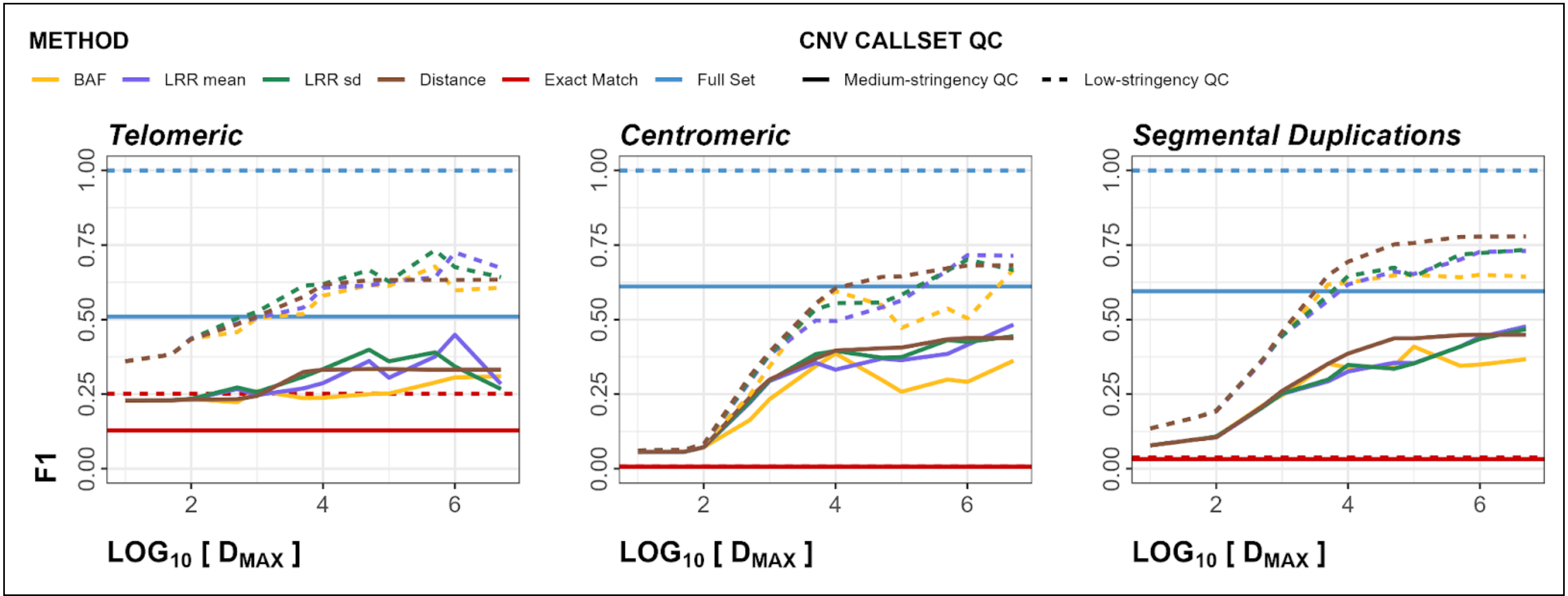


**Figure S30**. Within-Array Experiment F1 regional plots for both deletions and duplications. Panels show telomeric, centromeric, and segmental duplication regions in order, left to right. Dashed line represents low-stringency QC, solid line represents medium stringency QC.

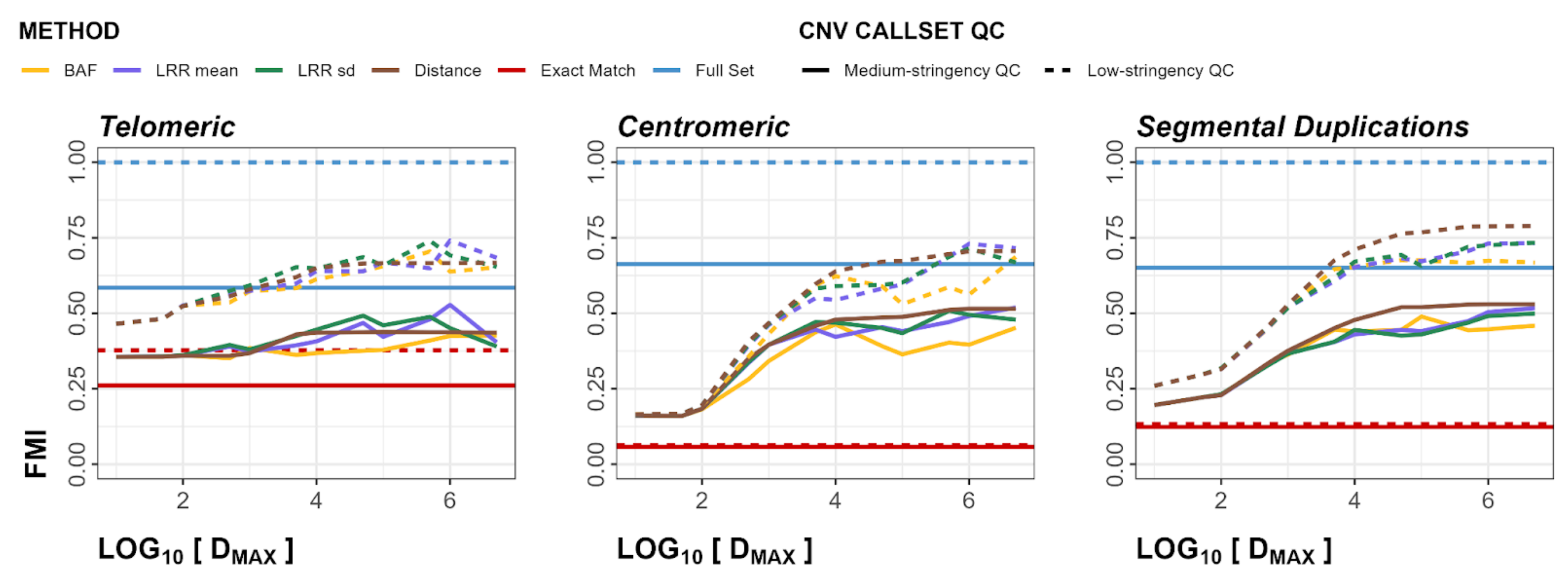


**Figure S31**. Within-Array Experiment FMI regional plots for both deletions and duplications. Panels show telomeric, centromeric, and segmental duplication regions in order, left to right. Dashed line represents low-stringency QC, solid line represents medium stringency QC.

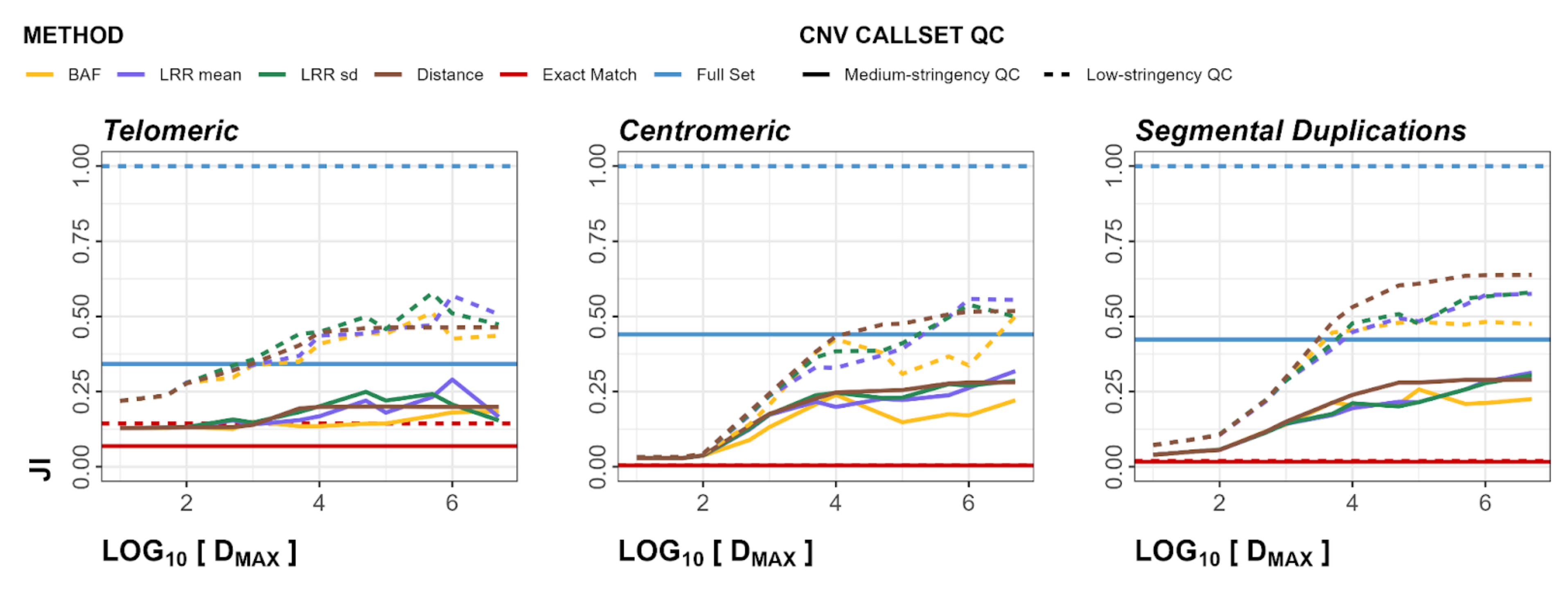


**Figure S32**. Within-Array Experiment JI regional plots for both deletions and duplications. Panels show telomeric, centromeric, and segmental duplication regions in order, left to right. Dashed line represents low-stringency QC, solid line represents medium stringency QC.

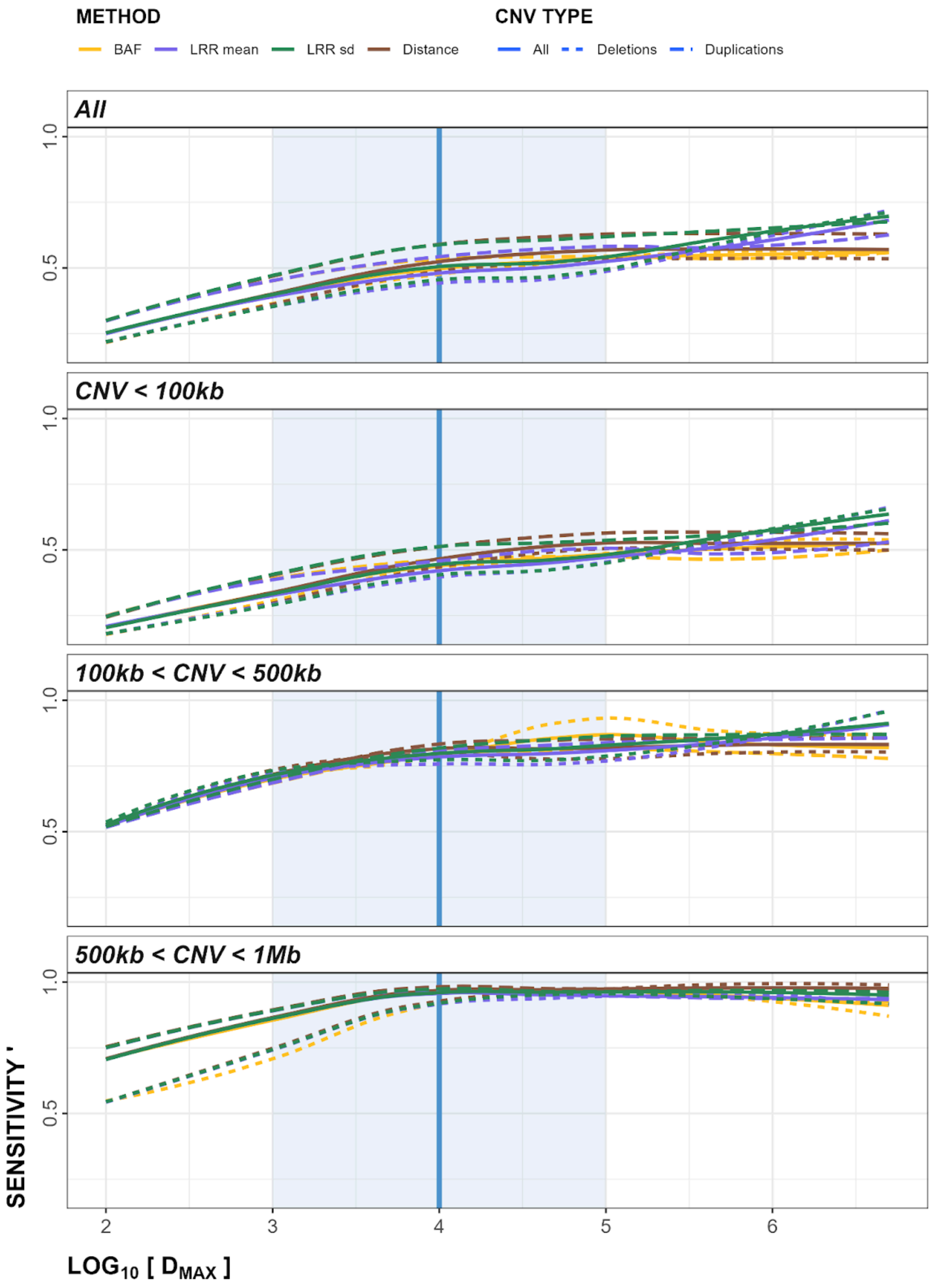


**Figure S33**. Within-Array Experiment plots of smoothed, scaled sensitivity (sensitivity') metrics to determine optimal $D_{MAX}$ parameter. The blue line represents $D_{MAX}$ = 10kb, the shaded blue region represents 1kb < $D_{MAX}$ < 100kb.

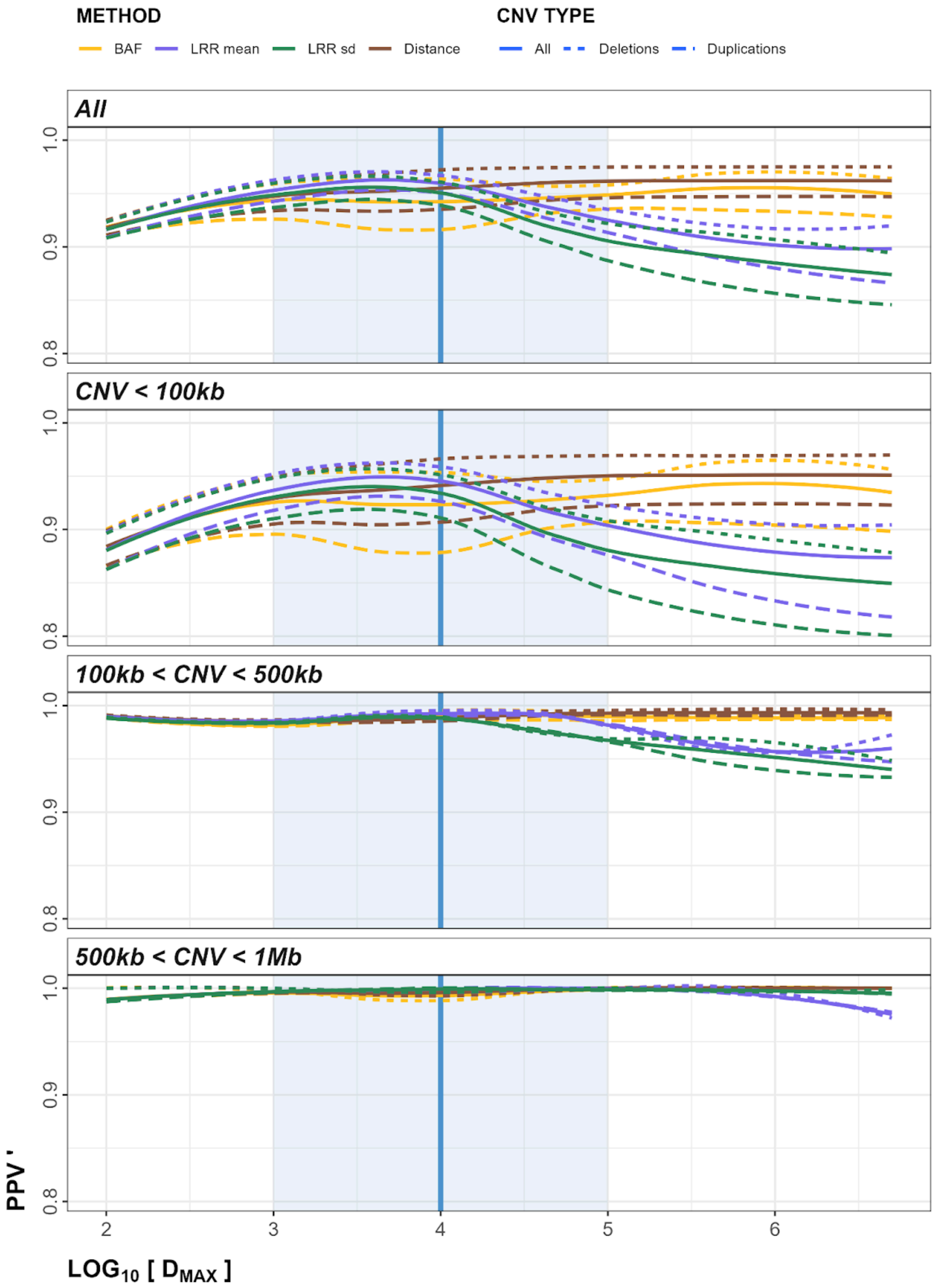


**Figure S34**. Within-Array Experiment plots of smoothed, scaled PPV (PPV') metrics to determine optimal $D_{MAX}$ parameter. The blue line represents $D_{MAX}$ = 10kb, the shaded blue region represents 1kb < $D_{MAX}$ < 100kb.

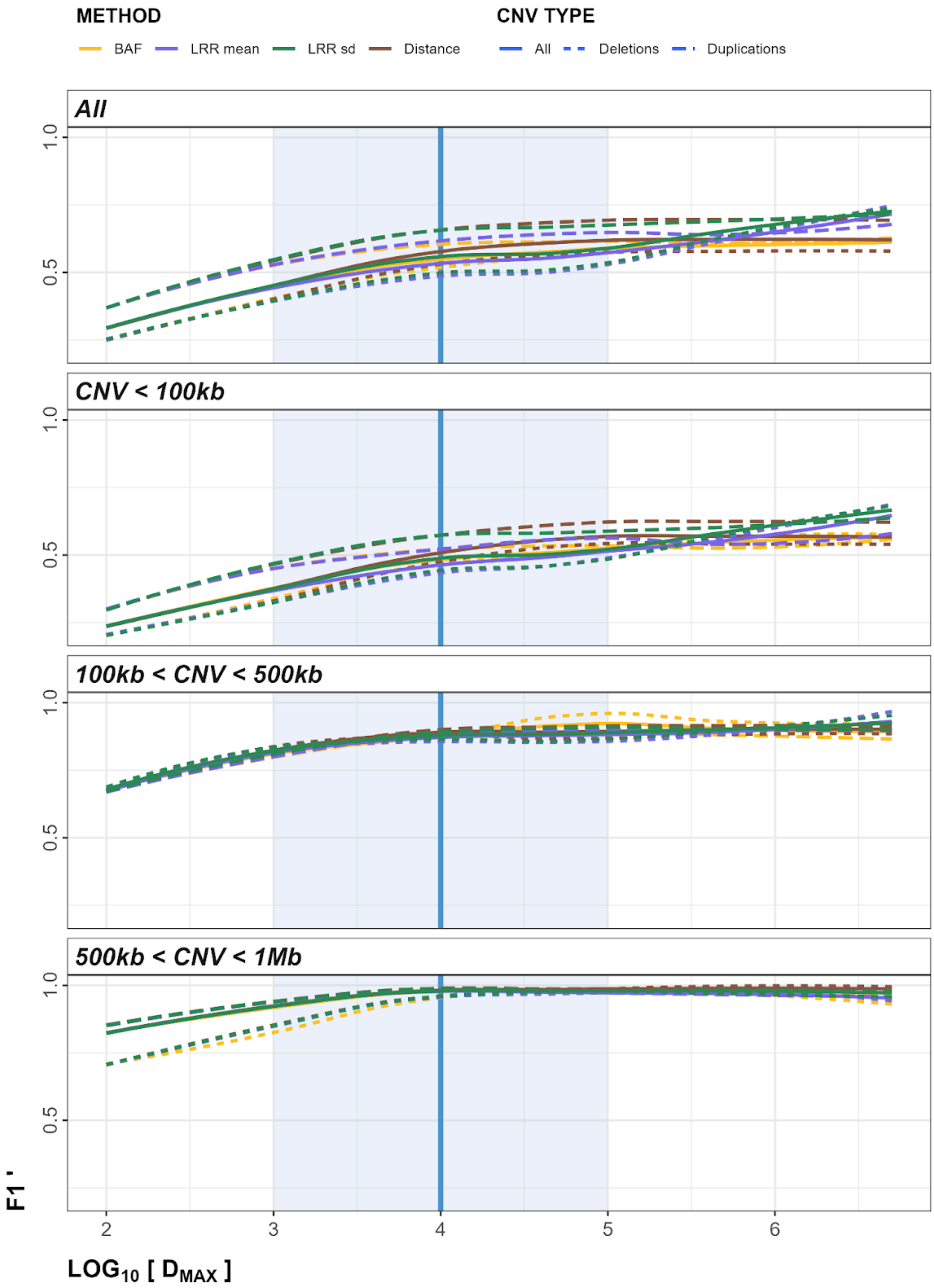


**Figure S35**. Within-Array Experiment plots of smoothed, scaled F1 (F1‘) metrics to determine optimal $D_{MAX}$ parameter. The blue line represents $D_{MAX}$ = 10kb, the shaded blue region represents 1kb < $D_{MAX}$ < 100kb.

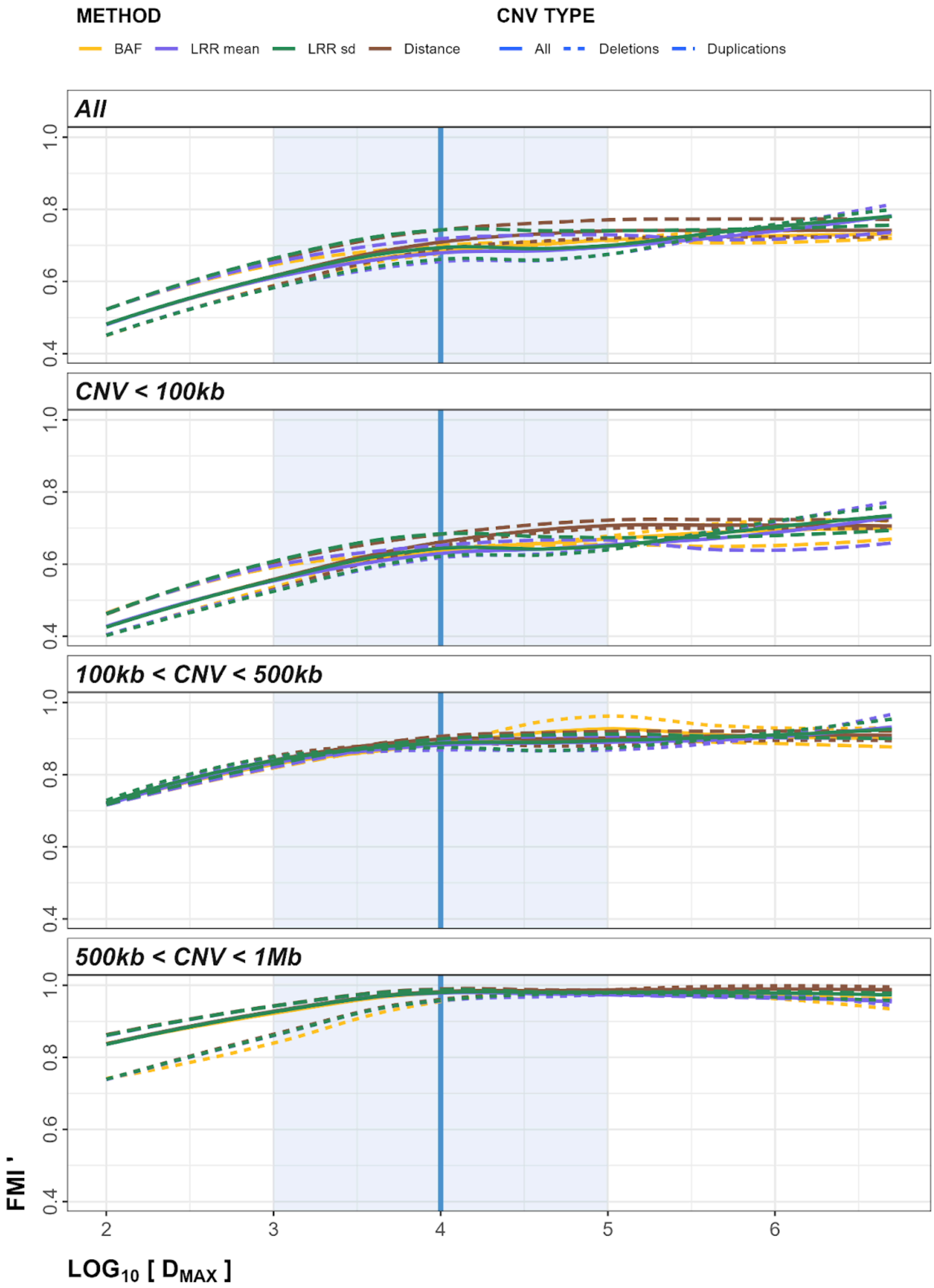


**Figure S36**. Within-Array Experiment plots of smoothed, scaled FMI (FMI') metrics to determine optimal $D_{MAX}$ parameter. The blue line represents $D_{MAX}$ = 10kb, the shaded blue region represents 1kb < $D_{MAX}$ < 100kb.

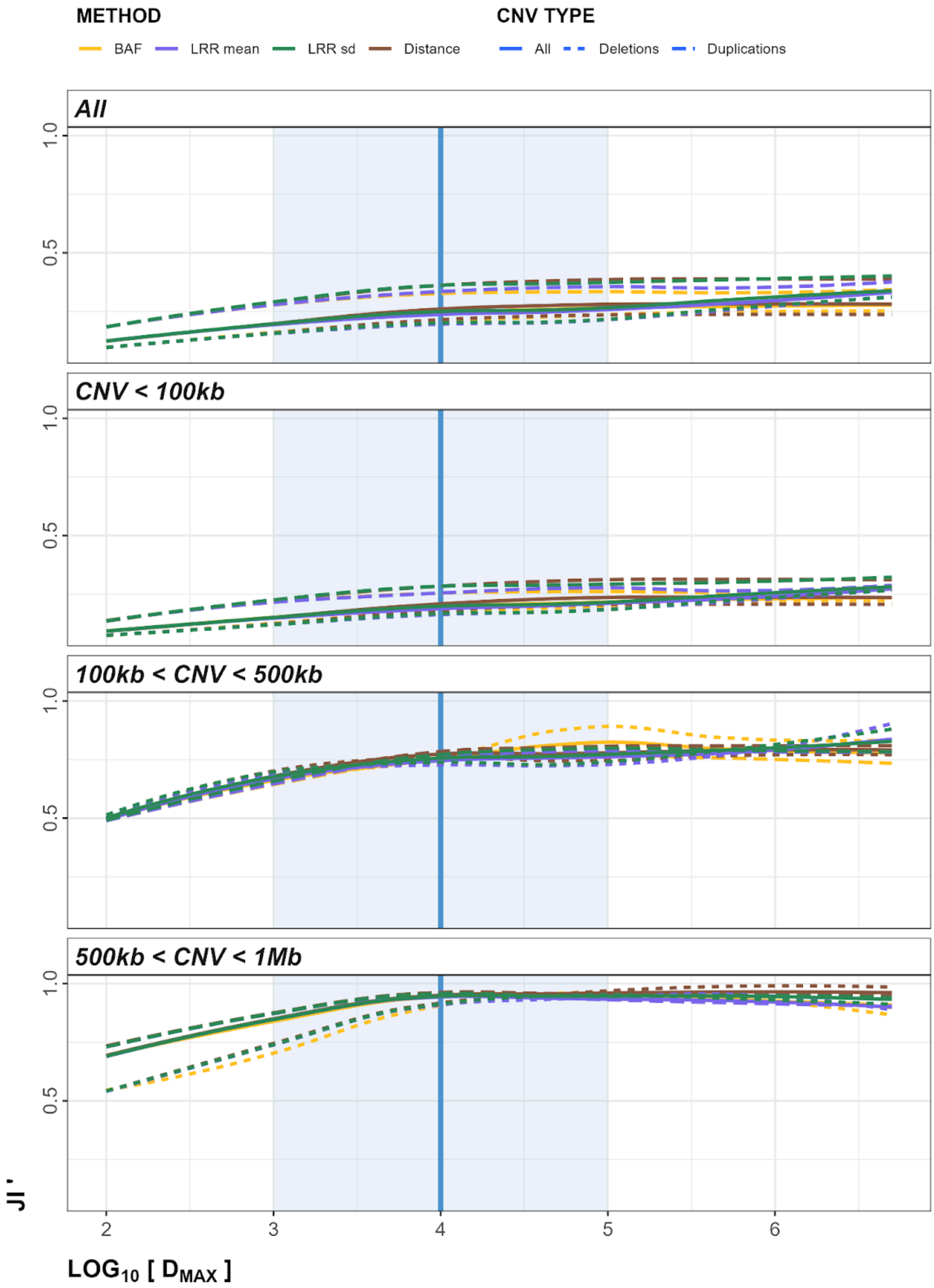


**Figure S37**. Within-Array Experiment plots of smoothed, scaled JI (JI') metrics to determine optimal $D_{MAX}$ parameter. The blue line represents $D_{MAX}$ = 10kb, the shaded blue region represents 1kb < $D_{MAX}$ < 100kb.

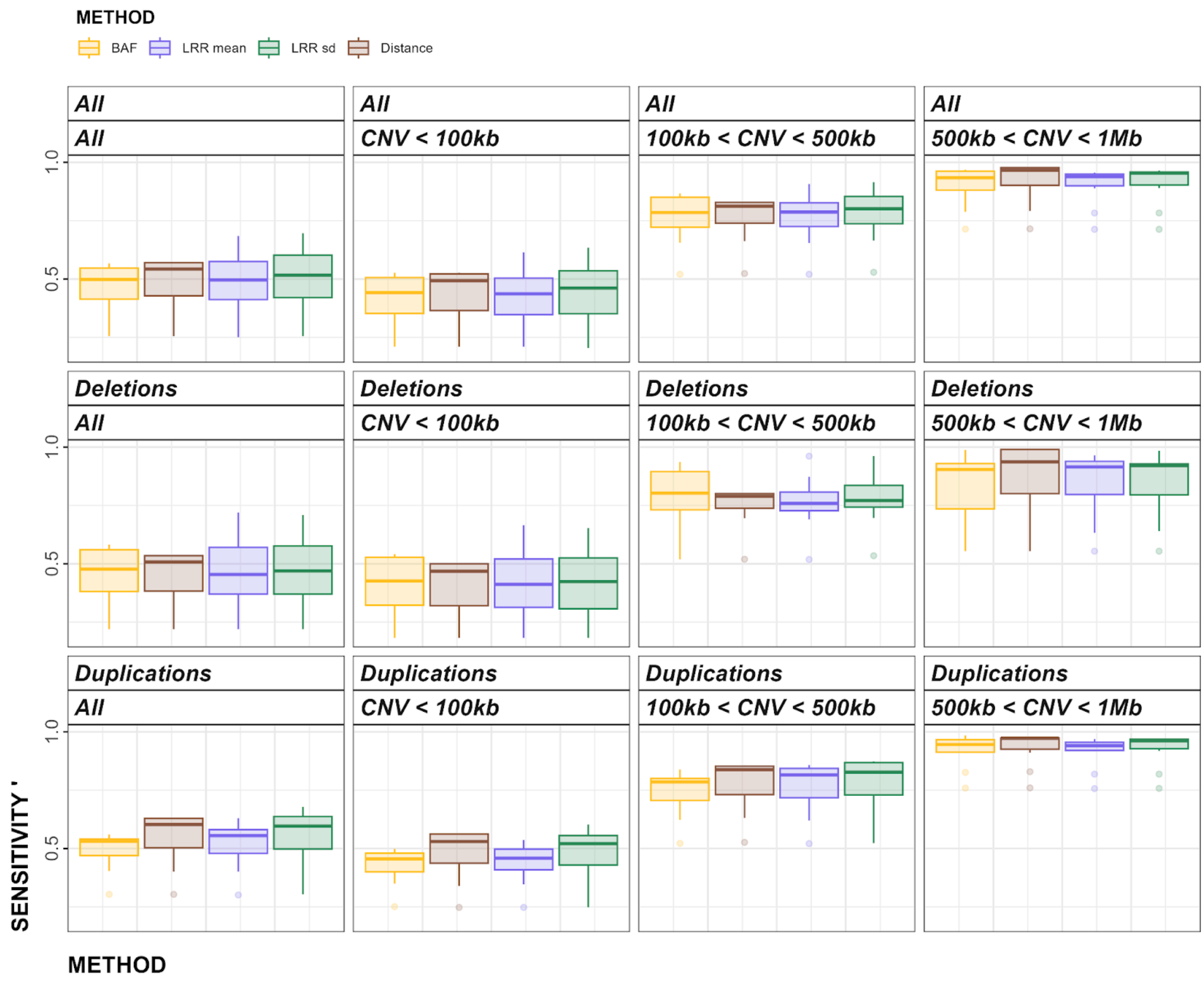


**Figure S38**. Within-Array Experiment plots of smoothed, scaled sensitivity (sensitivity’) metrics to determine optimal *Method* parameter.

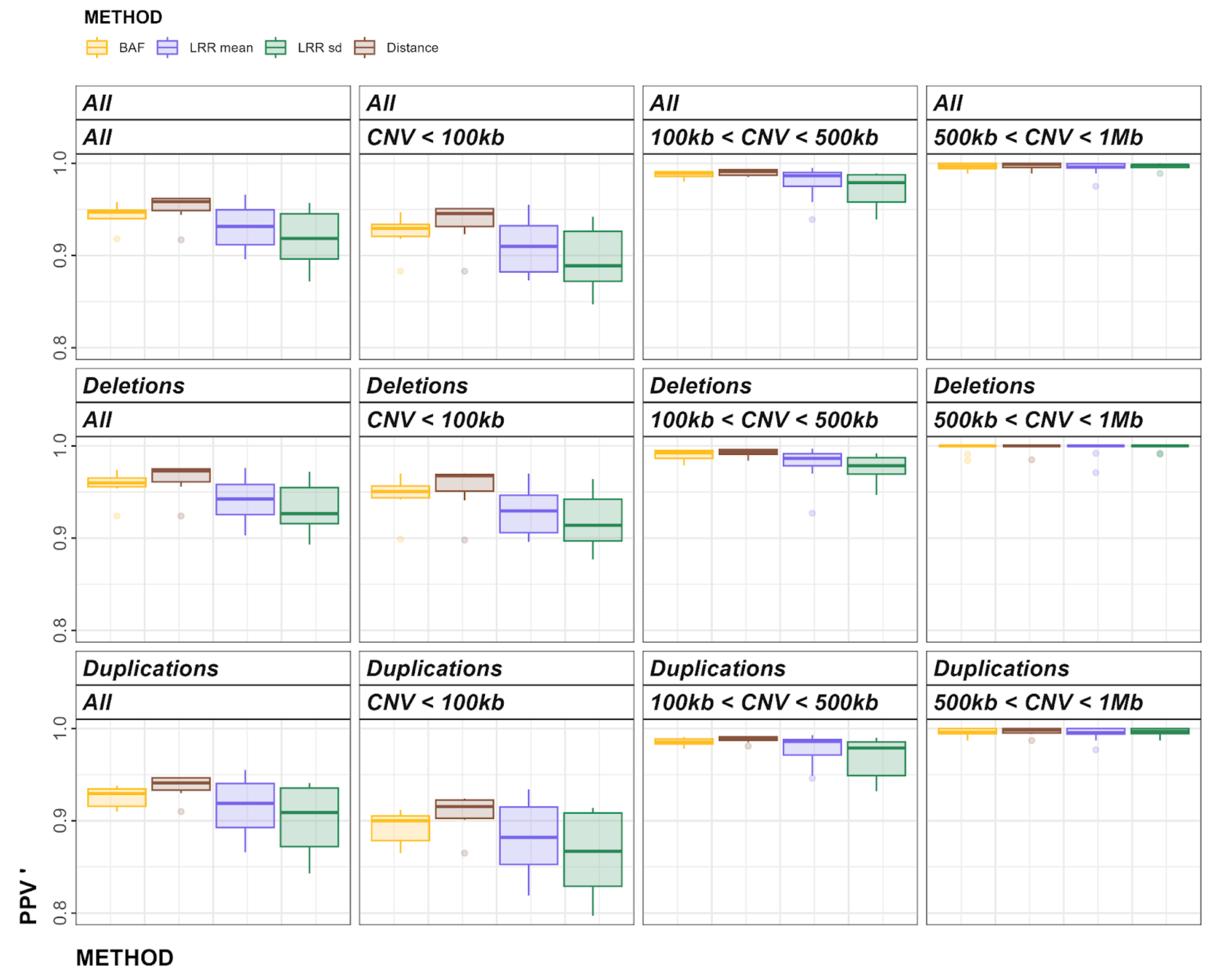


**Figure S39**. Within-Array Experiment plots of smoothed, scaled PPV (PPV’) metrics to determine optimal *Method* parameter.

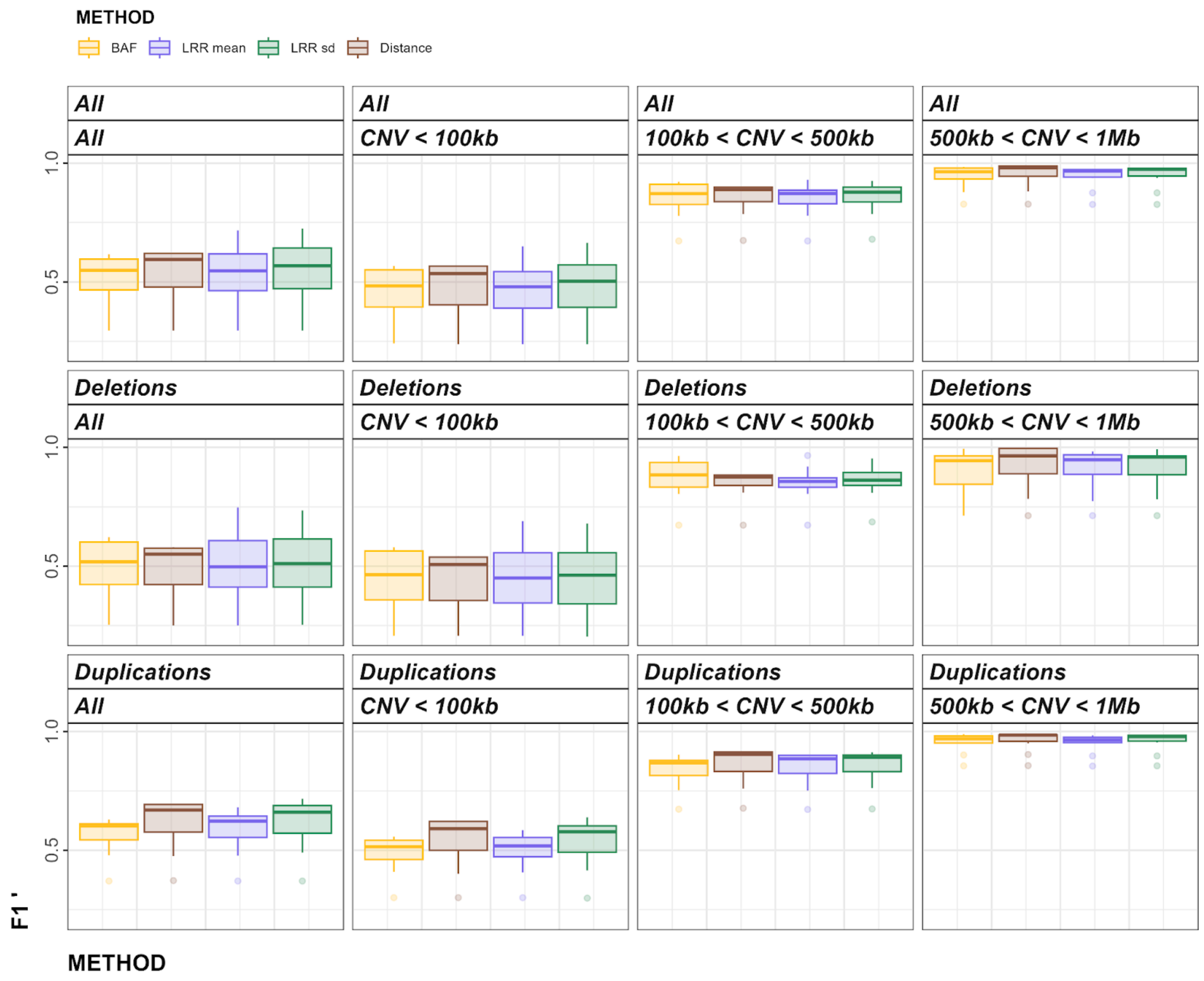


**Figure S40**. Within-Array Experiment plots of smoothed, scaled F1 (F1') metrics to determine optimal *Method* parameter.

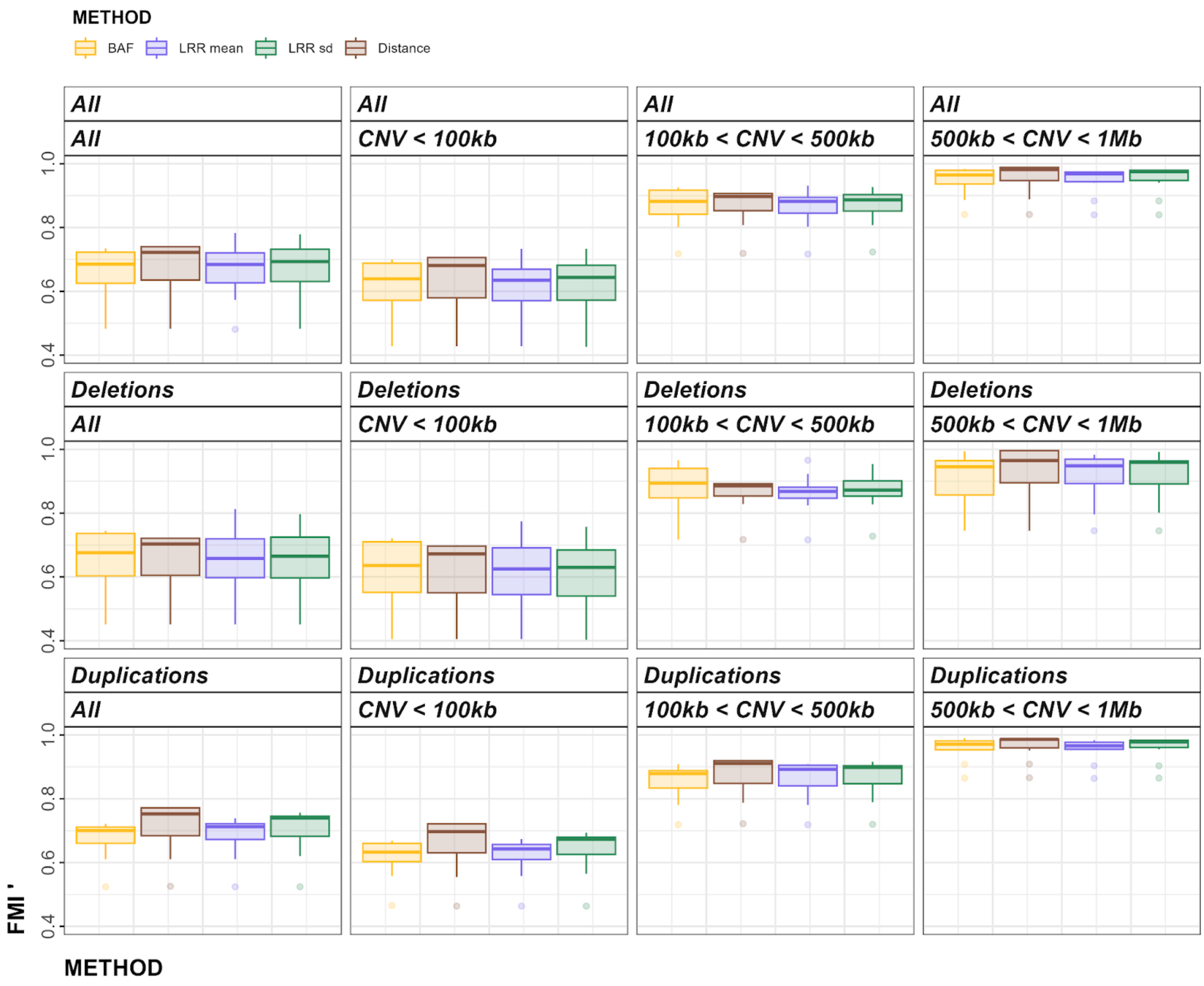


**Figure S41**. Within-Array Experiment plots of smoothed, scaled FMI (FMI’) metrics to determine optimal *Method* parameter.

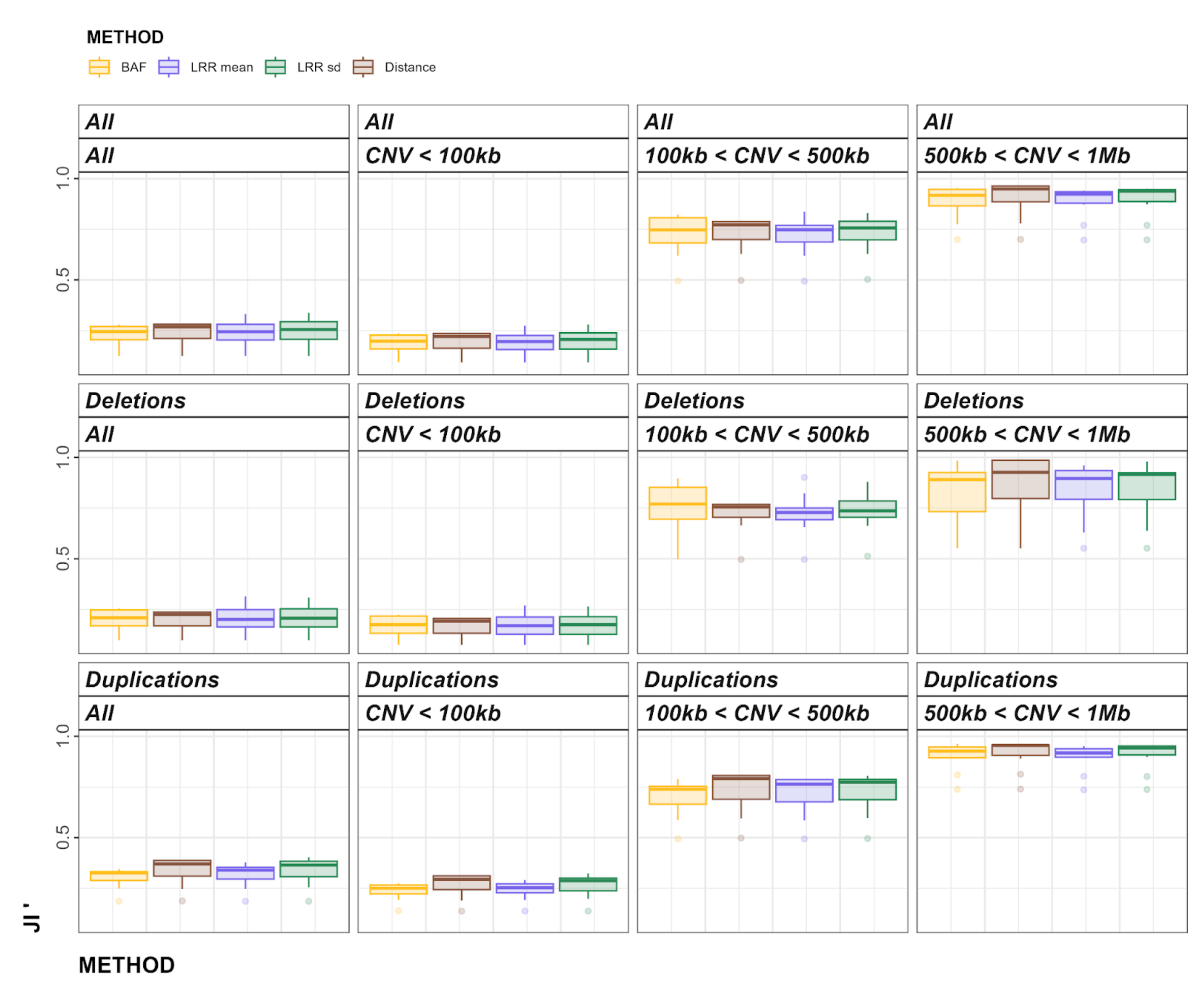


**Figure S42**. Within-Array Experiment plots of smoothed, scaled JI (JI') metrics to determine optimal *Method* parameter.

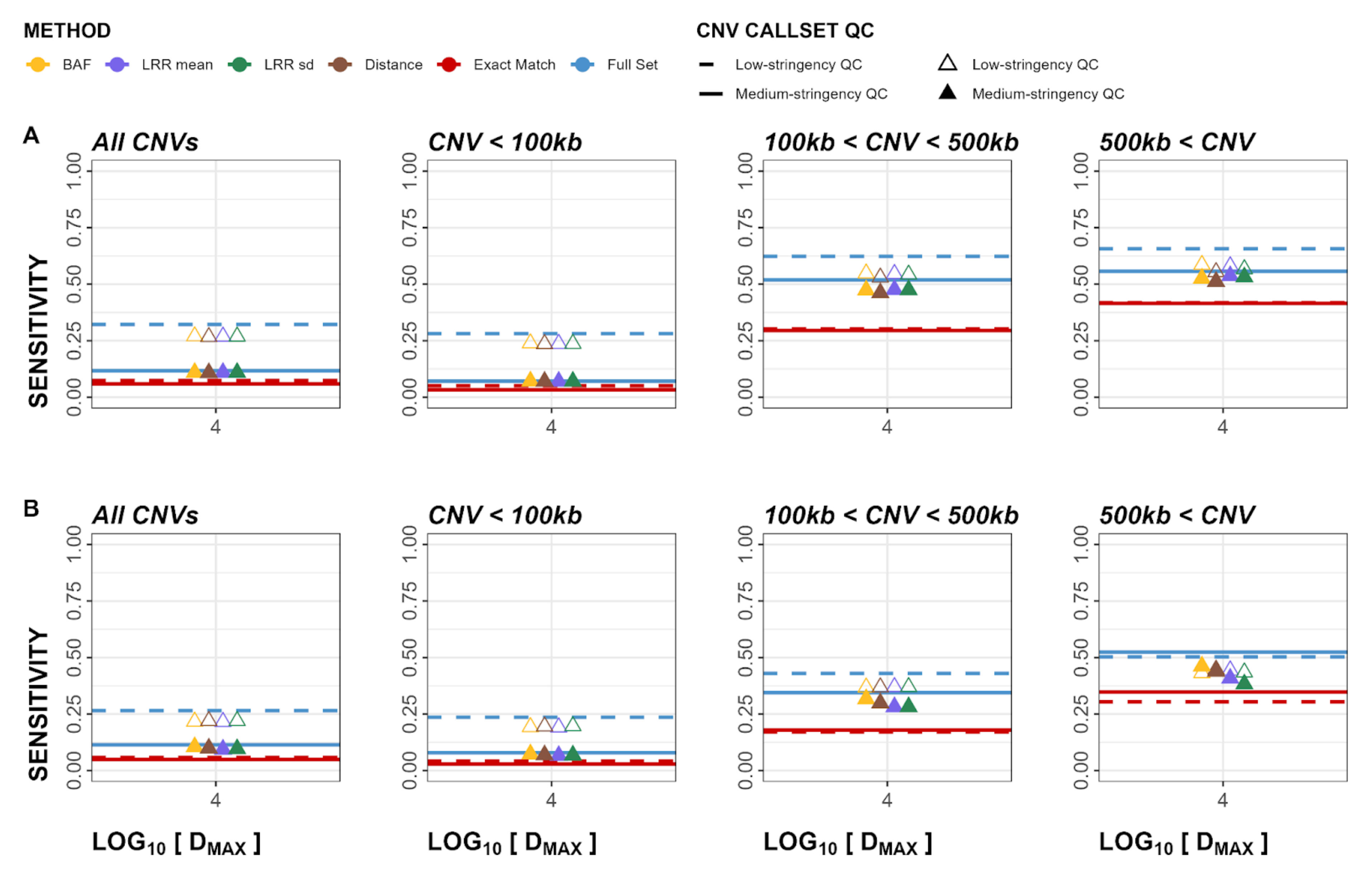


**Figure S43**. Cross-Array Experiment sensitivity plots for both deletions and duplications. Panel A represents calls in GSA validated in OEE. Panel B represents calls in OEE validated in GSA. Empty triangles and dashed lines represent low-stringency QC callsets, whereas solid lines and filled triangles represent medium-stringency QC callsets. GSA: Global Screening Array, OEE: Omni Express Exome array.

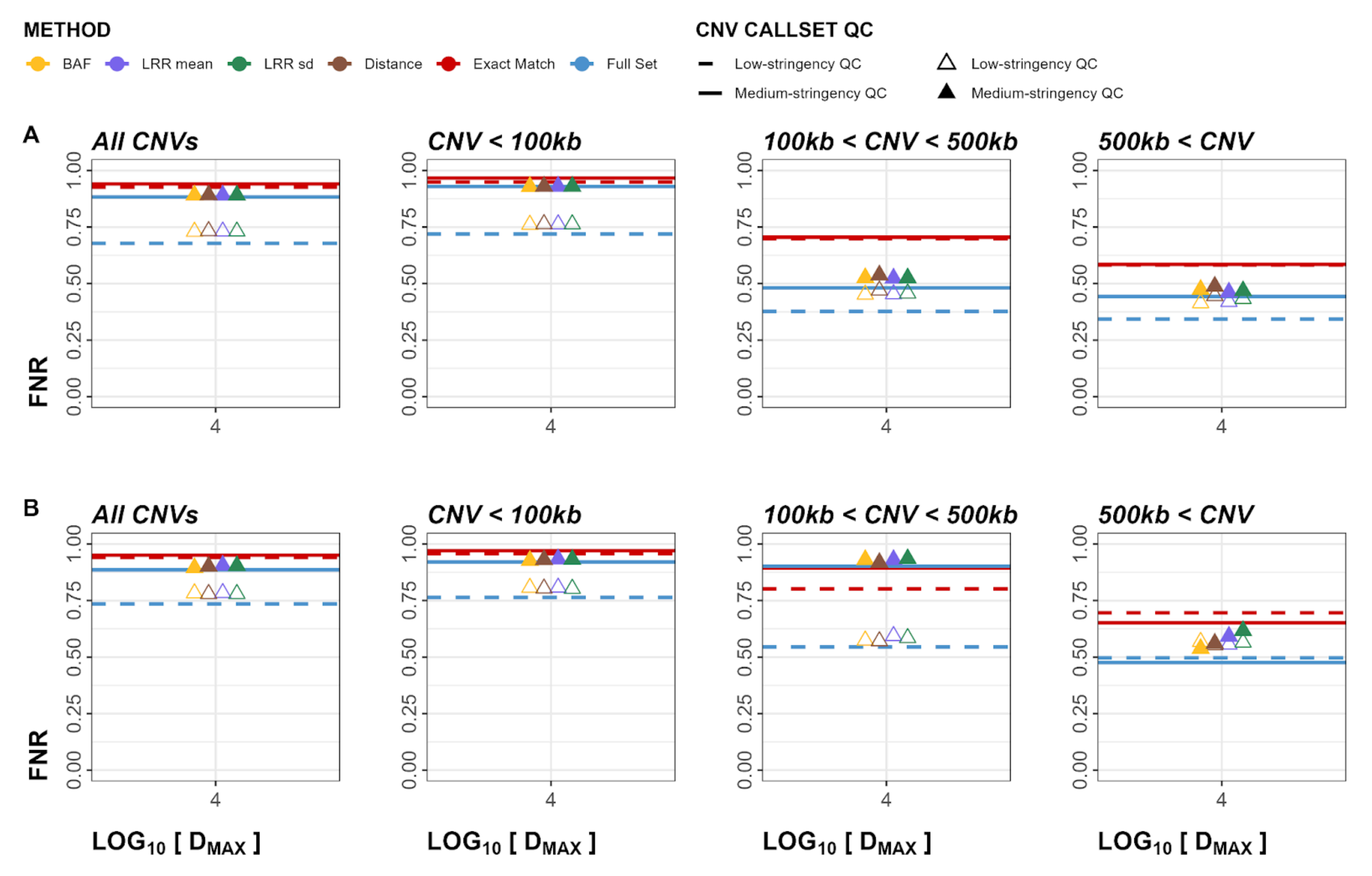


**Figure S44**. Cross-Array Experiment FNR plots for both deletions and duplications. Panel A represents calls in GSA validated in OEE. Panel B represents calls in OEE validated in GSA. Empty triangles and dashed lines represent low-stringency QC callsets, whereas solid lines and filled triangles represent medium-stringency QC callsets. GSA: Global Screening Array, OEE: Omni Express Exome array.

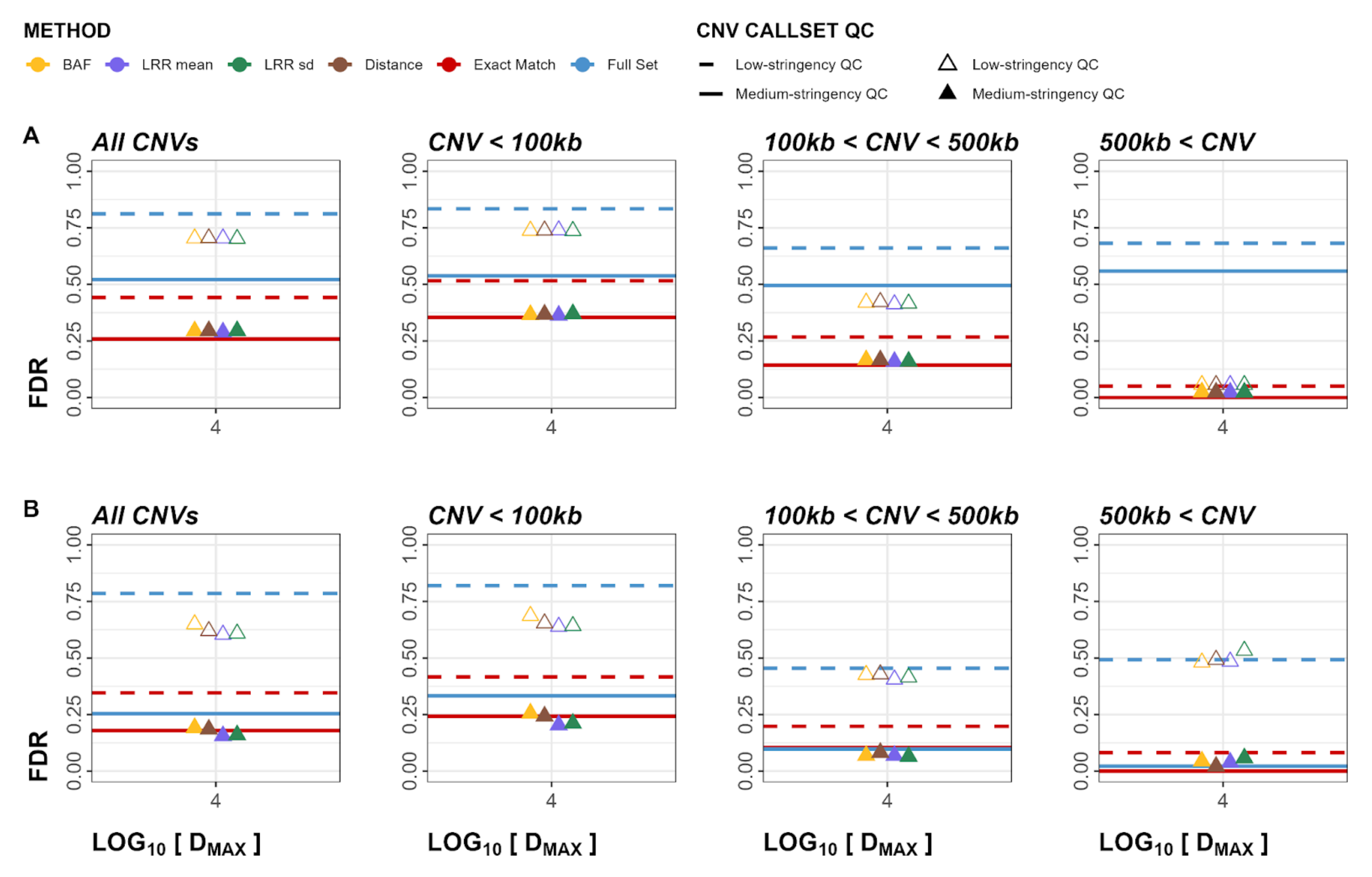


**Figure S45**. Cross-Array Experiment FDR plots for both deletions and duplications. Panel A represents calls in GSA validated in OEE. Panel B represents calls in OEE validated in GSA. Empty triangles and dashed lines represent low-stringency QC callsets, whereas solid lines and filled triangles represent medium-stringency QC callsets. GSA: Global Screening Array, OEE: Omni Express Exome array.

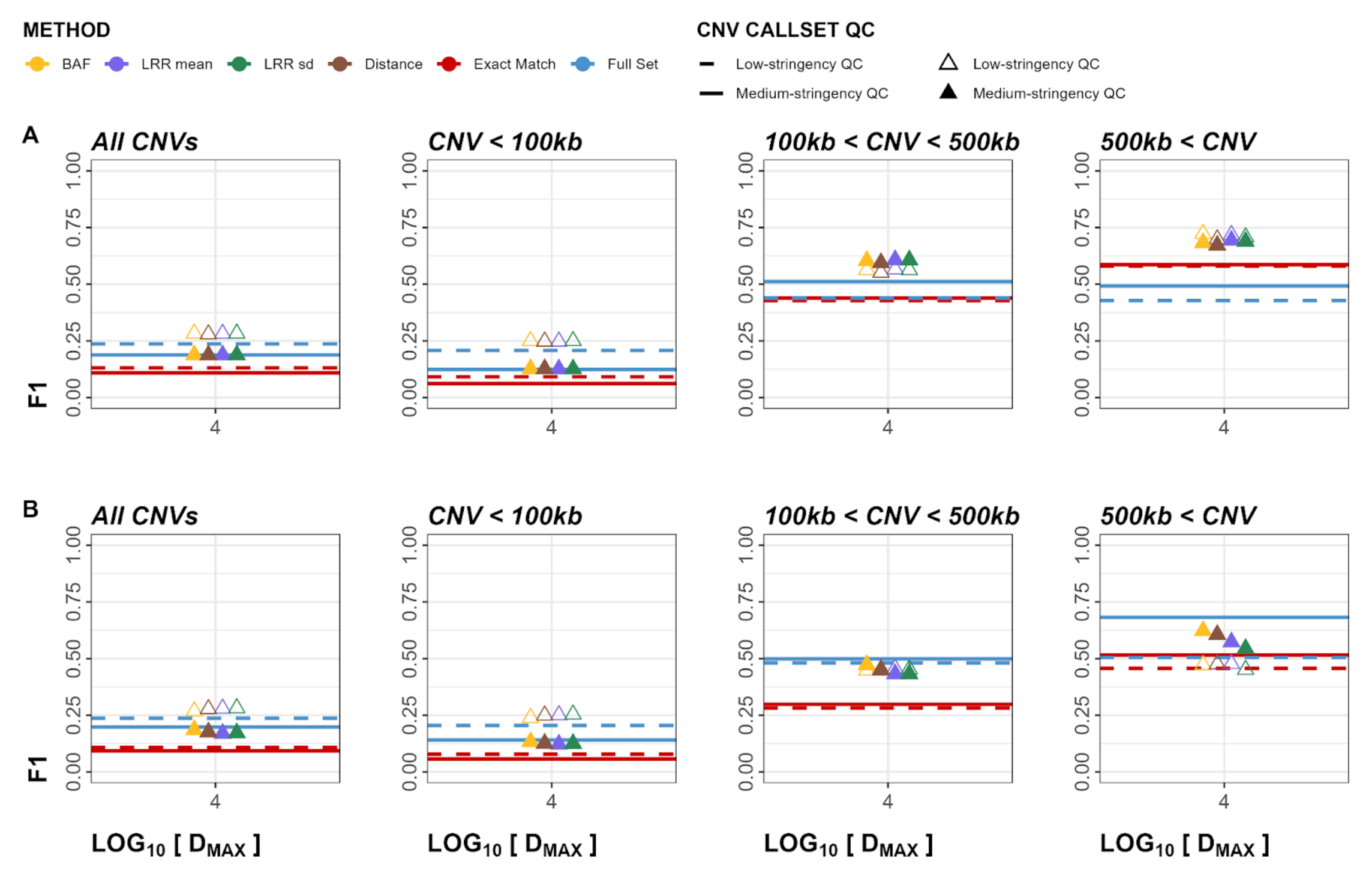


**Figure S46**. Cross-Array Experiment F1 plots for both deletions and duplications. Panel A represents calls in GSA validated in OEE. Panel B represents calls in OEE validated in GSA. Empty triangles and dashed lines represent low-stringency QC callsets, whereas solid lines and filled triangles represent medium-stringency QC callsets. GSA: Global Screening Array, OEE: Omni Express Exome array.

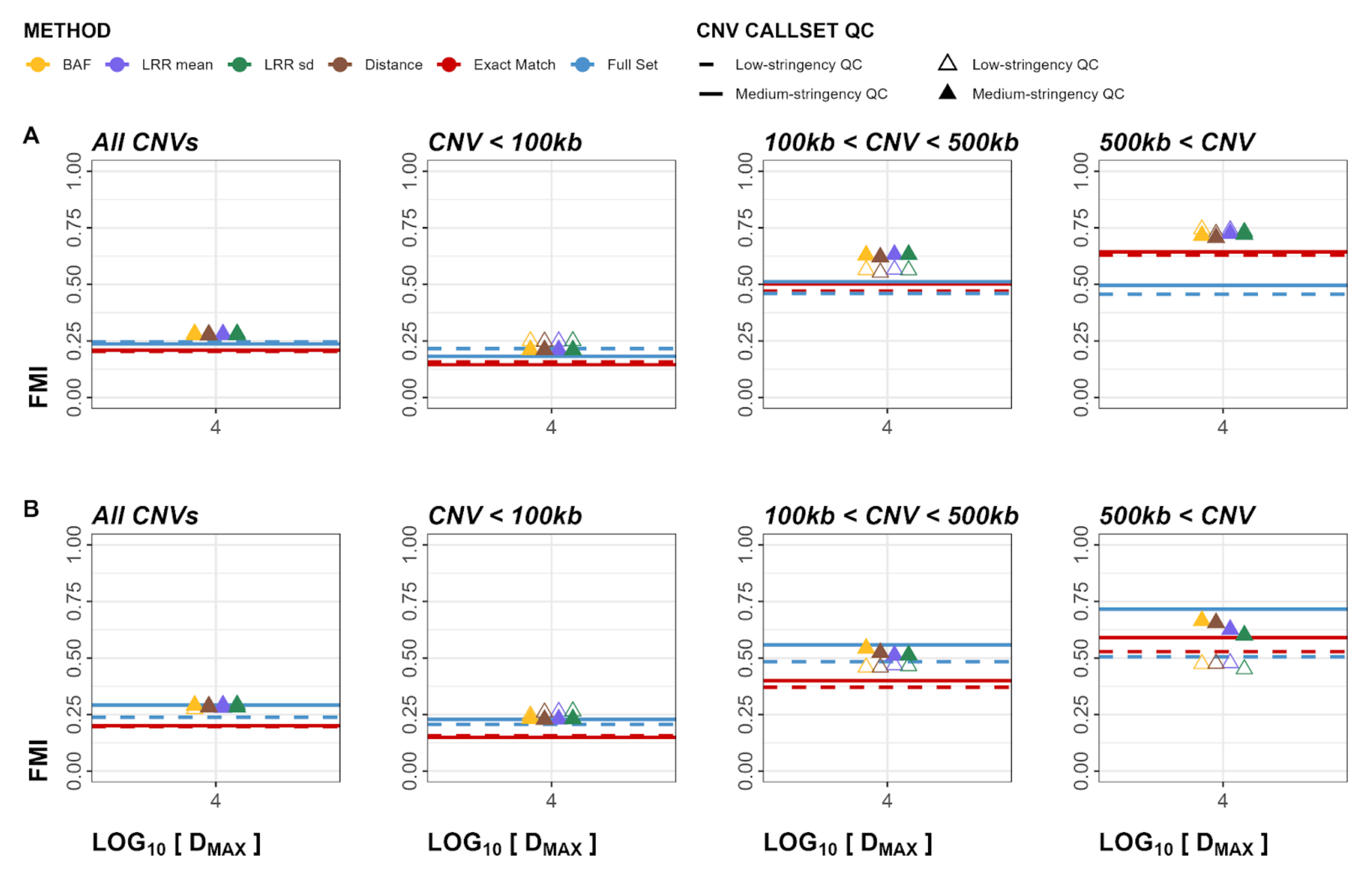


**Figure S47**. Cross-Array Experiment FMI plots for both deletions and duplications. Panel A represents calls in GSA validated in OEE. Panel B represents calls in OEE validated in GSA. Empty triangles and dashed lines represent low-stringency QC callsets, whereas solid lines and filled triangles represent medium-stringency QC callsets. GSA: Global Screening Array, OEE: Omni Express Exome array.

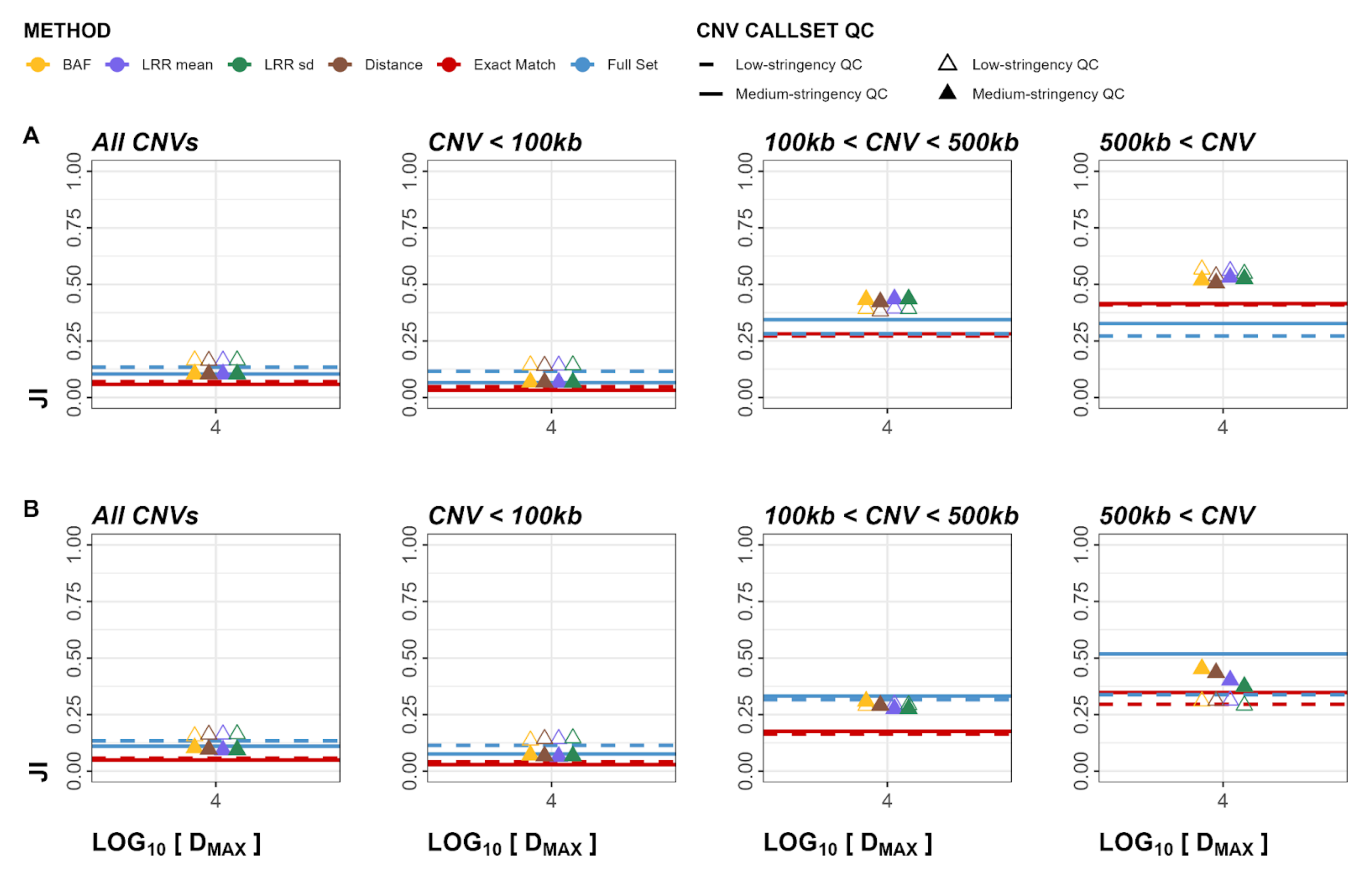


**Figure S48**. Cross-Array Experiment JI plots for both deletions and duplications. Panel A represents calls in GSA validated in OEE. Panel B represents calls in OEE validated in GSA. Empty triangles and dashed lines represent low-stringency QC callsets, whereas solid lines and filled triangles represent medium-stringency QC callsets. GSA: Global Screening Array, OEE: Omni Express Exome array.

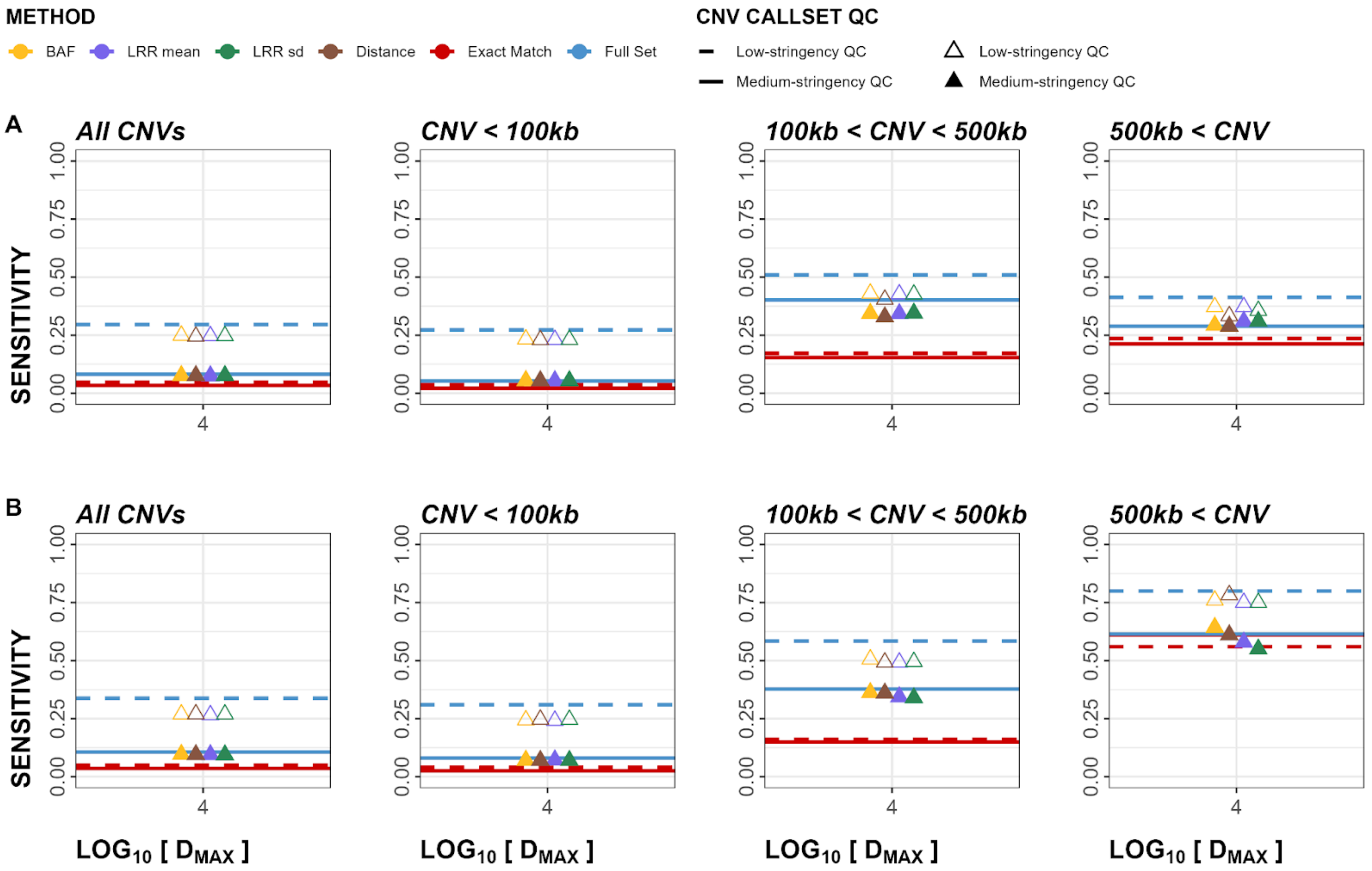


**Figure S49**. Cross-Array Experiment sensitivity plots for deletions. Panel A represents calls in GSA validated in OEE. Panel B represents calls in OEE validated in GSA. Empty triangles and dashed lines represent low-stringency QC callsets, whereas solid lines and filled triangles represent medium-stringency QC callsets. GSA: Global Screening Array, OEE: Omni Express Exome array.

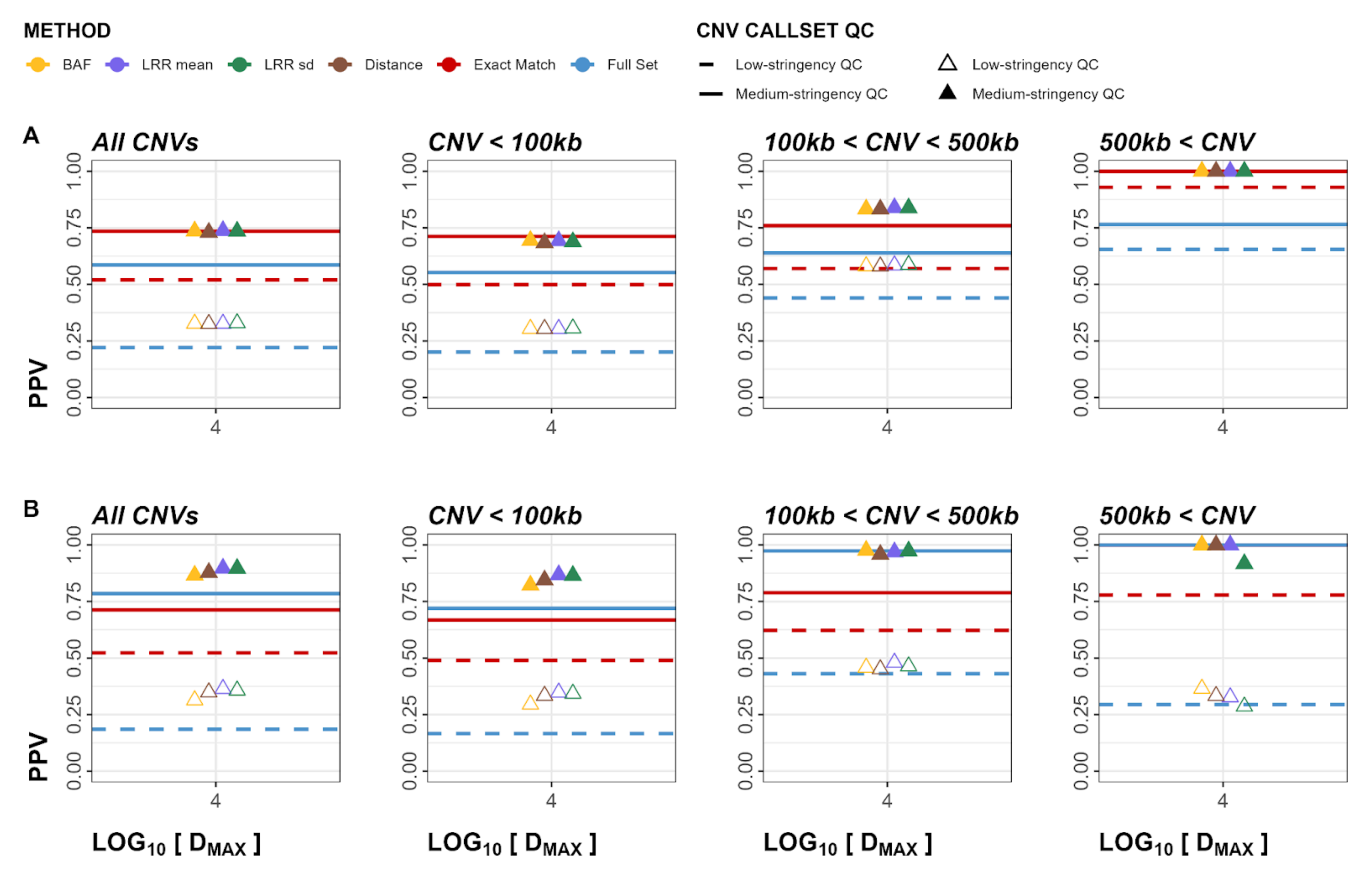


**Figure S50**. Cross-Array Experiment PPV plots for deletions. Panel A represents calls in GSA validated in OEE. Panel B represents calls in OEE validated in GSA. Empty triangles and dashed lines represent low-stringency QC callsets, whereas solid lines and filled triangles represent medium-stringency QC callsets. GSA: Global Screening Array, OEE: Omni Express Exome array.

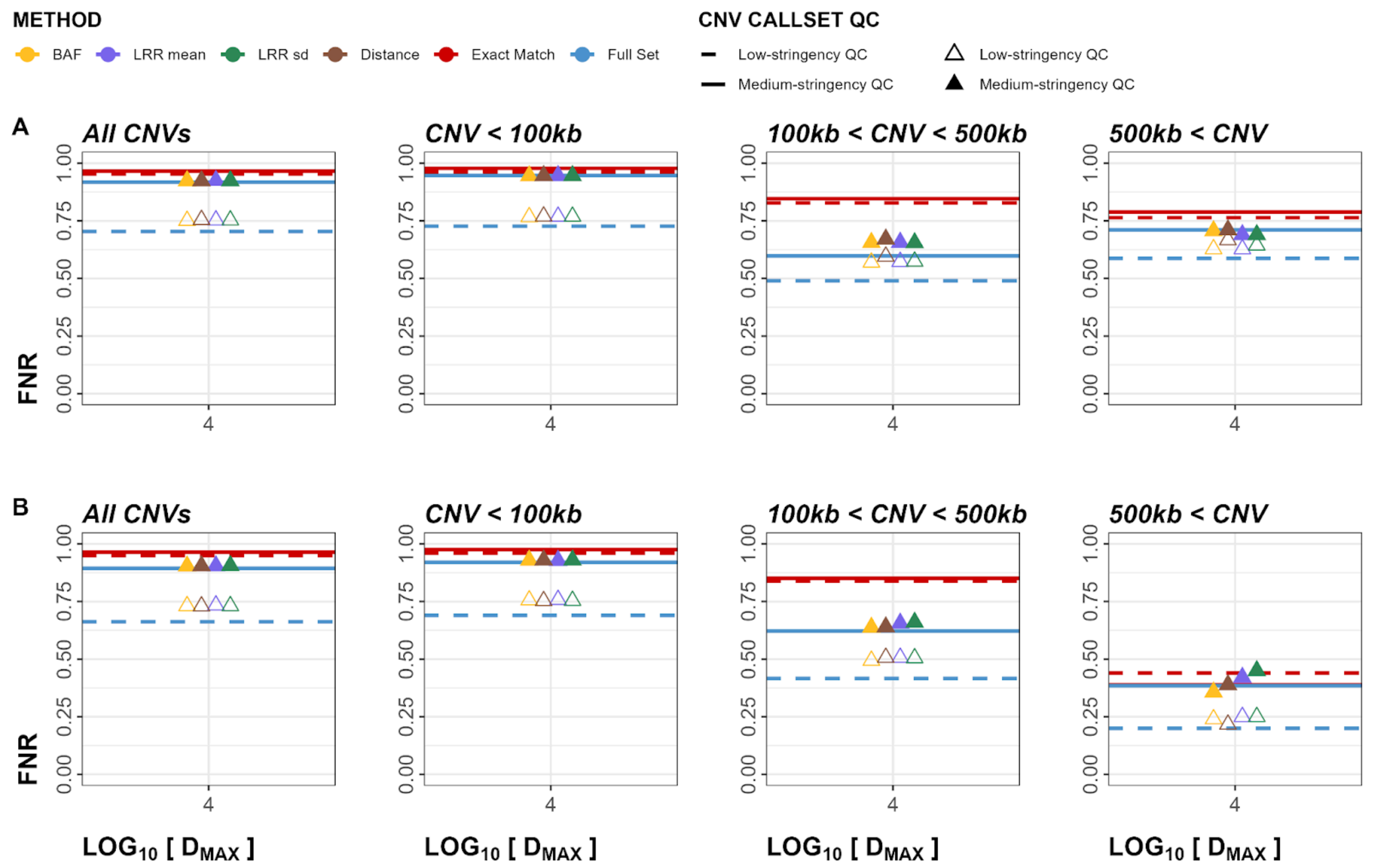


**Figure S51**. Cross-Array Experiment FNR plots for deletions. Panel A represents calls in GSA validated in OEE. Panel B represents calls in OEE validated in GSA. Empty triangles and dashed lines represent low-stringency QC callsets, whereas solid lines and filled triangles represent medium-stringency QC callsets. GSA: Global Screening Array, OEE: Omni Express Exome array.

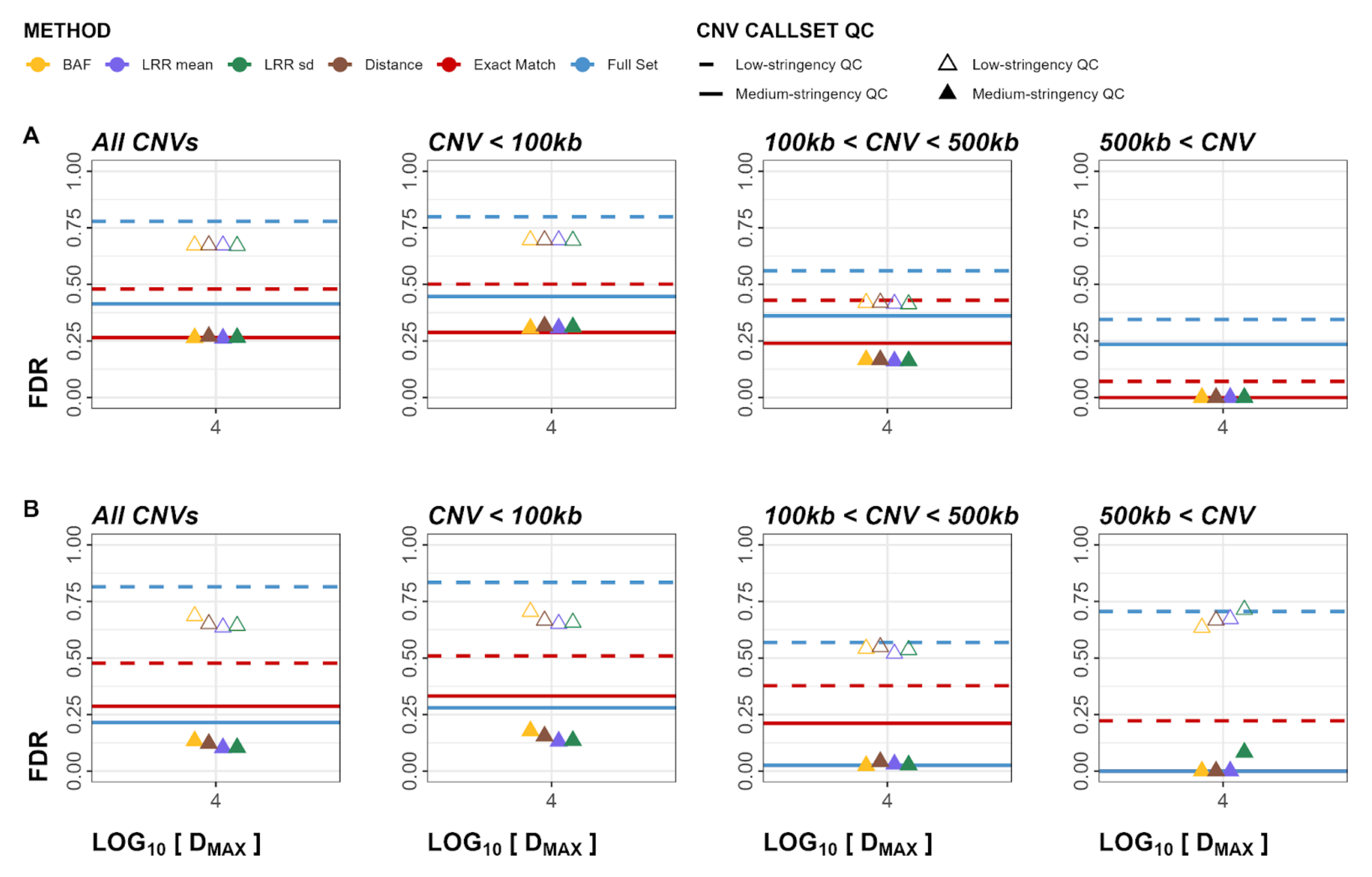


**Figure S52**. Cross-Array Experiment FDR plots for deletions. Panel A represents calls in GSA validated in OEE. Panel B represents calls in OEE validated in GSA. Empty triangles and dashed lines represent low-stringency QC callsets, whereas solid lines and filled triangles represent medium-stringency QC callsets. GSA: Global Screening Array, OEE: Omni Express Exome array.

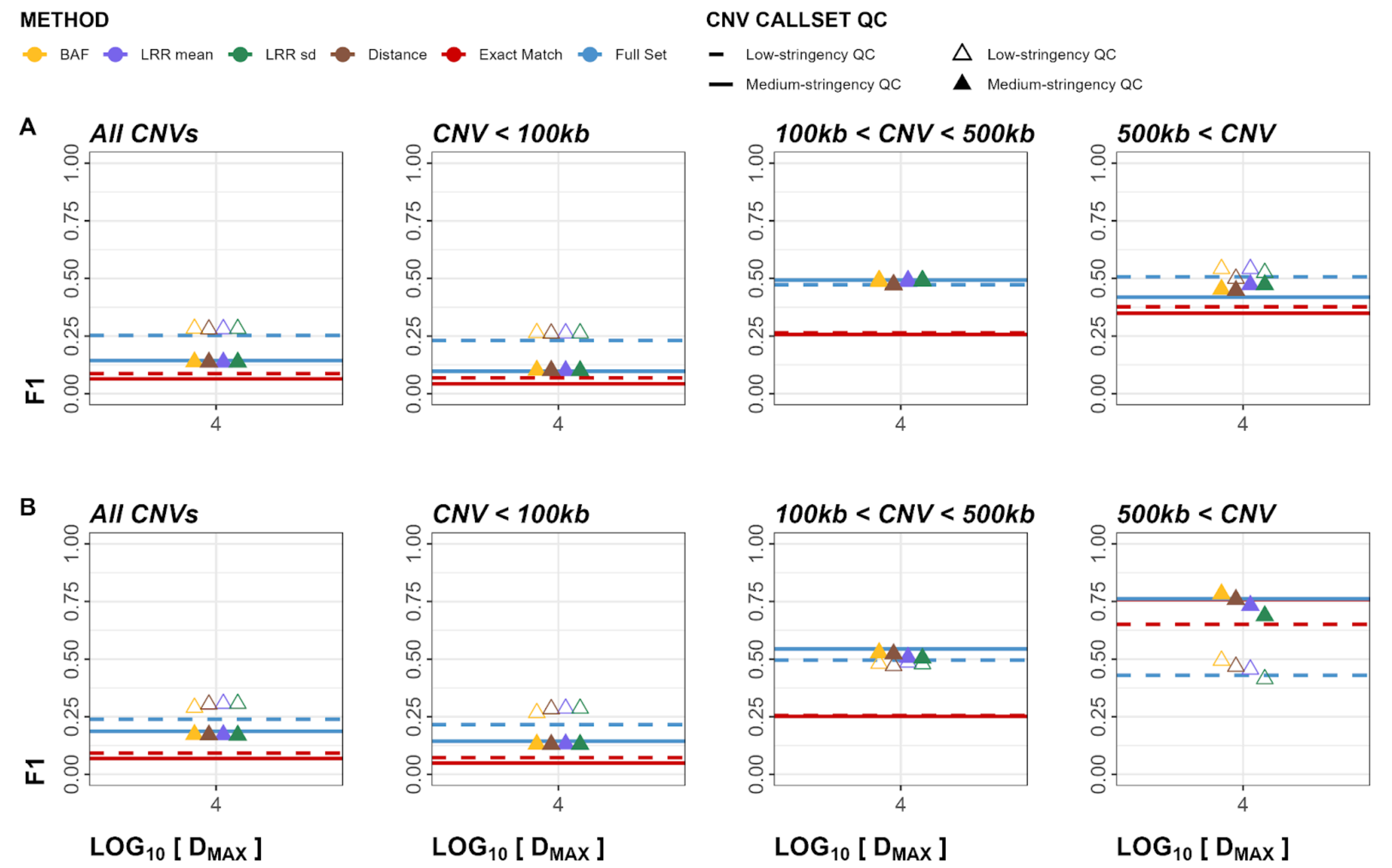


**Figure S53**. Cross-Array Experiment F1 plots for deletions. Panel A represents calls in GSA validated in OEE. Panel B represents calls in OEE validated in GSA. Empty triangles and dashed lines represent low-stringency QC callsets, whereas solid lines and filled triangles represent medium-stringency QC callsets. GSA: Global Screening Array, OEE: Omni Express Exome array.

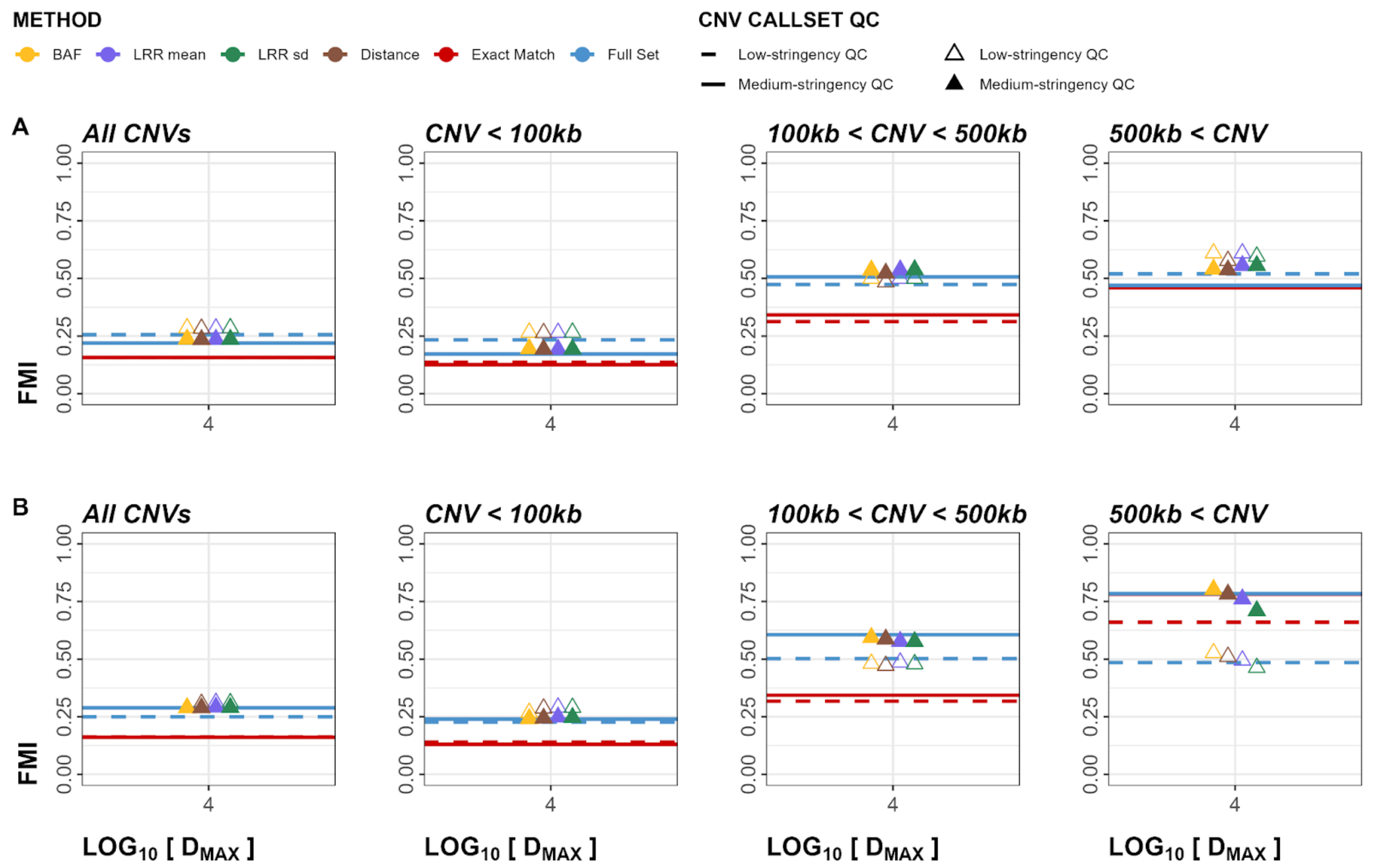


**Figure S54**. Cross-Array Experiment FMI plots for deletions. Panel A represents calls in GSA validated in OEE. Panel B represents calls in OEE validated in GSA. Empty triangles and dashed lines represent low-stringency QC callsets, whereas solid lines and filled triangles represent medium-stringency QC callsets. GSA: Global Screening Array, OEE: Omni Express Exome array.

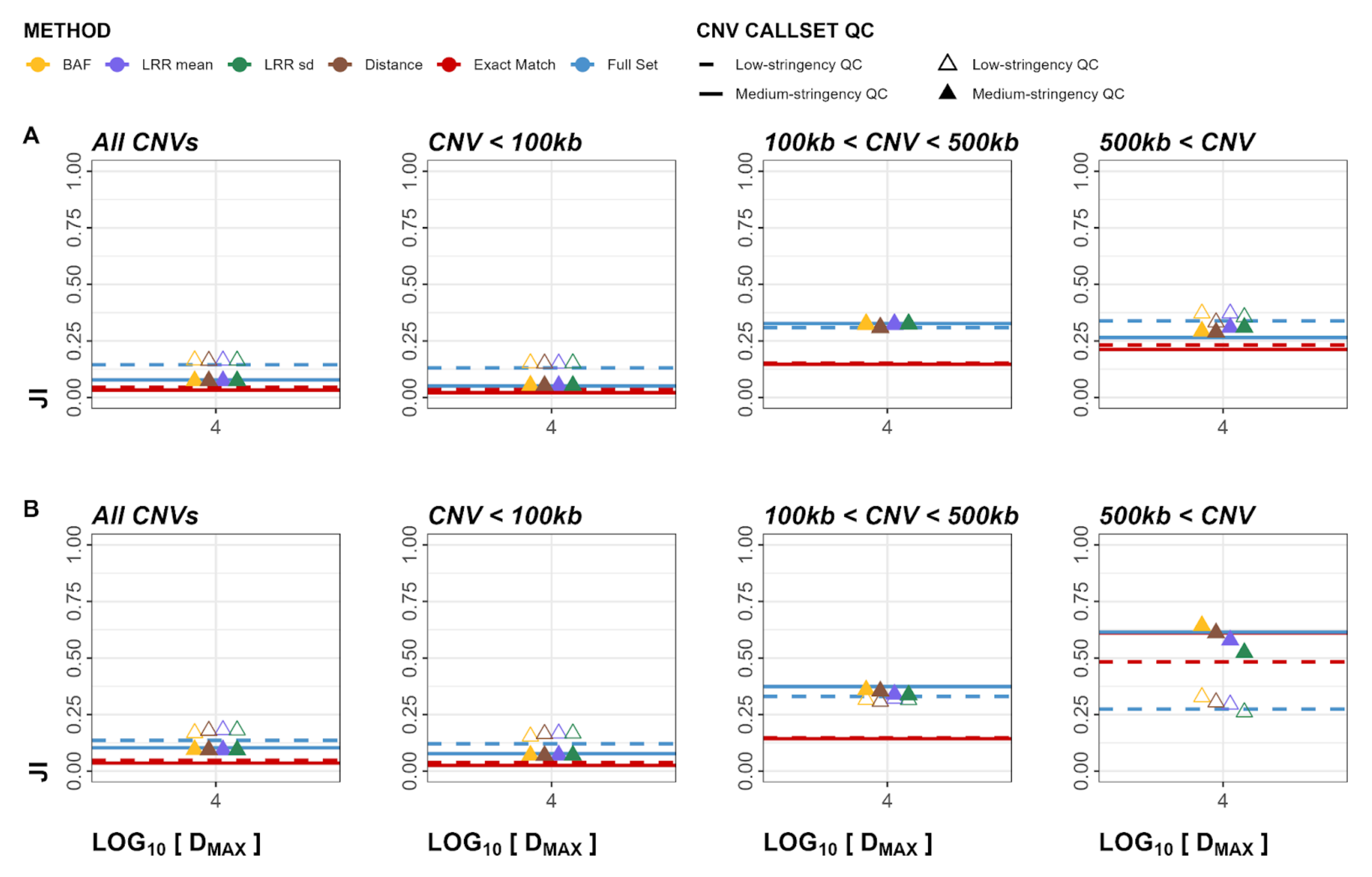


**Figure S55**. Cross-Array Experiment JI plots for deletions. Panel A represents calls in GSA validated in OEE. Panel B represents calls in OEE validated in GSA. Empty triangles and dashed lines represent low-stringency QC callsets, whereas solid lines and filled triangles represent medium-stringency QC callsets. GSA: Global Screening Array, OEE: Omni Express Exome array.

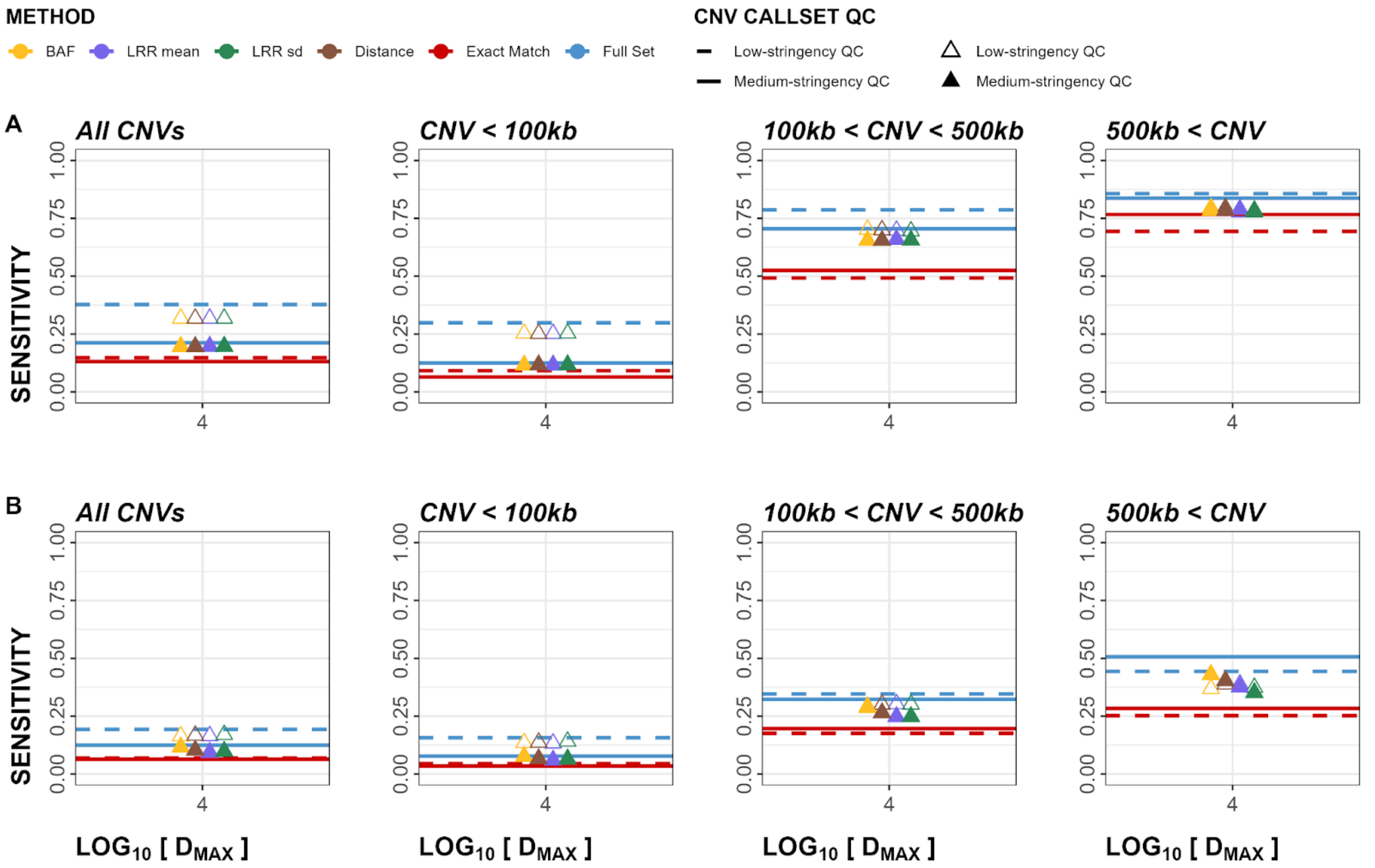


**Figure S56**. Cross-Array Experiment sensitivity plots for duplications. Panel A represents calls in GSA validated in OEE. Panel B represents calls in OEE validated in GSA. Empty triangles and dashed lines represent low-stringency QC callsets, whereas solid lines and filled triangles represent medium-stringency QC callsets. GSA: Global Screening Array, OEE: Omni Express Exome array.

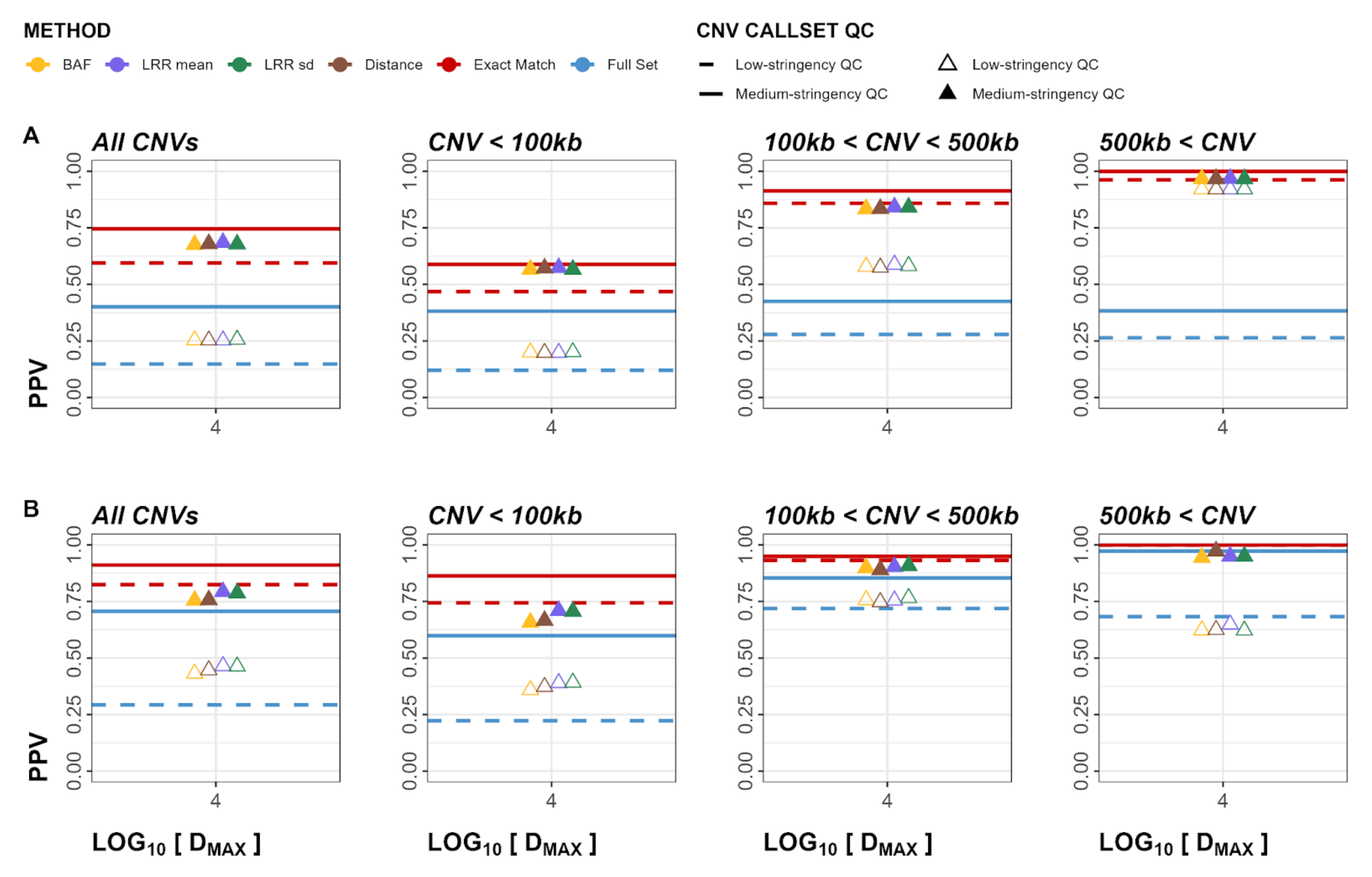


**Figure S57**. Cross-Array Experiment PPV plots for duplications. Panel A represents calls in GSA validated in OEE. Panel B represents calls in OEE validated in GSA. Empty triangles and dashed lines represent low-stringency QC callsets, whereas solid lines and filled triangles represent medium-stringency QC callsets. GSA: Global Screening Array, OEE: Omni Express Exome array.

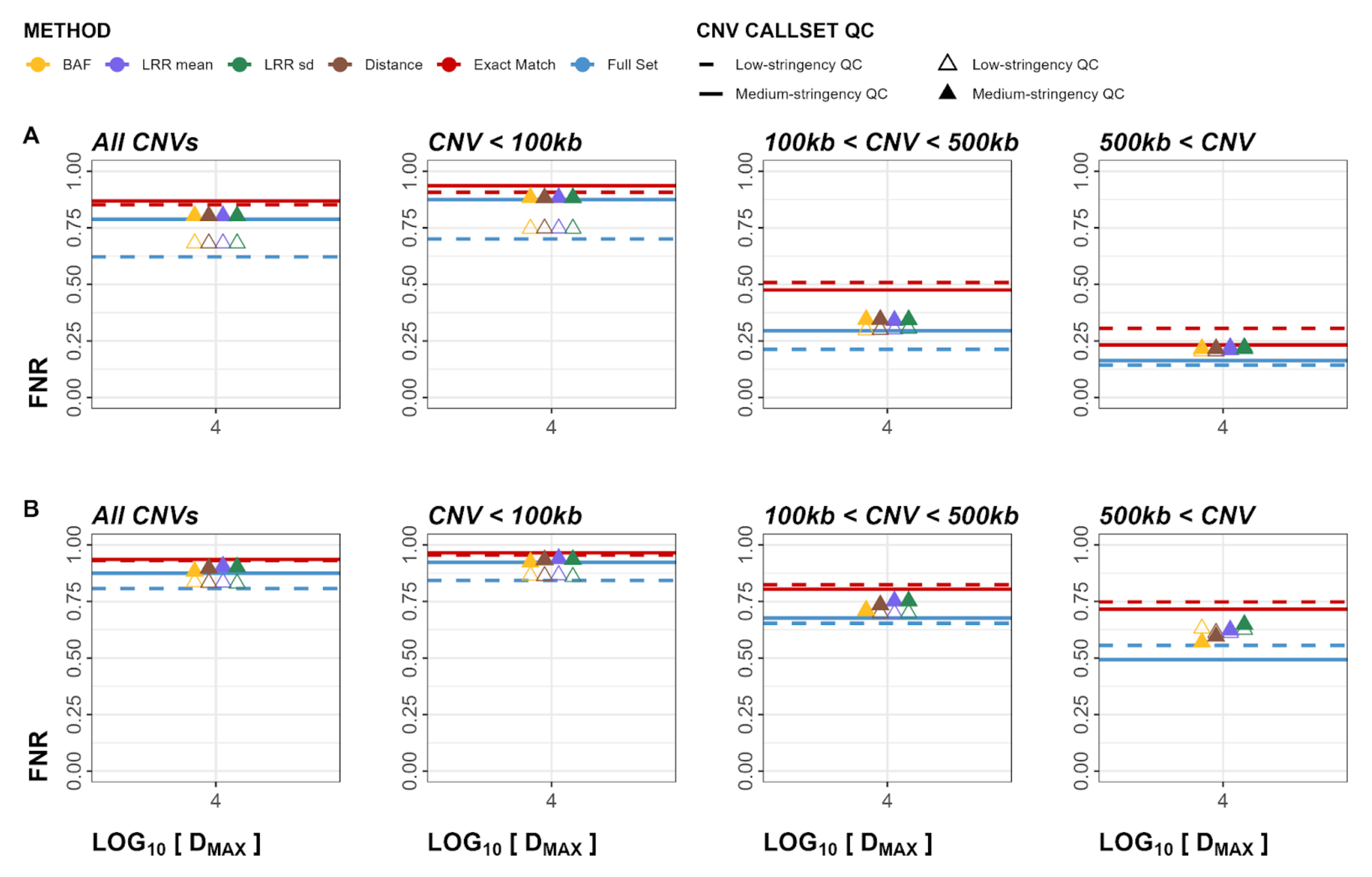


**Figure S58**. Cross-Array Experiment FNR plots for duplications. Panel A represents calls in GSA validated in OEE. Panel B represents calls in OEE validated in GSA. Empty triangles and dashed lines represent low-stringency QC callsets, whereas solid lines and filled triangles represent medium-stringency QC callsets. GSA: Global Screening Array, OEE: Omni Express Exome array.

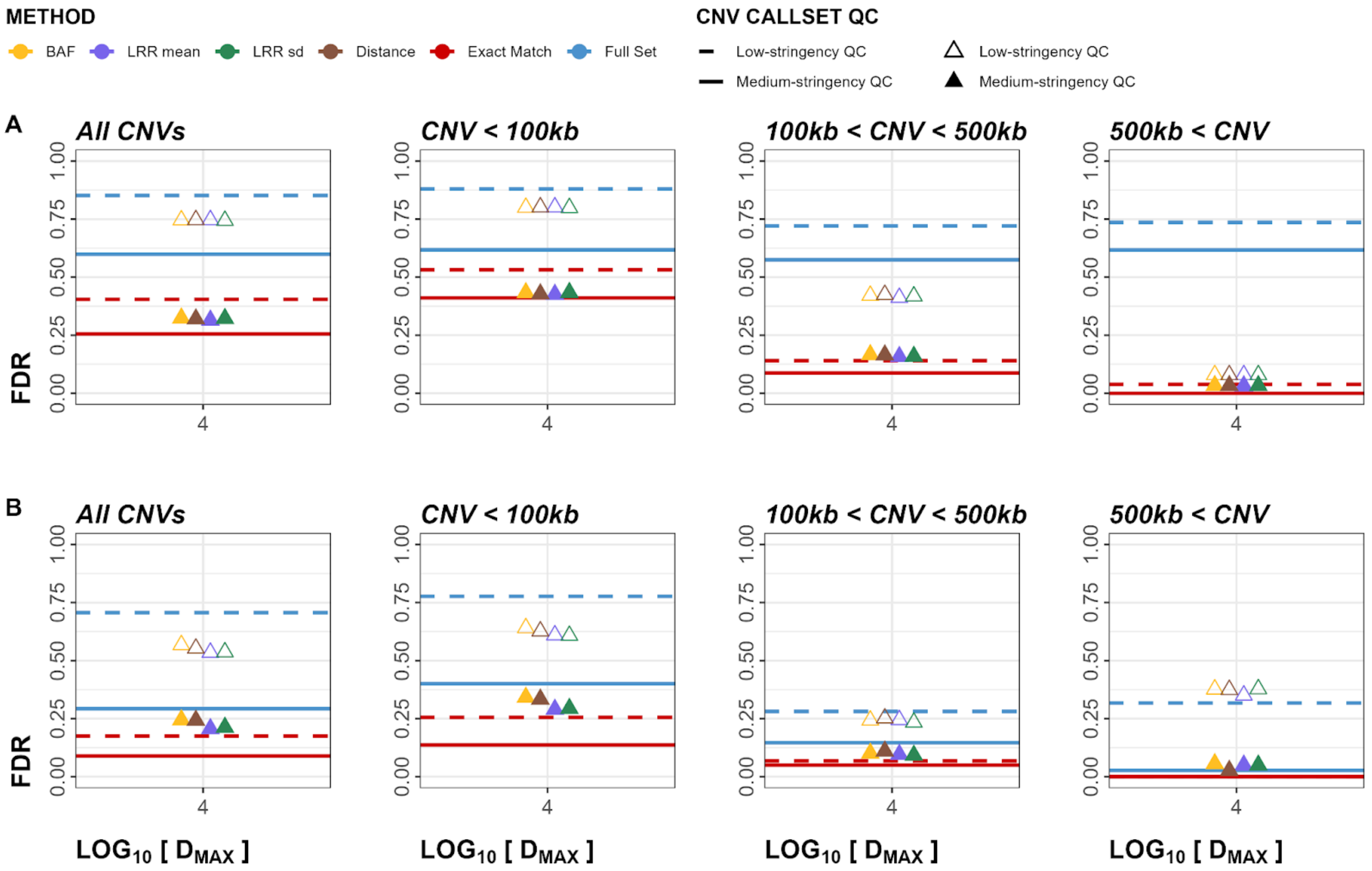


**Figure S59**. Cross-Array Experiment FDR plots for duplications. Panel A represents calls in GSA validated in OEE. Panel B represents calls in OEE validated in GSA. Empty triangles and dashed lines represent low-stringency QC callsets, whereas solid lines and filled triangles represent medium-stringency QC callsets. GSA: Global Screening Array, OEE: Omni Express Exome array.

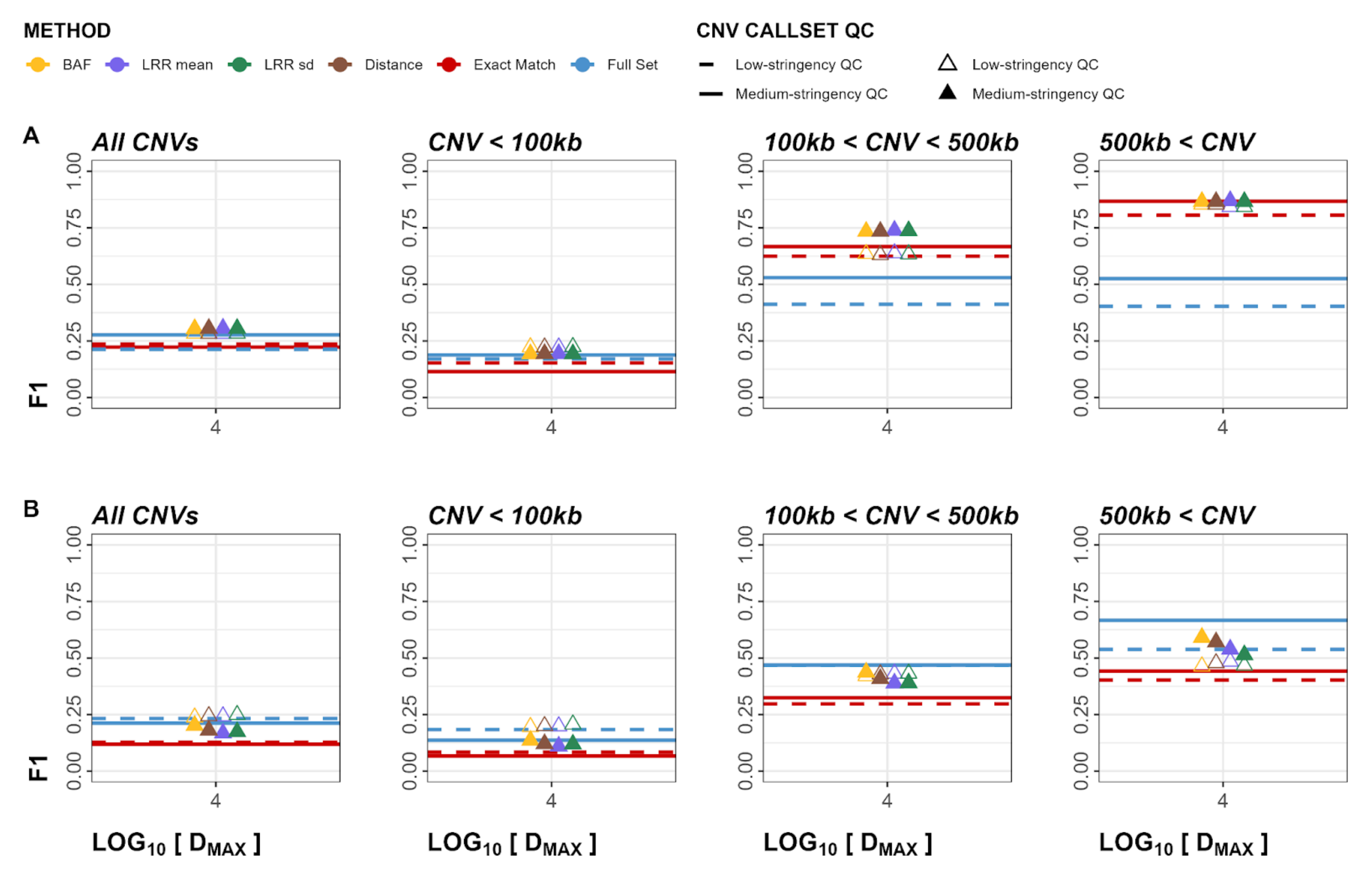


**Figure S60**. Cross-Array Experiment F1 plots for duplications. Panel A represents calls in GSA validated in OEE. Panel B represents calls in OEE validated in GSA. Empty triangles and dashed lines represent low-stringency QC callsets, whereas solid lines and filled triangles represent medium-stringency QC callsets. GSA: Global Screening Array, OEE: Omni Express Exome array.

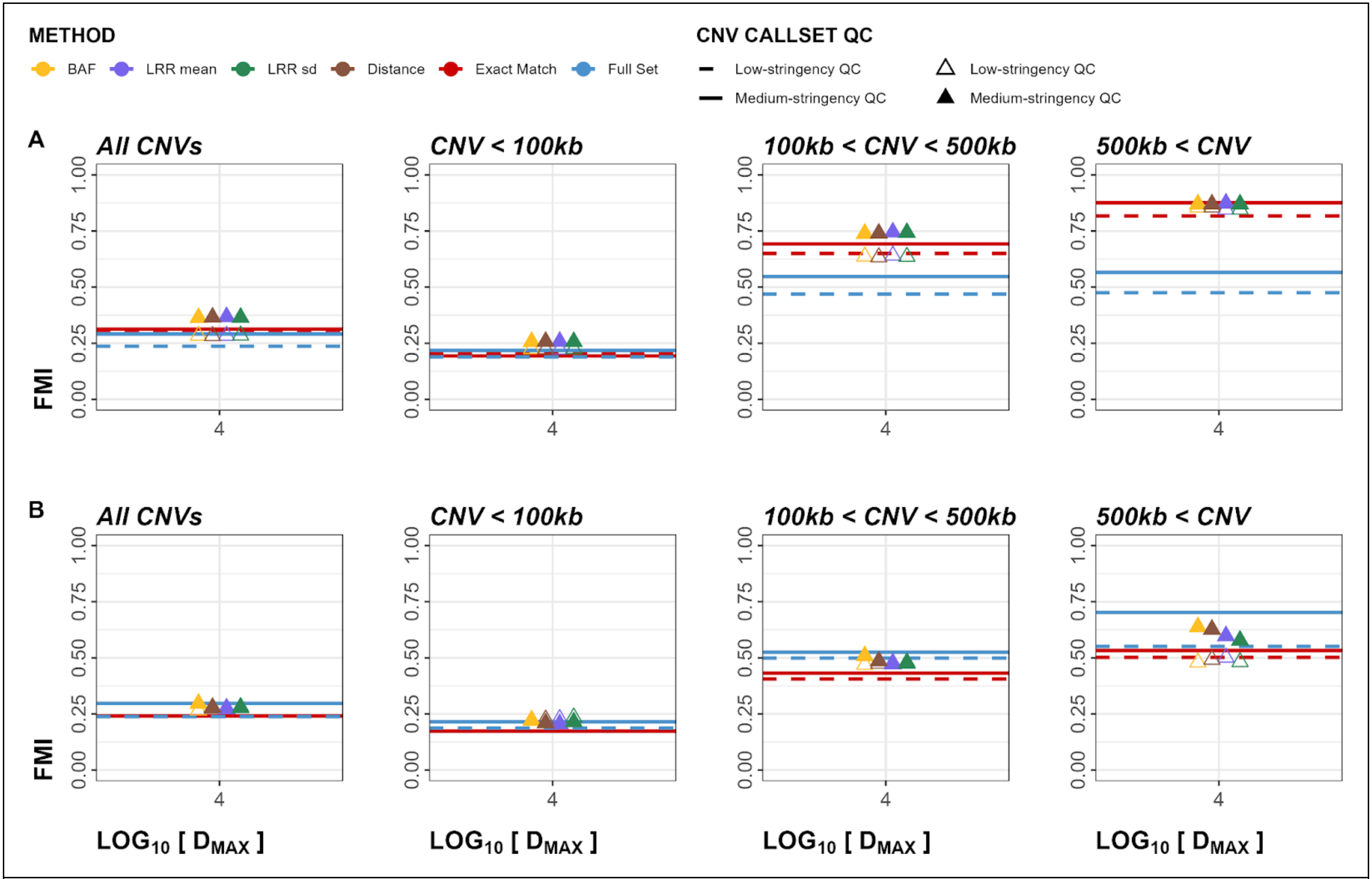


**Figure S61**. Cross-Array Experiment FMI plots for duplications. Panel A represents calls in GSA validated in OEE. Panel B represents calls in OEE validated in GSA. Empty triangles and dashed lines represent low-stringency QC callsets, whereas solid lines and filled triangles represent medium-stringency QC callsets. GSA: Global Screening Array, OEE: Omni Express Exome array.

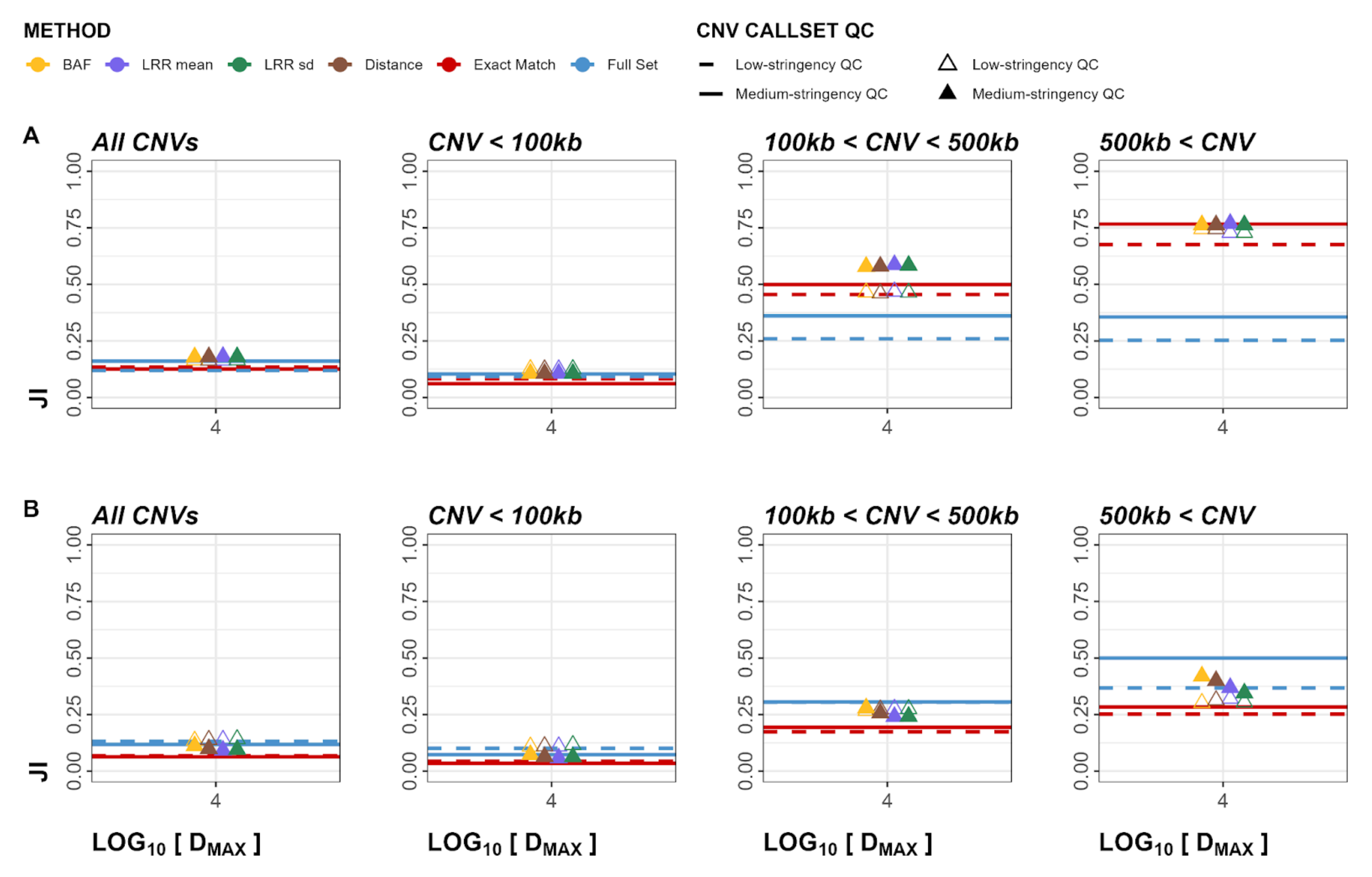


**Figure S62**. Cross-Array Experiment JI plots for duplications. Panel A represents calls in GSA validated in OEE. Panel B represents calls in OEE validated in GSA. Empty triangles and dashed lines represent low-stringency QC callsets, whereas solid lines and filled triangles represent medium-stringency QC callsets. GSA: Global Screening Array, OEE: Omni Express Exome array.

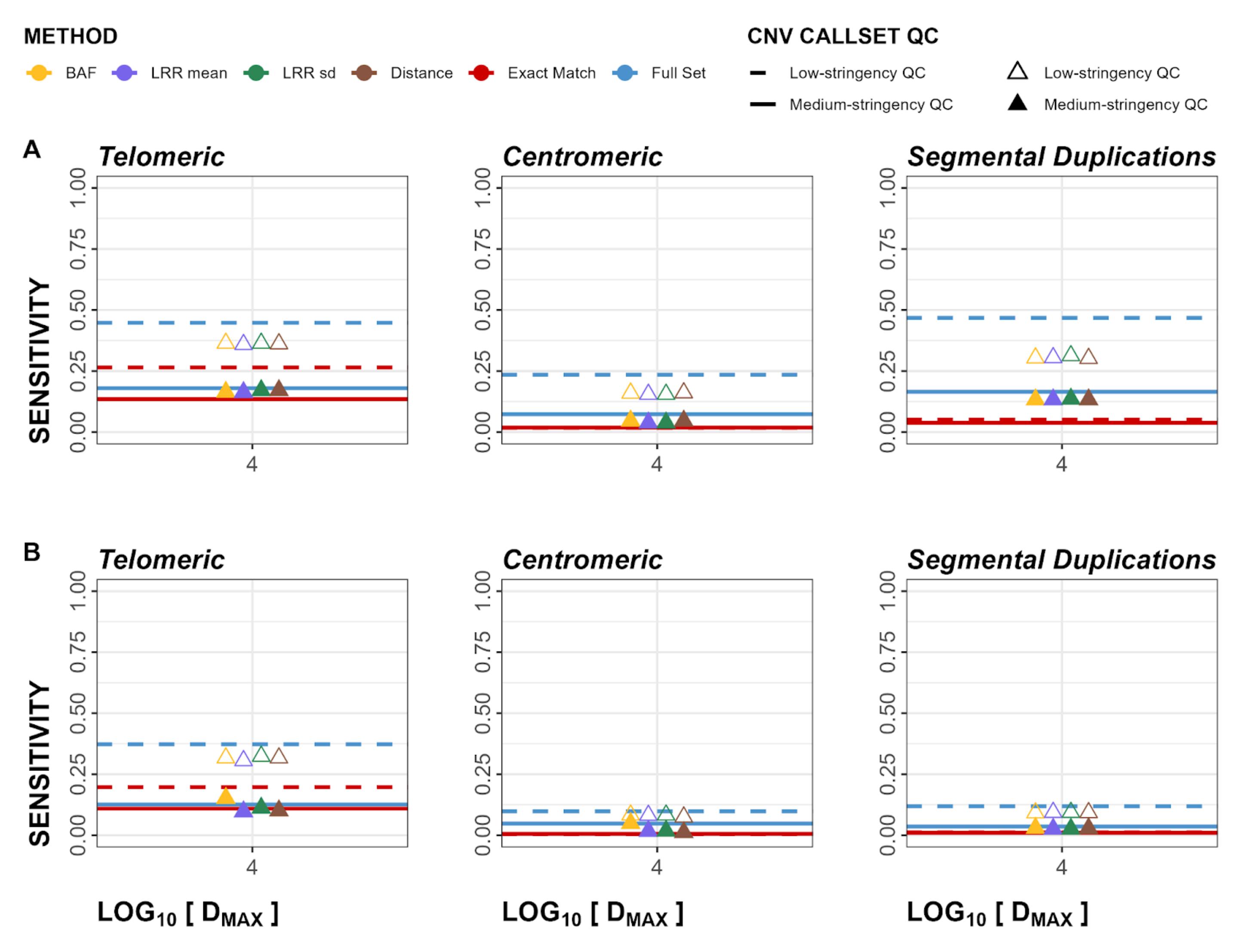


**Figure S63**. Cross-Array Experiment sensitivity regional plots for both deletions and duplications. Panels show telomeric, centromeric, and segmental duplication regions in order, left to right. Empty triangles and dashed lines represent low-stringency QC callsets, whereas solid lines and filled triangles represent medium-stringency QC callsets. Panel A represents calls in Global Screening Array (GSA) validated in Omni Express Exome (OEE) array. Panel  B represents calls in OEE validated in GSA.

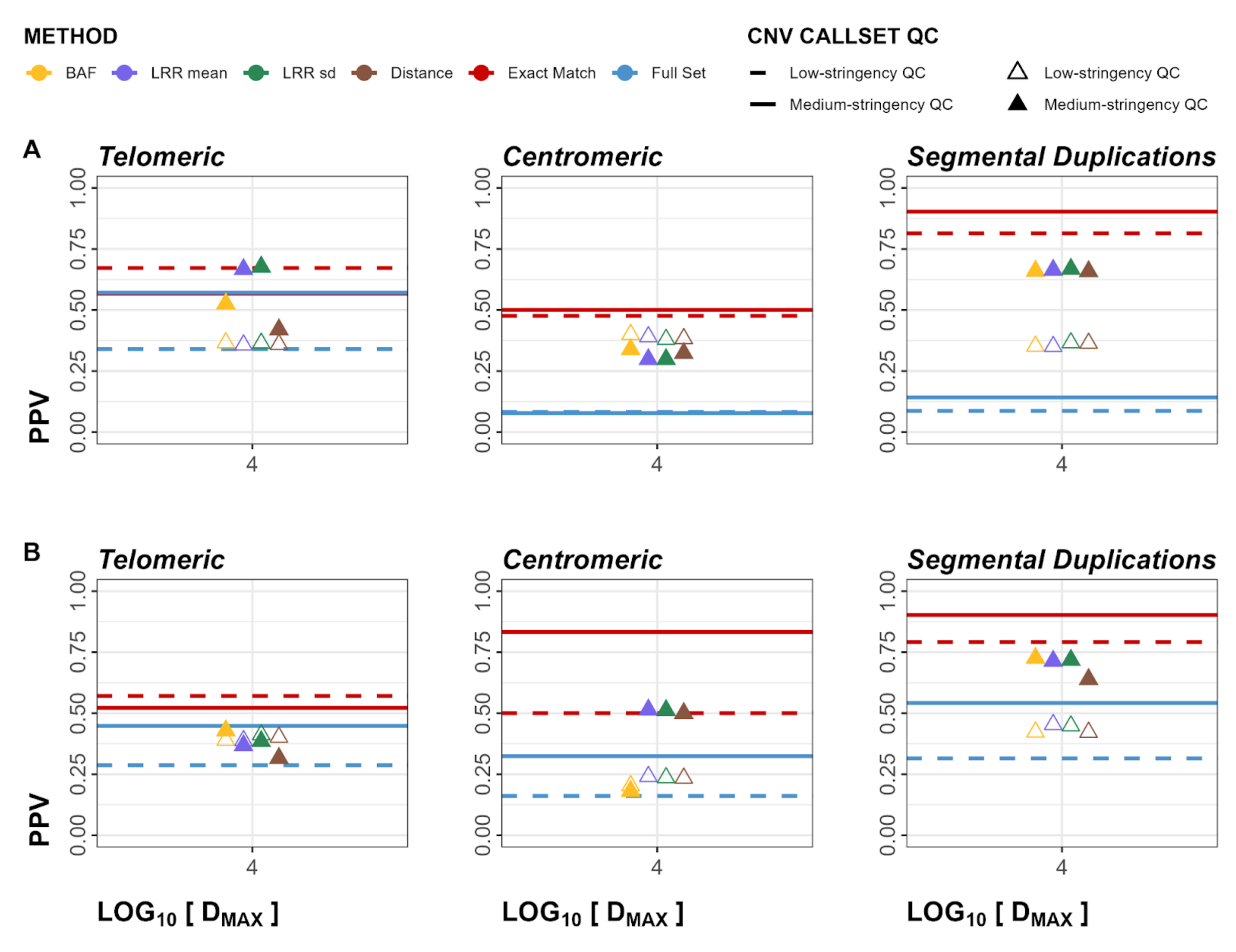


**Figure S64**. Cross-Array Experiment PPV regional plots for both deletions and duplications. Panels show telomeric, centromeric, and segmental duplication regions in order, left to right. Empty triangles and dashed lines represent low-stringency QC callsets, whereas solid lines and filled triangles represent medium-stringency QC callsets. Panel A represents calls in Global Screening Array (GSA) validated in Omni Express Exome (OEE) array. Panel B represents calls in OEE validated in GSA.

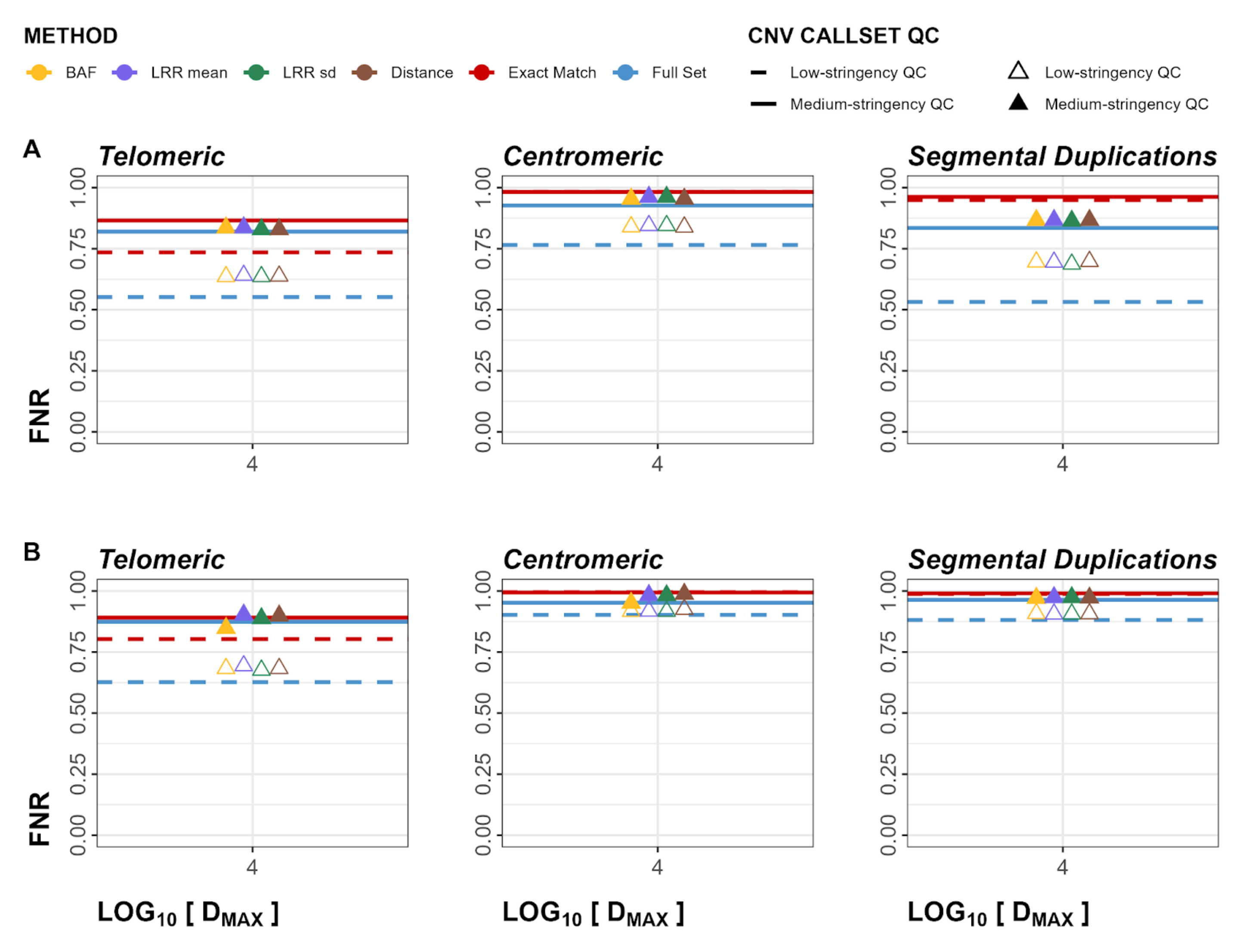


**Figure S65**. Cross-Array Experiment FNR regional plots for both deletions and duplications. Panels show telomeric, centromeric, and segmental duplication regions in order, left to right. Empty triangles and dashed lines represent low-stringency QC callsets, whereas solid lines and filled triangles represent medium-stringency QC callsets. Panel A represents calls in Global Screening Array (GSA) validated in Omni Express Exome (OEE) array. Panel  B represents calls in OEE validated in GSA.

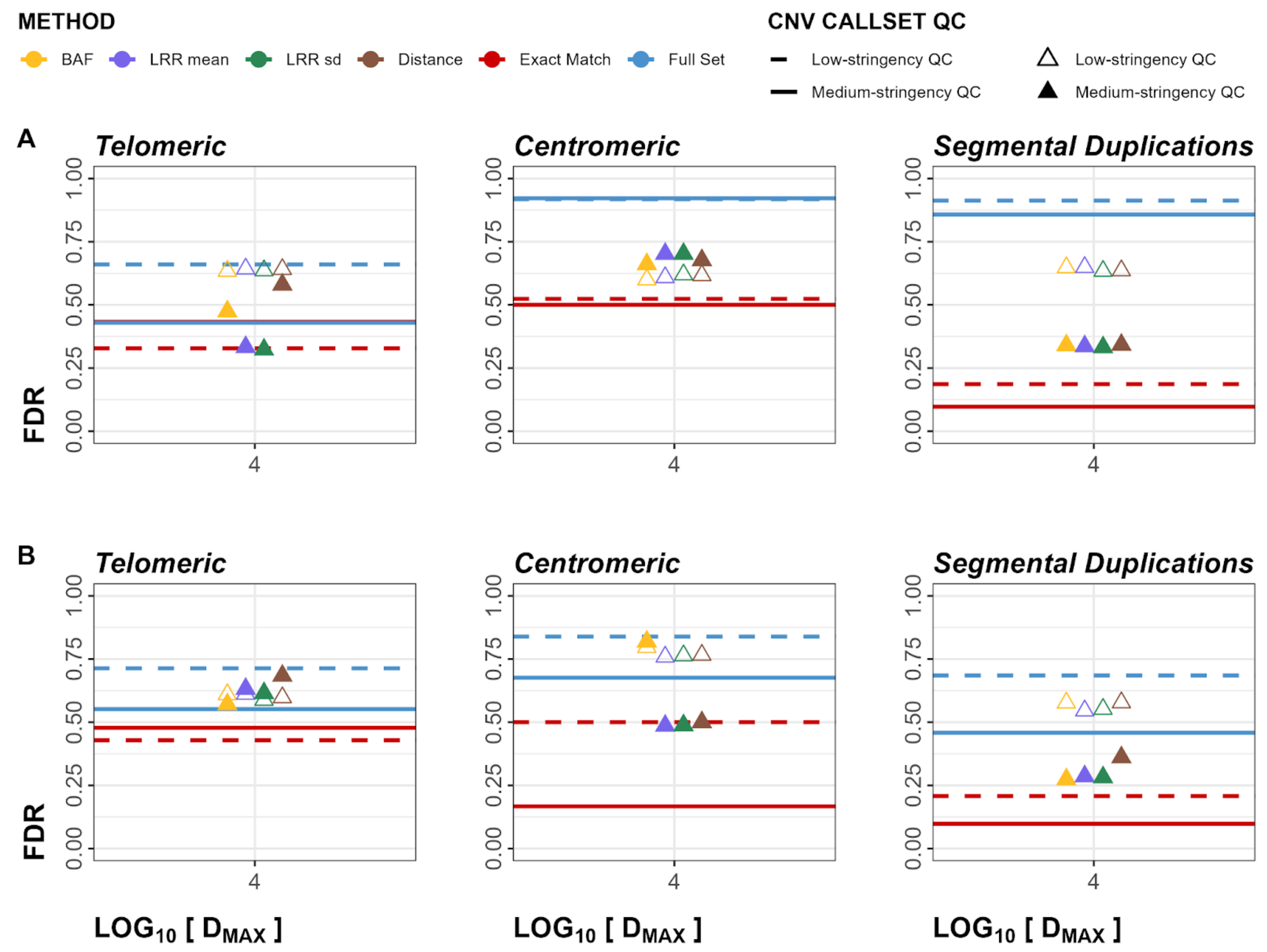


**Figure S66**. Cross-Array Experiment FDR regional plots for both deletions and duplications. Panels show telomeric, centromeric, and segmental duplication regions in order, left to right. Empty triangles and dashed lines represent low-stringency QC callsets, whereas solid lines and filled triangles represent medium-stringency QC callsets. Panel A represents calls in Global Screening Array (GSA) validated in Omni Express Exome (OEE) array. Panel  B represents calls in OEE validated in GSA.

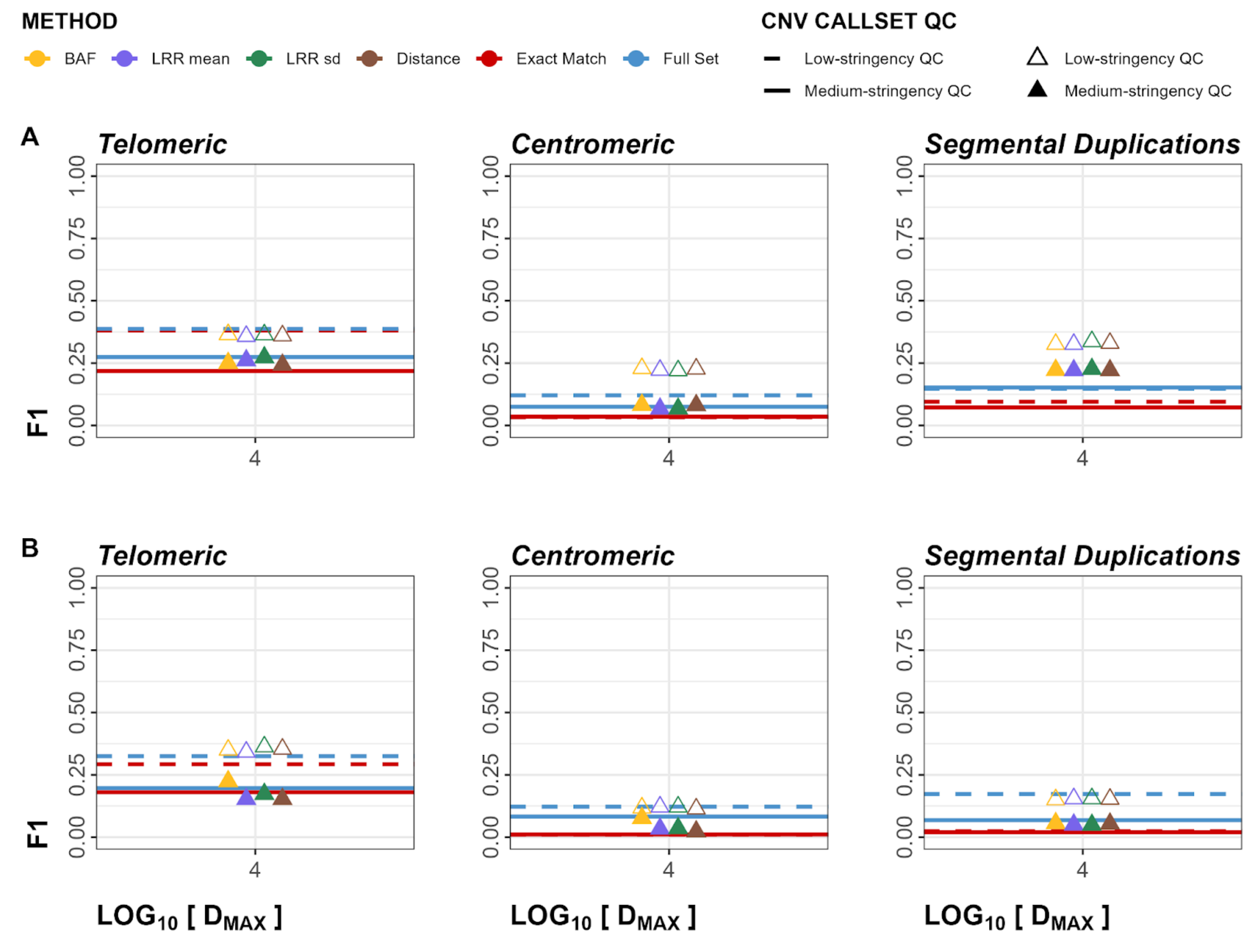


**Figure S67**. Cross-Array Experiment F1 regional plots for both deletions and duplications. Panels show telomeric, centromeric, and segmental duplication regions in order, left to right. Empty triangles and dashed lines represent low-stringency QC callsets, whereas solid lines and filled triangles represent medium-stringency QC callsets. Panel A represents calls in Global Screening Array (GSA) validated in Omni Express Exome (OEE) array. Panel B represents calls in OEE validated in GSA.

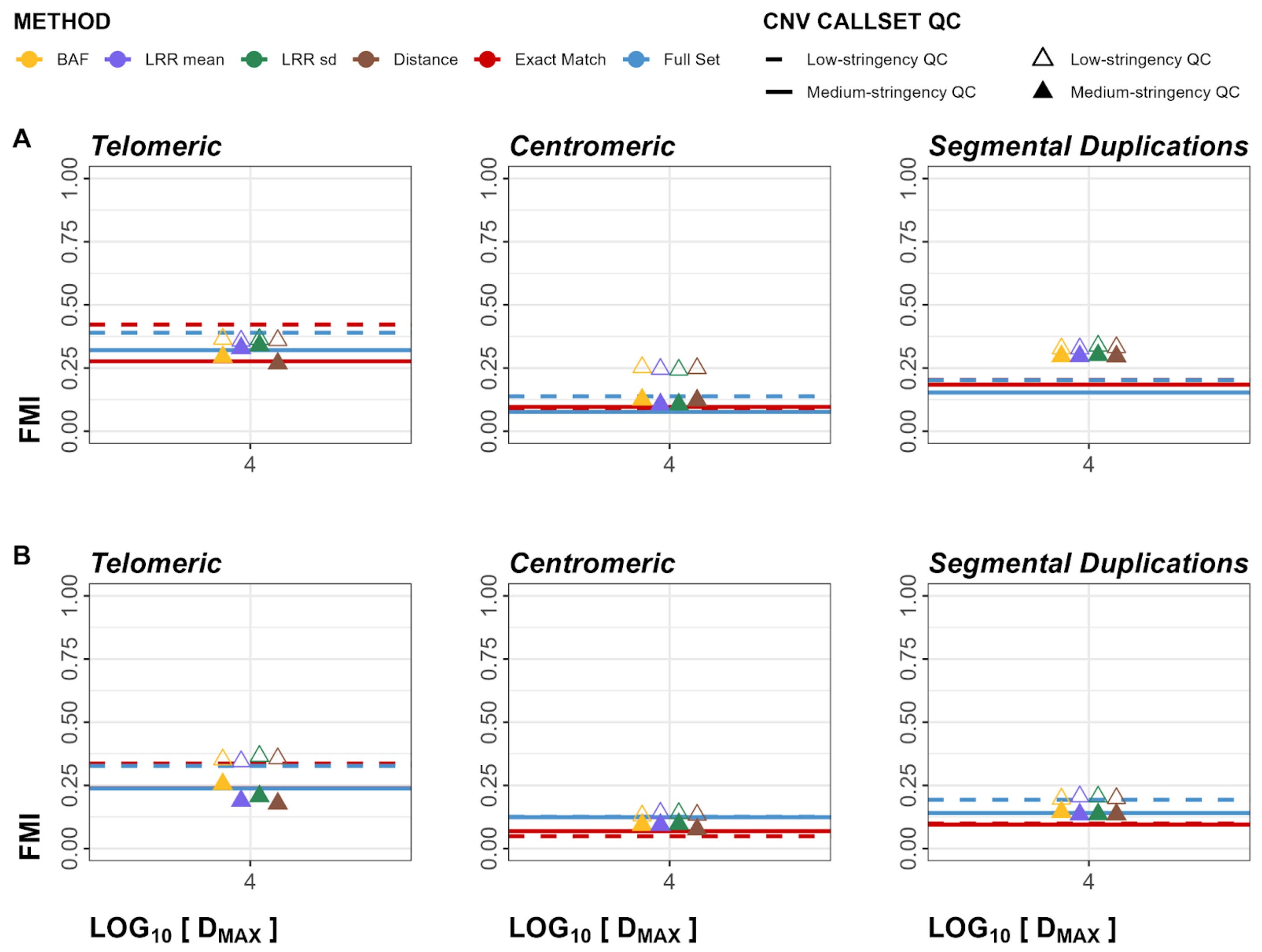


**Figure S68**. Cross-Array Experiment FMI regional plots for both deletions and duplications. Panels show telomeric, centromeric, and segmental duplication regions in order, left to right. Empty triangles and dashed lines represent low-stringency QC callsets, whereas solid lines and filled triangles represent medium-stringency QC callsets. Panel A represents calls in Global Screening Array (GSA) validated in Omni Express Exome (OEE) array. Panel B represents calls in OEE validated in GSA.

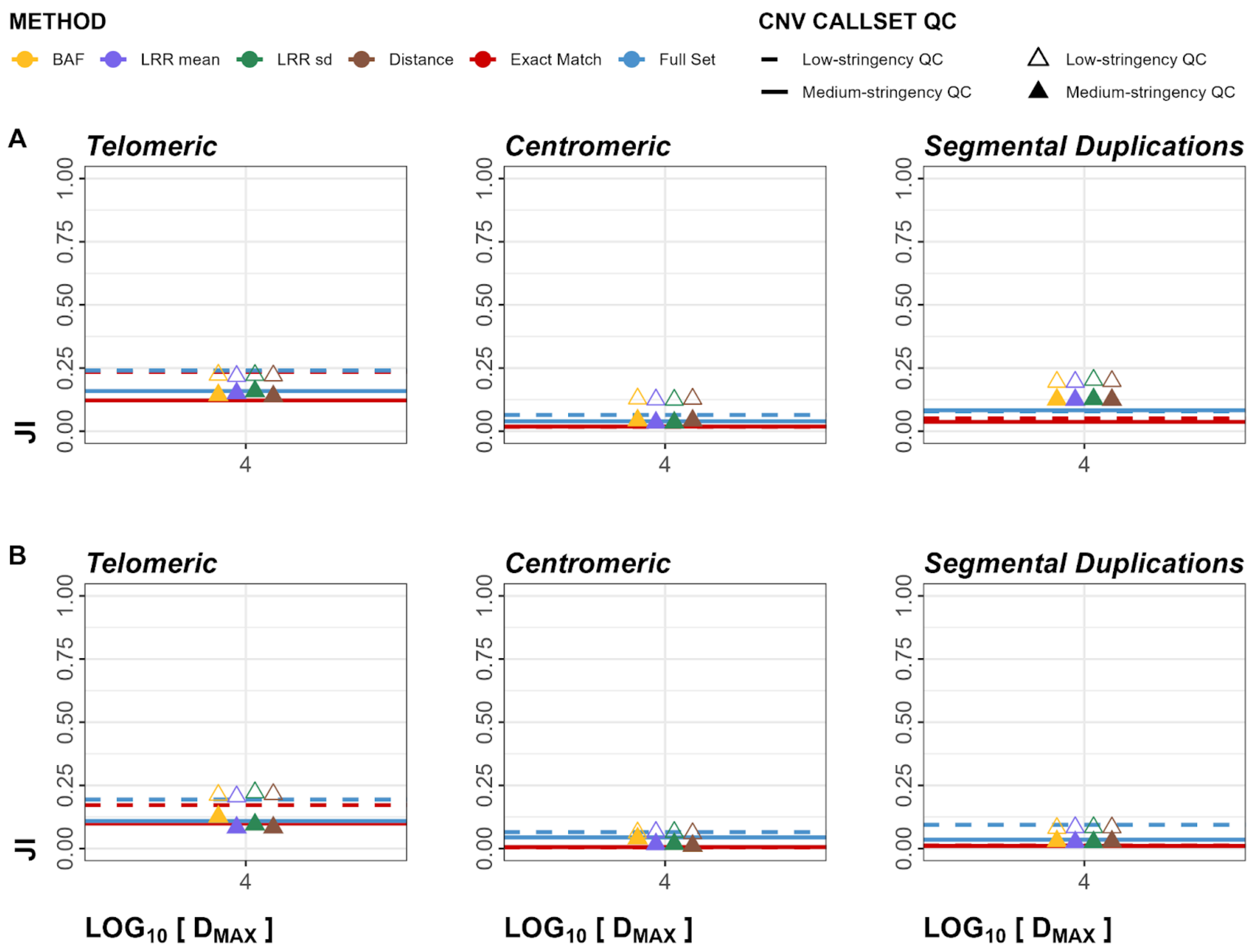


**Figure S69**. Cross-Array Experiment JI regional plots for both deletions and duplications. Panels show telomeric, centromeric, and segmental duplication regions in order, left to right. Empty triangles and dashed lines represent low-stringency QC callsets, whereas solid lines and filled triangles represent medium-stringency QC callsets. Panel A represents calls in Global Screening Array (GSA) validated in Omni Express Exome (OEE) array. Panel  B represents calls in OEE validated in GSA.

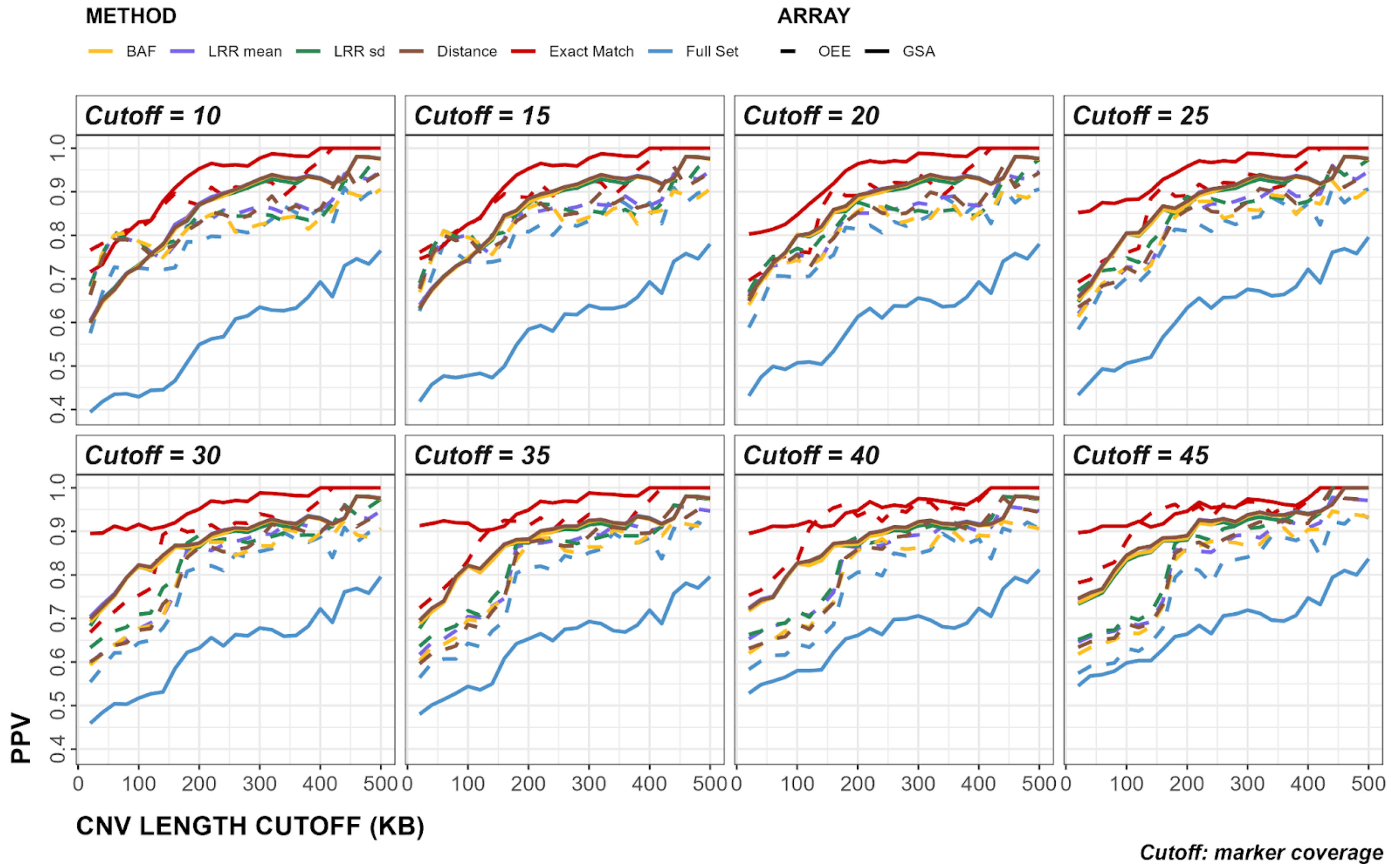


**Figure S70**. Cross-Array Experiment analysis of minimum cutoffs. Y-axes show PPV values, x-axes show minimum CNV length cutoffs, whereas each panel shows minimum marker coverage cutoffs. Solid and dashed lines show the results from GSA and OEE arrays, respectively. GSA: Global Screening Array, OEE: Omni Express Exome array.

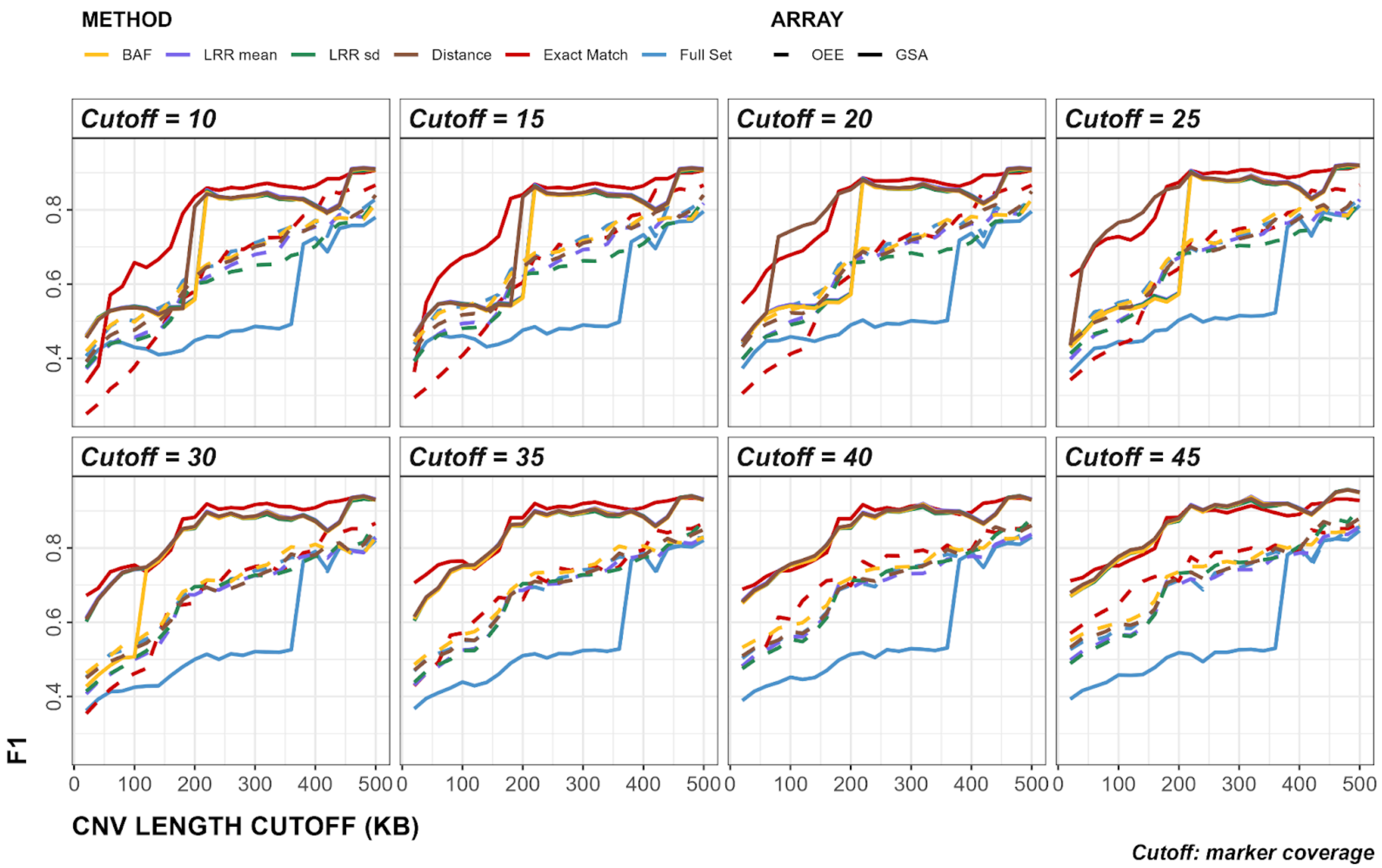


**Figure S71**. Cross-Array Experiment analysis of minimum cutoffs. Y-axes show F1 values, x-axes show minimum CNV length cutoffs, whereas each panel shows minimum marker coverage cutoffs. Solid and dashed lines show the results from GSA and OEE arrays, respectively. GSA: Global Screening Array, OEE: Omni Express Exome array.

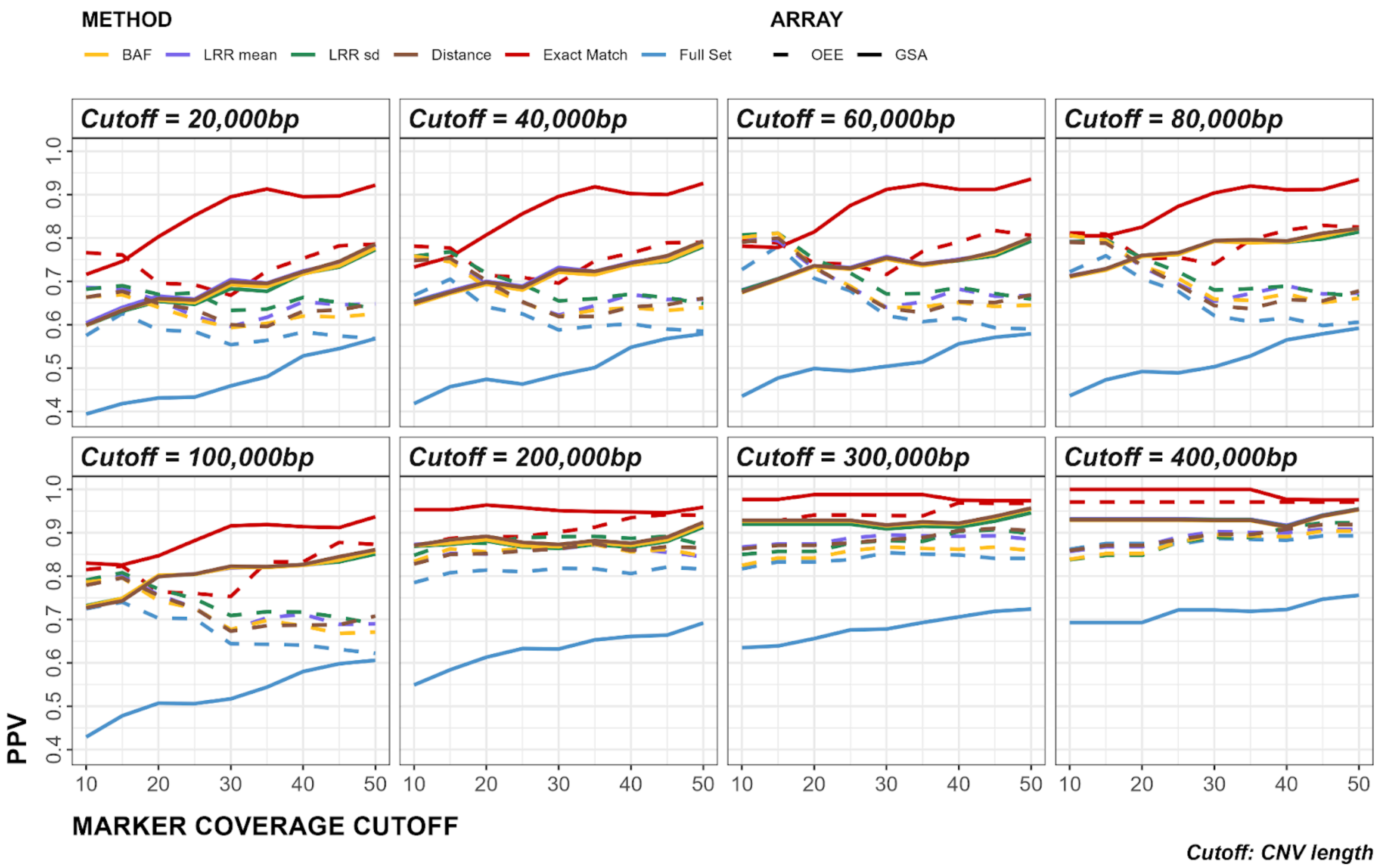


**Figure S72**. Cross-Array Experiment analysis of minimum cutoffs. Y-axes show PPV values, x-axes show minimum marker coverage cutoffs, whereas each panel shows minimum CNV length cutoffs. Solid and dashed lines show the results from GSA and OEE arrays, respectively. GSA: Global Screening Array, OEE: Omni Express Exome array.

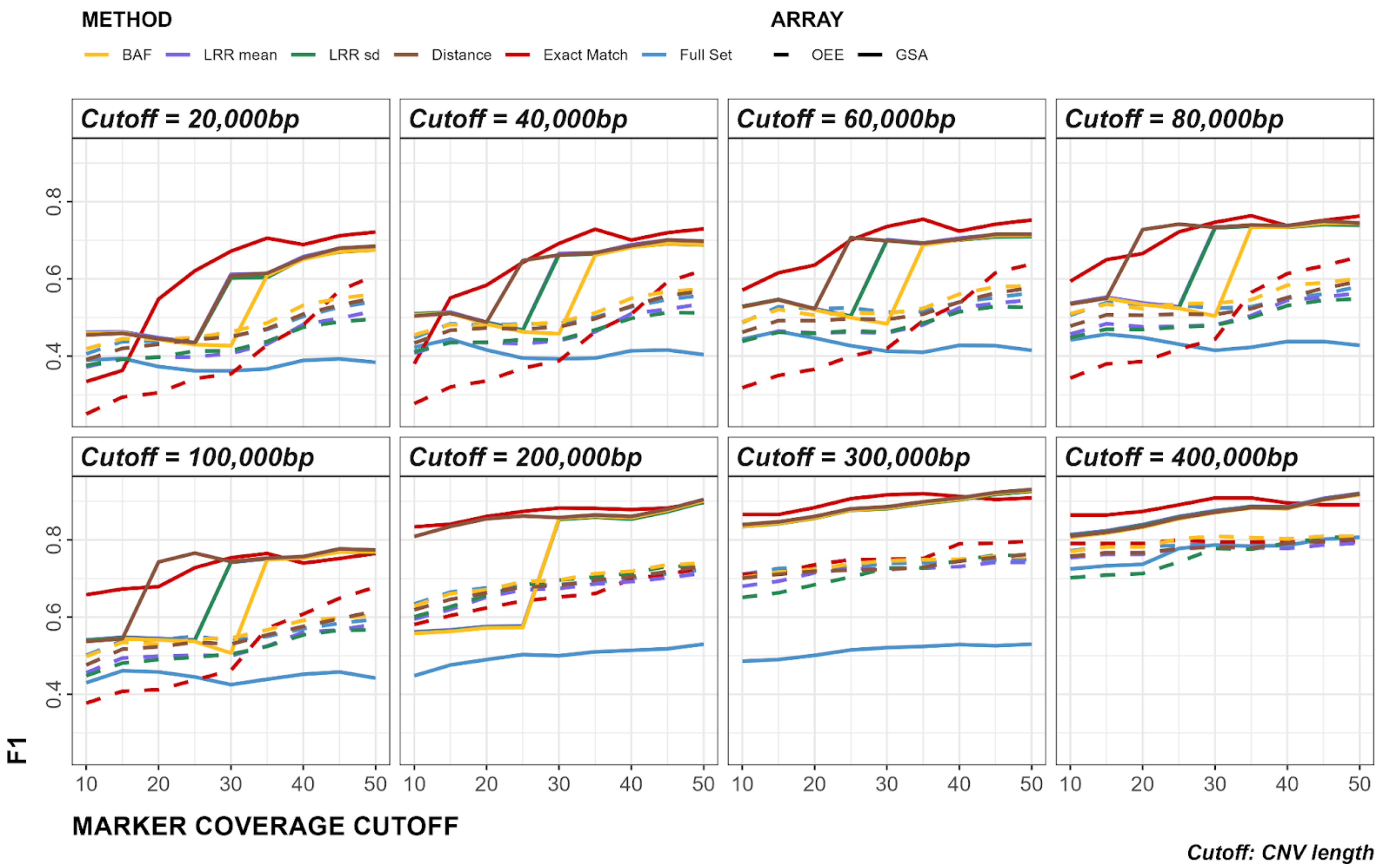


**Figure S73**. Cross-Array Experiment analysis of minimum cutoffs. Y-axes show F1 values, x-axes show minimum marker coverage cutoffs, whereas each panel shows minimum CNV length cutoffs. Solid and dashed lines show the results from GSA and OEE arrays, respectively. GSA: Global Screening Array, OEE: Omni Express Exome array.

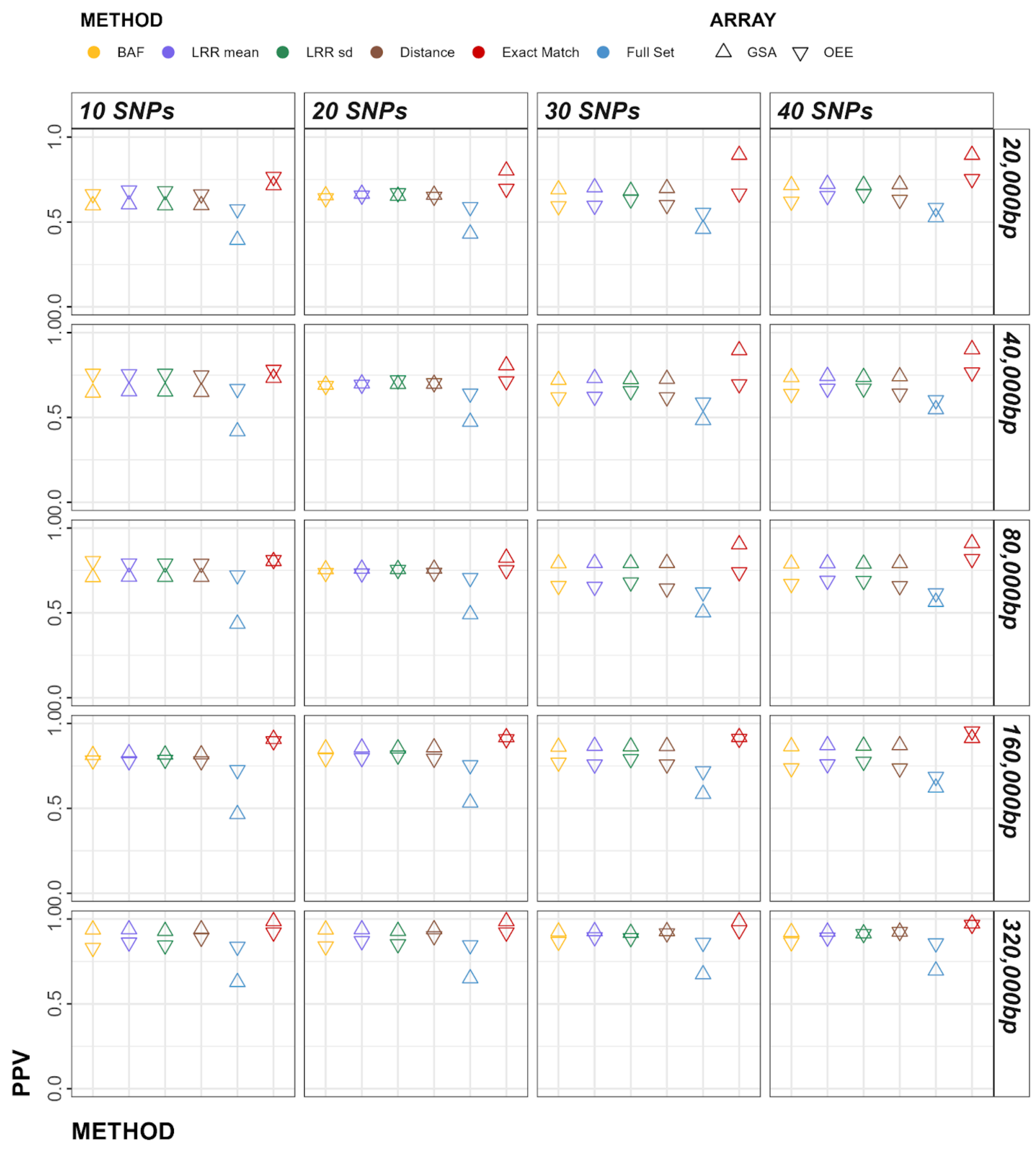


**Figure S74**. Cross-Array Experiment analysis of minimum cutoffs. Y-axes show PPV values, x-axes show individual *Method* parameters (also shown in color, *Full Set* and *Exact Match* included). Panels horizontally grow set *marker coverage cutoffs*, panels vertically grow in set *CNV length cutoffs*. GSA and OEE results are shown as triangles pointing up and down, respectively. GSA: Global Screening Array, OEE: Omni Express Exome array.

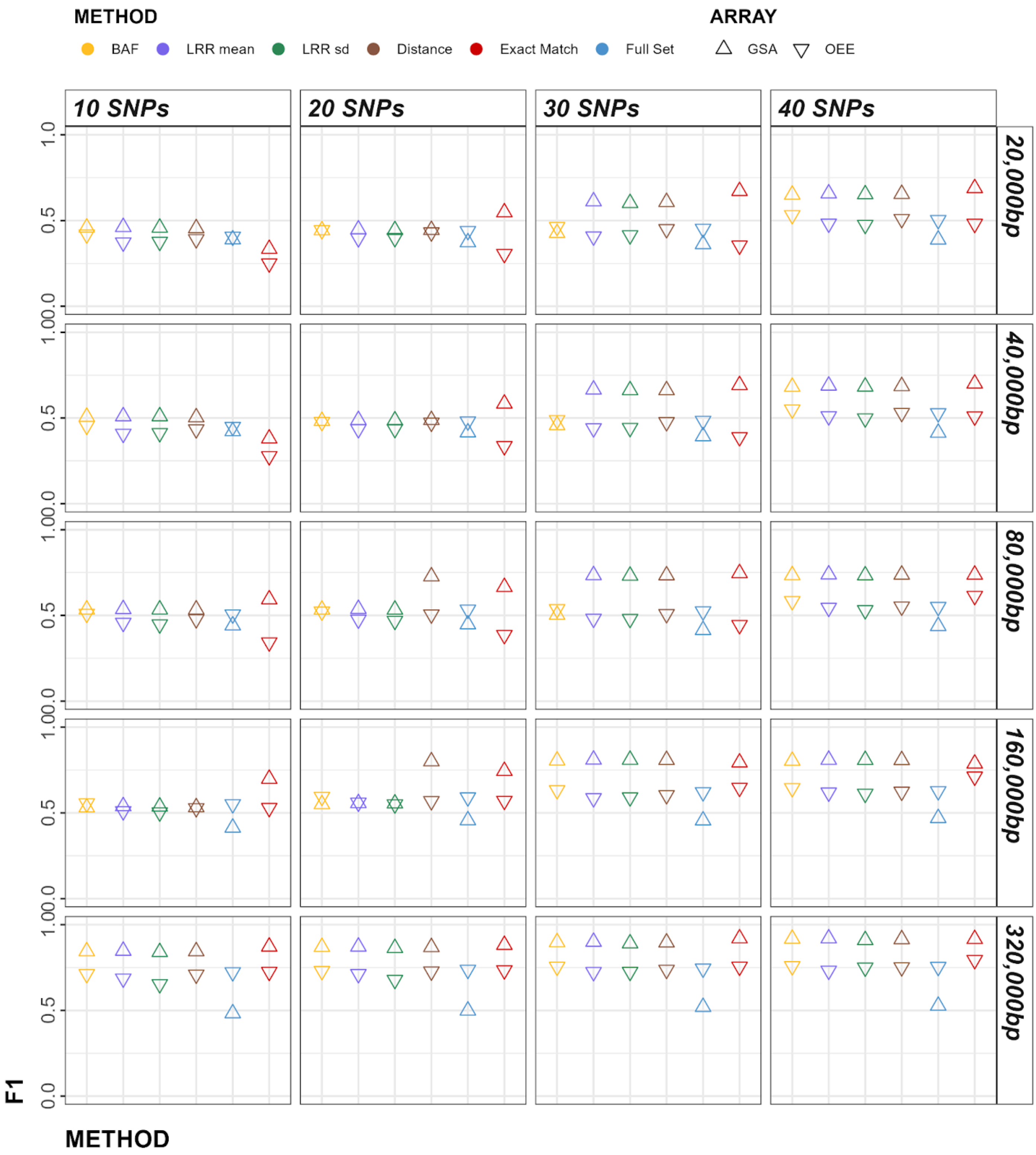


**Figure S75**. Cross-Array Experiment analysis of minimum cutoffs. Y-axes show F1 values, x-axes show individual *Method* parameters (also shown in color, *Full Set* and *Exact Match* included). Panels horizontally grow set *marker coverage cutoffs*, panels vertically grow in set *CNV length cutoffs*. GSA and OEE results are shown as triangles pointing up and down, respectively. GSA: Global Screening Array, OEE: Omni Express Exome array.

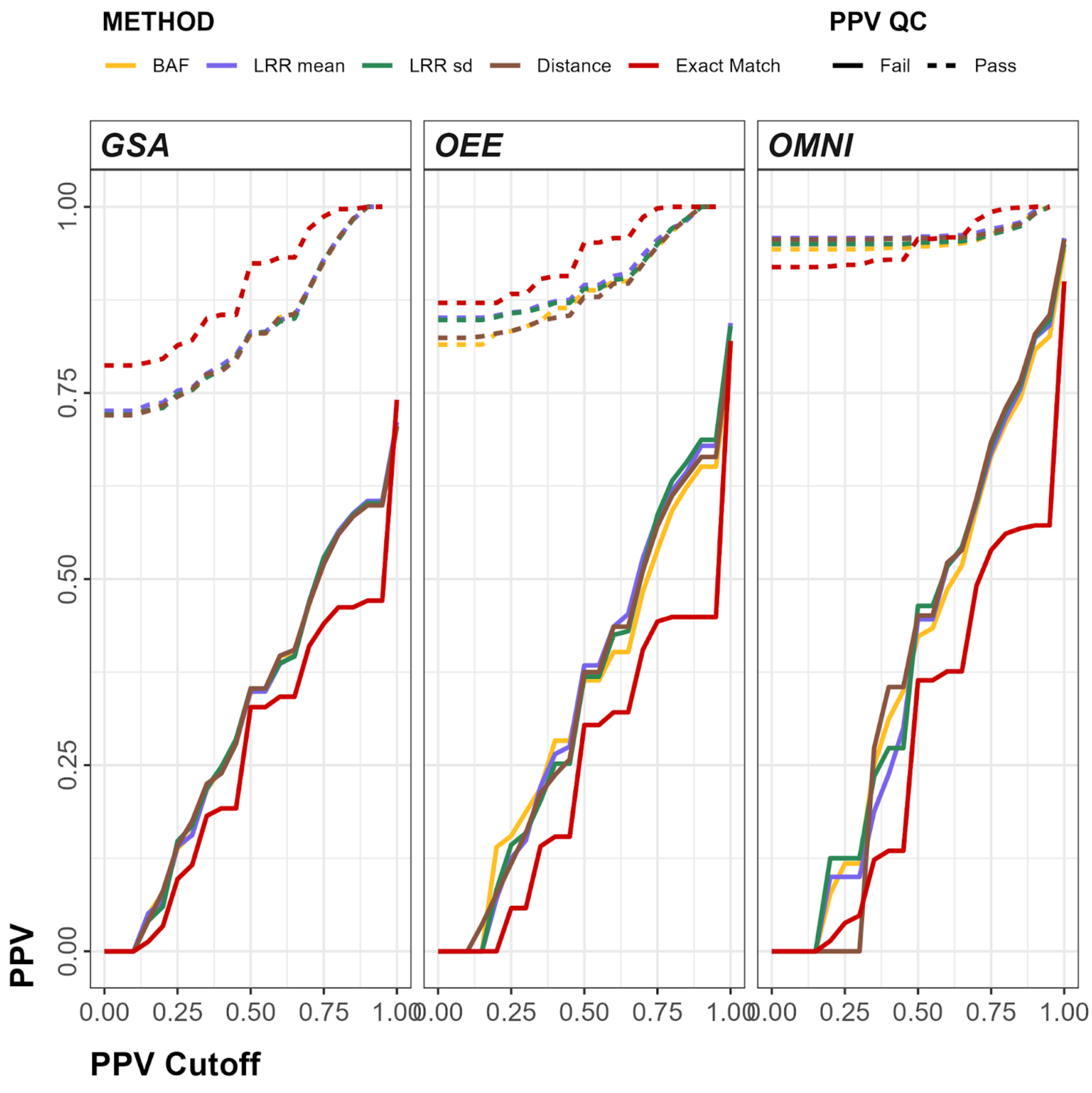


**Figure S76**. Callset PPV as a function of sample-wise PPV QC. The curves indicate callset PPV, after PPV QC. PPV QC passing (solid lines) and failing (dashed lines) along various PPV Cutoffs (ranging from 0 to 1), stratified by *Method* (color) and array (panels). GSA: Global Screening Array, OEE: Omni Express Exome array, OMNI: Omni2.5 array.

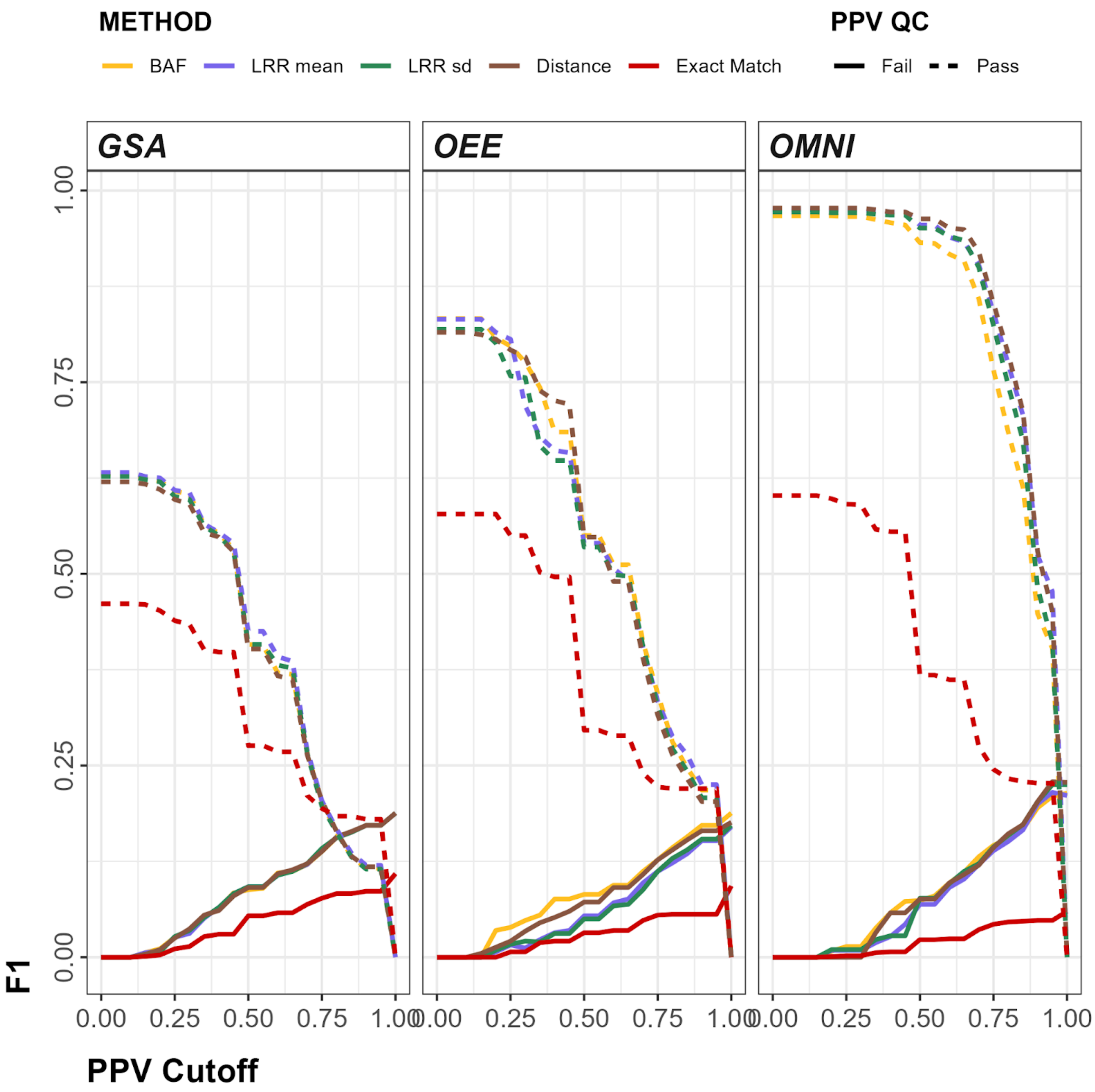


**Figure S77**. Callset F1 as a function of sample-wise PPV QC. The curves indicate callset F1, after PPV QC. PPV QC passing (solid lines) and failing (dashed lines) along various PPV Cutoffs (ranging from 0 to 1), stratified by *Method* (color) and array (panels). GSA: Global Screening Array, OEE: Omni Express Exome array, OMNI: Omni2.5 array.
